# Powerful flowers: Public perception of grassland aesthetics is strongly related to management and biodiversity

**DOI:** 10.1101/2025.02.24.639859

**Authors:** Valentin H. Klaus, Nathan Fox, Franziska J. Richter, Davide Andreatta, Abdesslam Chai-allah

**Author notes:** corresponding author: Valentin H. Klaus, University of Bochum. Shared last authorship (equal contributions) (Davide Andreatta, Abdesslam Chai-allah).

## Abstract

Temperate grasslands provide various cultural ecosystem services that are appreciated in diverse ways. Capturing these diverse appreciations requires different methodological approaches, such as questionnaire surveys and social media analyses. In this study, we combined the potential of both approaches to capture two aspects of what people appreciate in agricultural grasslands, i.e., the aesthetic quality of differently managed plant communities and the objects frequently found in grassland-based social media images. The two complementary approaches showed that people preferred colourful flower- and species-rich grasslands over grass-dominated and fertilised swards. Social media analysis highlighted that people mainly photographed flowers, followed by livestock and/or wildlife, but this depended also on the social media platform used.

In conclusion, people’s appreciation was clearly related to the intensity of grassland management and to the level of biodiversity, with a preference for extensively managed grasslands with diverse flowers and wildlife. Yet, we also found significant differences between (i) conservationists and agricultural professionals in the aesthetic appreciation of the plant communities, and (ii) between common visitors and naturalists in their social media content. Our results suggest that extensive management and ecological restoration can be used to increase cultural grassland ecosystem services by enhancing the richness of forbs, flowers and other attractive wildlife. Thus, targeted management is necessary to maintain and enhance the attractiveness of grassland landscapes and subsequently increase the health benefits that can be associated with these cultural grassland ecosystem services and human-nature contacts.

## 1. Introduction

Cultural ecosystem services (CES) and positive human-nature interactions are of great relevance for human wellbeing and people’s perception of nature and nature conservation, but due to methodological challenges and a lack of data they are widely understudied (Plieninger et al., 2013; Cheng et al., 2019). Insight into the drivers of CES and how these can be enhanced is required to guide land management in all major land use types, especially since intensification and global change threatens CES (Kosanic and Petzold, 2020; Straffelini et al., 2024). Grassland is a major land use type in many regions around the world and provides a wide range of relevant ecosystem services to local populations (Bengtsson et al., 2019). Intensification of grassland management as well as climate change were shown to threaten grassland CES (Allan et al., 2015; Schils et al., 2022; Straffelini et al., 2024). Yet, grasslands were frequently ignored by international strategies (Bardgett et al., 2021), highlighting that grassland CES are not only threatened but also widely overlooked. Thus, we urgently require a better understanding of how grasslands provide CES to be able to develop strategies to maintain and enhance these. Therefore, we must understand what exactly people find interesting and attractive in grasslands, and how this can be supported.

A major CES is the aesthetic value of an ecosystem, which is also closely related to other services such as recreation, tourism, culture and heritage, as well as landscape aesthetics (Bengtsson et al., 2019; Plieninger et al., 2013). Previous work found grasslands in the temperate zone to be hardly perceived to contribute to landscape-scale CES (Ridding et al., 2018; Fox et al., 2022a). However, Schils et al. (2022) found grasslands to contribute more to landscape aesthetics and recreation than forests or arable land. Furthermore, grasslands were found to be important landscape compartments as they enable landscape views (Fox et al., 2022b; Chai-allah et al., accepted). In addition, in a previous study, Richter et al. (2024) found extensive management and use as meadow (versus pasture) to result in higher perceived aesthetic quality, while organic farming did not have an impact on grassland aesthetics. Furthermore, a high diversity of plant species was found to be appreciated by visitors of temperate grasslands (Lindemann-Matthies et al., 2010). Yet, a mechanistic understanding of what people aesthetically prefer in permanent grasslands and how this can be enhanced by land management is still missing.

In grasslands, many aspects can contribute to the aesthetic value: the ecosystem itself, i.e., plant and animal communities and their characteristics, but also land management, such as grazing animals as well as objects such as single trees or old barns (Van Berkel and Verburg, 2014). Therefore, assessing the aesthetic value of grasslands requires a comprehensive view on the many characteristics of this ecosystem, and likely also different methodological approaches that allow insight into the aspects of aesthetic perception.

The most frequently employed approaches to understanding ecosystem aesthetics comprise questionnaire surveys that interview people either in front of the ecosystem in question (e.g., Lindemann-Matthies et al., 2010) or with images as visual stimuli (e.g., Fischer et al., 2018). The extraction of specific characteristics of images linked to respondents’ appreciation is typically carried out by hand as well as questionnaires to collect information about the actual appreciation (Richter et al., 2021; Wang et al., 2024). Examples of this include visually estimating flower cover and other structural properties in the field. However, this method is time-consuming and challenging to scale and standardise. Automatic spectral analysis of the images may be able to help get around this restriction since spectral features of images were shown to be associated with the composition and structure of plant communities (Torresani et al., 2024). However, the relationship between the spectral characteristics of grassland images and appreciation remains largely unexplored in the scientific literature.

More recently, studies used the vast crowdsourced data uploaded to social media and citizen science platforms (e.g., Havinga et al., 2020; Chai-allah et al., 2023; Ghermandi et al., 2023). Such approaches benefit from the large amount of data that is being uploaded, such as spatially explicit images, and employ advanced data science tools for content analysis (Fox et al., 2021; Schirpke et al., 2023). Combining both questionnaire surveys to indicate ‘stated-preferences’ and crowdsourced images analyses to indicate ‘revealed preference’ has the potential to considerably advance our understanding of grassland aesthetics (Cheng et al., 2019).

In this work, we explore what grassland characteristics are appreciated by people and how these are related to management measures (Figure 1). Here, we combine a questionnaire survey, an automatic spectral image analysis, and a crowdsourced image analysis to understand what people find attractive and of interest in agriculturally managed grasslands in Switzerland. Further, we conducted (i) an in-depth analysis of a structured questionnaire survey with people’s ratings of 92 grassland images taken along a land-use intensity gradient (Richter et al., 2024) and (ii) a social media-based image analysis extracting information on relevant objects that have been photographed in grasslands and uploaded to two crowdsourced data sources. Thus, we studied two presumably simultaneously relevant aspects of the aesthetic value of grasslands, i.e., people’s perception of the grassland plant community, as affected by different anthropogenic and environmental drivers, and the objects people photograph and share online. Some of the results of this questionnaire were used by Richter et al. (2024) to coarsely characterise the aesthetic value as an aspect of grassland multifunctionality. This analysis, however, goes much deeper and overcomes the purely descriptive approach of the previous study by conducting a mechanistic assessment of what drives people’s aesthetic rating of the plant communities.

**Figure 1.**
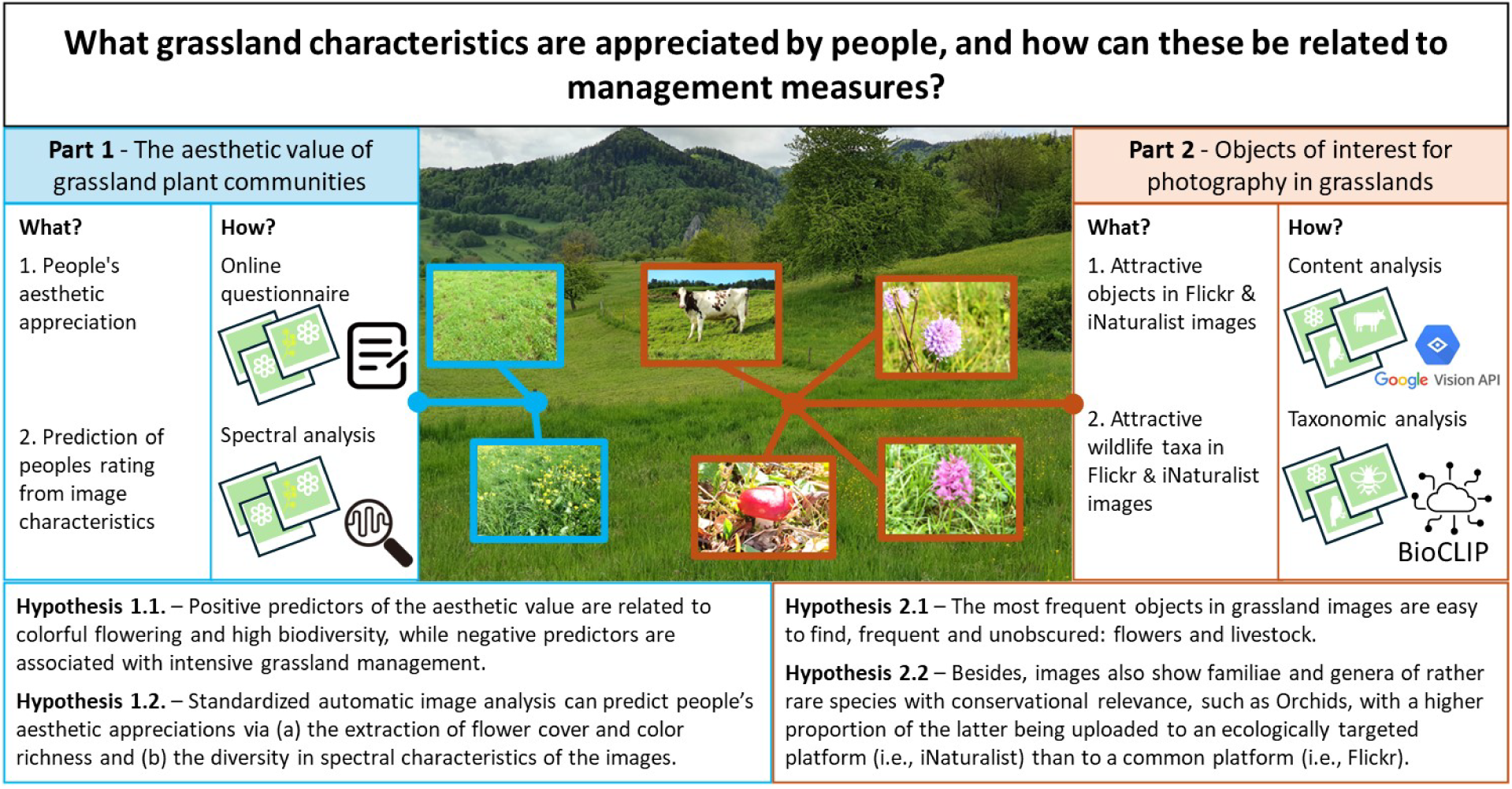
Conceptual figure highlighting the two approaches used in this study, the respective methods and hypotheses. While part 1 uses images (photographs) taken in a standardised way, the images in part 2 were uploaded by social media (i.e., Flickr and iNaturalist) users in the study region (Figure 2).

We defined two sets of hypotheses, as shown in Figure 1.

## 2. Materials and methods

Because of the multi-faceted and complex nature of CES, this work combines two complementary approaches assessing two different but presumable simultaneously relevant aspects of the aesthetic attractiveness of grasslands in the northwest of Switzerland (section 2.1). First, we assess the attractiveness of the vegetation matrix, i. e., the grassland plant community, via a photo-based structured survey on people’s aesthetic perceptions (section 2.2). Second, we assess objects of interest in grasslands with an analysis of images uploaded to two crowdsourced platforms (section 2.3).

### 2.1. Study system

We focus our work on the northwest of Switzerland (Figure 2), a grassland-dominated region with a landscape-scale land-use composition comparable to many other temperate European regions. Grassland dominates the two biogeographic subregions studied, i.e., the intensively used plateau, in which grassland is combined especially with arable land, and the mountain range “Swiss Jura”, composed of limestone ridges in which grassland and forest occur. While the plateau stretches between 400 to 800 m a.s.l., the peaks of the mountainous part reach up to 1200 to 1700 m a.s.l. We chose this region because its grassland types and their management are typical for central Europe. These grassland types are (i) productive sown leys (temporary grassland) that can be included in crop rotations, (ii) productive and intensively managed permanent grasslands and (iii) semi-natural grasslands, which are usually managed according to agri-environmental schemes that considerably restrict management intensity, especially fertilisation (Klaus et al., 2023; Lüscher et al., 2019). No alpine areas and no natural grasslands occur in the study region.

**Figure 2.**
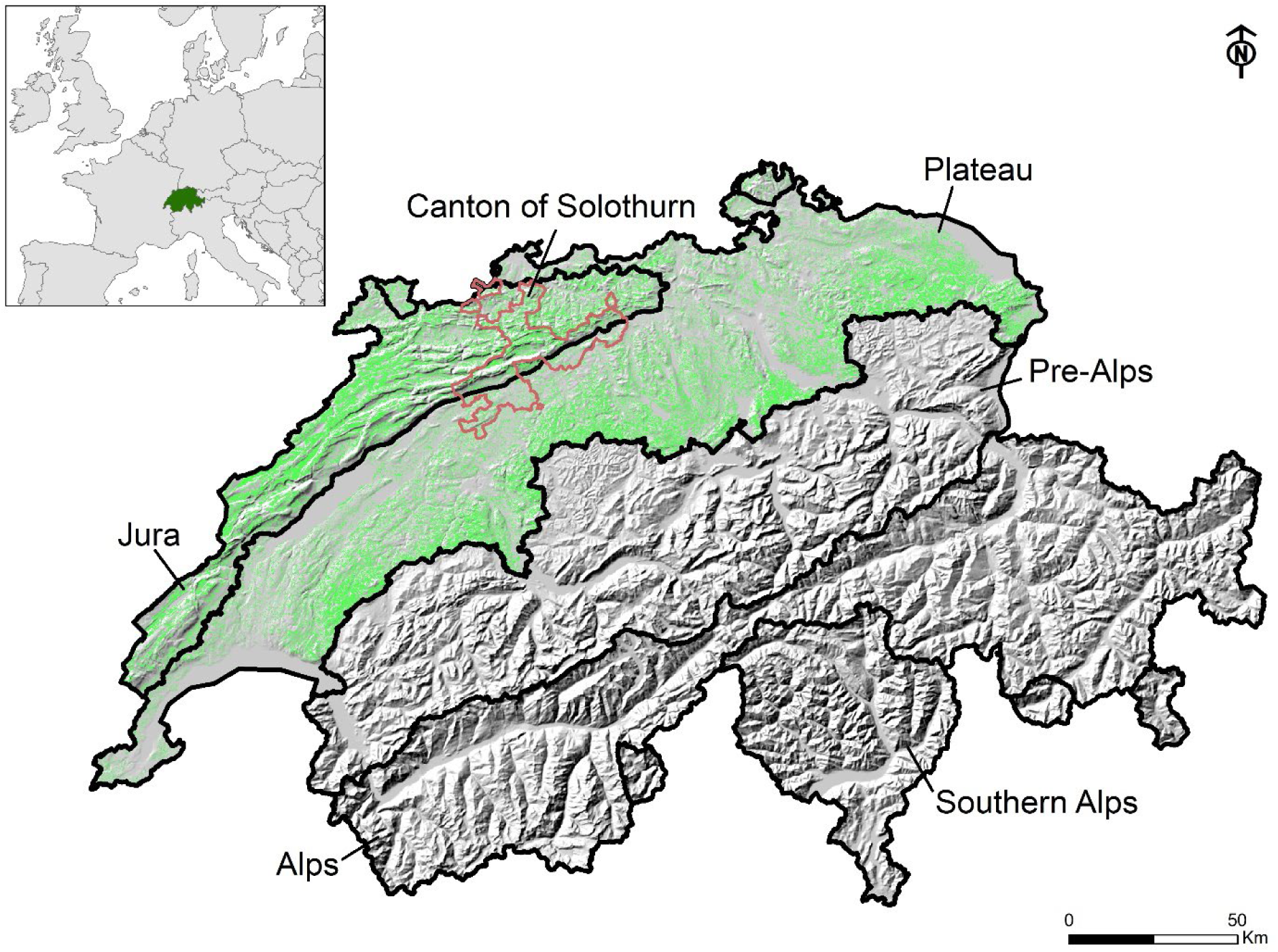
Study region composed of the two biogeographic subregions “Jura” and “Plateau” in northwestern Switzerland. In the two areas the distribution of grasslands is shown in green according to Huber et al. (2022). Shaded relief of Switzerland taken from swisstopo (2020). While the standardised images for part 1 were taken in the canton of Solothurn, indicated by a brown frame, images taken from social media platforms (part 2) originate from the whole study region.

### 2.2. Part 1 - structured survey on people’s aesthetic perceptions

#### 2.2.1. Questionnaire survey

To conduct a questionnaire survey on people’s aesthetic appreciation, we took 92 images of permanent grasslands in the canton of Solothurn, which is located right in the middle of the overall study area and covers both the Jura and the Plateau subregions (Figure 2). Images were taken from mid-May to mid-June 2021. Respective permanent grassland plots were selected along a gradient in management intensity based on the grassland typology provided by official agricultural statistics. The main four grassland types in this typology are meadows (mainly mown) and pastures (mainly grazed), which can be intensively managed (fertilisation allowed) or extensively managed (fertilisation prohibited; management according to the Swiss agri-environmental scheme “extensively managed pastures” and “extensively managed meadows”). For more details on the typology and the management of the grassland types see Klaus et al. (2023).

All images show a close-up of the plant community in an area of approximately 2 m × 2 m from the height of one meter above ground level during peak flowering in late spring, without any objects, such as livestock or other animals, or landscape views, in a resolution of 4032 x 3024 pixels. Most images were taken before harvest, but in twelve cases, grasslands were already grazed during the time of sampling. Eleven out of these twelve plots were pastures, which can generally be grazed early in the season (as from end of March). The resulting “signs of defoliation” were considered by using dummy coded variables in the statistical analysis (see section 2.2.2.). In addition to the permanent grasslands, two images of sown (temporary) grasslands at comparable phenological stage were added to the final set to also consider this grassland type, resulting in a total of 92 images included in the questionnaire.

A German-language online questionnaire was online from the 11^th^ of January to the 7^th^ of March 2022, asking people for their personal perception of the aesthetic quality of grassland plant communities shown on 92 images. Aesthetic quality was available as a 5-point Likert scale from 0 = *unattractive* to 5 = *attractive*. The survey was set up with QuestionPro (QuestionPro Inc, Austin, TX, United States), with ten of the 92 images randomly chosen per questionnaire. The link to the survey was distributed via different pathways, i.e., (i) via e-mail to professional stakeholders in agriculture, nature conservation, tourism, and/or botanical gardens in Switzerland, (ii) via German-language Twitter (now X) posts and mailing lists, and (iii) further private and professional networks. The survey text and all pictures are available in supplementary material 1. Since the survey was completely anonymous, it did not require an approval of the institution’s ethical board.

The average rating per plot was computed and used as the indicator for people’s aesthetic appreciation of the shown grassland plant community. We further asked about the professional work of the respondents, offering the categories “agriculture”, “ecology/nature conservation”, “both professional areas”, “neither of both, other professions or retired”. We therefore additionally computed the average aesthetic value for all images for three separate groups, omitting the group responding “both agriculture and ecology/nature conservation” because we expected them to overlap with the other categories (and due to very few responses in the category, see below). A linear mixed effects model was used to test for significant differences in aesthetic ratings among the professional groups, with plot as random factor.

#### 2.2.2. Explanatory variables of plant communities

To understand the drivers of the aesthetic appreciation of people, we recorded information on plant community composition and the agricultural management of the grasslands, and we extracted visible and spectral characteristics of the images. Together with taking the 92 images in permanent grasslands, we at the same time conducted vegetation records in two 2 m × 2 m areas in each site. We visually assessed all vascular plant species and their ground cover as well as the total ground cover of cryptogams. The two records were near the area shown on the image but were not necessarily identical due to logistic reasons. However, since the vegetation of the grasslands was widely homogeneous and any accidentally occurring clusters of weeds or other atypical vegetation were avoided, we consider the vegetation records representative for the plant communities shown on the images. Based on the vegetation records, we calculated three plant diversity metrics (i.e., number of vascular plant species, evenness and Shannon diversity) as well as the cover sums of all grass and all forb species, with the latter including legume species. Furthermore, as an indicator of the agricultural value of a plant community, we calculated abundance-weighted forage quality indicator values according to Briemle et al. (2002) based on plant indicator values. After the vegetation records, we sampled the above-ground plant biomass as a measure of productivity (i.e., peak standing crop) and measured the height of the vegetation by placing a laminated A4 sheet on top of the vegetation, parallel to the ground and then recording the height of the sheet with a ruler. Height measurements represent a mean of four replicates per plot taken in each corner of each vegetation record. Information on topographical features, i.e., slope and elevation of each plot, were extracted from a Digital Elevation Model (25-m grid resolution, European Union 2018) using QGIS (version 3.22). To gather detailed information on the specific management of the grasslands, we collected management data for 2021 and 2022 from farmers. We recorded the fertilisation intensity (sum of available nitrogen in organic and mineral fertiliser per ha per year applied as fertilizer – not including defecation of animals grazing on the area), the mowing frequency (number cuts per year), and the grazing intensity (livestock unit days per ha per year) following Blüthgen et al. (2012). Intensity measures were averaged for the two years to provide a stable mean. For further details on the images, the questionnaire and the vegetation records see Richter et al. (2024).

To gather information on the actual plant community visible in the photographs, one person coded information from all images, i.e., flower cover (coded as % and quasi-metric categories of *1* = 0-1%, *2* = 1-5%, *3* = 5-15%, *4* = more than 15% of image filled with flowers), excluding green and grassy flowers, and flower colour richness. We defined the colour classes as follows: yellow flowers (such as *Ranunculus* spp., *Rhinantus* spp., *Crepis biennis*, and *Taraxacum* sp.), white flowers (such as *Leucanthemum ircutianum*, *Capsella bursa-pastoris*, *Daucus carota*, *Trifolium repens*, *Galium mollugo*, and *Bellis perennis*), red flowers (such as *Trifolium pratense* and *Onobrychis viciifolia*), violet and similarly coloured flowers (e.g., *Knautia arvensis, Scabiosa columbaria,* and *Salvia pratensis*), and blue flowers (e.g., *Ajiuga repens*, and *Veronica* spp.). In addition, other image properties were binary coded (i.e., signs of defoliation (Yes/No), bare soil visible (Yes/No), litter layer visible (Yes/No), agricultural weeds prominently visible (Yes/No), and vegetation structure (Light/Dense)). Agricultural weeds found on the images were *Rumex obtusifolius*, *Colchicum autumnale*, *Senecio* spp., *Rhinanthus* spp., *Carlina* spp., *Cirsium* spp., *Anthriscus sylvestris*, and *Heracleum sphondylium*. To relate all these characteristics of the plant communities and associated measures to the aesthetic ratings, we used Spearman correlations for metric and quasi-metric data as well as ANOVAs for categorical data. These analyses were run for all responses together and separately for the professional groups. Diagnostic plots were used to ensure model assumptions were met.

To get insights into the most common taxa found in the different grasslands in the study area, and to compare this with the Results related to hypothesis 2.2, we in a first step used the “classification” function from the package “taxsize” (v. 0.9.102, Chamberlain and Szocs, 2013) to extract the higher-level taxonomy from the species present in the vegetation records. We then used this information to calculate the cover of a) different plant families and b) different plant genera per grassland (using the mean covers of the two plots per grassland), as well as the number of grasslands in which a) different plant families and b) different plant genera were present in the vegetation records. These results were visualized in the form of barplots using ggplot2 (v. 3.5.1, Wickham, 2016). This analysis was carried out in R (v. 4.4.2, R Core Team, 2024).

#### 2.2.3. Automatic extraction of flower and spectral diversity metrics

In addition to estimating flower cover and flower colour richness visually, we automatically extracted flower cover and flower colour richness through supervised image classificationto establish uniform methods for quantifying floral attributes that relate to people’s aesthetic appreciations. First, we defined the pixel classes as follows: green vegetation and soil, yellow flowers, white flowers, red flowers, violet flowers, and blue flowers. Then, we developed a labelled dataset by randomly selecting a 1000 pixels × 1000 pixels image patch (corresponding to around 50 cm × 50 cm at the ground level) in each image, plotting in RGB colours using the “plotRGB’’ function of the “raster” package (Hijmans, 2023), labelling around 50 pixels per patch by clicking on the image to retrieve the X and Y coordinates using the “locator” function of the “graphics’’ package, and assigning to each pixel the class to which it belongs. Since labelled pixel class frequency was very unbalanced with only a few violet and blue flower pixels available, additional image patches where violet and blue flowers occurred were identified in the RGB images, and pixels labelled. Next, we removed pixels too close to each other from the dataset, i.e., pixels falling in the same pixel after downscaling the images with an aggregation factor equal to 10. The resulting dataset consisted of 4832 labelled pixels, with 117 to 3315 pixels per class. The reflectance values in the red, green and blue bands of the images in labelled pixels were extracted to obtain a dataset reporting image ID, pixel class, XY coordinates, and reflectance values for each labelled pixel. Afterwards the dataset was split into training and validation by randomly assigning 66% of images for training, and 33% of images for validation. The training dataset was used to develop a random forest classifier using the “randomForest’’ function of the “randomForest” package (Liaw and Wiener, 2002) with default parameters. Classifier performance was assessed on the validation dataset by computing the overall accuracy, the Kappa accuracy, and the mean F1 score of the six classes (Congalton and Green, 2009). The compiled classifier was used to classify all images except for two, which were discarded since they had different dimensions. While deriving the flower cover from the classified image through image statistics was straightforward, defining the flower colour richness required the definition of the percentage threshold above which a flower class is to be considered as present in the image. We tested 2000 possible thresholds from 0% to 1% and chose the threshold resulting in the highest correlation between manually and automatically estimated flower colour richness.

Since we hypothesised that people’s appreciation could not only be affected by flower cover and flower colour richness, but also by the evenness of the distribution of the pixel classes, with higher diversity associated with higher appreciation, we applied two diversity metrics widely used in grassland ecology studies, Shannon and Simpson, to the frequency distribution of the pixel classes in each image. The six-pixel classes were considered as species, and pixel counts as species abundances. Diversity metrics were computed using the “diversity” function of the “vegan” package (Oksanen et al., 2022). The relationships between appreciation and flower cover and between appreciation and diversity metrics were assessed by fitting second-order polynomial regression models, since the relationships were not linear. Finally, since we hypothesised images with more differences in the reflectance spectra between the pixels (e.g., less homogeneously green) to be associated with higher people’s aesthetic rating, we assessed the diversity in spectral characteristics of all 92 images of permanent grasslands. The diversity in spectral characteristics of images is not unequivocally defined, but suitable metrics can be found in the field of remote sensing of terrestrial plant biodiversity (Wang and Gamon, 2019). The rationale behind spectral diversity metrics is that the increased spectral variability reflects an increased variety of habitats, species, and organs, since different habitat, species and organs respond in their own way to incoming solar radiation according to their pigment, water, and biochemical content, as well as leaf and canopy structure. The diversity metrics we applied here were: whole image standard deviation, moving window standard deviation, multidimensional Rao index. Since shadowed pixels may lead to noise rather than enhancing the information content at very-high spatial resolution, image resolution was reduced by applying a downscaling factor equal to 10 before computing the diversity metrics, as frequently applied in the optical diversity context (Rocchini et al., 2016).

Standard deviations were separately computed on each of the three bands (red, green, blue) and the resulting statistics were then averaged to obtain a standard deviation (SD) for each image. Computing standard deviations in square moving windows and not only in the whole image allowed us to measure the spatial variability occurring at the scale of interest (i.e., flower scale in our study), and that can therefore be attributed to flowers. A window of 5 pixels × 5 pixels was chosen, corresponding to around 2.5 cm × 2.5 cm. As described in Rocchini et al. (2021), we first computed a diversity map by assigning to each pixel the SD of reflectance values in a square moving window of five pixels, and then obtained a summary metric for each image by averaging the diversity map. Lastly, we computed the Rao’s index, which quantifies the difference in reflectance values between two pixels drawn randomly with replacement from a set of pixels by considering their abundance and their pairwise distance (Hauser et al., 2021; Perrone et al., 2024). Rao’s index is a multidimensional index, which means that the diversity in multiple variables can be evaluated at the same time. In our study, the three bands (red, green, blue) were used. Using the “paRao” function of the “rasterdiv” package, Rao’s index was calculated in square moving windows of five pixels, applying Euclidean distance as a distance metric (Rocchini et al., 2021).

### 2.3. Part 2 - Image analysis using crowdsourced platforms

#### 2.3.1. Collection of images and platforms involved

The second part of the study was to identify objects in grasslands that people are interested in. Here, we assumed that the content of images (objects) uploaded to online platforms is indicative of people’s self-reported appreciation (Lee et al., 2022). To gather publicly available images posted online and georeferenced within the administrative boundaries of the study region (Figure 2), two crowdsourced platforms Flickr and iNaturalist were selected. Both platforms are commonly used by people to share their interactions with plant and animal species (Havinga et al., 2020). Flickr is a social media platform where photographers share their visual appreciation of nature, including casual interactions with nature, including but not limited to photographs of species, whereas iNaturalist is a citizen science platform where users specifically focus on sharing their observations of species (Havinag et al., 2020). We retrieved all publicly shared georeferenced images and their associated metadata, including user and image ID, the date of taking the image and a URL link to the image using the “photosearcher” package in R (Fox et al., 2020) to access the Flickr Application Programming Interface (API) and the “rinat” package in R (Barve and Hart, 2017) for the iNaturalist API. The data collected covers the years 2004 to 2023 and, after removing duplicate images, results in 1,718,001 unique images from 27,949 users in Flickr and 143,850 images from 3745 users in iNaturalist in the whole study region. These images were filtered for their location in grasslands using a grassland habitats map from Huber et al. (2022), resulting in a total of 88,025 georeferenced images by 3,831 Flickr users and 10,772 georeferenced images shared by 725 iNaturalist users. These images were then analysed for their content, focusing on the objects and especially the biological species depicted, highlighting the self-reported species of users on both platforms as an indicator of revealed preference (Lee et al., 2022).

#### 2.3.2. Extraction of objects from images

The content analysis consists of two methodological steps. First, we filtered all images for those focusing on objects belonging to nature. Second, we analysed what exactly was on the images in terms of specific plant and animal species. This two-step approach helped to save computational effort, since only the zoomed-in images of natural objects derived by step one were used for the in-depth analysis. To extract only those images showing nature and natural objects, the content of the images was extracted using Google Cloud Vision (GCV, https://cloud.google.com/vision/) by accessing its API using the R package “imgrec” (Schwemmer, 2019). GCV is a pre-trained machine-learning algorithm that has demonstrated a high ability to detect both natural and non-natural features in social media images (Fox et al., 2021; Wilkins et al., 2022). To extract nature images, we first ran all images through the algorithm using the “label” feature, which returns labels describing the image content, including natural labels (e.g., grassland, landscape, flower) and non-natural labels (e.g., building, road, car). For each image, we extracted up to 20 labels, considering only those with a confidence score of 60% or higher (Gosal et al., 2019) and a total of at least 15 occurrences in the dataset. Second, we adopted the framework proposed by Chai-allah et al. (accepted), classifying the GCV labels into non-natural and natural labels, with the latter being classified as landscape (e.g., landscape, mountain, sky) or objects/species (e.g., flower, bird, insect). The full label classification is listed in Table S3 in supplementary material 2. From nature images (i.e., images with more than 50% nature labels), we selected those with more than 50% object labels to build up our species appreciation dataset. Finally, since geolocations can be prone to error (Zielstra and Hochmair, 2013), we limited our analysis to images in grasslands by selecting only images with the presence of grassland-related labels (i.e., grass, grassland, meadow, and pasture), resulting in a final dataset of 5631 images shared by 808 Flickr users (mean = 6.9) and 4944 shared by 451 iNaturalist users (mean = 10.9).

We ran the GCV again to detect objects using the “object” feature. This feature has better capabilities to accurately detect the species present in the images compared to the “label” feature (https://cloud.google.com/vision/docs/features-list). For each image, we retrieved up to three unique objects. The confidence score for the retrieved objects ranged from 0.5 to 0.99 (mean = 0.79), which can be considered high. Thus, we decided to not further exclude any objects based on the confidence score. However, we limited our analysis to objects that occurred at least twice in total and excluded generic objects (i.e., animal and plant), objects describing pet and zoo animals (e.g., cat, dog, and brown bear) and non-natural objects (e.g., person, bicycle and car). The complete list of included and excluded objects can be found in Table S4 in supplementary material 2.

Since the hierarchical functioning of the GCV algorithm can classify an image using multiple related objects, such as “bird” as the first object and “swan” as the second object, or one object, simply “bird” or “swan”, a pattern predominantly occurring for birds and insects, we decided to simplify the classifications by grouping objects. All bird-related objects (i.e., bird, duck, sparrow, goose, swan, falcon, raven, eagle, magpie, owl, blue jay and woodpecker) were merged into the category “bird” and all insect-related objects (i.e., insect, moths and butterflies, butterfly, dragonfly, beetle, bee, ladybug and caterpillar) into the category “insect”. We extracted only unique objects returned by GCV, meaning within an image, three objects belonging to the group “bird” would be reduced to one “bird” object. This step of the analysis revealed a general overview of the objects on the images taken without diving into a taxonomic analysis.

#### 2.3.3. Extraction of wildlife from images

To better understand the wildlife shown in the images, we carried out a taxonomy analysis using the computer vision model BioCLIP (Stevens et al., 2024). It is derived from the CLIP (Contrastive Language– Image Pre-training) model developed by OpenAI and trained on a large dataset of images including those from the Encyclopedia of Life project (https://eol.org/), which is a comprehensive collection of images and information about various species. Since the identification error of the model increases with increasing taxonomic detail, from kingdom to species (Stevens et al., 2024), we extracted different taxonomic levels of the objects found in the images. For each retained image, we first performed a taxonomic analysis at the family level. For this, only families identified with a confidence score of 60% or higher were considered. Next, we performed a genus-level analysis on these pre-filtered images, again retaining only those genera identified with a confidence score of 60% or higher.

## 3. Results

### 3.1. Predictors of people’s aesthetic appreciation of the plant communities (Hypothesis 1.1)

In total, 522 participants answered the questionnaire, which led to, on average, 55.3 ratings per grassland image (min = 39, max = 73). The questionnaire was answered by 47.9% female and 48.1% male respondents (1% diverse, 3.1% NA), which were mainly from Switzerland (48.9%) and Germany (44.3%), plus few from Austria (3.1%) and further countries or NA (3.8%). Age distribution of the respondents was reported in steps of ten years, with most people being quite evenly distributed between 20 and 70 years (Figure S1 in supplementary material 1). Yet, few younger (1.1% 18-20 years) and some older people (3.4% 70-90 years) were also responding (2.5% NA). Of all people responding on the profession (3.3% NA), 23.2% stated a relation to agriculture, 22.4% to ecology/nature conservation, 8.6% to both professional areas, and 42.5% to “neither of both, other professions or retired”.

People’s ratings of the aesthetic value of the 92 images of grassland plant communities varied widely, from min 1.79 to max 4.54 (mean = 3.15 and SD = 0.65; allowed range 0 = *unattractive* to 5 = *attractive*; Figure 3). Basic statistics of the perception of the three professional groups “agriculture”, “ecology/nature conservation”, and “neither of both, other professions or retired” (simplified to “*other*” in what follows) were highly similar (Table S1 in supplementary material 1) and mean values did generally not significantly differ (linear mixed model, p > 0.2). The aesthetic values of all respondents correlated strongly with those of all separate professional groups (r_Spearman_ 0.87 to 0.96, p < 0.001; Table 1). This is why the following is focused primarily on all responses together. Yet, ratings of people working in agriculture and in ecology/nature conservation were less closely correlated (r_Spearman_ 0.64, but p < 0.001; Table 1; Figure S2 in supplementary material 1), highlighting that the aesthetic rating somewhat varied among professional groups. For example, professionals related to ecology/nature conservation rated images of unfertilised grasslands on average a bit higher than agricultural professionals (3.82 vs. 3.42), and vice versa for intensively managed, fertilised grasslands (2.70 vs. 2.96; Figure S2 in supplementary material 1).

**Figure 3.**
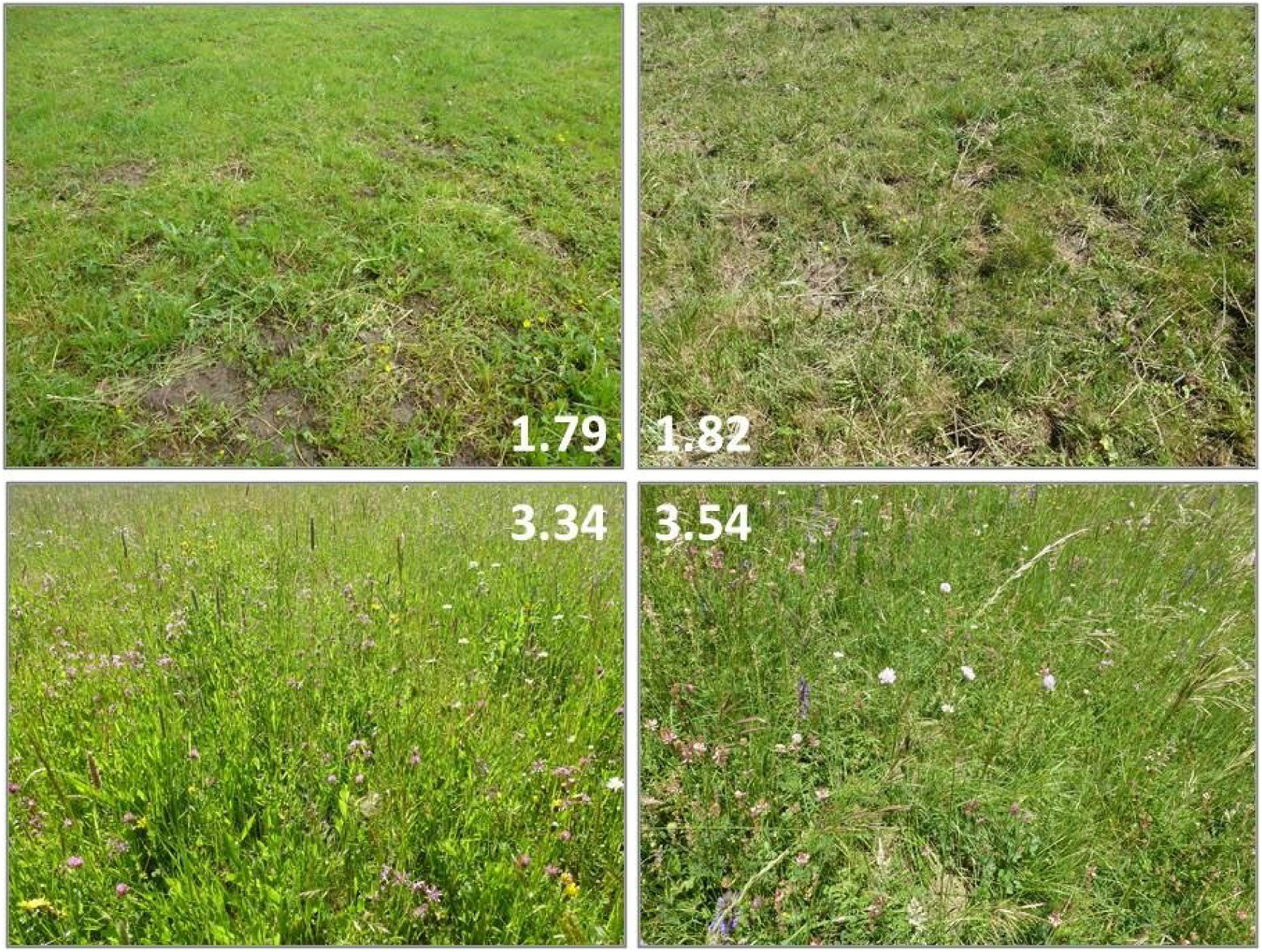
The four images with lowest (upper panel) and highest (lower panel) average ratings considering all responses (n = 522). Images that show signs of use (defoliation) and bare soil were perceived as particularly unattractive, while high flower cover and colour richness were perceived as attractive (Table 1, Table S2 in supplementary material 1).

**Table 1.** Spearman correlations between aesthetic ratings of different professional groups (in grey above) and with various grassland characteristics. Blue colours show positive predictors and red ones negative. While flower cover and colour richness have been extracted directly from the images used in the survey, all other information is based on field samplings and a land-use questionnaire with farmers. Significant levels: “***” = p < 0.001, “**” = p < 0.01, “*” = p < 0.05, “(.)” = p < 0.1, and “n.s.” = p > 0.1.

|  | Ratings all | Ratings agriculture | Ratings ecology | Ratings other |
| --- | --- | --- | --- | --- |
| Ratings agriculture | **0.87 \*\*\*** | - |  |  |
| Ratings ecology | **0.89 \*\*\*** | **0.64 \*\*\*** | - |  |
| Ratings other | **0.96 \*\*\*** | **0.78 \*\*\*** | 0.83 *** | - |
| Categ. flower cover (picture) | **0.66 \*\*\*** | **0.38 \*\*** | **0.66 \*\*\*** | **0.72 \*\*\*** |
| Flower cover perc. (picture) | **0.6 \*\*\*** | **0.35 \*\*** | **0.59 \*\*\*** | **0.65 \*\*\*** |
| Number flower colors (picture) | **0.6 \*\*\*** | **0.46 \*\*\*** | **0.61 \*\*\*** | **0.56 \*\*\*** |
| Stand height | **0.6 \*\*\*** | **0.65 \*\*\*** | **0.4 \*\*\*** | **0.6 \*\*\*** |
| Cover sum forbs | **0.5 \*\*\*** | **0.29 \*** | **0.52 \*\*\*** | **0.52 \*\*\*** |
| Number plant species | **0.42 \*\*\*** | 0.22 (.) | **0.56 \*\*\*** | **0.37 \*\*** |
| Shannon index plants | **0.42 \*\*\*** | 0.23 (.) | **0.52 \*\*\*** | **0.41 \*\*\*** |
| Evenness plants | **0.32 \*** | 0.2 n.s. | **0.35 \*\*** | **0.34 \*\*** |
| Cutting frequency | 0.18 n.s. | **0.27 \*** | 0.02 n.s. | 0.21 (.) |
| Slope | 0.12 n.s. | 0.02 n.s. | 0.21 (.) | 0.11 n.s. |
| Peak standing crop | 0.1 n.s. | 0.26 (.) | -0.11 n.s. | 0.14 n.s. |
| Cover sum cryptogams | -0.01 n.s. | -0.15 n.s. | 0.23 (.) | -0.07 n.s. |
| Elevation | -0.13 n.s. | -0.08 n.s. | -0.11 n.s. | -0.11 n.s. |
| Fertilisation intensity | **-0.33 \*** | -0.11 n.s. | **-0.51 \*\*\*** | **-0.3 \*** |
| Grazing intensity | **-0.48 \*\*\*** | **-0.44 \*\*\*** | **-0.43 \*\*\*** | **-0.48 \*\*\*** |
| Forage value indicator | **-0.48 \*\*\*** | -0.21 (.) | **-0.63 \*\*\*** | **-0.45 \*\*\*** |
| Cover sum grasses | **-0.49 \*\*\*** | -0.22 (.) | **-0.65 \*\*\*** | **-0.45 \*\*\*** |

Correlation analysis revealed many positive and negative predictions of the aesthetic value of the plant communities (Table 1). The strongest positive predictors based on all responses were flower cover, flower colour richness, stand height, the cover sum of forbs and plant diversity metrics (Shannon diversity and species richness). A weak but significantly positive effect of weeds being visible in the images was found for the professional groups “ecology/nature conservation” and “other”, but not for “agriculture” (Table S2 in supplementary material 1). Furthermore, and in contrast to the other professional groups, ratings of people related to agriculture were most positively correlated with stand height but much less so with flowers, forbs, and plant diversity (Table 1). The strongest negative predictors of all ratings were cover sum of grasses, forage value indicator, grazing intensity, and fertilisation intensity. While a high cover of grasses was obviously directly visible in the images, and strongly negatively related with the cover of forbs (r_Spearman_ = −0.57, p < 0.001), the other measures cannot be seen on the images but are indicators for intensive management, which generally reduces positive predictors such as flower cover and plant diversity (e.g., correlations of fertilisation intensity with flower cover and plant species richness: r_Spearman_ = −0.34, p < 0.001 and r_Spearman_ = −0.7, p < 0.001, respectively). However, agricultural professionalś ratings showed less strong negative relationships with the formerly mentioned factors except for grazing intensity (Table 1). While grazing intensity and, for most groups, also fertilisation intensity were clearly negatively related to aesthetic ratings, cutting frequency was either insignificant or weakly positively related to aesthetic ratings (Table 1). This might, however, be indirectly driven by a strong negative correlation between cutting frequency and grazing intensity (r_Spearman_ = −0.72, p < 0.001), meaning that if grasslands were not intensively grazed, they had to be frequently cut.

In addition, signs of defoliation, bare soil, and a thick litter layer in the images were also strongly negatively related to the aesthetic ratings of all groups (Table S2 in supplementary material 1). There was, however, one notable exception: A litter layer did not significantly decrease the aesthetic ratings of people related to ecology/nature conservation. Note that those twelve images showing signs of defoliation and potentially a lot of bare ground were mainly pastures, which were stocked particularly early, explaining the negative effect of grazing intensity on the aesthetic value.

### 3.2. Spectral characteristics related to people’s aesthetic perception (Hypothesis 1.2)

The standardised automated process to extract flower cover and flower colour richness produced accurate and insightful results, reflecting the clear correlations with people’s aesthetic appreciation. The validation process demonstrated high accuracy of the RF classifier, with an overall accuracy of 0.95, a Kappa accuracy of 0.88, and a mean F1 score of 0.85. The automatically extracted flower cover for each image showed high correlations of 0.79 (p<0.001) when compared to manually coded data (Figure S3a in supplementary material 1). However, the automatically extracted flower cover values were significantly lower than those estimated manually, indicating a perceptual bias in human evaluation. We calculated flower colour richness for each image by counting the number of distinct flower classes that covered more than 0.005% of the image area, i.e., 600 pixels. This specific threshold yielded the highest correlation between manually and automatically estimated flower colour richness (r_Pearson_ = 0.64; p <0.001; Fig. S3b in supplementary material 1). High flower cover was associated with high appreciation scores, as in the correlation analysis previously shown in Table 1. However, at high levels of flower cover, this was only true when combined with high flower colour richness, as shown both by the manual and automatic estimates (Figure 4a). Images with very high flower cover (over 5%) typically featured uniform yellow blooms (such as dominance of *Taraxacum officinale* or *Ranunculus acris*) and received moderate appreciation scores. In contrast, the most highly appreciated images had moderate flower cover but displayed a flower colour richness between 4 and 5. Shannon and Simpson diversity indices of pixel classes frequency distribution explained a large portion of the variability in aesthetic appreciation (R^2^ of 0.42 and 0.43, respectively). The high correlation of these indices with flower cover (r_Pearson_ 0.98 and 1.00, respectively; both with p<0.001), confirmed that images with more evenly distributed class frequencies were those with higher flower cover, which were generally more appreciated by viewers. The three spectral diversity metrics were highly correlated with each other (r_Pearson_ between 0.862 and 0.999, p < 0.001), uncorrelated to flower colour richness, and significantly positively correlated to flower cover (r_Pearson_ between 0.43 and 0.46; p <0.001). However, the spectral diversity metrics were not correlated to people’s aesthetic appreciation (p > 0.05, Figure 4b).

**Figure 4.**
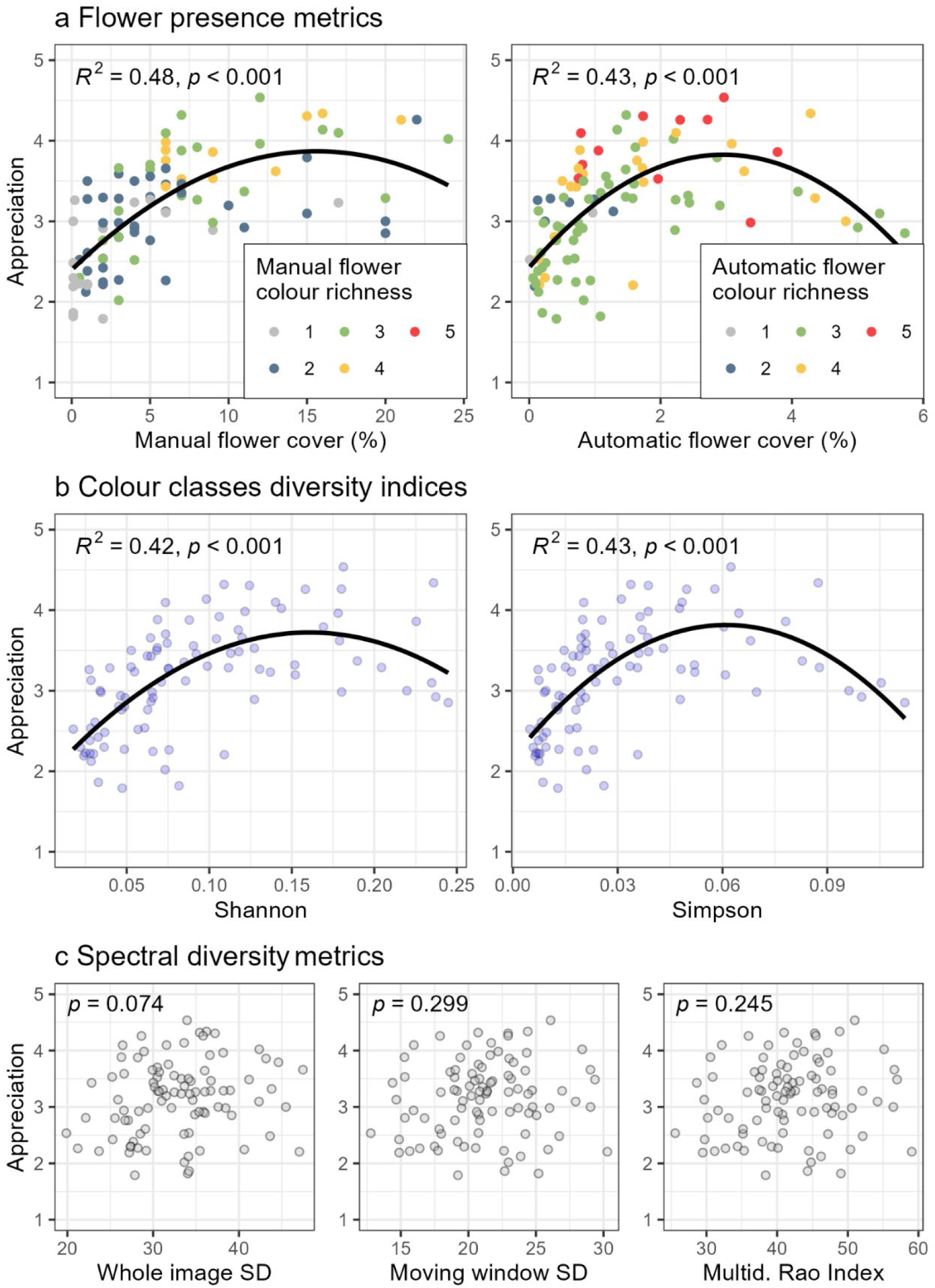
Relationships between aesthetic people’s appreciation and image flower presence metrics (a), colour classes diversity indices (b) and spectral diversity metrics (c). Flower cover, colour richness and the Shannon and Simpson indices of colour classes frequency distribution were associated with appreciation (aesthetic ratings), whereas the metrics measuring spectral diversity were not.

### 3.3. Frequent objects in grassland images (Hypothesis 2.1)

Flowers were the most photographed objects on both platforms, with 2,500 images on iNaturalist (> 56% of all images) and 2,198 images on Flickr (> 46% of all images; Figure 5). Flowers were, for instance, photographed two times more often on iNaturalist and 3.6 times more often on Flickr than insects, the second most popular object on both platforms. Birds appear as the third most photographed object on both platforms, with a similar number of images on both platforms. However, they were photographed 2.8 times less often than insects on iNaturalist (439 versus 1,230) and 1.2 times less often than insects on Flickr (510 versus 612). Following flowers, insects and birds, Flickr users frequently took images of livestock, i.e., cattle, horses, goats and sheep, reflecting a much greater interest in domestic livestock than iNaturalist users. Deer and frogs were similarly photographed on both platforms and on iNaturalist there are also images of mushrooms, lizards, spiders, and snails in the top 10 objects. Over 60% of Flickr users and 64% of iNaturalist users included in our study uploaded at least one image of flower(s), with an average of 5.1 flower images per user on Flickr and 9.5 on iNaturalist. Similarly, the average number of images per user for insects and birds was higher on iNaturalist compared to Flickr.

**Figure 5.**
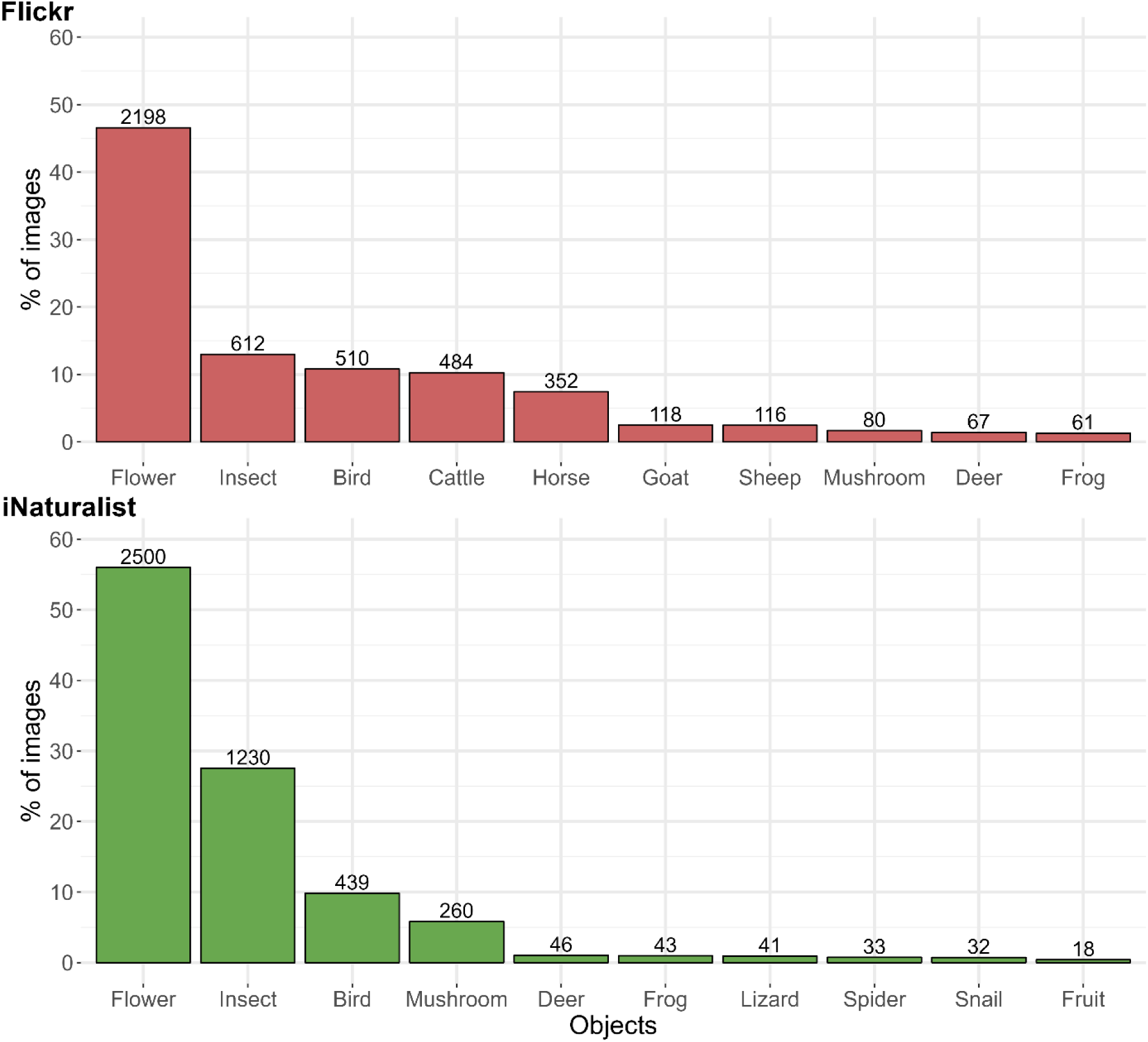
The ten most frequently photographed objects in Flickr and iNaturalist. The complete list of objects can be found in Table S4 in supplementary material 2. Total number of images included n = 4719 (Flickr) and n = 4464 (iNaturalist).

Images uploaded followed a distinct seasonal distribution. On both platforms, flowers and birds were photographed mainly during spring (March to May), while insects were photographed more often during the summer (June to August). On Flickr, cattle were photographed fairly consistently throughout autumn, spring, and summer, but there was a large drop-off in the winter, with only 14 out of 484 cattle images taken. Cattle were photographed by 25% of all Flickr users, while this was not relevant for iNaturalist users. Meanwhile, on iNaturalist, 76% of mushroom images were taken in autumn (Table S5 in supplementary material 2).

### 3.4. Wildlife in grassland images of Flickr vs. iNaturalist users (Hypothesis 2.2)

To understand differences in the content of images uploaded to a common, non-ecologically targeted platform (Flickr) and an ecologically focused platform (iNaturalist), we extracted familiae and genera of the wildlife depicted. Plant families make up the majority of the most frequently photographed wildlife familiae, accounting for 55% of the top 20 most photographed familiae on Flickr and 65% on iNaturalist (Figure 6). Asteraceae and Orchidaceae were the most photographed plant familiae on both Flickr and iNaturalist, followed by Rosaceae on Flickr and Fabaceae on iNaturalist. Papaveraceae were popular on Flickr but not on iNaturalist, whereas Lamiaceae was popular only on iNaturalist. While most top 20 plant familiae were forbs or familiae with mainly woody species (Rosaceae, i.e., roses, and Caprifoliaceae, i.e., honeysuckle), Poaceae (Gramineae, grasses) were also listed in both platforms, showing that not only plants with colourful flowers were among the frequently photographed taxa.

**Figure 6.**
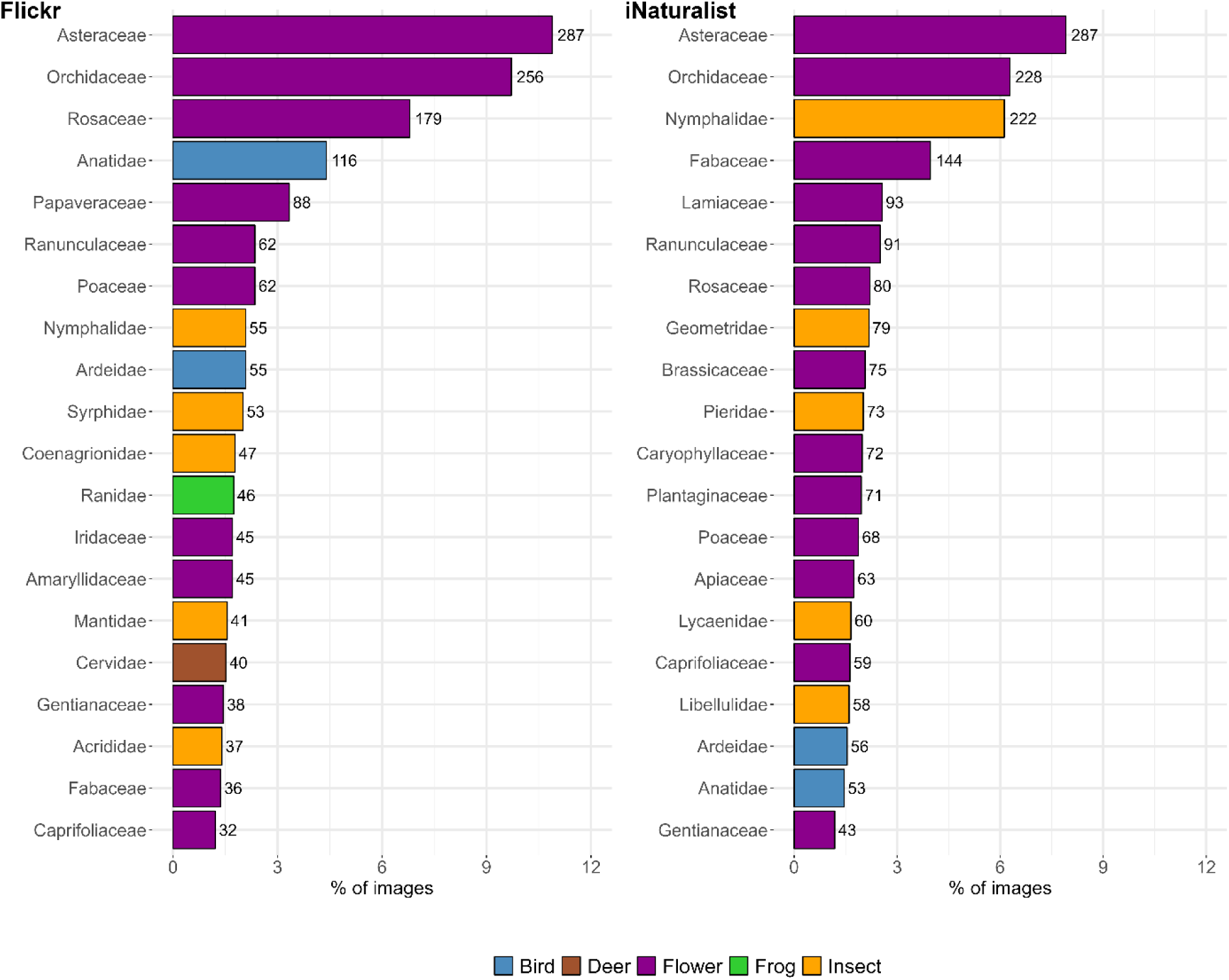
The 20 most frequently photographed familiae in Flickr and iNaturalist wildlife images. The number given at each bar indicates the total number of identifications of this kind. Total number of images included n = 2636 (Flickr) and n = 3628 (iNaturalist). The complete list of observed familiae can be found in Table S6 in supplementary material 2.

However, compared to the mean cover of plant families from the vegetation records in the canton of Solothurn (grasslands from which the pictures were taken, hypotheses 1.1 and 1.2), Poaceae were highly under-represented in both platforms (supplementary material 2, Figure S4). In the studied grasslands, Poaceae were the family with by far the highest mean cover per plot (61 %), followed by Fabaceae, Asteraceae and Ranunculaceae (all around 6-8%). Orchidaceae, the second most often photographed plant family on both flickr and iNaturalist, showed a very low cover in the studied plots (2.3 %), and was only present in four of the grasslands.

Five insect familiae ranked in the top 20 on both Flickr and iNaturalist, but with a higher proportion of all images on iNaturalist. Among these five popular insect familiae, Nymphalidae (brush-footed butterflies) were the most photographed on both platforms. However, they were about four times more frequent on iNaturalist compared to Flickr. The four other familiae show a difference in interest between Flickr and iNaturalist, with Flickr users photographing various families Syrphidae (hoverflies), Coenagrionidae (damselflies), Mantidae (praying mantis), and Acrididae (grasshoppers and locusts). In contrast, iNaturalist users primarily photographed further butterfly familiae (Geometridae, Pieridae, and Lycaenidae) and dragonflies (Libellulidae).

Vertebrates were rather rare within the top 20 families. Only two bird families made it into the top 20, both on Flickr and iNaturalist. These were Anatidae (water birds including ducks, geese, and swans) and Ardeidae (herons), with Anatidae being photographed more frequently on Flickr than iNaturalist. Frogs (family Ranidae) and deer (family Cervidae) appeared in the top 20 on Flickr, but not on iNaturalist.

The preference of Flickr and iNaturalist users for photographing plant species was also evident at the genus level (Figure 7). Among the top 20 most popular wildlife genera, about two third or more were plant genera, i.e., 65% on Flickr and 70% on iNaturalist. Although both Flickr and iNaturalist users showed a preference for photographing plants, the specific genera differed. Flickr users most often photographed *Papaver*, *Taraxacum*, and *Prunus*. Two orchid genera (*Ophrys* and *Spiranthes*) were also popular. In comparison, the most photographed genera on iNaturalist were *Orchis* (an orchid genera), *Veronica*, *Cirsium*, *Dactylorhiza*, and *Ranunculus*. Besides the woody species belonging to *Prunus*, which were ranked third on Flickr, non-forb plant genera, with respect to the species occurring in grasslands the study region, were quite rare. Yet, *Carex* (sedges) were ranked 11th on iNaturalist.

**Figure 7.**
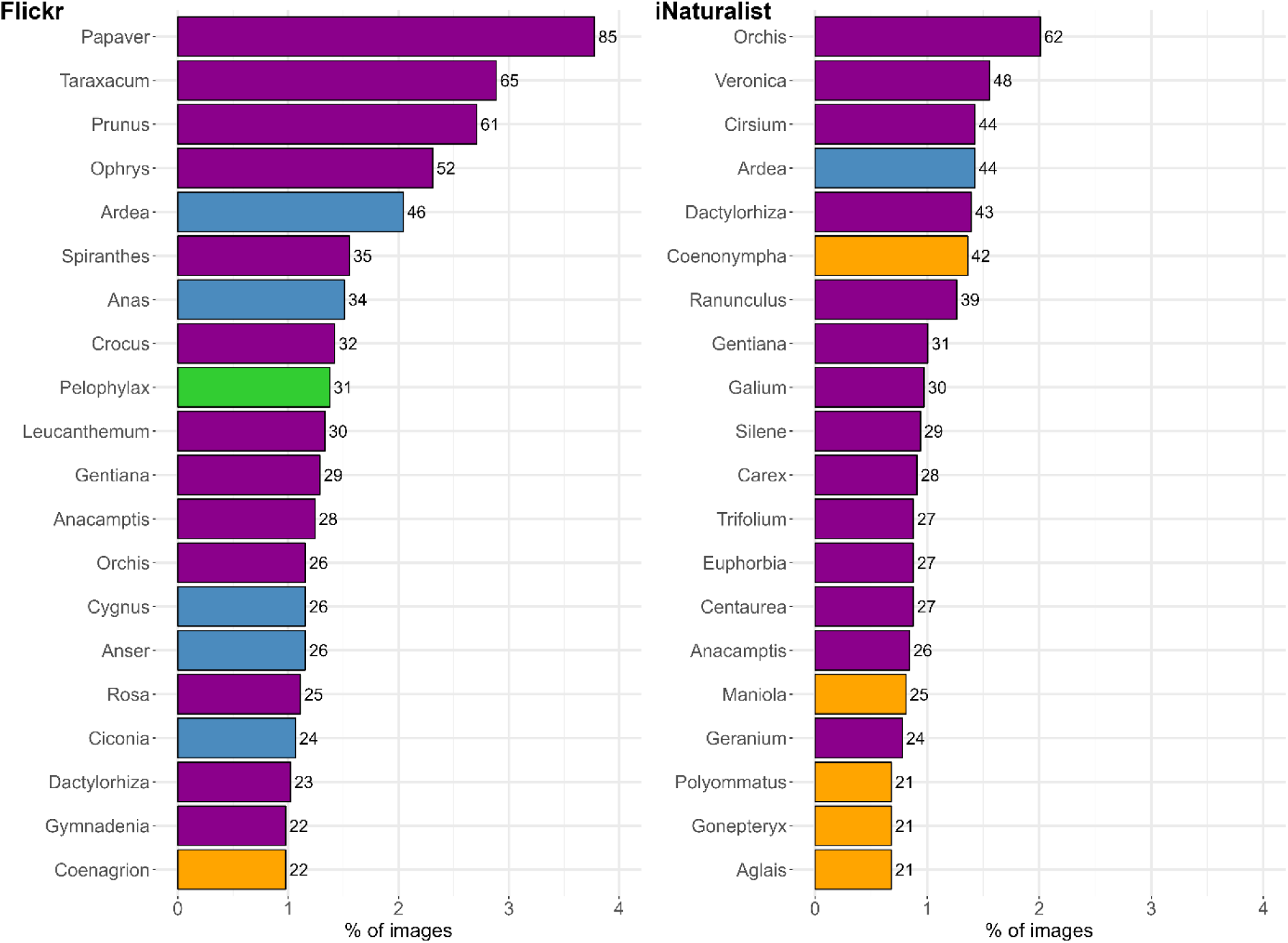
The 20 most frequently photographed genera in Flickr and iNaturalist wildlife images. The number given at each bar indicates the total number of identifications of this kind. Total number of images included n = 2251 (Flickr) and n = 3084 (iNaturalist). The complete list of observed genera can be found in Table S7 in supplementary material 2.

Compared to the cover of different plant genera from the vegetation records of the grasslands studied in the canton of Solothurn, grass genera were (like already for the families) underrepresented on Flickr and iNaturalist. The three grass genera *Poa, Lolium,* and *Festuca* had the highest mean cover (supplementary material 2, Figure S5). These were followed by *Trifolium* and *Ranunculus*, which were at the same time present in the highest number of grasslands. However, *Taraxacum* and *Veronica*, genera that were often photographed on Flickr and iNaturalist, respectively, also had rather high mean covers (3.8% and 1.3%, within top 10 and top 20, respectively) and were present in many of the grasslands (81 and 83, respectively).

Bird genera were the in total second most popular group on Flickr, accounting for 25% of the top 20 wildlife genera, while only one insect genus (*Coenagrion*, a genus of damselflies) appeared on this list. The top five bird genera photographed by Flickr users were mostly large water birds, with *Ardea* appearing at the top, followed by *Anas* and *Cygnus*. In contrast, insect genera made up 25% of the top 20 on iNaturalist, while in turn only one bird genus (*Ardea*, a genus of herons) made it into the top-20-list. In addition, the frog genus *Pelophylax* (a genus of true frogs) was among the top 20 wildlife genera on Flickr. Meanwhile, iNaturalist users showed a clear preference for photographing butterflies, as the five insect genera among the top 20 were butterfly genera. The top three of these were *Coenonympha*, *Maniola*, and *Polyommatus*.

## 4. Discussion

Our study identified clear differences in people’s aesthetic appreciation of different plant communities and of different objects (as well as taxonomic units) in permanent grasslands. These different levels of appreciation (or potentially interest, in case of the images from social media and citizen science platforms; Ghermandi et al., 2023) can be seen as indication for differences in the CES and in human-nature interactions taking place within the grasslands studied. Furthermore, we found biodiversity to be directly and indirectly linked to appreciation and CES, which will be discussed in the following.

### 4.1. Drivers of people’s aesthetic perception of grassland plant communities

#### 4.1.1. Positive and negative characteristics of the grasslands and their management (Hypothesis 1.1)

Our study largely confirmed the first hypotheses by proving colourful flower- and forb-rich grasslands to be most attractive to people, while grass-dominated and defoliated swards were not favoured. Our results are in line with previous work showing colourful flowering grasslands and similar grassy habitats to be highly attractive to humans in rural (e.g., Junge et al., 2009; Lindemann-Matthies et al., 2010; Nowak-Olejnik et al., 2020) and urban settings (e.g., Lindemann-Matthies and Brieger, 2016; Southon et al., 2017; Hoyle et al., 2018). Our study extends these findings by adding a more mechanistic land-use and biodiversity context (see also Richter et al., 2024). People’s appreciation was clearly linked to the level of management intensity, which strongly determined the perceived attractiveness and, in addition, generally drives the level of biodiversity found in agricultural grasslands. Extensively managed grasslands can harbour not only high numbers of potentially flowering forbs but also a high diversity of other plants and animals, especially when managed at low intensity for decades (Isselstein et al., 2005; Klaus et al., 2023). In contrast, intensively managed and fertilised grasslands, especially when grazed early in the season and dominated by grasses and plants with high forage value, are usually low in biodiversity of many taxa (Allan et al., 2014). Moreover, intensification generally decreases flower colour diversity and usually leads to early defoliation inhibiting a long and diverse flowering aspect (Binkenstein et al., 2013; Johansen et al., 2019). Though there can potentially also be many flowers in such stands, they are usually dominated by single species indicative of nutrient rich conditions and disturbance, such as dandelion (*Taraxcum* spec., yellow flowering aspect; Martinková et al., 2009) or tall growing Apiaceae (e.g., *Heracleum sphondylium*, *Anthriscus sylvestris*, white flowering aspect; Sheppard, 1991). This dominance of one or few highly similar species potentially causing a vast but monochromic flowering aspect is likely to be the factor explaining why at high levels of flower cover people’s aesthetic ratings stagnated or even tended to decrease (Figure 4a).

We specifically found signs of disturbance, such as short grass and bare soil, to be perceived as unattractive. This is in line with Junge et al. (2015) that found early stages (March) and post-harvest stages (summer) with a lot of bare soil, short grass and a brownish appearance to be strongly disliked by people. The same study also found that extensively managed grasslands were preferred over intensively managed ones (Junge et al., 2015). While grazing intensity was strongly negatively related to perceived aesthetic quality, cutting frequency was not such a strong predictor. This result should be treated with caution since in our study grasslands with signs of defoliation and bare soil were all intensively grazed grasslands, while cut grasslands (meadows) were not yet harvested. Thus, the results likely depend on the seasonality of the images presented and could change if these were taken a bit later (end of May), when intensive meadows would also be cut, revealing mostly unattractive short grass and bare ground (Junge et al., 2015).

Differences in the aesthetic ratings of plant communities as derived from different professional groups were found but did not compromise the general patterns described before. Yet, appreciation was to some degree also linked to professional expertise. For example, nature conservationists rated unfertilised grasslands more positively than agricultural professionals, most likely because conservationists acknowledge the biodiversity value in such unfertilised stands and relate less to agricultural productivity (Peter et al., 2022). In addition, agricultural professionals were less negative about grasslands with intensive management than all other groups, potentially because productive grasslands are still a main pillar of grass-based food production in Europe (Isselstein et al., 2005). On a different spatial scale, a similar situation was found by Junge et al. (2011), with farmers rating the aesthetic quality of a more intensively used (grassland) landscape higher than non-farmers. Potentially surprisingly, when an agriculturally relevant weed species was visible in the image, this was slightly but significantly positively related to the aesthetic ratings of people belonging to ecology/nature conservation and also other persons. This was not the case for agricultural professionals. The reason behind the positive relation between weeds and aesthetics might be that, in addition to their general value for general biodiversity and pollinators, some agricultural weed species can show intensive and attractive flowering aspects (e.g., species of Rhinanthus, i.e., rattles, and Cirsium, i.e., thistles; Balfour and Ratnieks, 2022). This is underlined by the genus of *Cirsium* being among the most frequently photographed plant genera in the second part of this study (Figure 7). These results highlight the need to protect colourful flowering extensively managed grasslands in recreational landscapes.

Our study underlines the value of biodiverse grasslands for human-nature interactions. Similar to Lindemann-Matthies et al. (2010) and Nowak-Olejnik et al. (2020), we found plant diversity to be directly related to people’s aesthetic ratings and, consequently, to the CES provided. While in the urban context, plant diversity and flower (colour) richness can be decoupled from native biodiversity by using seed mixtures of (partially) non-native species (Jiang and Yuan, 2017; Hoyle et al., 2017, 2018), this is not well possible in agriculturally managed permanent grasslands. At least in the atlantic, alpine and mediterranean biogeographical regions of Europe, the latter are clearly dominated by native species (Axmanová et al., 2021), of which many became to some degree rare or endangered. Thus, in the rural context, the aesthetic appreciation clearly favours extensively managed semi-natural grasslands with high forb richness and, naturally related, high biodiversity. This calls for a protection and, in face of the EÚs Nature Restoration Law (Regulation 2024/1991), restoration of forb-rich semi-natural grasslands, especially in areas that are accessible to people.

#### 4.1.2. Spectral characteristics related to people’s aesthetic perception (Hypothesis 1.2)

We applied several standardised methods to automatically extract information from images, including i) automatic flower cover and colour richness classification, ii) pixel class diversity indices, and iii) spectral diversity metrics, and we evaluated their capability in predicting people’s aesthetic appreciations. These methods worked well, showing a high potential for practical application in similar cases. Notably, our case study revealed an interplay between flower cover and flower colour richness in predicting aesthetic ratings, which accounts for the non-linear relationship observed between flower cover and appreciation. The analysis of the Shannon and the Simpson indices, based on the frequency distribution of the pixel classes, with aesthetic ratings did not yield additional insight, since both were strongly correlated with flower cover. As mentioned before, these correlations can be attributed to the dominance of few flowering species so that increased flower cover led to a more even distribution of pixels among various classes (i.e., higher Shannon and Simpson indices). Conversely, low flower cover means most pixels were classified as “green vegetation or soil,” leading to an uneven class distribution (i.e., lower Shannon and Simpson indices).

The spectral diversity metrics were positively but only weakly correlated with flower cover, as in Perrone et al. (2024), and not significantly related to aesthetic ratings of the grassland plant communities. This is probably because spectral diversity is predominantly influenced by light conditions (Arroyo-Mora et al., 2021), grassland vertical structure (Conti et al., 2021), the presence of dead biomass (litter, Rossi et al., 2022), and differences in the colours of vegetative organs (Binkenstein et al., 2013). Notably, flowers constituted less than 1% of pixels in over half of the images analysed. This finding suggests that spectral diversity is no suitable predictor of grassland aesthetic ratings and cannot be used to ease and standardise the assessment of vegetation aesthetics, falsifying our second hypothesis. However, the RF model’s high accuracy, along with the strong correlation between automated and manual estimates of flower cover and flower colour richness, confirmed the reliability of the classification of grassland images. This ability to rapidly classify many grassland images creates new possibilities for ecological research, allowing for more extensive and frequent site monitoring (Andreatta et al., 2023; Torresani et al., 2023).

### 4.2. What people like to photograph in grasslands

#### 4.2.1. Frequent objects in social media derived grassland images (Hypothesis 2.1)

Our results show that flowers are the most frequently photographed objects in grasslands, reflecting their cultural significance. This was consistent with our third hypothesis and reflects people’s deep connection with flowering plants, which elicit positive emotional responses (Havinga et al., 2024). The vibrant colours and visual appeal of flowers, especially in contrast to the dominant green of grasslands, naturally draw attention and contribute to the attractiveness of a habitat (Mou et al., 2023). Flowers are more abundant and easier to photograph than birds or insects, as they are static and pose no perceived threat (Austen et al., 2021). Though less photographed, birds and insects still captivate visitors, highlighting both their ecological importance and the enjoyment people find in spotting them.

In addition to flowers, the presence of different kinds of livestock was appreciated by people, confirming previous research on people’s preferences in agricultural landscapes using surveys (Junge et al., 2015) or social media (Chai-allah et al., accepted). These studies found cattle to be valued by people assessing landscape aesthetics. However, the fact that they were photographed exclusively on Flickr versus iNaturalist highlights that people’s interests and motivations often influence what they share online during outdoor experiences (Di Cecco et al., 2021). It is likely that Flickr users are more inclined to share photographs of landscapes and aesthetic appeal symbolising rural identity, such as cows in a field, whereas iNaturalist users are more inclined to share close-up images of specific species lacking contextual context of the landscape (Lopez et al., 2020). However, beside livestock, the objects more frequently included in the top 20 in both platforms were highly similar, indicating that availability (abundance) and commonness of the objects also played a major role in determining their frequency on the images, a consistent pattern also in previous research (Havinga et al., 2023).

#### 4.2.2. Differences in wildlife taxa photographed by common visitors and naturalists (Hypothesis 2.2)

Despite the general patterns and similarities found before, our analysis also revealed differences between users of the two platforms, confirming our fourth hypothesis. These differences in occurrence and abundance ranking imply that commonness is not the only factor driving the frequency of what is photographed. While commonness is clearly a driver of how often an object is photographed and uploaded to a social media platform (Havinga et al., 2023), some of the taxa photographed here are not very common or even particularly rare such as orchid species (Orchidaceae in families and genera of *Orchis*, *Ophrys*; *Dactylorhiza*, and *Anacamptis*; Djordjević and Tsiftsis, 2022). This could be confirmed via a comparison with the mean cover as well as occurrence of families and genera on the grasslands used for the image analysis, where Poaceae were much more dominant, and Orchidaceae on the other hand were much rarer than the pictures uploaded to Flickr and iNaturalist would suggest.

Analysis at the taxonomic levels of family and genus revealed that common visitors (Flickr users) and naturalists (iNaturalist users) expressed different appreciations for species, which most likely reflects their main motivations (Havinga et al., 2023). There is an expected bias for plants with colourful flowers or leaves (August et al., 2020; White et al., 2023). Flickr images include some more generalist taxa that are easily recognizable or aesthetically pleasing, such as species of *Prunus* (cherries, plums etc.), *Taraxacum* (dandelion), and *Papaver* (poppy), but also ducks and herons. These are quite common and not of a specific ecological interest, at least in terms of nature conservation. In contrast, iNaturalist images included species that may not be commonly perceived as particularly beautiful, such as species of *Cirsium* (thistles) and *Carex* (sedges). This may be because iNaturalist users tend to consider the visual appeal of species together with their ecological significance. Most notably, there is a high frequency of orchid species on iNaturalist and, with a lower ranking, also on Flickr. These species can be seen as quite rare and not very easy to (accidentally) find in the study area. They are also of high relevance for nature conservation, due to their natural rarity and anthropogenic decline (Kull et al., 2016). Thus, besides a potentially positive effect of positive interactions with nature on people’s ecological standpoint and pro-environmental behaviour (Mackay and Schmitt, 2019), such eco-touristic activities can also harm populations of rare and endangered species (Ballantyne and Pickering, 2012). These findings also suggest that exploring users of broader social media, such as of Flickr, can complement citizen science approaches in the attempt to understand how people interact with certain species from a conservation perspective, and to identify locations of potentially damaging eco-tourism.

The familiae and genera of grassland-related wildlife that was found to be attractive to people, has two slightly contradicting implications. First, in line with the results obtained for species-rich plant communities, many of the species included in the attractive wildlife depend on species-rich, extensively managed grasslands. Examples include plant species of *Gentiana* (gentian) and all orchids (Djordjević and Tsiftsis, 2022) but also most butterfly species (Öckinger et al., 2006), which depend on late (or incomplete) cutting or grazing allowing plants and animals to flower and reproduce. Since most of these species are directly dependent on grassland habitats, they underline the suitability of our approach to identify typical grasslands-based wildlife that is attractive to people. On the other hand, some rather common and not at all grassland-typical wildlife was frequently included, especially in images from Flickr. Examples are *Papaver* (poppy) species, which in the study area only occur in arable ecosystems, and classical waterfowl (ducks, swans, herons), which likely belong to ponds surrounded by grassy strips. The images might originate from people using grasslands only as a spot to observe neighbouring habitat types. Having both rare and common wildlife frequently photographed shows that what is considered attractive by people can be very diverse and our landscapes therefore need to accommodate many different interests and values.

While crowdsourced platforms offer fresh insights into how people value biodiversity, they come with biases that must be acknowledged (August et al., 2020; Di Cecco et al., 2021). The users of these platforms may overrepresent certain socio-demographic groups, influenced by technology access or the users targeted by each platform (Ghermandi et al., 2023; Venter et al., 2023). Though these platforms are often biased towards accessible areas (e.g. nature trails), this fine-scale spatial bias does not necessarily mean that recorded taxa are unrepresentative of the wider area (Geurts et al., 2023). Though season colour variation in plants may impact aesthetic perception (Campbell-Arvai et al., 2024), our study focused on people’s appreciation of species without accounting for seasonal changes. Future research should explore how seasonality, especially the flowering season, affects people’s appreciation of different kinds of nature.

### 4.3 Joint discussion and implications for landscape management

We identified different grassland characteristics that contribute to an attractive grassland landscape. Both approaches used in this study revealed the aesthetic value of extensively managed, species-rich grasslands that contain various flowers and wildlife. We found high plant diversity and many familiae and genera that belong to such ancient, nutrient-poor grassland ecosystems to be directly linked to people’s aesthetic preferences and human-nature interactions. This calls for protecting and enhancing these grasslands in regions where CES and ecotourism need to be increased, e.g., in the surroundings of a new settlement. Agricultural policies can support such aesthetically attractive but economically not viable agricultural habitats via different pathways, such as with direct payments for (cultural) ecosystems services (e.g., Farley and Costanza, 2010) and result-based schemes that account for the abundance of particularly attractive species such as orchids (e.g., Šumrada et al., 2021) and for extensive grazing with livestock (Matzdorf et al., 2008). However, as specified above, increased human-nature interaction can also cause issues, for example, when ecotourism gets in conflict with the protection of rare and endangered species (Ballantyne and Pickering, 2012).

Since flowers appeared to be a particular part of grassland CES, while signs of harvesting were particularly unattractive, suitable measures to improve the aesthetic quality of intensively used grassland landscapes are delaying or diversifying harvest dates, and, even easier to implement, leaving stripes of vegetation with high forb cover uncut. This doesn’t even need to compromise agricultural production, as shown by Ravetto Enri et al. (2017), which developed a rotational grazing system, which, during the main flowering period, temporarily excludes a parcel from grazing. They achieved enhanced flower resources without reducing farm-scale production (Ravetto Enri et al., 2017). In addition, measures of ecological restoration that increase forb (species) richness in de-intensified but species-poor plant communities can also increase the aesthetic value of a late-cut grassland, highlighting co-production of CES and nature conservation (Bullock et al., 2021). However, particularly rare species such as orchids are usually not directly included in such a restoration approach, highlighting the importance of prioritising the conservation of ancient species-rich grasslands.

Although our study did not compare the relative importance of the two aspects assessed, i.e., plant communities and objects/wildlife, we assume that the effect is additive, with attractive objects in attractive plant communities to be the most valued case. Future research might want to explore this and the CES of other agricultural habitat types in more detail.

## 5. Conclusions

Our study has shown several biodiversity-related features of agricultural grasslands to be attractive to people. This knowledge allows us to specifically manage land for CES and human-nature interactions by supporting extensive management and boosting the ecological restoration of grasslands. The attractive features we identified can be used as indicators for grassland CES and inform result-based agri-environment schemes, for example, to financially remunerated farmers for the ecosystem services provided by their land. Future research can add to this by extending our approach to other land use types and landscape-scale evaluations. Since the people’s perceptions and interests were quite diverse, favouring both rare and endangered as well as common wildlife, our landscapes need to be diverse and accommodate a range of different habitats.

## Supporting information

Pictures used in questionnaire survey (part of supplementary materials)

## Acknowledgements

We are grateful to all people involved in the ServiceGrass project, especially Andreas Lüscher, Nina Buchmann, Nadja El Benni, Pierrick Jan as well as all student helpers involved in the study. We further want to thank all farmers, who granted access to their grassland parcels, all people who answer the online questionnaire, and all Flickr and iNaturalist users that unconsciously supported our study.

## Data availability statement

Data will be shared upon request. Code will be made available upon acceptance.

Please note: For the purpose of easing the peer review, we include the supplementary material in the main file.

## Supplementary material 1

### Questionnaire text

Questionnaire text translated to English (see below for the original German-language questionnaire and all pictures included in this study):

#### Introductory text

“With this questionnaire we would like to understand what characteristics of grassland are considered beautiful in the sense of contributing to an attractive landscape. Therefore, we would like you to rate the following ten photos of grasslands in terms of their perceived beauty. Please try to focus only on the plant communities and not on the technical quality or perspective of the photos.

Thank you very much for your support of our research!

The results of this survey will be used anonymously for the ServiceGrass research project by ETH Zurich and Agroscope to investigate socially relevant ecosystem services. Ecosystem services are benefits grassland provides to humans, e.g. it stores carbon, prevents erosion, provides habitat for animals … and can look beautiful!

By completing this survey, you agree that your answers may be used anonymously for research in the ServiceGrass project of ETH Zurich and Agroscope.”

#### Start of questionnaire

“Now the photos will follow. Please rate the grasslands freely according to your gut feeling and personal aesthetic perception.

How attractive do you find this grassland? (“not beautiful” to “very beautiful” as a five-point Likert-scale question; series of ten pictures)”

#### End of questionnaire

“Thank you for your evaluation of the grasslands! Now just a few brief personal details: Which country do you live in? (Switzerland, Germany, Austria, other)

What is your age? (in steps of 10 years)

Which gender do you consider yourself to be? (diverse, female, male)

Is your professional activity related to the following fields? (agriculture, ecology or conservation, none of these/other/not working)”

### Example of Questionnaire in German language

Submitted as separate file

### All pictures used for the questionnaire

Submitted as separate file

Supplementary material 2

---

**Figure S1.**
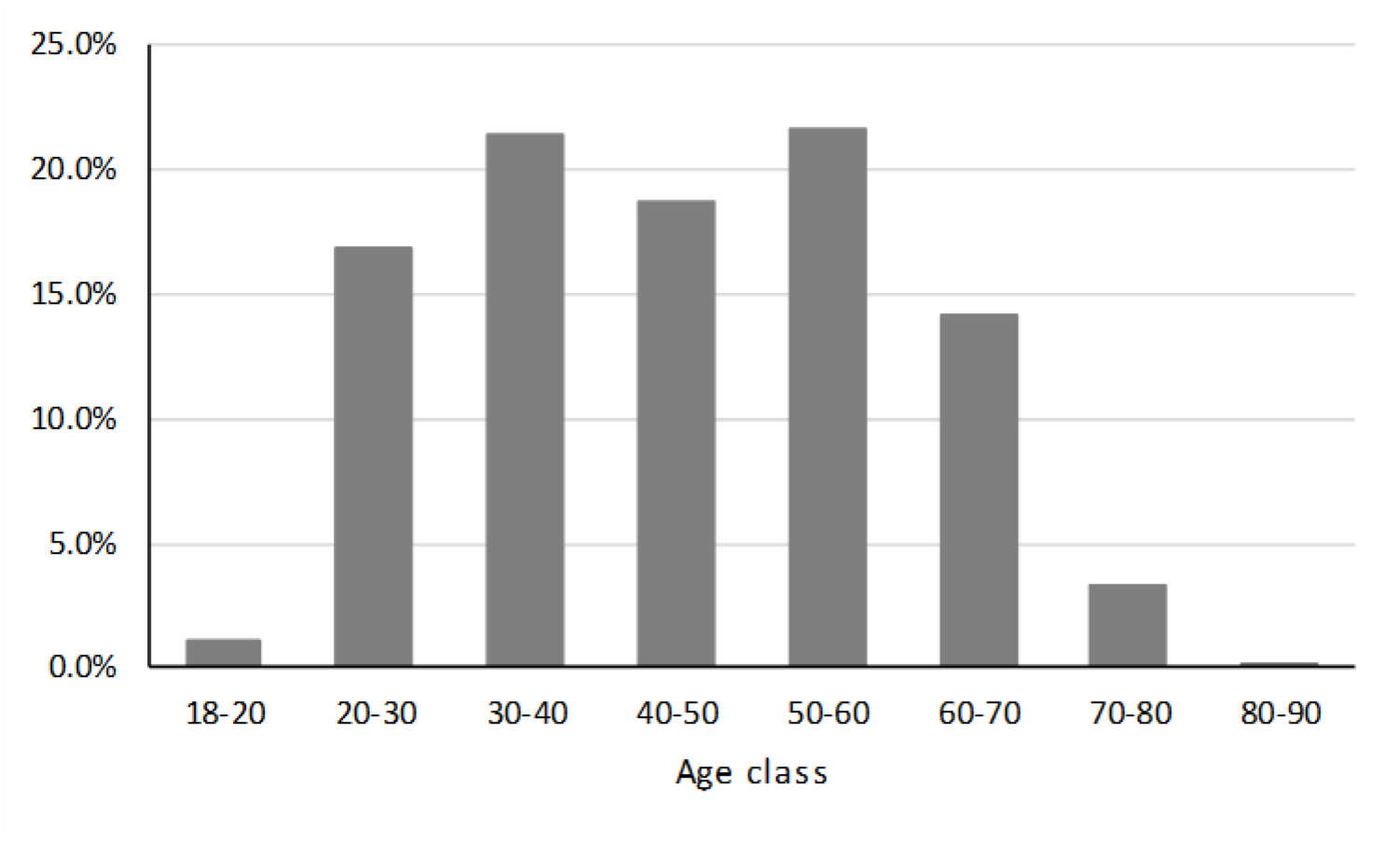
Distribution of age classes of survey participants (n = 522)

**Figure S2.**
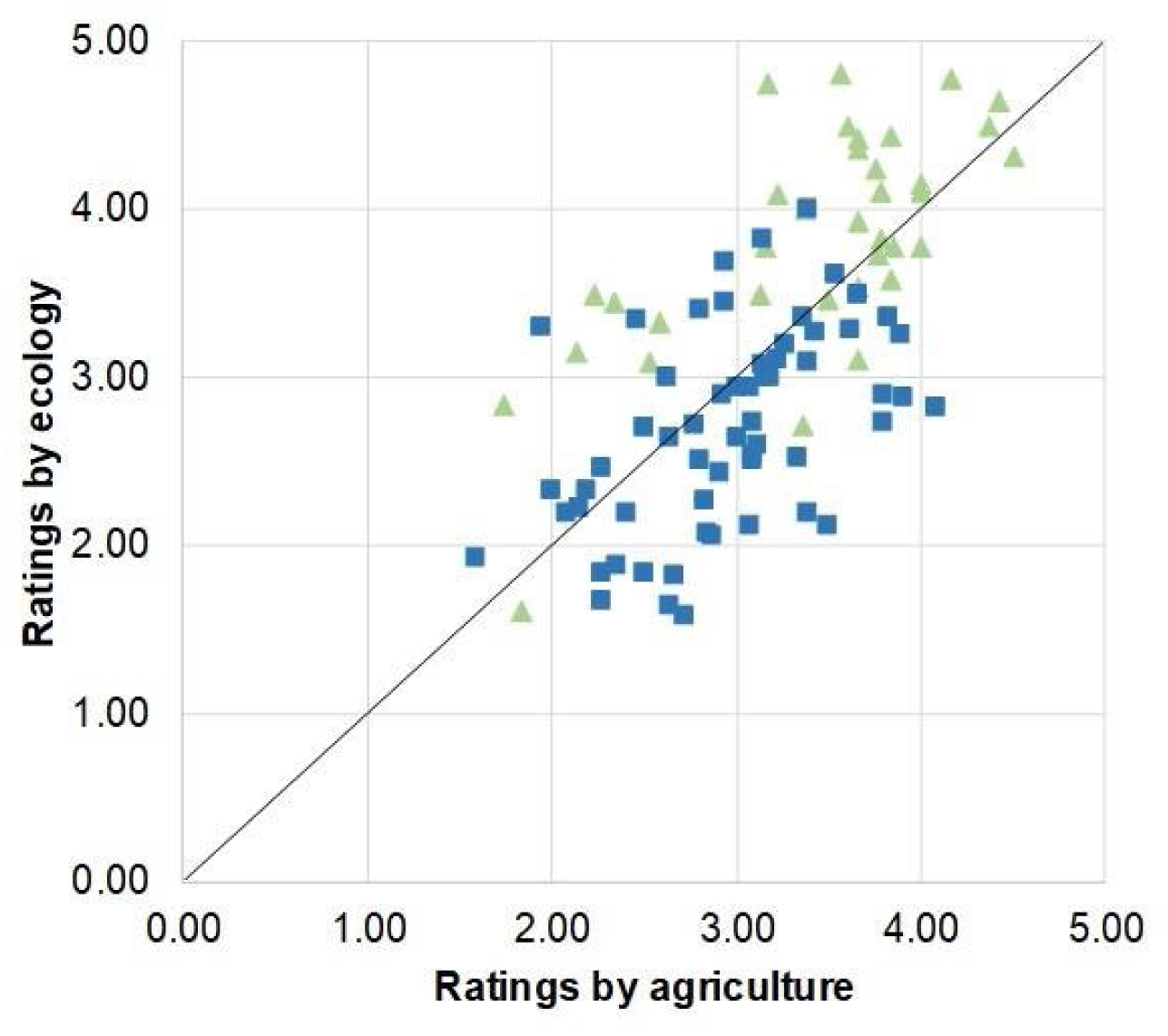
Aesthetic ratings of 92 grassland images by agricultural and ecological professionals. Blue squares depict fertilised and intensively managed grasslands, and green triangles depict extensive grasslands in which no fertilisation takes place. The black line represents the 1:1-line.

**Figure S3.**
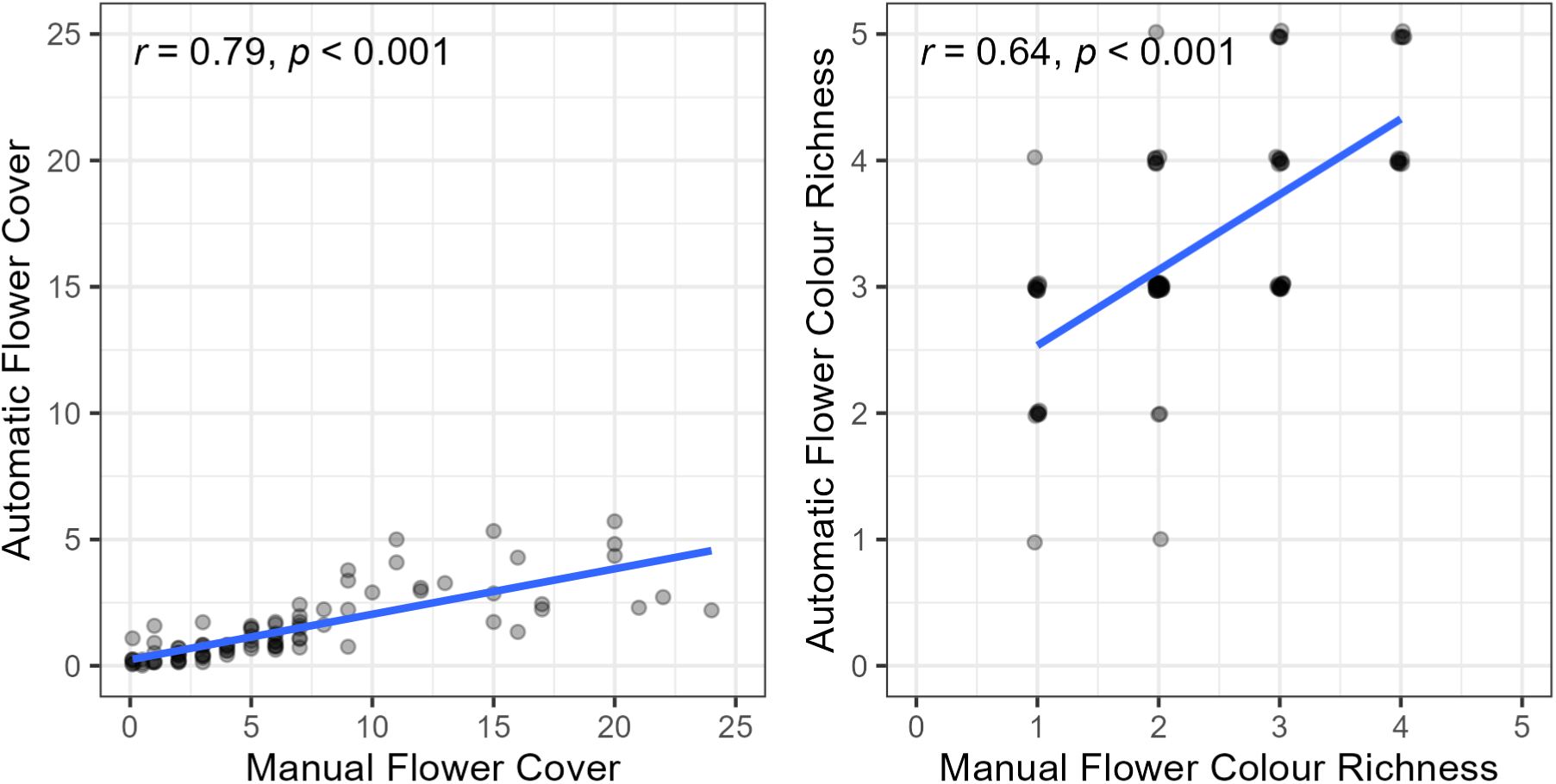
Relationships between automatically and manually estimated flower cover and flower colour richness. Pearson Correlation Coefficients and p-values are reported in the top-left corner of each panel. A small amount of jittering was applied to reduce overlap.

**Figure S4.**
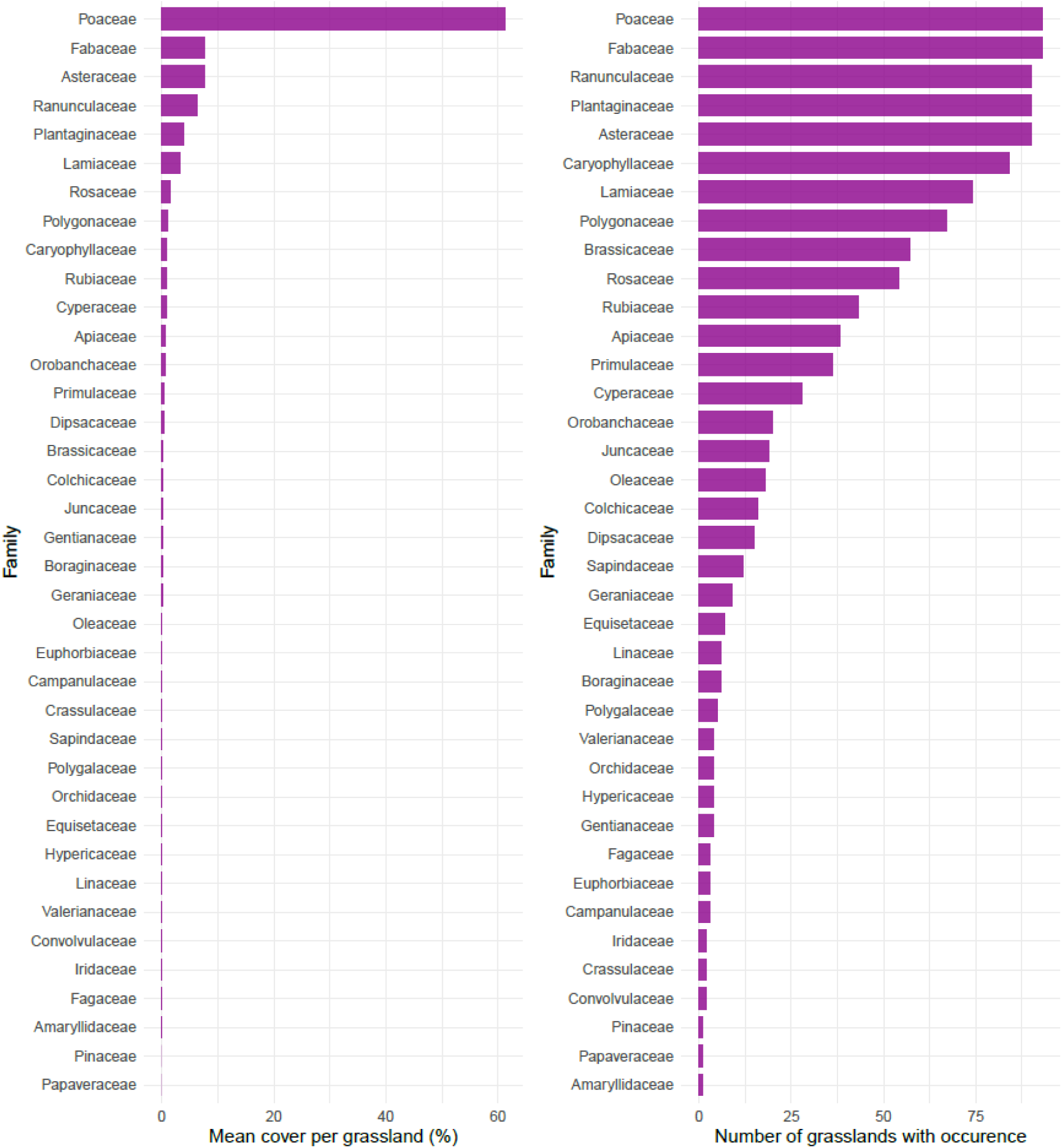
Mean cover of different plant families per grassland (left) and number of grasslands in which different plant families occurred (right) for the 92 grasslands from which the images for the image analysis were taken.

**Figure S5.**
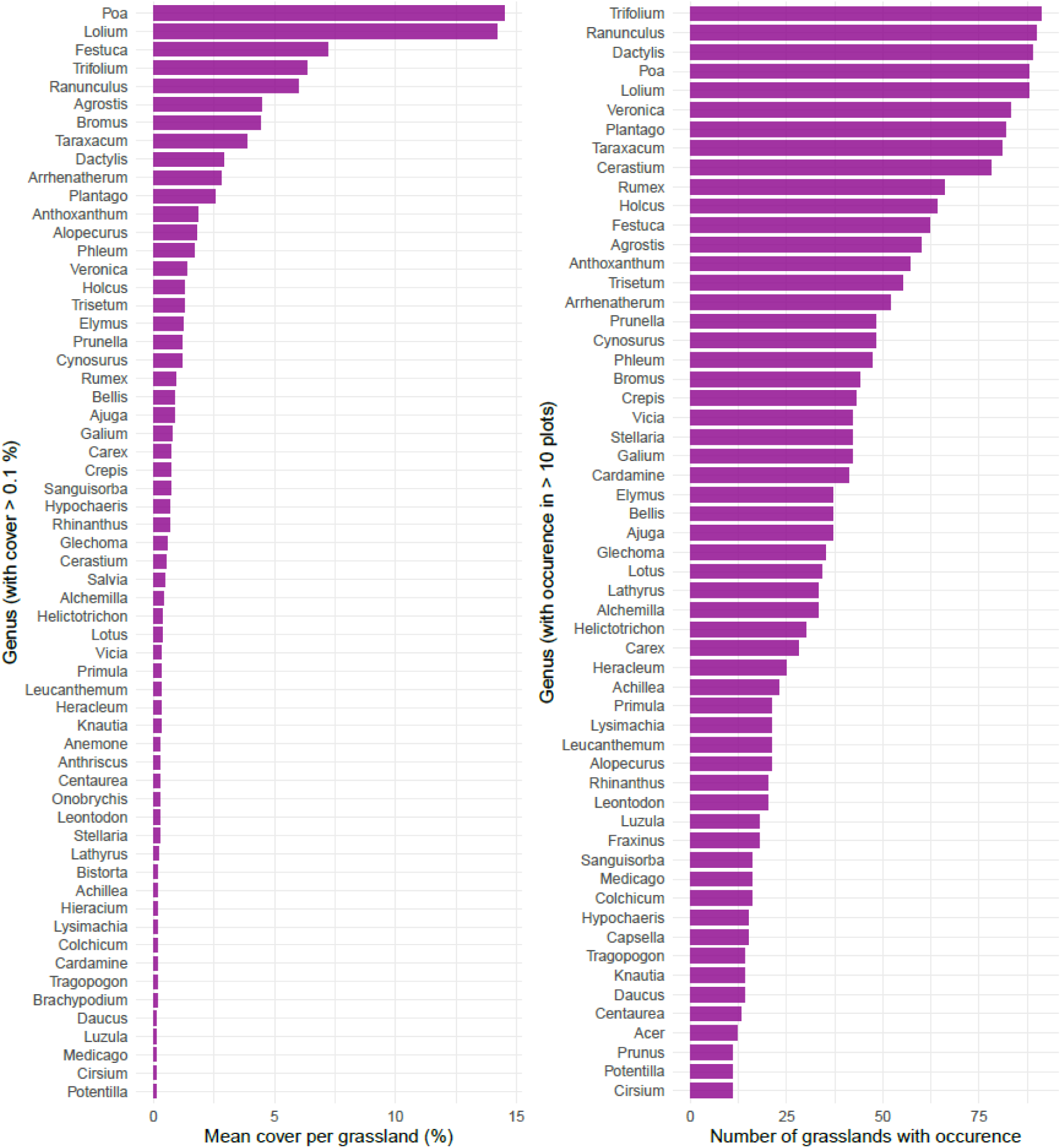
Mean cover of different plant genera per grassland (left) and number of grasslands in which different plant genera occurred (right) for the 92 grasslands from which the images for the image analysis were taken. Only genera with a cover > 0.1% and genera occurring in > 10 grasslands are shown, respectively.

**Table S1.** Basic statistics of responses given for all respondents and separated into three groups related to professional work. Responses with NA and mixed professions were not analysed separately (n = 62) but are included in all ratings.

|  | **All ratings (n = 522)** | **Ratings from agriculture (n = 121)** | **Ratings from ecology/nature conservation (n = 117)** | **Ratings from neither of both, other professions, retired (n = 222)** |
| --- | --- | --- | --- | --- |
| **Mean value** | 3.15 | 3.13 | 3.12 | 3.20 |
| **Standard deviation** | 0.65 | 0.65 | 0.82 | 0.70 |
| **Min** | 1.79 | 1.58 | 1.59 | 1.60 |
| **Max** | 4.54 | 4.50 | 4.80 | 4.57 |

**Table S2.** Results from ANOVAs relating the aesthetic rating (total and of different professional groups) to different plant community characteristics visible on the images (manually dummy coded). Significant levels: “***” = p < 0.001, “**” = p < 0.01, “*” = p < 0.05, “.” = p < 0.01 (all other p > 0.01).

| Ratings | Signs of defoliation (yes vs. no) |  |  |  |  |  | Bare soil visible (yes vs. no) |  |  |  |  |  |
| --- | --- | --- | --- | --- | --- | --- | --- | --- | --- | --- | --- | --- |
|  | Model p | Model R<sup>2</sup> | Estimate | SE | t value | Signif. | Model p | Model R<sup>2</sup> | Estimate | SE | t value | Signif. |
| All | < 0.001 | 0.33 | -1.12 | 0.17 | -6.71 | *** | < 0.001 | 0.17 | -0.61 | 0.14 | -4.34 | *** |
| Agriculture | < 0.001 | 0.23 | -0.93 | 0.18 | -5.21 | *** | < 0.001 | 0.28 | -0.77 | 0.13 | -5.93 | *** |
| Ecology | < 0.001 | 0.22 | -1.17 | 0.23 | -5.15 | *** | 0.021 | 0.06 | -0.44 | 0.19 | -2.35 | * |
| Other | < 0.001 | 0.36 | -1.26 | 0.18 | -7.16 | *** | < 0.001 | 0.15 | -0.61 | 0.15 | -4.00 | *** |
| Ratings | Weed prominently visible (yes vs. no) |  |  |  |  |  | Vegetation structure (light vs. dense) |  |  |  |  |  |
|  | Model p | Model R<sup>2</sup> | Estimate | SE | t value | Signif. | Model p | Model R<sup>2</sup> | Estimate | SE | t value | Signif. |
| All | 0.111 | 0.03 | 0.23 | 0.14 | 1.61 |  | 0.078 | 0.03 | 0.25 | 0.14 | 1.78 | . |
| Agriculture | 0.580 | 0.00 | -0.08 | 0.15 | -0.56 |  | 0.956 | 0.00 | -0.01 | 0.14 | -0.06 |  |
| Ecology | 0.007 | 0.08 | 0.49 | 0.18 | 2.77 | ** | < 0.001 | 0.12 | 0.61 | 0.17 | 3.60 | *** |
| Other | 0.021 | 0.06 | 0.36 | 0.15 | 2.34 | * | 0.138 | 0.02 | 0.23 | 0.15 | 1.50 |  |
| Ratings | Litter visible (yes vs. no) |  |  |  |  |  |  |  |  |  |  |  |
|  | Model p | Model R<sup>2</sup> | Estimate | SE | t value | Signif. |  |  |  |  |  |  |
| All | < 0.001 | 0.14 | -0.93 | 0.24 | -3.90 | *** |  |  |  |  |  |  |
| Agriculture | < 0.001 | 0.20 | -1.12 | 0.23 | -4.85 | *** |  |  |  |  |  |  |
| Ecology | 0.196 | 0.02 | -0.42 | 0.32 | -1.30 |  |  |  |  |  |  |  |
| Other | < 0.001 | 0.17 | -1.11 | 0.25 | -4.36 | *** |  |  |  |  |  |  |

**Table S3.** Overview of the labels generated by the feature “label” of the Google Cloud Vision algorithm for Flickr and iNaturalist photographs and their classification. This classification was adapted from Chai-allah et al. (submitted). Only labels with frequency > 15 were included.

| Category | Sub-category | Labels |
| --- | --- | --- |
| Nature | Landscape | aeolian landform, agriculture, arroyo, badlands, batholith, beach, bedrock, biome, body of water, cave, chaparral, coast, coastal and oceanic landforms, creek, crop, deciduous, ecoregion, escarpment, fell, fen, field, fluid, fluvial landforms of streams, forest, freshwater marsh, garden, geological phenomenon, geology, glacial landform, glacier, grassland, hay, headland, highland, hill, hinterland, ice cap, jungle, lacustrine plain, lake, landscape, limestone, loch, logging, marsh, massif, meadow, monolith, mountain, mountain range, mountain river, mountainous landforms, natural environment, natural landscape, natural material, nature, nature reserve, northern hardwood forest, ocean, old-growth forest, outcrop, pasture, pebble, plain, plant community, plateau, pond, prairie, reservoir, ridge, riparian forest, riparian zone, river island, rock, rural area, salt marsh, sand, seep, shore, shrubland, slope, snow, soil, speleothem, spruce-fir forest, stalactite, stalagmite, steppe, stream, summit, tarn, temperate broadleaf and mixed forest, temperate coniferous forest, terrain, thatching, tidal marsh, tropical and subtropical coniferous forests, underwater, valley, water, water feature, water resources, watercourse, waterfall, waterway, wave, wetland, wilderness, woodland |
|  | Objects/species | accipitridae, accipitriformes, acmon blue, adonis blue, agaric, agaricaceae, agaricomycetes, aglais, aglais io, algae, allium, alpaca, american black bear, american black duck, american coot, amphibian, anaxyrus, animal migration, annual plant, ant, antelope, anthriscus, antler, apatura, aphantopus, aquatic plant, arachnid, araneus, argiope, argynnis, aricia, arthropod, bald eagle, band winged grasshoppers, beak, bear, bee, bee balm, beetle, berry, bird, bird migration, bird nest, bird of prey, bison, blossom, blowflies, bolete, boloria, bombyliidae, botany, bovine, branch, brimstones, brown argus, brown bear, bud, bufo, bug, bull, bullfrog, bumblebee, bumblebee orchid, burdock, buttercup, butterfly, buzzard, cabbage butterfly, calf, calidrid, camomile, caterpillar, celastrina, chamaemelum nobile, charadriiformes, cherry blossom, chickadee, chicken, ciconiiformes, claw, cleavers, coenagrion, coenonympha tullia, colias, colias croceus, colias hyale, colias sareptensis, coltsfoot, comb, comma, common blue, condor, conifer, corn poppy, cow-goat family, crane-like bird, creeping thistle, crepis paludosa, cretan crocus, cricket-like insect, crow, crow-like bird, cupido (butterfly), cut flowers, dactylorhiza praetermissa, dairy cow, daisy, daisy family, damselfly, dandelion, deer, distaff thistles, domestic pig, dowitcher, dragon lizard, dragonflies and |
|  |  | damseflies, dragonfly, drosophila, duck, ducks, geese and swans, dung beetle, eagle, eastern tent caterpillar, edible mushroom, egret, elapidae, elk, emberizidae, european marsh thistle, european robin, evergreen, falcon, falconiformes, fawn, feather, felidae, feral goat, fern, ferns and horsetails, fin, finch, fire salamander, fish crow, flightless bird, flock, floribunda, flower, flowering plant, fly orchid, forb, forget-me-not, fowl, fox, french lavender, fritillaria, frog, fruit, fungus, fur, galliformes, garden white, gatekeeper (butterfly), gentiana, goat, goat-antelope, goats, gonepteryx cleopatra, gonepteryx rhamni, grapevine family, grass, grass family, grass snake, grasshopper, grazing, great blue heron, great heron, great spangled fritillary, greater scaup, green-veined white, grizzly bear, ground beetle, groundcover, grove, gull, halictidae, hare, harrier, hawk, hawker dragonflies, heath fritillary, hedge, hemp family, hen-of-the-wood, herb, herbaceous plant, herd, heron, hierochloe, high brown fritillary, hipparchia, holly blue, honeybee, horse, horse flies, house fly, hybrid clover, hybrid tea rose, hypericum, iguana, inonotus, insect, invertebrate, khorasan wheat, kingcup, kit fox, ladybug, larch, large skipper, large tortoiseshell, large white, lari, larva, lasiommata, leaf, leafhopper, leptidea, lesser yellowlegs, livestock, lizard, llama, locust, lotus, lotus family, lycaena, lymantriidae, lymnaeidae, lynx, malus, mammal, mane, maniola, mantidae, mantis, mare, marguerite daisy, marine biology, marine invertebrates, marine mammal, marsupial, mayflies, mazarine blue, meadow brown, medicinal mushroom, megachilidae, melanargia, melanargia galathea, melitaea, membrane-winged insect, mole salamander, morning glory, moth, moths and butterflies, mountain cottontail, mushroom, mustelidae, narcissus, natural foods, neotinea ustulata, nest, net-winged insects, new caledonian crow, nightshade family, non-vascular land plant, northern brown argus, northern leopard frog, noxious weed, nymphalis, nymphalis xanthomelas, ochlodes, ophrys, orange sulphur, orb-weaver spider, orchids of the philippines, organism, oriental poppy, osprey, ostrich fern, owl, ox, oxeye daisy, pack animal, papilio, papilio machaon, pararge, parasite, peafowl, pearl-bordered fritillar, pedicel, pelecaniformes, peppered moth, perching bird, perennial plant, pest, petal, phasianidae, piciformes, pincushion flower, plant, plant pathology, plant stem, plebejus, poales, pollen, pollinator, polygonia, polyommatus, polyporales, poultry, produce, pyronia cecilia, rabbit, rabbits and hares, rallidae, raven, red bugs, red fox, reef, rein, reindeer, reptile, rodent, rosa × centifolia, rosa arkansana, rosa canina, rose, rose family, rubus, russula integra, saffron crocus, salamander, salamandra, sandpiper, scaled reptile, scentless plant bugs, sea eagle, sea snail, seabird, seaduck, sharp shinned hawk, sheep, shell, shield bugs, shortstraw pine, shrike, shrub, silver-studded blue, silver-washed fritillary, silvery blue, skink, skipper (butterfly), small blue, small heath (butterfly), small pearl-bordered butterfly, small tortoiseshell, snail, snails and slugs, snout, snow crocus, soldier beetle, songbird, sorrel, southern dogface, southern |
|  |  | leopard frog, sow thistles, sparrow, spear thistle, spider, spider web, spotted salamander, spring crocus, squirrel, stallion, stock dove, subshrub, suidae, sunflower, swallowtail butterfly, swan, sweet grass, sweet peas, swift fox, tangle-web spider, teasel, terrestrial animal, terrestrial plant, thistle, thorns, spines, and prickles, thymelicus, tiger salamander, toad, tommie crocus, tortoise, tree, true frog, true salamanders and newts, true toad, trunk, tundra swan, turtle, twig, underwing moths, valerian, vanessa (butterfly), vascular plant, vegetable, vegetation, vertebrate, vicuña, vitis, vulture, wall lizard, wasp, water bird, water lily, waterfowl, weevil, western tiger swallowtail, wheat, whiptail, white-tailed deer, white stork, wild carrot, wildflower, wildlife, wing, wolf, wolf spider, wood, woodpecker, woody plant, working animal |
| Non-nature | Non-nature | abdomen, accessory fruit, accordion, accordionist, acoustic guitar, acting, active shirt, active shorts, adaptation, adventure, advertising, aerial photography, aerobatics, aerospace engineering, aerospace manufacturer, aerostat, afterglow, agricultural machinery, air racing, air show, air sports, air travel, aircraft, aircraft engine, airline, airliner, airplane, airport, airport apron, airship, alcohol, alcoholic beverage, alley, alligator, alloy wheel, aluminium, amber, american alligator, american crocodile, american crow, ancient history, animal product, animal shelter, animal sports, antique, antique car, apartment, aqua, aquarium, aqueduct, arbor day, arcade, arch, arch bridge, archaeological site, architecture, arctic, arctic ocean, arecales, arête, arm, armored car, army, art, art paint, artifact, artificial flower, artist, artwork, ash, asphalt, astronomical object, astronomy, athlete, athletic shoe, athletics, atlas, atmosphere, atmospheric phenomenon, audience, audio equipment, auto part, auto racing, automotive carrying rack, automotive decal, automotive design, automotive exhaust, automotive exterior, automotive fog light, automotive fuel system, automotive light bulb, automotive lighting, automotive mirror, automotive parking light, automotive side-view mirror, automotive side marker light, automotive tail & brake light, automotive tire, automotive wheel system, automotive window part, aviation, azure, baby, baby & toddler clothing, baby carriage, baby products, backlighting, backpack, bag, baked goods, baking, balance, ball, ball game, balloon, baluster, band plays, bangs, bank, banner, barbed wire, barechested, barefoot, barware, baseball, baseball cap, baseball equipment, baseball field, baseball glove, baseball park, baseball player, baseball protective gear, baseball uniform, basket, bassist, bat-and-ball games, bathroom, batting helmet, beacon, beam, beam bridge, beard, beauty, beef, beer, beer glass, beige, belt, bench, bengal tiger, bermuda shorts, bicycle, bicycle accessory, bicycle chain, bicycle clothing, bicycle fork, bicycle frame, bicycle handlebar, bicycle helmet, bicycle hub, bicycle jersey, bicycle part, bicycle saddle, bicycle seatpost, bicycle stem, bicycle tire, bicycle wheel, bicycle wheel rim, bicycles--equipment and supplies, big cats, billboard, biplane, bird's-eye view, bit, black, black-and-white, |
|  |  | black hair, blazer, blimp, block party, blond, blue, blue-collar worker, board game, boardsport, boardwalk, boat, boats and boating--equipment and supplies, body jewelry, bonfire, book, boot, bottle, bouquet, bowl, box, box girder bridge, brand, brass instrument, brassiere, bread, brick, brickwork, bridal accessory, bridal clothing, bridal party dress, bride, bridge, bridle, bronze, bronze sculpture, brown, brown hair, building, building material, bulldozer, bullet train, bumper, bus, business jet, button, byzantine architecture, cabinetry, cable, cable-stayed bridge, cable car, cage, cake, calabaza, calm, camelid, camera, camera accessory, camera lens, cameras & optics, camouflage, campfire, camping, canal, candle, canopy, canopy walkway, cap, cape, car, car seat, car seat cover, carbon, carbonated soft drinks, cargo aircraft, cargo pants, carmine, carriage, cart, carving, cash crop, castle, ceiling, ceiling fixture, celebrating, celestial event, cello, cemetery, center console, ceramic, cg artwork, chain-link fencing, chair, champagne stemware, championship, chandelier, channel, chapel, charcoal, cheek, cheetah, chemical compound, chess, chessboard, chest, child, chimney, chin, chinese architecture, choreography, christmas, christmas decoration, christmas ornament, christmas tree, church, chute, circle, citrus, city, cityscape, classic, classic car, classical architecture, classical music, classical sculpture, cleanliness, cleat, clock, close-up, clothing, cloud, coat, cobblestone, cockpit, cocktail, coffee cup, coffee table, collaboration, collar, college baseball, college softball, colorfulness, colubridae, column, combat sport, combat vehicle, comfort, comfort food, commemorative plaque, commercial building, commercial vehicle, communication device, community, compact car, compact van, companion dog, competition, competition event, composite material, compost, computer, computer hardware, computer keyboard, computer monitor, concert, concrete, concrete bridge, condominium, cone, confectionery, constellation, construction, construction equipment, construction worker, contact sport, conversation, cooking, cookware and bakeware, cool, corn on the cob, costume, costume hat, cottage, couch, countertop, cowboy hat, crane, crankset, creative arts, crescent, crew, cricket, cricket cap, crocodile, crocodilia, croft, cross, cross-country equestrianism, cross-country skiing, crowd, cuisine, culinary art, cumulus, cup, cupboard, curtain, customer, cutlery, cycle sport, cycling, cycling shoe, cycling shorts, cylinder, cymbal, cynara, dairy, dance, darkness, dawn, daylighting, daytime, decoration, deejay, delicacy, denim, design, desk, dessert, devil's bridge, dew, diagram, digital camera, digital slr, dirt road, dirt track racing, disc dog, disc jockey, dish, dishware, display device, distilled beverage, ditch, diwali, dock, document, dog agility, dog collar, dog sports, dog supply, dome, door, door handle, downhill ski binding, downtown, drawer, drawing, dress, dress shirt, drink, drinking, drinking establishment, drinking water, drinkware, driveway, drizzle, drone, drop, drum, drum stick, drumhead, drummer, drums, dusk, ear, earrings, eclipse, elbow, electric blue, electric guitar, electric locomotive, electrical supply, |
|  |  | electrical wiring, electricity, electronic device, electronic instrument, electronic keyboard, electronic musical instrument, elephants and mammoths, emblem, emergency service, emergency vehicle, employment, endurance sports, enduro, engineer, engineering, english pleasure, english riding, entertainment, environmental art, equestrian, equestrian helmet, equestrian sport, equestrianism, equitation, erosion, evening, event, eventing, exercise, exhaust system, extreme sport, eye, eye glass accessory, eyebrow, eyelash, eyewear, facade, face, facial expression, facial hair, factory, family car, fan, farmer, farmhouse, farmworker, fashion, fashion accessory, fashion design, fast food, fault, fedora, fence, fender, festival, fictional character, fiddle, fighter aircraft, fighter pilot, film camera, fines herbes, finger, finger food, finial, fire, firefighter, fireplace, fireworks, fixed link, fixture, flag, flag of the united states, flagstone, flame, flap, flash photography, flat panel display, flat racing, flesh, flight, floor, flooring, floral design, floristry, flower arranging, flowerpot, fly, flying boat, flying disc, flying disc freestyle, fog, folding chair, folk instrument, folk wrestling, font, food, food craving, foot, football, football equipment, football player, footwear, forehead, fork, formal wear, formation, fountain, fourball, free reed aerophone, freeway, freezing, freight car, fried food, friendship, frisbee games, frost, fuel tank, full-size car, full moon, fun, function hall, furniture, gadget, galaxy, games, garden buildings, garden roses, garden tool, gardening, garnish, gas, gauge, gazebo, general aviation, gesture, girder bridge, glass, glass bottle, glasses, glider, gliding, glove, gluten, goal, goggles, gold, golf, golf bag, golf ball, golf cart, golf club, golf course, golf equipment, golfer, gothic architecture, government agency, gown, graffiti, grand prix motorcycle racing, graphic design, graphics, grappling, grave, green, grey, grille, grilling, ground attack aircraft, group a, groupset, guard rail, guitar, guitar accessory, guitarist, hair, hairstyle, hall, halter, hand, handbag, handle, handrail, handwriting, hangar, happy, harbor, hard hat, hardtop, hardwood, harvester, hat, haze, head, head restraint, headgear, headlamp, headpiece, headquarters, headstone, hearing, heat, helicopter, helicopter rotor, helmet, high-speed rail, high-visibility clothing, highway, hiking, hiking equipment, hill station, historic site, history, holiday, holy places, home, home appliance, home door, home fencing, hood, horizon, horn, horse and buggy, horse supplies, horse tack, horse trailer, hot air balloon, hot air ballooning, house, household hardware, houseplant, hub gear, hubcap, human, human body, human leg, human settlement, hurdle, hut, idiophone, illustration, individual sports, indoor games and sports, industry, infantry, inflatable, infrastructure, ingredient, interaction, interior design, intersection, intrusion, iris, iron, island, isle of man tt, jacket, japanese architecture, jaw, jazz pianist, jeans, jersey, jet aircraft, jet engine, jewellery, job, jockey, joint, jumping, junk food, kangaroo, keyboard, keyboard player, kick, kit car, kitchen, kitchen & dining room table, kitchen appliance, kitchen stove, kitchen utensil, kite, kite sports, klippe, |
|  |  | knee, knee pad, knife, knit cap, kodiak bear, ladder, lake district, laminate flooring, lamp, land lot, land vehicle, landing, landmark, landscape lighting, landscaping, lane, lap, laptop, laugh, lavender, law enforcement, lawn, lawn ornament, layered hair, leash, leather jacket, lectern, leg, leisure, lens, lens flare, leopard, light, light aircraft, light commercial vehicle, light fixture, lighthouse, lighting, lightning, line, line art, linens, lion, lip, liqueur, liquid, liver, living room, local food, lockheed martin, locking hubs, locomotive, log cabin, logo, long hair, luggage and bags, lumber, lunar eclipse, luxury vehicle, machine, macro photography, magenta, major appliance, makeover, manor house, map, marathon, marines, market, marketplace, masai lion, mast, match play, material property, measuring instrument, meat, mechanical fan, medieval architecture, meeting, megalith, melting, membranophone, memorial, mesh, metal, meteorological phenomenon, metropolis, metropolitan area, microphone, microphone stand, mid-size car, midnight, military, military aircraft, military camouflage, military helicopter, military officer, military organization, military person, military uniform, military vehicle, milky way, mirror, mirrorless interchangeable-lens camera, mist, mixed-use, mixing bowl, mixture, mobile phone, mode of transport, model aircraft, moisture, molding, monochrome, monochrome photography, monoplane, monument, moon, moonlight, morning, motocross, motor glider, motor vehicle, motorcycle, motorcycle accessories, motorcycle fairing, motorcycle helmet, motorcycle racer, motorcycle racing, motorcycling, motorsport, mountaineer, moustache, mouth, multi-sport event, multimedia, mural, muscle, music, music artist, music stand, music venue, musical ensemble, musical instrument, musical instrument accessory, musical keyboard, musician, mythology, nail, narrow-body aircraft, naval architecture, nebula, neck, necklace, neighbourhood, neon, net, net sports, new year, new year's eve, new years day, nickel, night, non-commissioned officer, nonbuilding structure, northern harrier, nose, notchback, number, nunatak, oar, observation tower, off-road racing, office equipment, office ruler, office supplies, official, one-piece garment, open-wheel car, optical instrument, orange, orator, organ, ornament, outdoor bench, outdoor furniture, outdoor play equipment, outdoor recreation, outdoor shoe, outdoor structure, outdoor table, outer space, outerwear, output device, overhead power line, overpass, paddle, paint, painting, palm tree, paper, paper product, parachute, parachuting, paragliding, parallel, parish, parking, party, party supply, passenger, passenger car, passenger ship, path, pattern, paw, pedestrian, people, people in nature, people on beach, percussion, percussion mallet, percussionist, performance, performance art, performance car, performing arts, performing arts center, peripheral, personal computer, personal luxury car, personal protective equipment, pet supply, photograph, photographer, physical fitness, pianist, piano, picnic table, picture frame, pier, pilot, pink, pink family, pint, pipe, piste, place of worship, plaid, |
|  |  | planet, plank, plantation, plaster, plastic, plastic bottle, plate, platter, play, player, playground, playing in the snow, playing sports, plucked string instruments, plumbing, plumbing fixture, plywood, podium, point-and-shoot camera, polar ice cap, pole, pollution, polo shirt, pop music, porcelain, porch, pork, portrait photography, poster, pottery, precipitation, precision sports, presentation, primate, product, professional golfer, projection screen, projector accessory, propeller, propeller-driven aircraft, property, public address system, public event, public space, public speaking, public transport, public utility, publication, purple, pyramid, quadrathlon, race car, race track, racing, radiator, radio-controlled aircraft, radio-controlled helicopter, radio-controlled toy, railroad car, railway, rainbow, rallycross, rallying, ranch, rangpur, rapid, real estate, rear-view mirror, recipe, recital, recreation, rectangle, red, red meat, red sky at morning, reed, reed instrument, reflection, reflex camera, relief, religious institute, religious item, residential area, resort town, retail, rice, riding boot, rim, road, road bicycle, road surface, roadster, rock-climbing equipment, rock concert, rolling, rolling stock, romance, roof, room, rope, rose order, rotor, rotorcraft, rubble, rugby, rugby short, ruins, ruler, running, runway, sacred lotus, saddle, sail, sailboat, sailing, saltwater crocodile, sandal, sash window, saxophone, saxophonist, scarf, science, scrap, screenshot, sculpture, security, selfie, serpent, serveware, service, serving tray, shack, shade, shadow, sharing, shed, shelf, shelving, shepherd, shin guard, ship, shirt, shoe, shorts, shoulder, show jumping, siberian tiger, sidewalk, siding, sign, sign language, signage, singer, singing, single-lens reflex camera, sink, sitting, skateboard deck, sketch, ski, ski binding, ski boot, ski equipment, ski helmet, ski pole, skier, skiff, skiing, skin, sky, skyscraper, sled, sledding, sleeve, slug, smile, smoke, snake, snapshot, sneakers, snow leopard, snowboard, snowboarding, soccer, soccer ball, soccer kick, soccer player, social group, sock, soft drink, solar energy, soldier, solution, solvent, song, sound, space, spandex, speaker, spectacle, speech, speed golf, speedometer, sphere, spire, splint boots, split-rail fence, spoke, spokesperson, spoon, sport utility vehicle, sport venue, sporting group, sports, sports car, sports car racing, sports equipment, sports gear, sports jersey, sports toy, sports uniform, sportswear, spring, square, squash, squeezebox, stable, stadium, stage, stage equipment, stained glass, stairs, standing, staple food, star, stately home, statue, steak, steam, steam engine, steel, steeple, steeplechase, steering part, steering wheel, stemware, step cutting, stick and ball sports, still life photography, stock photography, stone carving, stone wall, storm, stove, street, street art, street furniture, street light, street sign, string instrument, string instrument accessory, stunt, stunt performer, style, suburb, sugar house, suit, summer, sun, sun hat, sunglasses, sunlight, sunrise, sunset, superbike racing, superfood, supermoto, supersonic aircraft, suspension bridge, sweatpant, sweetness, swimming pool, swimwear, symbol, symmetry, synthetic rubber, t-shirt, table, tablecloth, tabletop |
|  |  | game, tableware, tachometer, tail, takeoff, tap, tar, tarpaulin, tartan, team, team sport, technology, telecommunications engineering, television, temperature, temple, tent, terrace, textile, theater curtain, theatre, thigh, thoroughfare, throat, thumb, thunder, thunderstorm, tie, tied-arch bridge, tiger, tile flooring, tints and shades, tire, tire care, titanium, toddler, tomtom drum, tool, tooth, tour bus service, touring car racing, tourism, tournament, tower, tower block, town, toy, toy airplane, toy dog, toy vehicle, track, track racing, tractor, tradesman, tradition, traffic sign, trail, trailer, train, train station, trainer, tram, transmission tower, transmitter station, transparency, transparent material, transport hub, travel, travel trailer, tread, triangle, tripod, tropics, trousers, truck, trunks, truss bridge, tunnel, turbine, turboprop, turret, twinjet, ultimate, ultralight aviation, ultramarathon, umbrella, uniform, universe, urban area, urban design, urtica, vacation, van, varnish, vault, vegan nutrition, vehicle, vehicle brake, vehicle door, vehicle registration plate, vest, viaduct, village, vintage car, vintage clothing, viola, violet, violin, violin family, violist, violone, vision care, visual arts, visual effect lighting, volleyball, wagon, waist, walking, walking shoe, walkway, wall, waste, waste container, waste containment, watch, water bottle, water transportation, watercolor paint, watercraft, wedding ceremony supply, wedding dress, wedge, western riding, wheel, whiskers, white, white-collar worker, whole food, whole grain, wide-body aircraft, wind, wind farm, wind instrument, wind turbine, wind wave, windbreaker, windmill, window, window treatment, windscreen wiper, windshield, windsports, wine, wine bottle, wine glass, winter, winter sport, wire, wire fencing, women's football, wood flooring, wood stain, woodwind instrument, woolen, workwear, world, world rally car, woven fabric, wrestling, wrinkle, wrist, writing, yellow, youth, zeppelin |

**Table S4.**
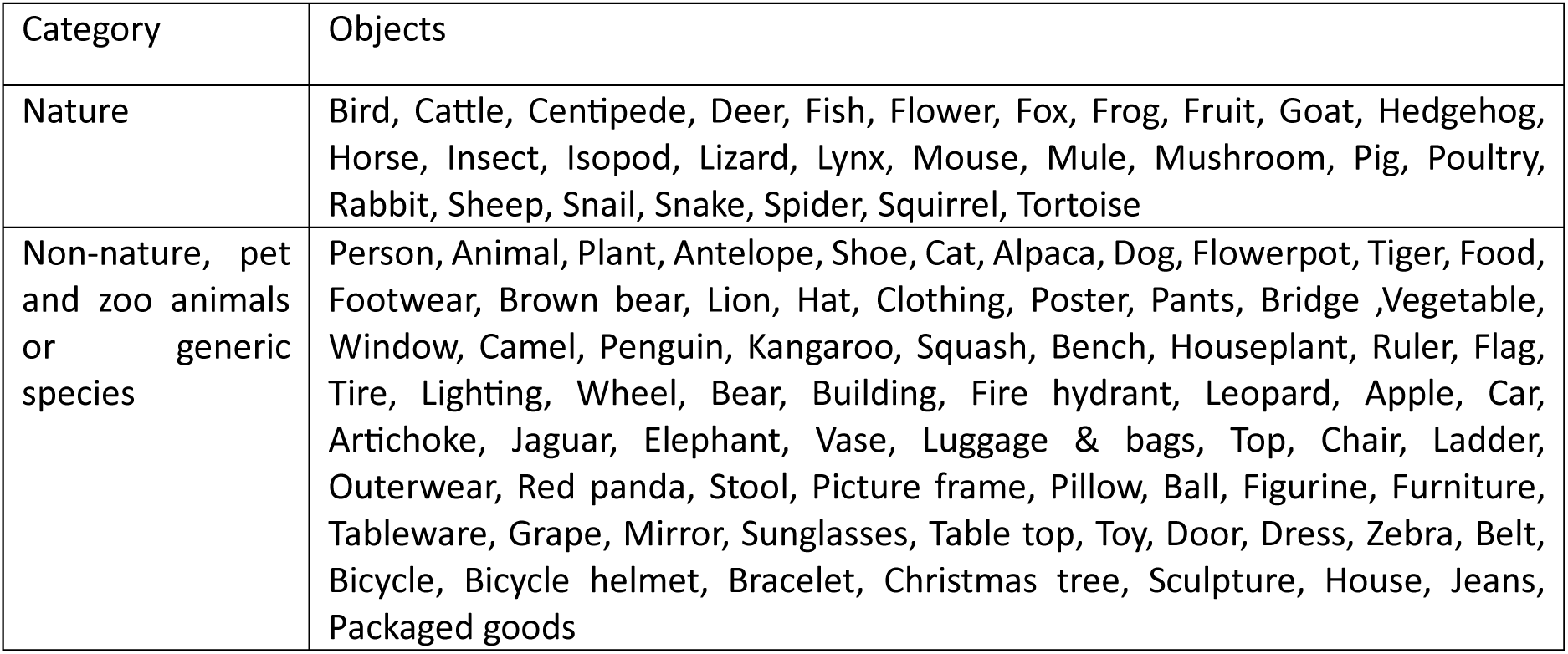
Overview of the extracted objects using the feature “object” of the Google Cloud Vision algorithm for Flickr and iNaturalist photographs of species. Only objects with frequency > 1 were included.

| Category | Objects |
| --- | --- |
| Nature | Bird, Cattle, Centipede, Deer, Fish, Flower, Fox, Frog, Fruit, Goat, Hedgehog, Horse, Insect, Isopod, Lizard, Lynx, Mouse, Mule, Mushroom, Pig, Poultry, Rabbit, Sheep, Snail, Snake, Spider, Squirrel, Tortoise |
| Non-nature, pet and zoo animals or generic species | Person, Animal, Plant, Antelope, Shoe, Cat, Alpaca, Dog, Flowerpot, Tiger, Food, Footwear, Brown bear, Lion, Hat, Clothing, Poster, Pants, Bridge ,Vegetable, Window, Camel, Penguin, Kangaroo, Squash, Bench, Houseplant, Ruler, Flag, Tire, Lighting, Wheel, Bear, Building, Fire hydrant, Leopard, Apple, Car, Artichoke, Jaguar, Elephant, Vase, Luggage & bags, Top, Chair, Ladder, Outerwear, Red panda, Stool, Picture frame, Pillow, Ball, Figurine, Furniture, Tableware, Grape, Mirror, Sunglasses, Table top, Toy, Door, Dress, Zebra, Belt, Bicycle, Bicycle helmet, Bracelet, Christmas tree, Sculpture, House, Jeans, Packaged goods |

**Table. S5.** Summary of the top 10 most frequently photographed objects, their distribution per user and their seasonal distribution.

| Platform | Species | Number of images | Number of users | Images per user (mean) | Seasonal distribution (N° of images) |  |  |  |
| --- | --- | --- | --- | --- | --- | --- | --- | --- |
|  |  |  |  |  | Spring | Summer | Autumn | Winter |
| Flickr | Flower | 2198 | 430 | 5,11 | 977 | 729 | 393 | 99 |
|  | Insect | 612 | 147 | 4,16 | 164 | 331 | 111 | 6 |
|  | Bird | 510 | 141 | 3,61 | 191 | 163 | 103 | 53 |
|  | Cattle | 484 | 178 | 2,71 | 134 | 154 | 182 | 14 |
|  | Horse | 352 | 65 | 5,41 | 98 | 194 | 41 | 19 |
|  | Goat | 118 | 50 | 2,36 | 29 | 27 | 40 | 22 |
|  | Sheep | 116 | 50 | 2,32 | 21 | 20 | 35 | 40 |
|  | Mushroom | 80 | 52 | 1,53 | 13 | 8 | 45 | 14 |
|  | Deer | 67 | 39 | 1,71 | 11 | 14 | 33 | 9 |
|  | Frog | 61 | 28 | 2,17 | 27 | 30 | 4 | 0 |
| iNaturalist | Flower | 2500 | 263 | 9,5 | 1105 | 981 | 383 | 31 |
|  | Insect | 1230 | 157 | 7,83 | 450 | 611 | 156 | 13 |
|  | Bird | 439 | 78 | 5,62 | 154 | 108 | 97 | 80 |
|  | Mushroom | 260 | 40 | 6,5 | 34 | 23 | 197 | 6 |
|  | Deer | 46 | 15 | 3,06 | 16 | 7 | 11 | 12 |
|  | Frog | 43 | 29 | 1,48 | 25 | 10 | 8 | 0 |
|  | Lizard | 41 | 33 | 1,24 | 19 | 13 | 8 | 1 |
|  | Spider | 33 | 19 | 1,73 | 6 | 21 | 6 | 0 |
|  | Snail | 32 | 22 | 1,45 | 16 | 16 | 0 | 0 |
|  | Fruit | 18 | 11 | 1,63 | 1 | 9 | 8 | 0 |

**Table S6.**
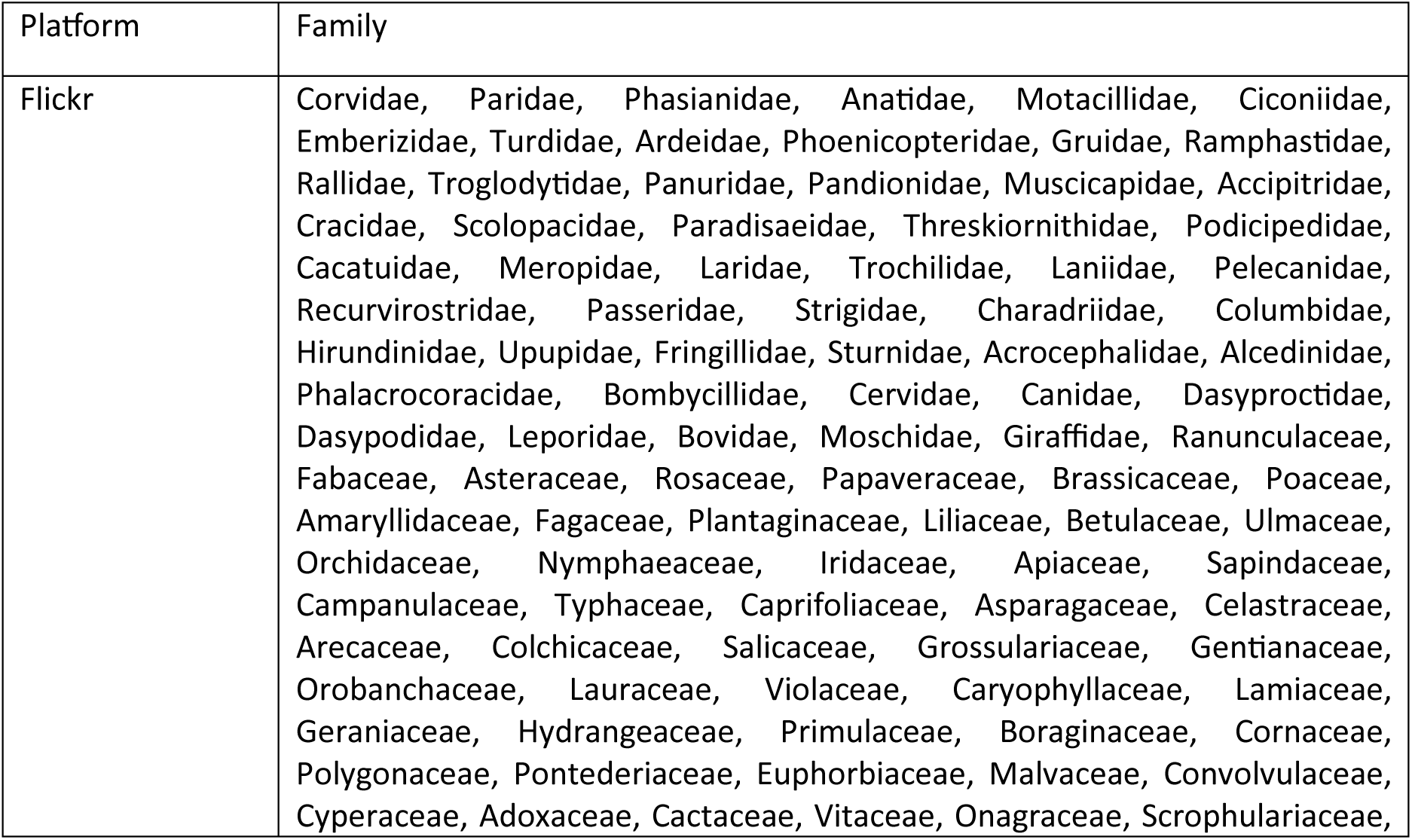
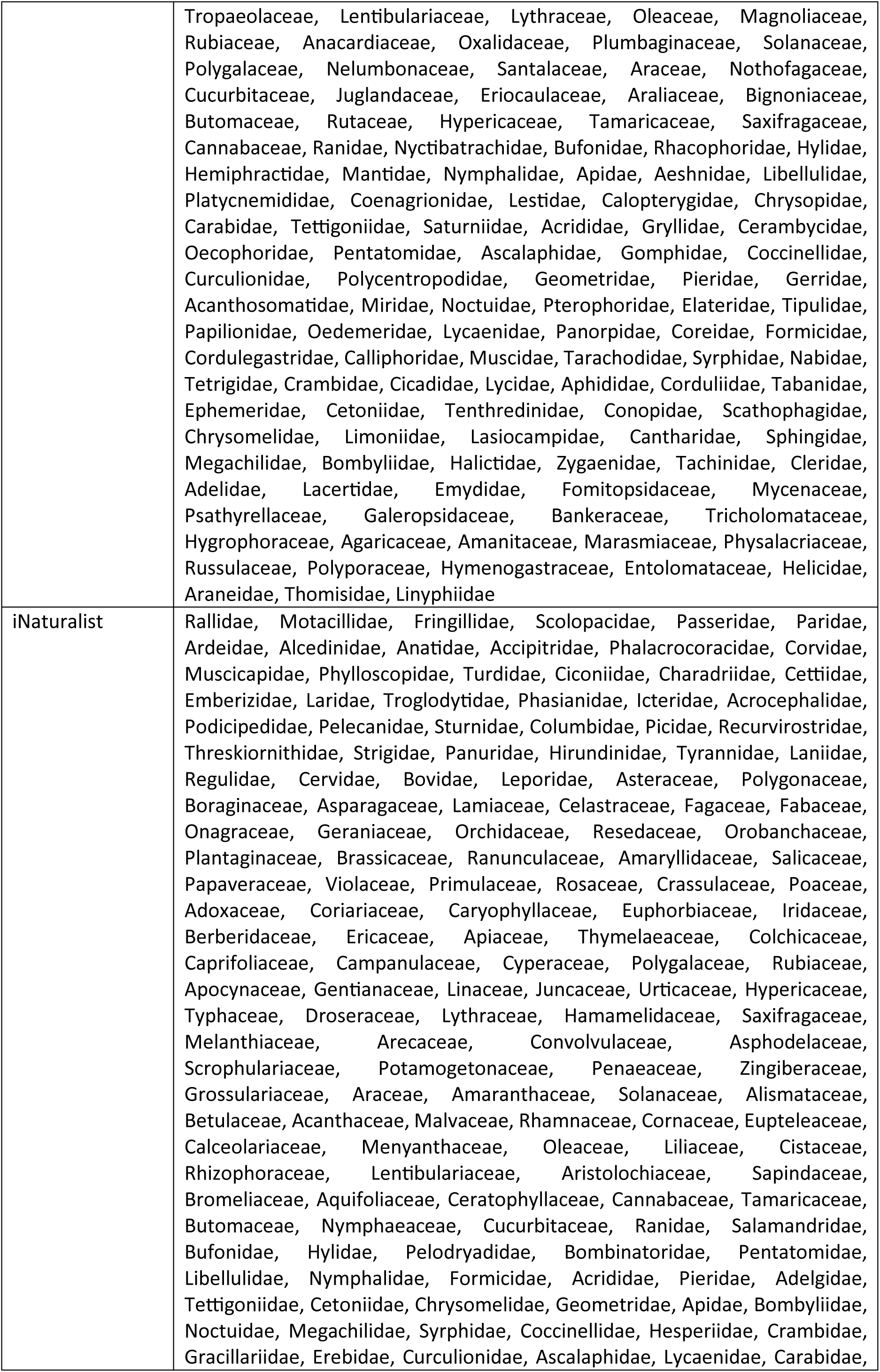
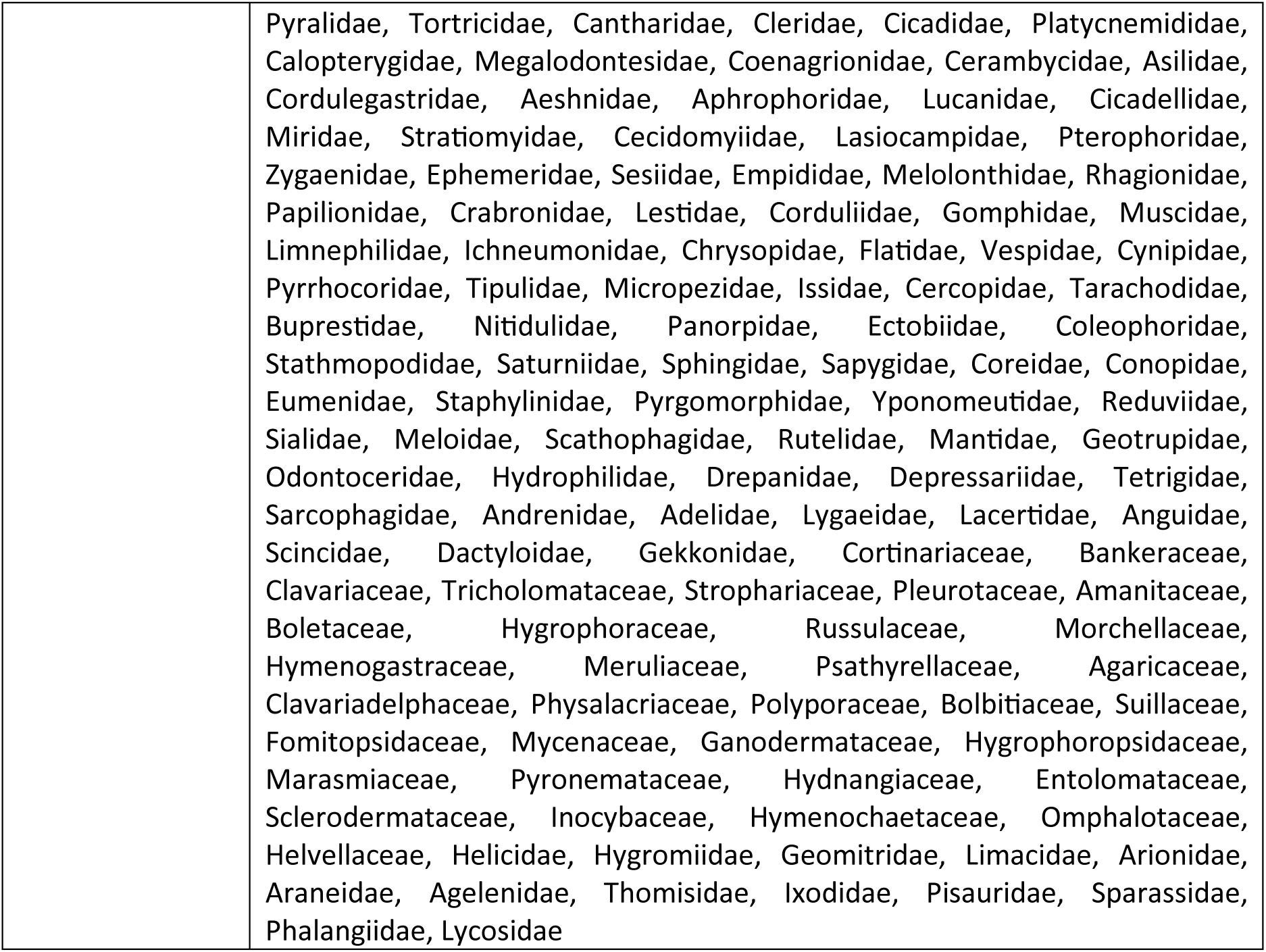
Overview of the extracted families using the BioCLIP algorithm for Flickr and iNaturalist photographs of the most photographed wildlife species.

**Table S7.**
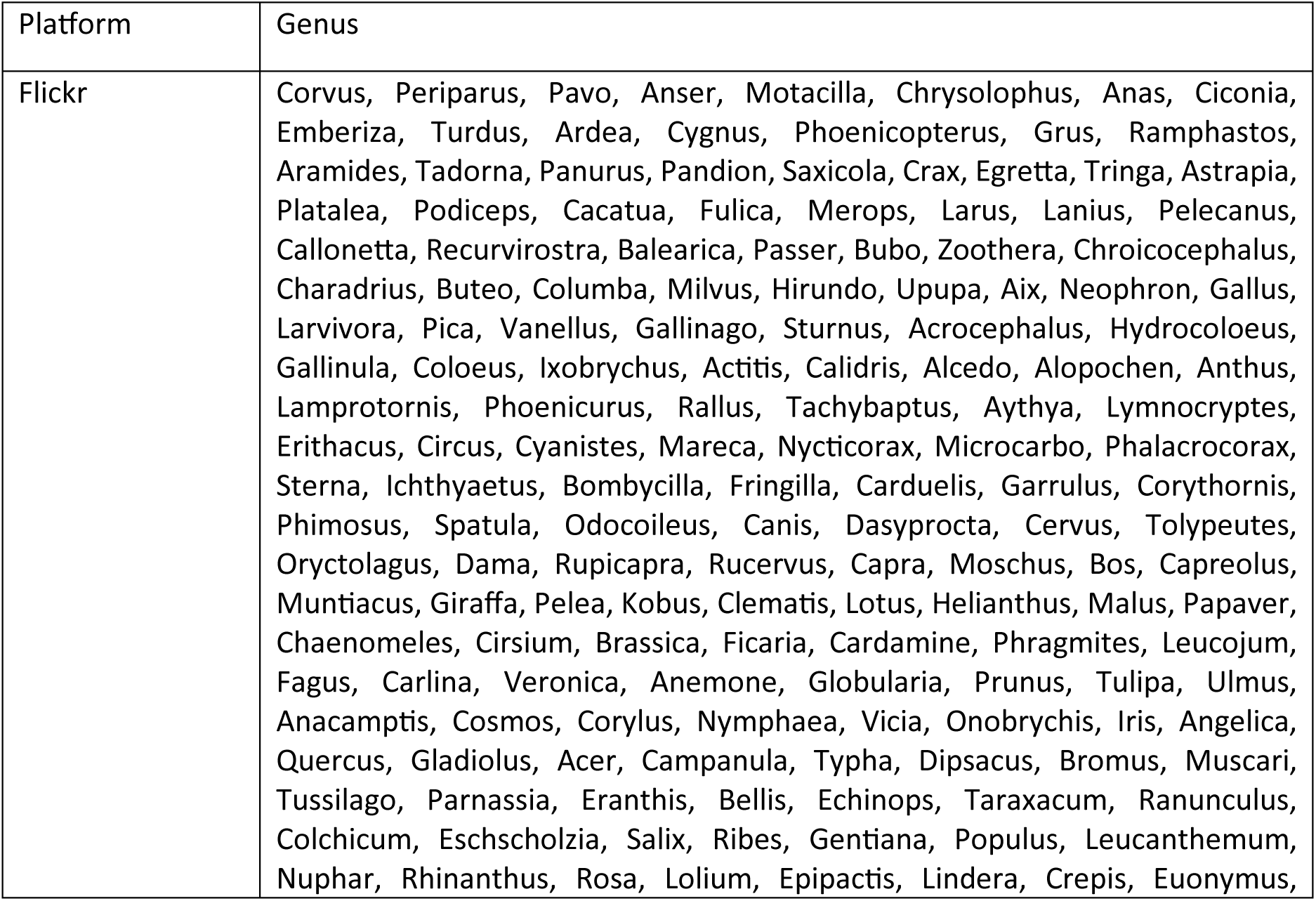
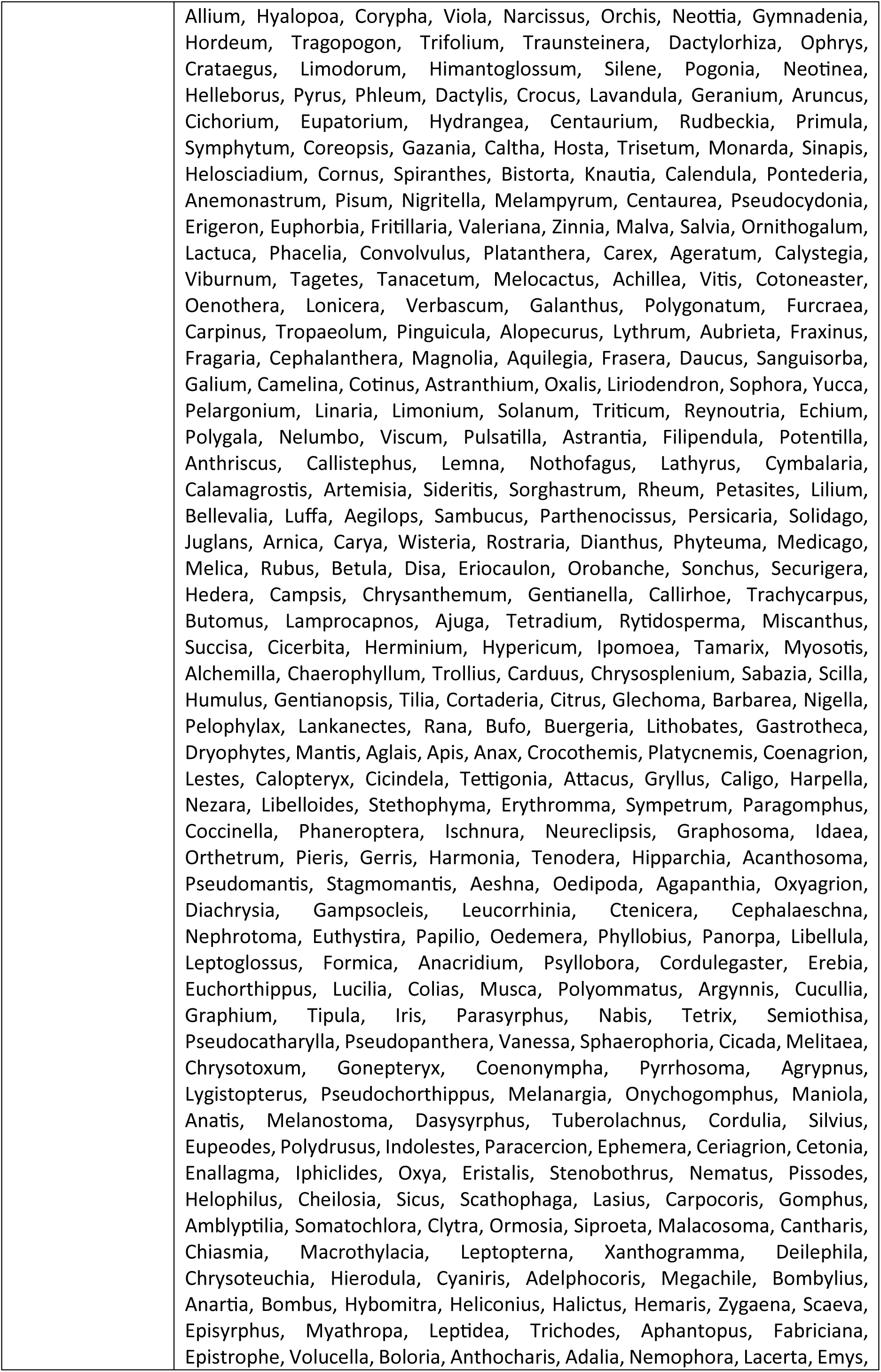
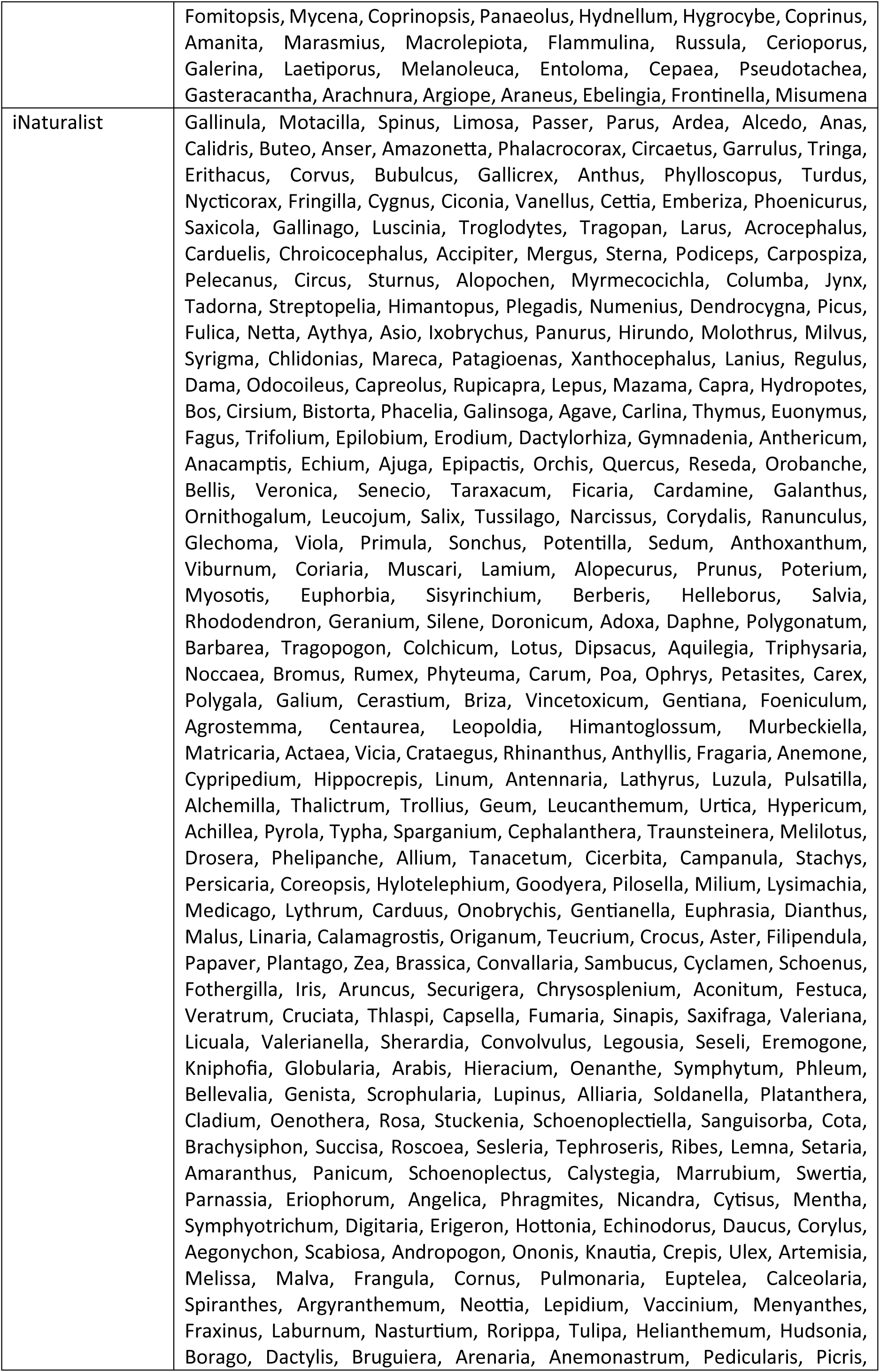
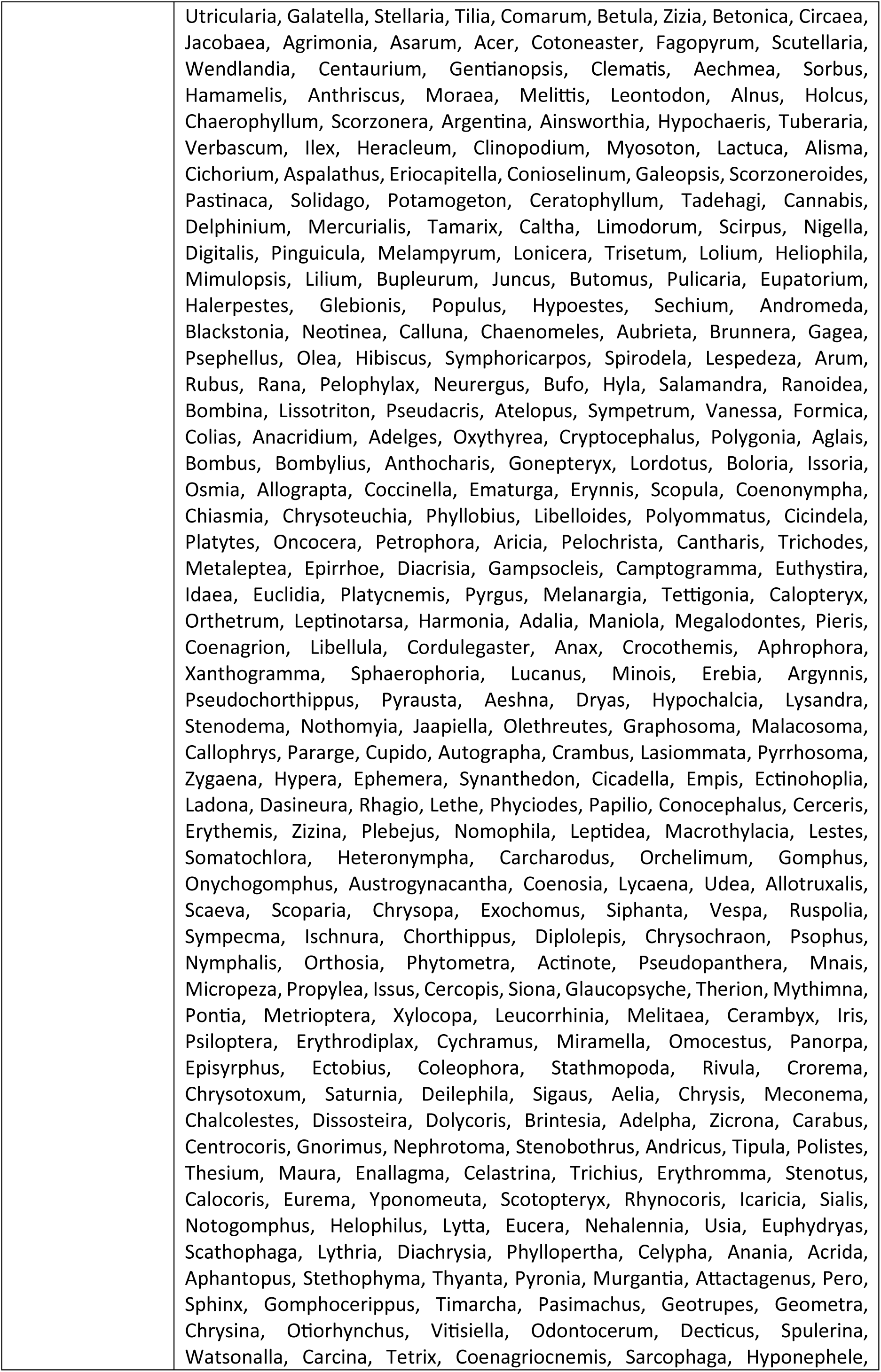
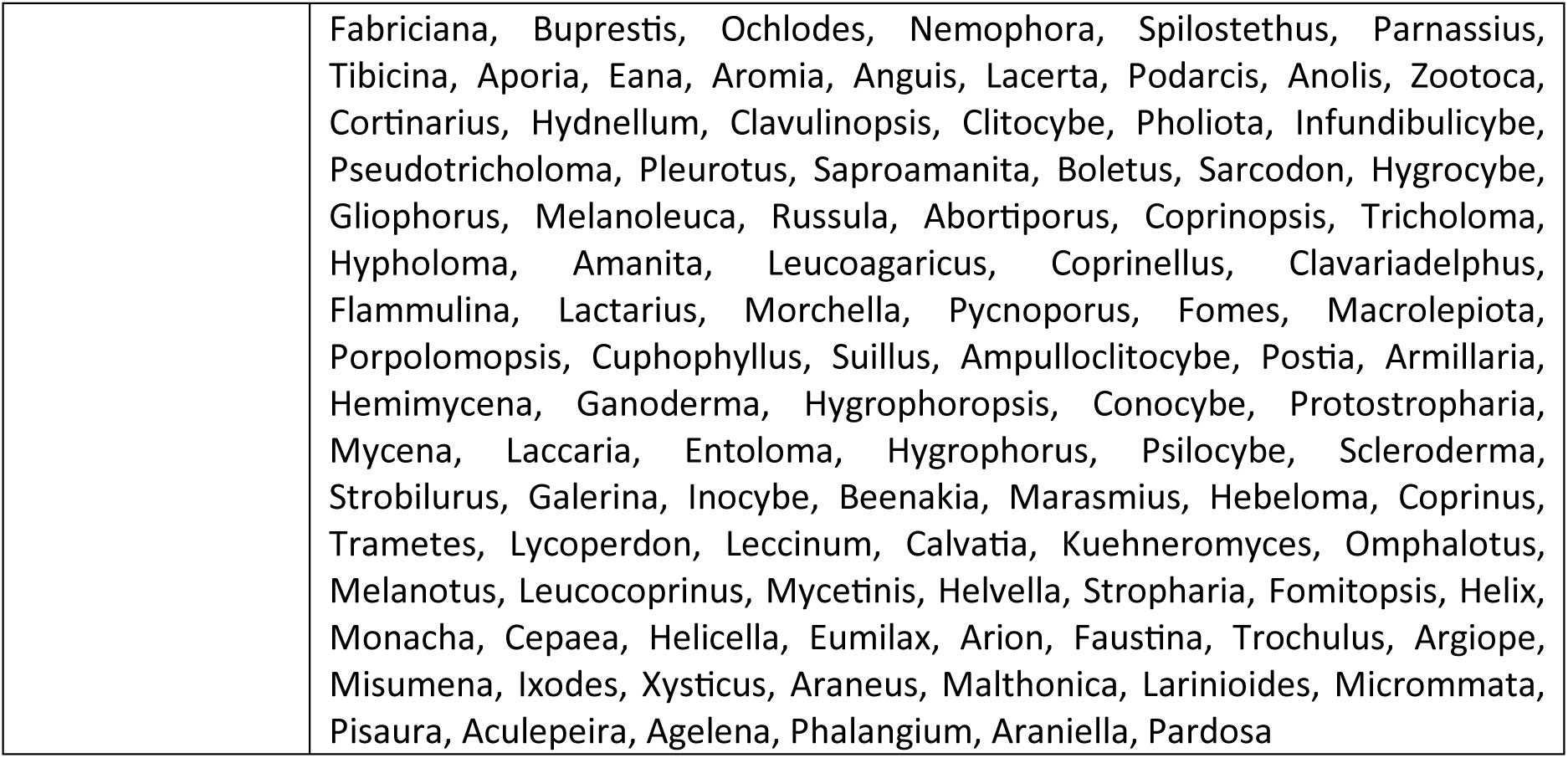
Overview of the extracted genus using the BioCLIP algorithm for Flickr and iNaturalist photographs of the most photographed wildlife species

## References

1. Allan, E., Bossdorf, O., Dormann, C.F., Prati, D., Gossner, M.M., Tscharntke, T., Blüthgen, N., Bellach, M., Birkhofer, K., Boch, S., Böhm, S., Börschig, C., Chatzinotas, A., Christ, S., Daniel, R., Diekötter, T., Fischer, C., Friedl, T., Glaser, K., Hallmann, C., Hodac, L., Hölzel, N., Jung, K., Klein, A.M., Klaus, V.H., Kleinebecker, T., Krauss, J., Lange, M., Morris, E.K., Müller, J., Nacke, H., Pašalić, E., Rillig, M.C., Rothenwöhrer, C., Schall, P., Scherber, C., Schulze, W., Socher, S.A., Steckel, J., Steffan-Dewenter, I., Türke, M., Weiner, C.N., Werner, M., Westphal, C., Wolters, V., Wubet, T., Gockel, S., Gorke, M., Hemp, A., Renner, S.C., Schöning, I., Pfeiffer, S., König-Ries, B., Buscot, F., Linsenmair, K.E., Schulze, E.-D., Weisser, W.W., Fischer, M., 2014. Interannual variation in land-use intensity enhances grassland multidiversity. Proc. Natl. Acad. Sci. 111 (1), 308–313.

2. Allan, E., Manning, P., Alt, F., Binkenstein, J., Blaser, S., Blüthgen, N., Böhm, S., Grassein, F., Hölzel, N., Klaus, V.H., Kleinebecker, T., Morris, E.K., Oelmann, Y., Prati, D., Renner, S.C., Rillig, M.C., Schaefer, M., Schloter, M., Schmitt, B., Schöning, I., Schrumpf, M., Solly, E., Sorkau, E., Steckel, J., Steffen-Dewenter, I., Stempfhuber, B., Tschapka, M., Weiner, C.N., Weisser, W.W., Werner, M., Westphal, C., Wilcke, W., Fischer, M., 2015. Land use intensification alters ecosystem multifunctionality via loss of biodiversity and changes to functional composition. Ecol. Lett. 18 (8), 834–843.

3. Andreatta, D., Bachofen, C., Dalponte, M., Klaus, V.H., Buchmann, N., 2023. Extracting flowering phenology from grassland species mixtures using time-lapse cameras. Remote Sens. Environ. 298, 113835.

4. Arroyo-Mora, J.P., Kalacska, M., Løke, T., Schläpfer, D., Coops, N.C., Lucanus, O., Leblanc, G., 2021. Assessing the impact of illumination on UAV pushbroom hyperspectral imagery collected under various cloud cover conditions. Remote Sens. Environ. 258.

5. August, T. A., Pescott, O. L., Joly, A., & Bonnet, P., 2020. AI naturalists might hold the key to unlocking biodiversity data in social media imagery. Patterns. 1, 100116

6. Austen, G.E., Dallimer, M., Irvine, K.N., Maund, P.R., Fish, R.D., Davies, Z.G., 2021. Exploring shared public perspectives on biodiversity attributes. People Nat. 3 (4), 901–913.

7. Axmanová, I., Kalusová, V., Danihelka, J., Dengler, J., Pergl, J., Pyšek, P., Večeřa, M., Attorre, F., Biurrun, I., Boch, S., Conradi, T., Gavilán, R.G., Jiménez-Alfaro, B., Knollová, I., Kuzemko, A., Lenoir, J., Leostrin, A., Medvecká, J., Moeslund, J.E., Obratov-Petkovic, D., Svenning, J.-C., Tsiripidis, I., Vassilev, K., Chytrý, M., 2021. Neophyte invasions in European grasslands. J. Veg. Sci. 32 (2), e12994.

8. Balfour, N.J., Ratnieks, F.L., 2022. The disproportionate value of ‘weeds’ to pollinators and biodiversity. J. Appl. Ecol. 59 (5), 1209–1218.

9. Ballantyne, M., Pickering, C., 2012. Ecotourism as a threatening process for wild orchids. J. Ecotourism 11 (1), 34–47.

10. Bardgett, R.D., Bullock, J.M., Lavorel, S., Manning, P., Schaffner, U., Ostle, N., Chomel, M., Durigan, G., Fry, E, L., Johnson, D., Lavallee, J.M., Le Provost, G., Luo, S., Png, K., Sankaran, M., Hou, X., Zhou, H., Ma, L., Ren, W., Li, X., Ding, Y., Li, Y., Shi, H., 2021. Combatting global grassland degradation. Nat. Rev. Earth Environ. 2, 720–735.

11. Barve, V., Hart, E., 2017. rinat: Access iNaturalist Data Through APIs. R package. https://cran.r-project.org/package=rinat

12. Bengtsson, J., Bullock, J.M., Egoh, B., Everson, C., Everson, T., O’Connor, T., O’Farrell, P.J., Smith, H.G., Lindborg, R., 2019. Grasslands—more important for ecosystem services than you might think. Ecosphere 10 (2), e02582.

13. Binkenstein, J., Renoult, J.P., Schaefer, H.M., 2013. Increasing land-use intensity decreases floral colour diversity of plant communities in temperate grasslands. Oecologia 173 (2), 461–471.

14. Blüthgen, N., Dormann, C.F., Prati, D., Klaus, V.H., Kleinebecker, T., Hölzel, N., Alt, F., Boch, S., Gockel, S., Hemp, A., Müller, J., Nieschulze, J., Renner, S.C., Schöning, I., Schumacher, U., Socher, S.A., Wells, K., Birkenhofer, K., Buscot, F., Oelmann, Y., Rothenwöhrer, C., Scherber, C., Tscharntke, T., Weiner, C.N., Fischer, M., Kalko, E.K.V., Linsenmair, K.E., Schulze, E.-D., Weisser, W.W., 2012. A quantitative index of land-use intensity in grasslands: Integrating mowing, grazing and fertilization. Basic Appl. Ecol. 13 (3), 207–220.

15. Briemle, G., Nitsche, S., Nitsche, L., 2002. Nutzungswertzahlen für Gefäßpflanzen des Grünlandes. Schriftenr. Vegetationskd. 38 (2), 203–225.

16. Bullock, J.M., McCracken, M.E., Bowes, M.J., Chapman, R.E., Graves, A.R., Hinsley, S.A., Hutchins, M.G., Nowakowski, M., Nicholls, D.J.E., Oakley, S., Old, G.H., Ostle, N.J., Redhead, J.W., Woodcock, B.A., Bedwell, T., Mayes, S., Robinson, V.S., Pywell, R.F., 2021. Does agri-environmental management enhance biodiversity and multiple ecosystem services?: A farm-scale experiment. Agric. Ecosyst. Environ. 320, 107582.

17. Campbell-Arvai, V., Serrano Vergel, R., Lindquist, M., Fox, N., Van Berkel, D., 2024. Tree selection for a virtual urban park: Comparing aided and unaided decision-making to support public engagement in greenspace design. Urban For. Urban Green. 99, 128447.

18. Chai-allah, A., Fox, N., Brunschwig, G., Bimonte, S., Joly, F., 2023. A trail-based approach using crowdsourced data to assess recreationists’ preferences for landscape. Landsc. Urban Plan. 233, 104700.

19. Chai-allah, A., Hermes, J., Zander, V., Joly, F., Brunschwig, G., Bimonte, S., Fox, N., in press. Assessing recreationists’ preferences of the landscape and species using crowdsourced images and machine learning. Landsc. Urban Plan. Accepted and in press

20. Chamberlain, E. and Szocs, E., 2013. taxize - taxonomic search and retrieval in R. F1000Research, 2:191. http://f1000research.com/articles/2-191/v2.

21. Cheng, X., Van Damme, S., Li, L., Uyttenhove, P., 2019. Evaluation of cultural ecosystem services: a review of methods. Ecosyst. Serv. 37, 100925. 10.1016/j.ecoser.2019.100925.

22. Congalton, R.G., Green, K., 2009. Assessing the Accuracy of Remotely Sensed Data: Principles and Practices (2nd ed.). CRC Press.

23. Conti, L., Malavasi, M., Galland, T., Komárek, J., Lagner, O., Carmona, C.P., de Bello, F., Rocchini, D., Šímová, P., 2021. The relationship between species and spectral diversity in grassland communities is mediated by their vertical complexity. Appl. Veg. Sci. 24 (3).

24. Di Cecco, G.J., Barve, V., Belitz, M.W., Stucky, B.J., Guralnick, R.P., Hurlbert, A.H., 2021. Observing the observers: how participants contribute data to iNaturalist and implications for biodiversity science. Bioscience. 71, 1179–1188.

25. Djordjević, V., Tsiftsis, S., 2022. The role of ecological factors in distribution and abundance of terrestrial orchids. Orchids phytochemistry, biology and horticulture: fundamentals and applications 3–72. Springer International Publishing.

26. European Union, 2018. Copernicus DEM - Global and European Digital Elevation Model: EEA-10: European coverage. Copernicus Land Monitoring Service, European Environment Agency (EEA) 10.5270/ESA-c5d3d65

27. Farley, J., Costanza, R., 2010. Payments for ecosystem services: from local to global. Ecol. Econ. 69 (11), 2060–2068.

28. Fischer, L.K., Honold, J., Cvejić, R., Delshammar, T., Hilbert, S., Lafortezza, R., Nastran, M., Nielsen, A.B., Pintar, M., van der Jagt, A.P.N., Kowarik, I., 2018. Beyond green: Broad support for biodiversity in multicultural European cities. Glob. Environ. Change 49, 35–45.

29. Fox, N., August, T., Mancini, F., Parks, K.E., Eigenbrod, F., Bullock, J. M., Sutter, L., Graham, L.J., 2020. “photosearcher” package in R: An accessible and reproducible method for harvesting large datasets from Flickr. SoftwareX, 12, 100624.

30. Fox, N., August, T., Mancini, F., Parks, K.E., Eigenbrod, F., Bullock, J.M., Sutter, L., Graham, L.J., 2021. Enriching social media data allows a more robust representation of cultural ecosystem services. Ecosyst. Serv. 50, 101328.

31. Fox, N., Chamberlain, B., Lindquist, M., Van Berkel, D., 2022a. Understanding landscape aesthetics using a novel viewshed assessment of social media locations within the Troodos UNESCO Global Geopark, Cyprus. Front. Environ. Sci. 10, 884115.

32. Fox, N., Graham, L.J., Eigenbrod, F., Bullock, J.M., Parks, K.E., 2022b. Geodiversity supports cultural ecosystem services: an assessment using social media. Geoheritage 14 (1), 27.

33. Geurts, E.M., Reynolds, J.D., Starzomski, B.M., 2023. Turning observations into biodiversity data: Broadscale spatial biases in community science. Ecosphere 14, e4582.

34. Ghermandi, A., Langemeyer, J., Van Berkel, D., Calcagni, F., Depietri, Y., Vigl, L.E., Fox, N., Havinga, I., Jäger, H., Kaiser, N., Karasov, O., McPhearson, T., Podschun, S., Ruiz-Frau, A., Sinclair, M., Venohr, M., Wood, S.A., 2023. Social media data for environmental sustainability: A critical review of opportunities, threats, and ethical use. One Earth 6 (3), 236–250.

35. Gosal, A.S., Geijzendorffer, I.R., V’aclavík, T., Poulin, B., Ziv, G., 2019. Using social media, machine learning and natural language processing to map multiple recreational beneficiaries. Ecosyst. Serv. 38, 100958.

36. Hauser, L.T., Timmermans, J., van der Windt, N., Sil, Â.F., César de Sá, N., Soudzilovskaia, N.A., van Bodegom, P.M., 2021. Explaining discrepancies between spectral and in-situ plant diversity in multispectral satellite earth observation. Remote Sens. Environ. 265, 112684.

37. Havinga, I., Bogaart, P.W., Hein, L., Tuia, D., 2020. Defining and spatially modelling cultural ecosystem services using crowdsourced data. Ecosyst. Serv. 43, 101091.

38. Havinga, I., Marcos, D., Bogaart, P., Massimino, D., Hein, L., Tuia, D., 2023. Social media and deep learning reveal specific cultural preferences for biodiversity. People Nat. 5(3), 981–998.

39. Havinga, I., Marcos, D., Bogaart, P., Tuia, D., Hein, L., 2024. Understanding the sentiment associated with cultural ecosystem services using images and text from social media. Ecosyst. Serv. 65, 101581.

40. Hijmans, R.J., 2023. raster: Geographic analysis and modeling with raster data. R package version 3.6–26. https://CRAN.R-project.org/package=raster

41. Hoyle, H., Hitchmough, J., Jorgensen, A., 2017. All about the ‘wow factor’? The relationships between aesthetics, restorative effect and perceived biodiversity in designed urban planting. Landsc. Urban Plan. 164, 109–123.

42. Hoyle, H., Norton, B., Dunnett, N., Richards, J.P., Russell, J.M., Warren, P., 2018. Plant species or flower colour diversity? Identifying the drivers of public and invertebrate response to designed annual meadows. Landsc. Urban Plan. 180, 103–113.

43. Huber, N., Ginzler, C., Pazúr, R., Descombes, P., Baltensweiler, A., Ecker, K., Meier, E., Price, B., 2022. Distribution maps of permanent grassland habitats for Switzerland. EnviDat. https://www.doi.org/10.16904/envidat.341.

44. Isselstein, J., Jeangros, B., Pavlu, V., 2005. Agronomic aspects of biodiversity targeted management of temperate grasslands in Europe–a review. Agron. Res. 3 (2), 139–151.

45. Jiang, Y., Yuan, T., 2017. Public perceptions and preferences for wildflower meadows in Beijing, China. Urban For. Urban Green. 27, 324–331.

46. Johansen, L., Westin, A., Wehn, S., Iuga, A., Ivascu, C.M., Kallioniemi, E., Lennartsson, T., 2019. Traditional semi-natural grassland management with heterogeneous mowing times enhances flower resources for pollinators in agricultural landscapes. Global Ecol. Conserv. 18, e00619.

47. Junge, X., Jacot, K.A., Bosshard, A., Lindemann-Matthies, P., 2009. Swiss people’s attitudes towards field margins for biodiversity conservation. J. Nat. Conserv. 17 (3), 150–159.

48. Junge, X., Lindemann-Matthies, P., Hunziker, M., Schüpbach, B., 2011. Aesthetic preferences of non-farmers and farmers for different land-use types and proportions of ecological compensation areas in the Swiss lowlands. Biol. Conserv. 144 (5), 1430–1440.

49. Junge, X., Schüpbach, B., Walter, T., Schmid, B., Lindemann-Matthies, P., 2015. Aesthetic quality of agricultural landscape elements in different seasonal stages in Switzerland. Landsc. Urban Plan. 133, 67–77.

50. Klaus, V.H., Jehle, A., Richter, F., Buchmann, N., Knop, E., Lüscher, G., 2023. Additive effects of two agri-environmental schemes on plant diversity but not on productivity indicators in permanent grasslands in Switzerland. J. Environ. Manage. 348, 119416.

51. Kosanic, A., Petzold, J., 2020. A systematic review of cultural ecosystem services and human wellbeing. Ecosyst. Serv. 45, 101168.

52. Kull, T., Selgis, U., Peciña, M.V., Metsare, M., Ilves, A., Tali, K., Sepp, K., Kull, K., Shefferson, R.P., 2016. Factors influencing IUCN threat levels to orchids across Europe on the basis of national red lists. Ecol. Evol. 6 (17), 6245–6265.

53. Lee, H., Seo, B., Cord, A.F., Volk, M., Lautenbach, S., 2022. Using crowdsourced images to study selected cultural ecosystem services and their relationships with species richness and carbon sequestration. Ecosyst. Serv. 54, 101411.

54. Liaw, A., Wiener, M., 2002. Classification and Regression by randomForest. R News, 2 (3), 1822.

55. Lindemann-Matthies, P., Junge, X., Matthies, D., 2010. The influence of plant diversity on people’s perception and aesthetic appreciation of grassland vegetation. Biol. Conserv. 143 (1), 195–202.

56. Lindemann-Matthies, P., Brieger, H., 2016. Does urban gardening increase aesthetic quality of urban areas? A case study from Germany. Urban For. Urban Green. 17, 33–41.

57. Lopez, B., Minor, E., Crooks, A., 2020. Insights into human-wildlife interactions in cities from bird sightings recorded online. Landsc. Urban Plan. 196, 103742.

58. Lüscher, A., Grieder, C., Huguenin-Elie, O., Klaus, V., Reidy, B., Schneider, M.K., Schubiger, F., Suter, D., Suter, M., Kölliker, R., 2019. Grassland systems in Switzerland with a main focus on sown grasslands. Grassland Sci. Eur. 24, 3–16.

59. Mackay, C.M., Schmitt, M.T., 2019. Do people who feel connected to nature do more to protect it? A meta-analysis. J. Environ. Psychol. 65, 101323.

60. Martinková, Z., Honek, A., Pekár, S., 2009. Seed availability and gap size influence seedling emergence of dandelion (Taraxacum officinale) in grasslands. Grass Forage Sci. 64 (2), 160–168.

61. Matzdorf, B., Kaiser, T., Rohner, M.S., 2008. Developing biodiversity indicator to design efficient agri-environmental schemes for extensively used grassland. Ecol. Indic. 8 (3), 256–269.

62. Mou, N., Wang, J., Zheng, Y., Zhang, L., Makkonen, T., Yang, T., Niu, J., 2023. Flowers as attractions in urban parks: Evidence from social media data. Urban For. Urban Green. 82, 127874.

63. Nowak-Olejnik, A., Mocior, E., Hibner, J., Tokarczyk, N., 2020. Human perceptions of cultural ecosystem services of semi-natural grasslands: The influence of plant communities. Ecosyst. Serv. 46, 101208.

64. Öckinger, E., Eriksson, A.K., Smith, H.G., 2006. Effects of grassland abandonment, restoration and management on butterflies and vascular plants. Biol. Conserv. 133 (3), 291–300.

65. Oksanen, J., Blanchet, F.G., Kindt, R., Legendre, P., Minchin, P.R., O’hara, R.B., Simpson, G.L., Solymos, P., Stevens, M.H.H., Wagner, H., 2022. ‘vegan’, Community Ecology Package. https://CRAN.R-project.org/package=vegan

66. Perrone, M., Conti, L., Galland, T., Komárek, J., Lagner, O., Torresani, M., Rossi, C., Carmona, C. P., de Bello, F., Rocchini, D., Moudrý, V., Šímová, P., Bagella, S., Malavasi, M., 2024. “Flower power”: How flowering affects spectral diversity metrics and their relationship with plant diversity. Ecol. Inform. 81, 102589.

67. Peter, S., Le Provost, G., Mehring, M., Müller, T., Manning, P., 2022. Cultural worldviews consistently explain bundles of ecosystem service prioritisation across rural Germany. People Nat. 4 (1), 218–230.

68. Plieninger, T., Dijks, S., Oteros-Rozas, E., Bieling, C., 2013. Assessing, mapping, and quantifying cultural ecosystem services at community level. Land Use Policy 33, 118–129.

69. Ravetto Enri, S., Probo, M., Farruggia, A., Lanore, L., Blanchetete, A., Dumont, B., 2017. A biodiversity-friendly rotational grazing system enhancing flower-visiting insect assemblages while maintaining animal and grassland productivity. Agric. Ecosyst. Environ. 241, 1–10.

70. R Core Team, 2024. R: A language and environment for statistical computing. R Foundation for Statistical Computing, Vienna, Austria. https://www.R-project.org/

71. Richter, F., Jan, P., El Benni, N., Lüscher, A., Buchmann, N., Klaus, V.H., 2021. A guide to assess and value ecosystem services of grasslands. Ecosyst. Serv. 52, 101376.

72. Richter, F., Suter, M., Lüscher, A., Buchmann, N., El-Benni, N., Feola-Conz, R., Hartmann, M., Jan, P., Klaus, V.H., 2024. Effects of management practices on the ecosystem service multifunctionality of temperate grasslands. Nat. Commun. 15, 3829.

73. Ridding, L.E., Redhead, J.W., Oliver, T.H., Schmucki, R., McGinlay, J., Graves, A.R., Morris, J., Bradbury, R.B., King, H., Bullock, J.M., 2018. The importance of landscape characteristics for the delivery of cultural ecosystem services. J. Environ. Manage. 206, 1145–1154.

74. Rocchini, D., Boyd, D.S., Féret, J.B., Foody, G.M., He, K.S., Lausch, A., Nagendra, H., Wegmann, M., Pettorelli, N., 2016. Satellite remote sensing to monitor species diversity: potential and pitfalls. Remote Sens. Ecol. Conserv. 2 (1), 25–36.

75. Rocchini, D., Thouverai, E., Marcantonio, M., Iannacito, M., Da Re, D., Torresani, M., Bacaro, G., Bazzichetto, M., Bernardi, A., Foody, G. M., Furrer, R., Kleijn, D., Larsen, S., Lenoir, J., Malavasi, M., Marchetto, E., Messori, F., Montaghi, A., Moudrý, V., Naimi, B., Ricotta, C., Rossini, M., Santi, F., Santos, M.J., Schaepman, M.E., Schneider, F.D., Schuh, L., Silvestri, S., Ŝímová, P., Skidmore, A.K., Tattoni, C., Tordoni, E., Vicario, S., Zannini, P., Wegmann, M., 2021. rasterdiv—An Information Theory tailored R package for measuring ecosystem heterogeneity from space: To the origin and back. Methods Ecol. Evol. 12 (6), 1093–1102.

76. Rossi, C., Kneubühler, M., Schütz, M., Schaepman, M.E., Haller, R.M., Risch, A.C., 2022. Spatial resolution, spectral metrics and biomass are key aspects in estimating plant species richness from spectral diversity in species-rich grasslands. Remote Sens. Ecol. Conserv. 8 (3), 297–314.

77. Schils, R.L.M., Bufe, C., Rhymer, C.M., Francksen, R.M., Klaus, V.H., Abdalla, M., Milazzo, F., Lellei-Kovács, E., ten Berge, H., Bertora, C., Chodkiewicz, A., Dǎmǎtîrcǎ, C., Feigenwinter, I., Fernández-Rebollo, P., Ghiasi, S., Hejduk, S., Hiron, M., Janicka, M., Pellaton, R., Smith, K.E., Thorman, R., Vanwalleghem, T., Williams, J., Zavattaro, L., Kampen, J., Derkx, R., Smith, P., Whittingham, M.J., Buchmann, N., Price, J.P.N., 2022. Permanent grasslands in Europe: Land use change and intensification decrease their multifunctionality. Agric. Ecosyst. Environ. 330, 107891.

78. Schirpke, U., Ghermandi, A., Sinclair, M., Van Berkel, D., Fox, N., Vargas, L., Willemen, L., 2023. Emerging technologies for assessing ecosystem services: A synthesis of opportunities and challenges. Ecosyst. Serv. 63, 101558.

79. Schwemmer, C., 2019. imgrec: An Interface for Image Recognition. R package version 0.1, 1. https://github.com/cschwem2er/imgrec

80. Sheppard, A.W., 1991. Heracleum sphondylium L. J. Ecol. 79 (1), 235–258.

81. Southon, G.E., Jorgensen, A., Dunnett, N., Hoyle, H., Evans, K.L., 2017. Biodiverse perennial meadows have aesthetic value and increase residents’ perceptions of site quality in urban green-space. Landsc. Urban Plan. 158, 105–118.

82. Stevens, S., Wu, J., Thompson, M.J., Campolongo, E.G., Song, C.H., Carlyn, D.E., Dong, L., Dahdul, W.M., Stewart, C., Berger-Wolf, T., Chao, W.-L., Su, Y., 2024. BioCLIP: A Vision Foundation Model for the Tree of Life. Proceedings of the IEEE/CVF Conference on Computer Vision and Pattern Recognition, 19412–19424.

83. Straffelini, E., Luo, J., Tarolli, P., 2024. Climate change is threatening mountain grasslands and their cultural ecosystem services. Catena 237, 107802.

84. Šumrada, T., Vreš, B., Čelik, T., Šilc, U., Rac, I., Udovč, A., Erjavec, E., 2021. Are result-based schemes a superior approach to the conservation of High Nature Value grasslands? Evidence from Slovenia. Land Use Policy, 111, 105749.

85. Swisstopo (Eds.), 2020. The high precision digital elevation model of Switzerland swissALTI3D (2m). Swiss Federal Office of Topography. www.swisstopo.admin.ch/en/geodata/height/alti3d.html#technische_details

86. Torresani, M., Kleijn, D., de Vries, J.P.R., Bartholomeus, H., Chieffallo, L., Cazzolla Gatti, R., Moudrý, V., da Re, D., Tomelleri, E., Rocchini, D., 2023. A novel approach for surveying flowers as a proxy for bee pollinators using drone images. Ecol. Indic. 149, 110123.

87. Torresani, M., Rossi, C., Perrone, M., Hauser, L.T., Féret, J.-B., Moudrý, V., Simova, P., Ricotta, C., Foody, G.M., Kacic, P., Feilhauer, H., Malavasi, M., Tognetti, R., Rocchini, D., 2024. Reviewing the Spectral Variation Hypothesis: Twenty years in the tumultuous sea of biodiversity estimation by remote sensing. Ecol. Inform. 82, 102702.

88. Van Berkel, D.B., Verburg, P.H., 2014. Spatial quantification and valuation of cultural ecosystem services in an agricultural landscape. Ecol. Indic. 37, 163–174.

89. Wang, R., Gamon, J.A., 2019. Remote sensing of terrestrial plant biodiversity. Remote Sens. Environ. 231.

90. Wang, L., Huang, L., Cao, W., Zhai, J., Fan, J., 2024. Assessing grassland cultural ecosystem services supply and demand for promoting the sustainable realization of grassland cultural values. Sci. Total Environ. 912, 169255.

91. Wickham, H., 2016. Ggplot2: Elegant graphics for data analysis (2^nd^ ed.). Springer International Publishing.

92. Wilkins, E.J., Van Berkel, D., Zhang, H., Dorning, M.A., Beck, S.M., Smith, J.W., 2022. Promises and pitfalls of using computer vision to make inferences about landscape preferences: Evidence from an urban-proximate park system. Landsc. Urban Plan. 219, 104315.

93. Zielstra, D., Hochmair, H.H., 2013. Positional accuracy analysis of Flickr and Panoramio images for selected world regions. J. Spat. Sci. 58, 251–273.

