## Supplementary figures and images for "Powerful flowers: Public perception of grassland aesthetics is strongly related to management and biodiversity"

### Pictures used in questionnaire survey (part of supplementary materials)

1

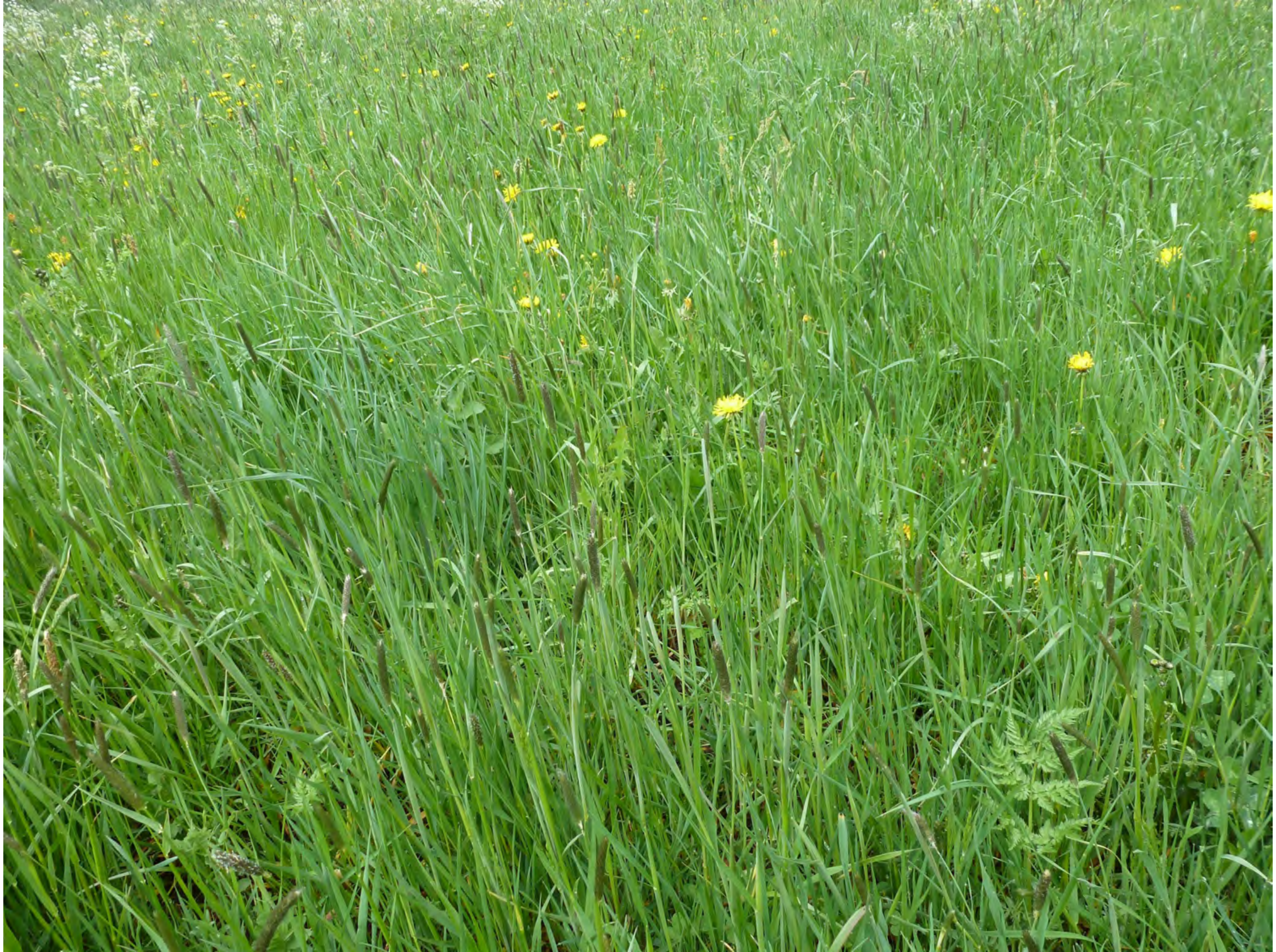

2

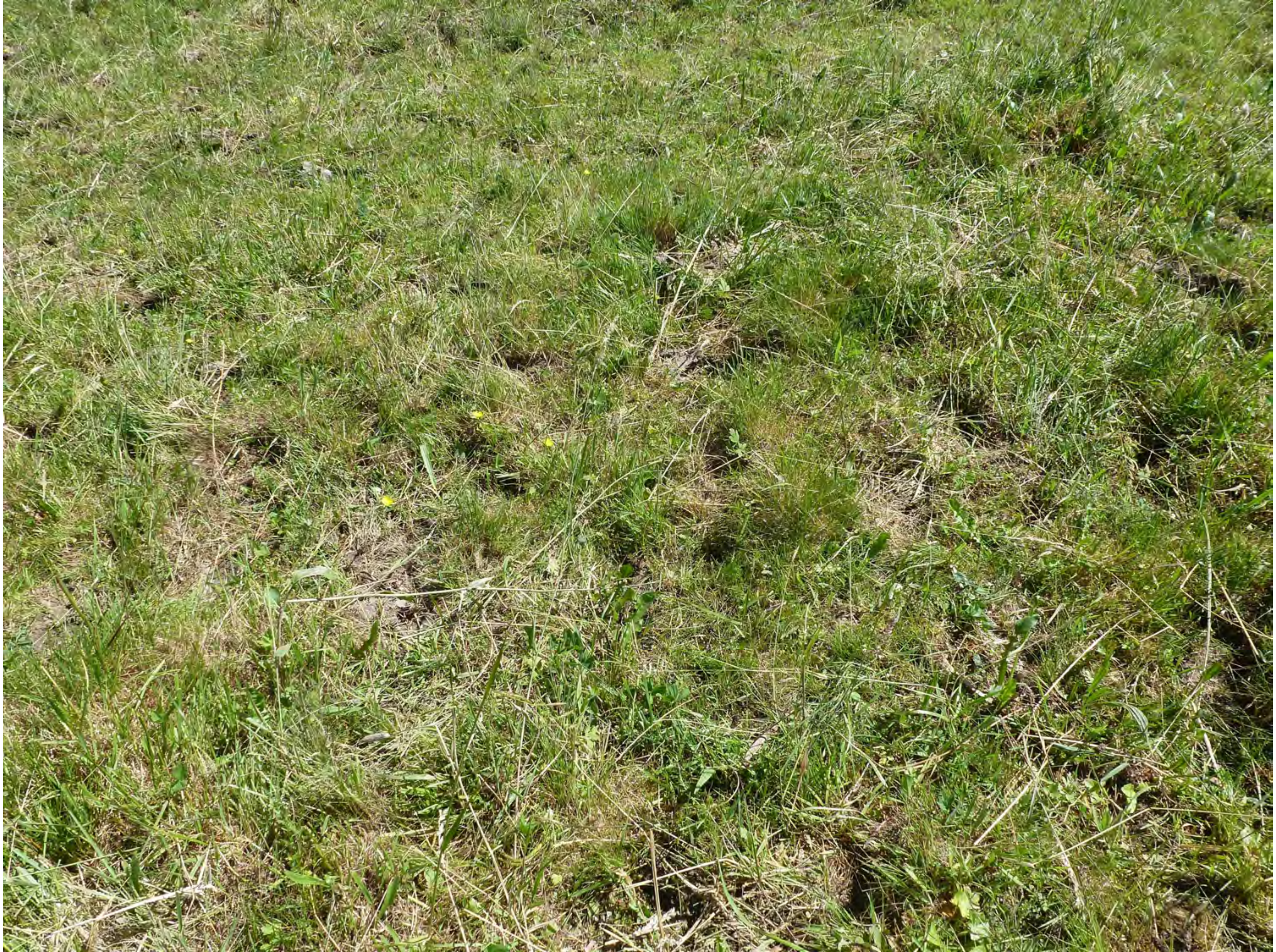

3

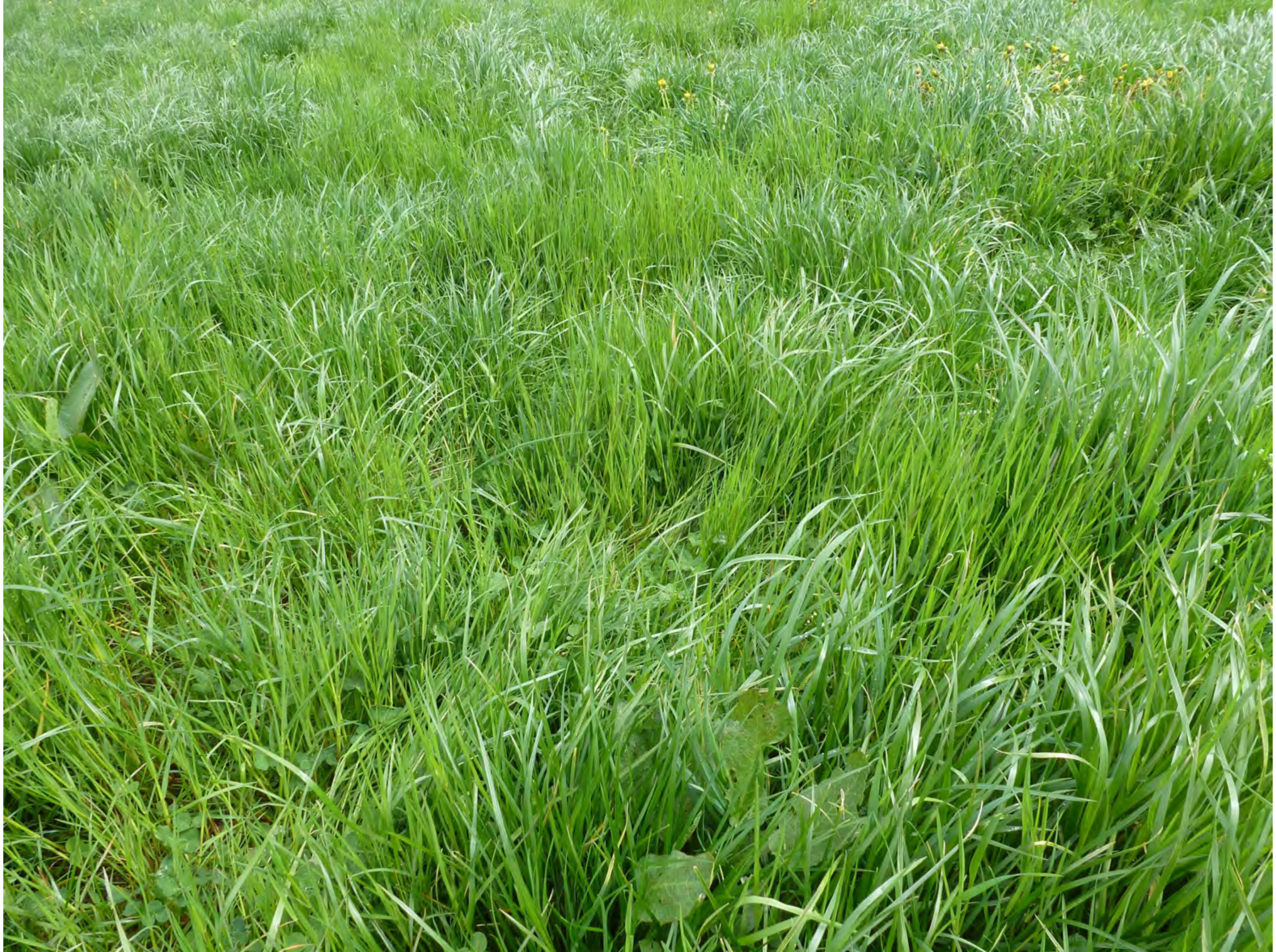

4

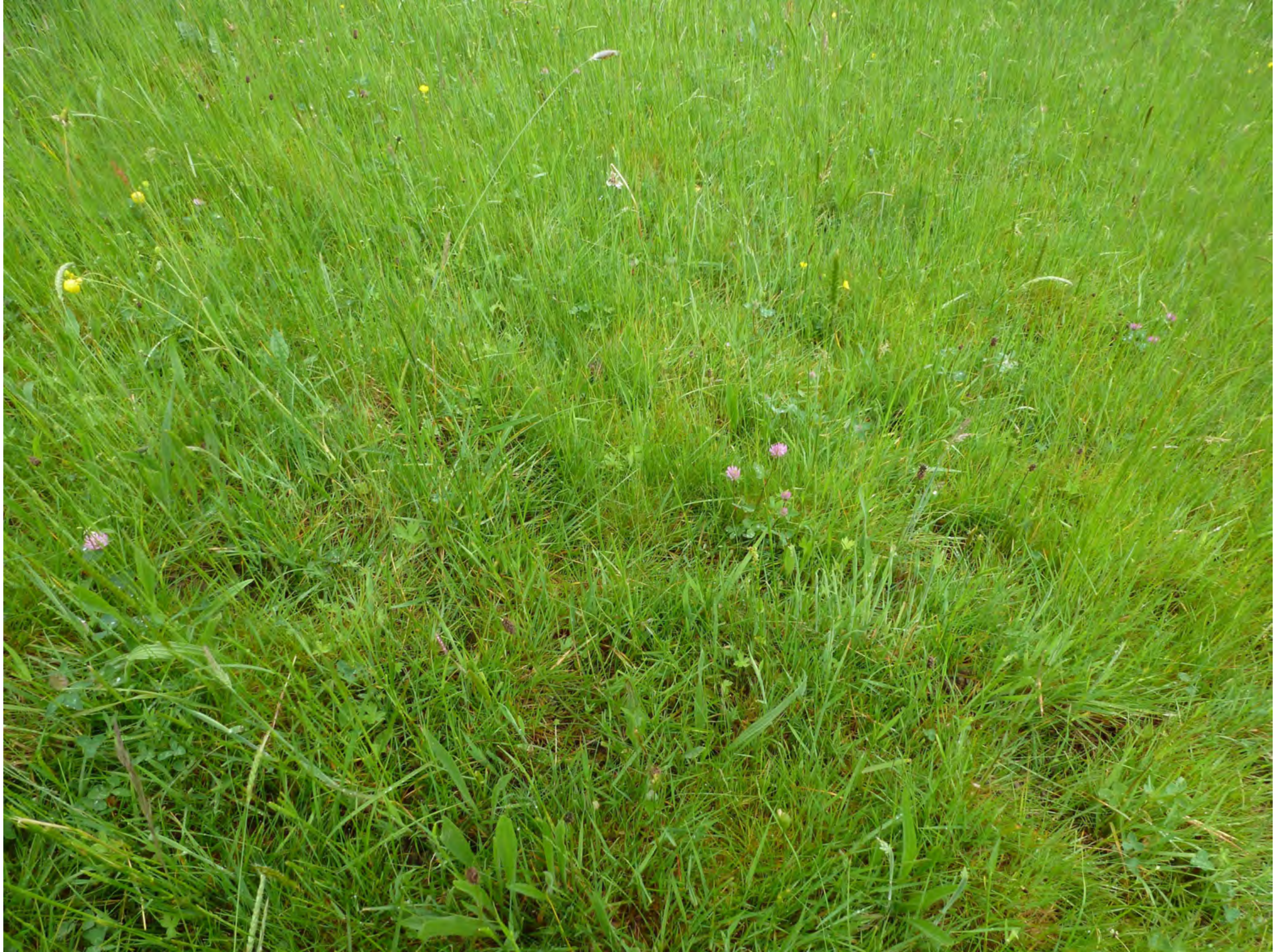

5

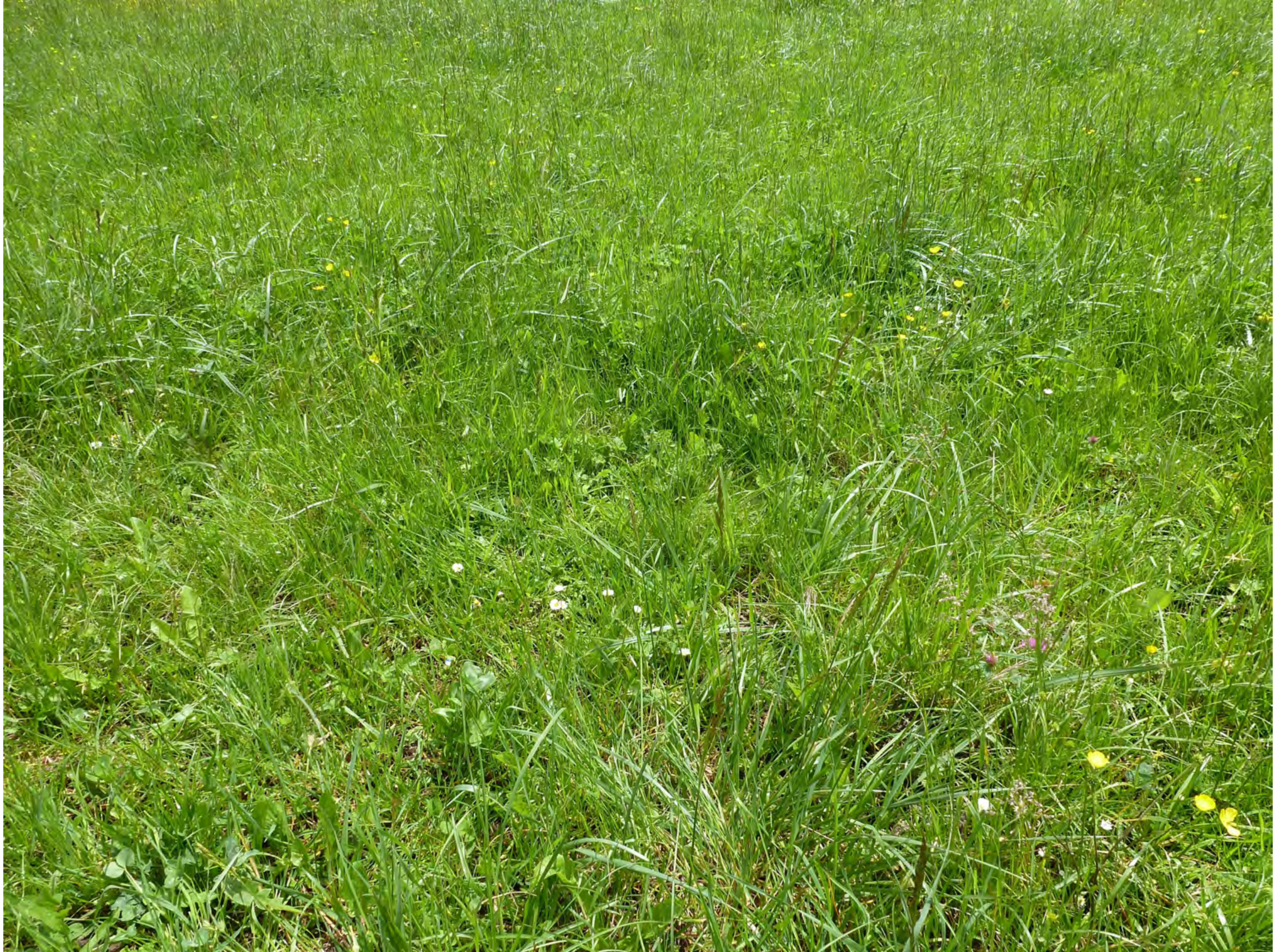

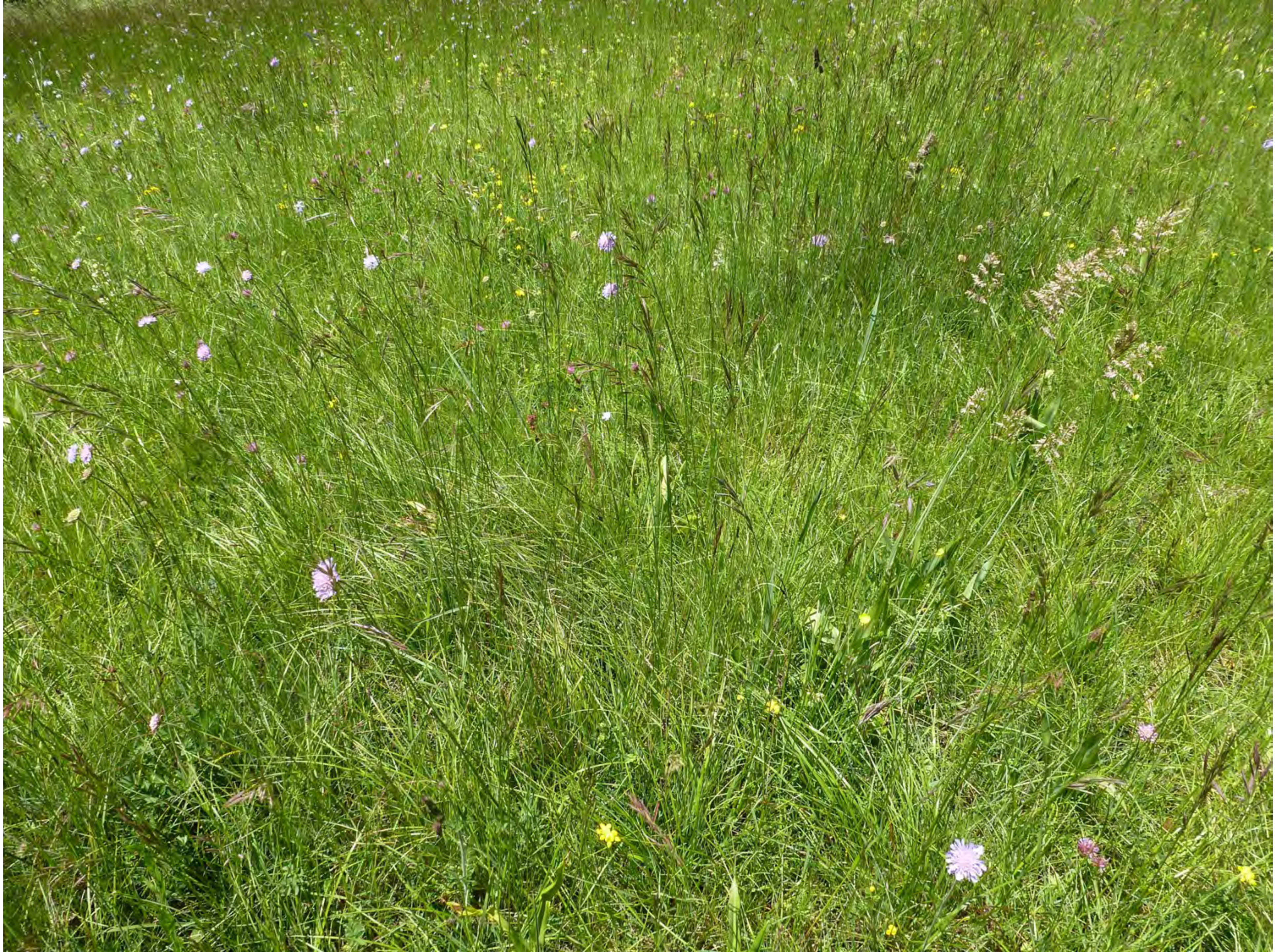

7

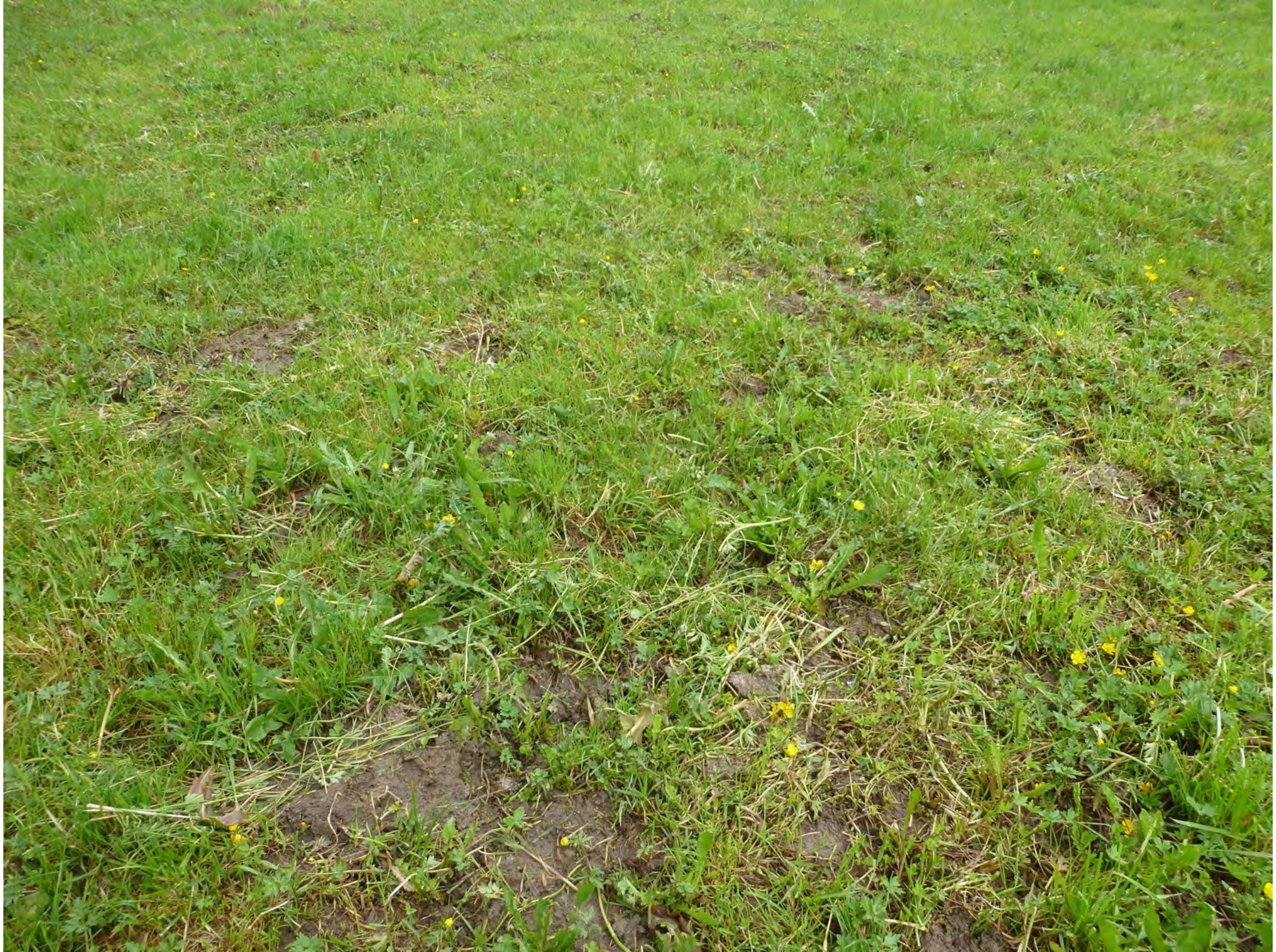

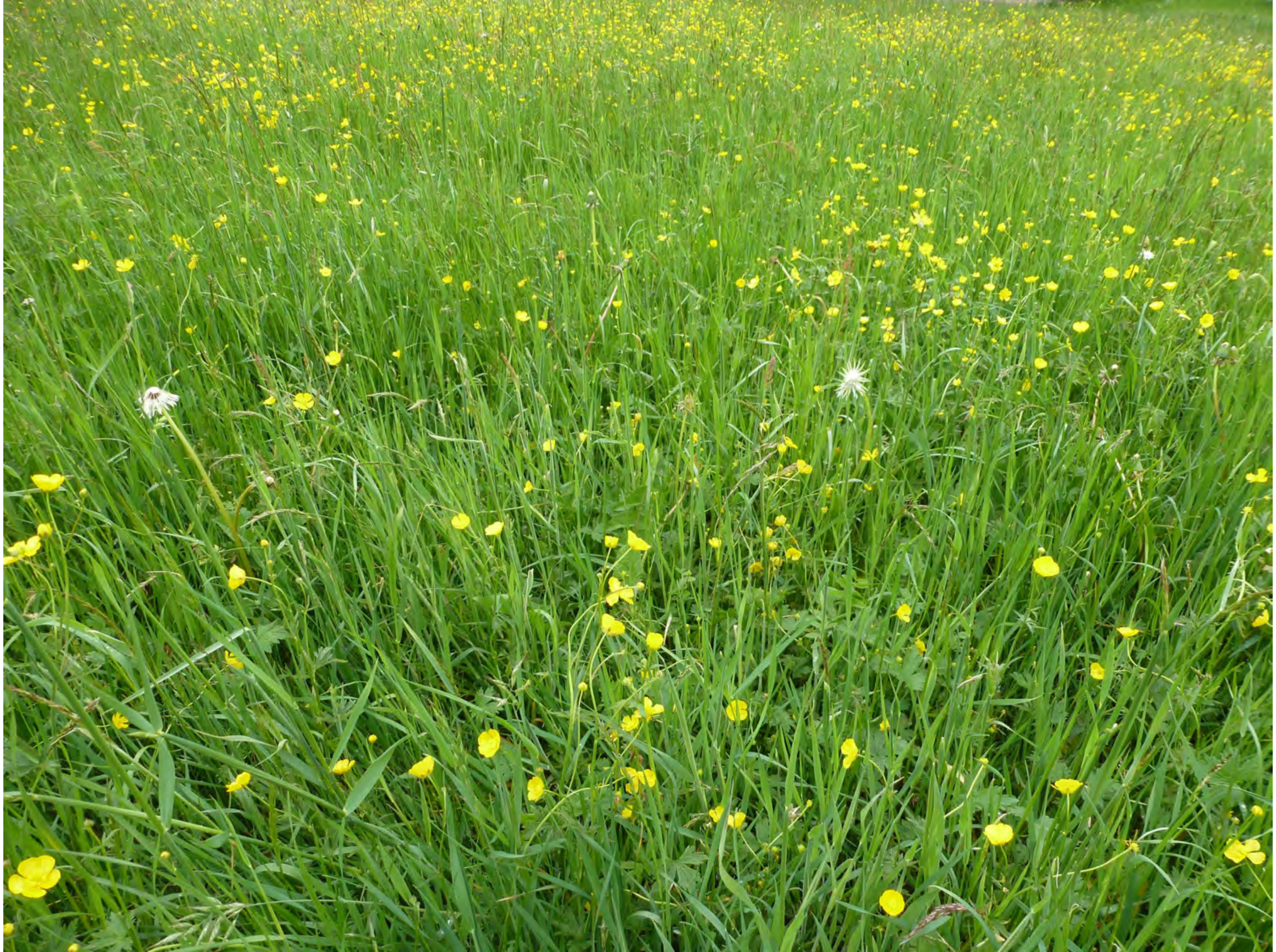

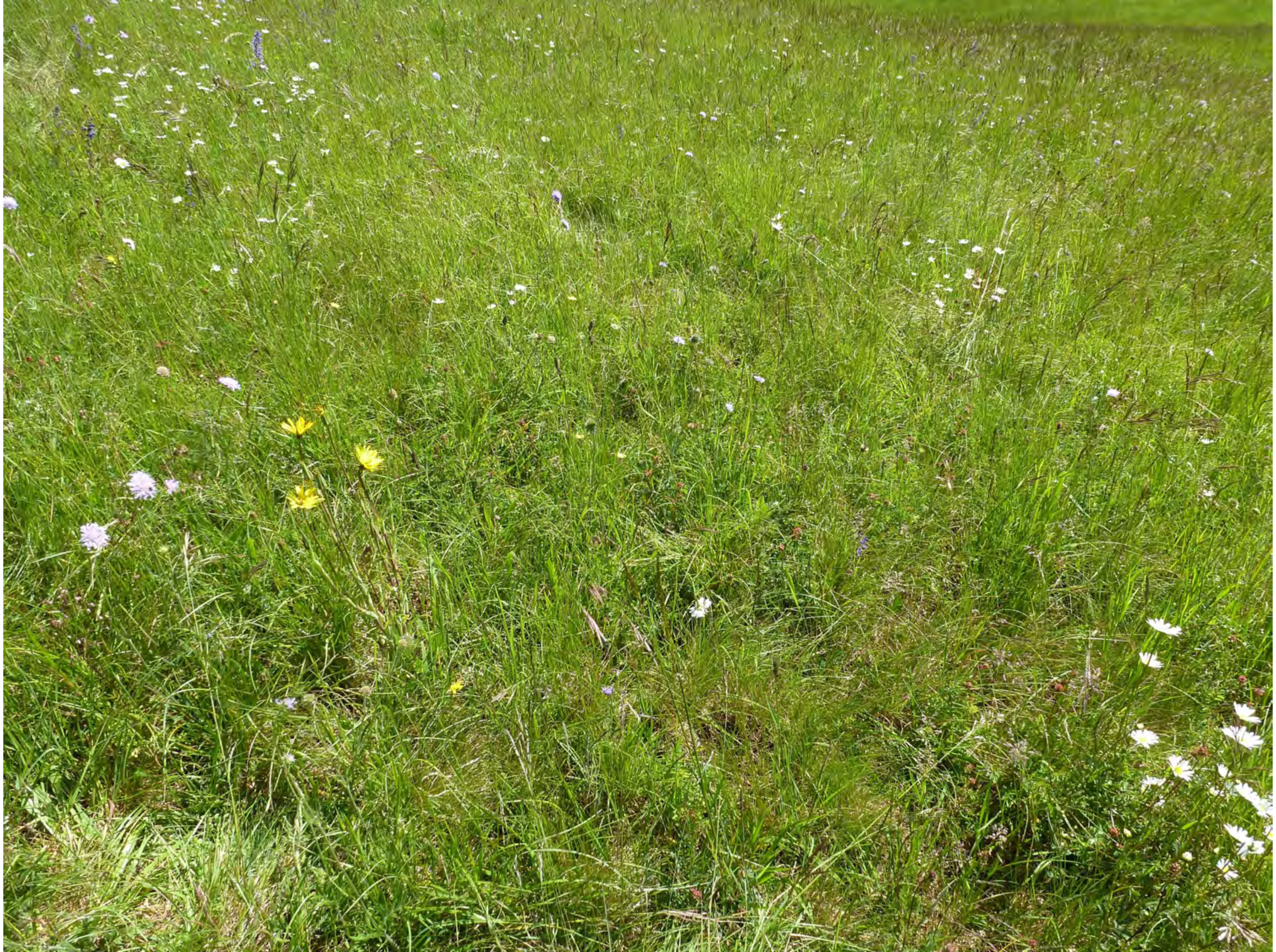

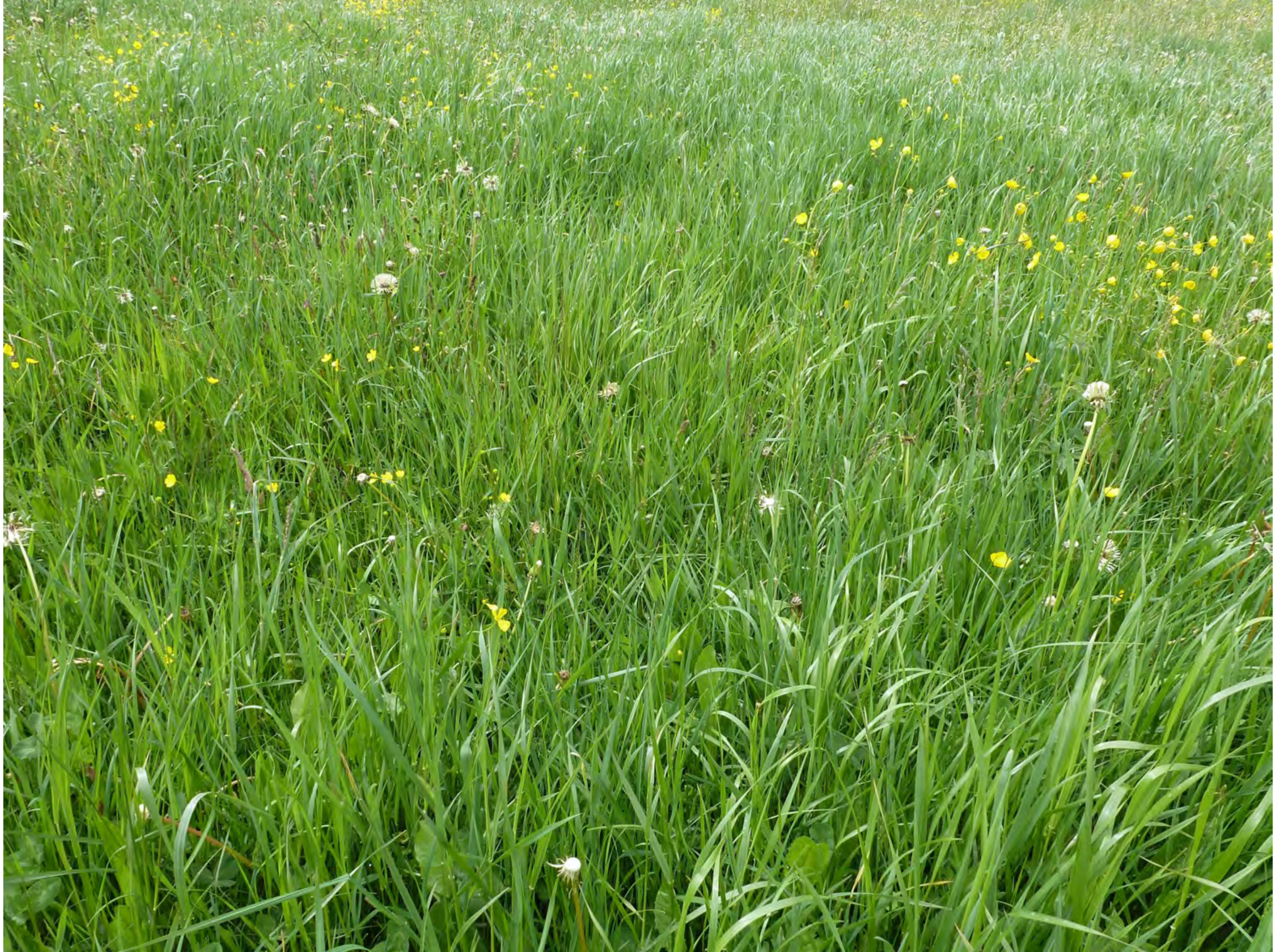

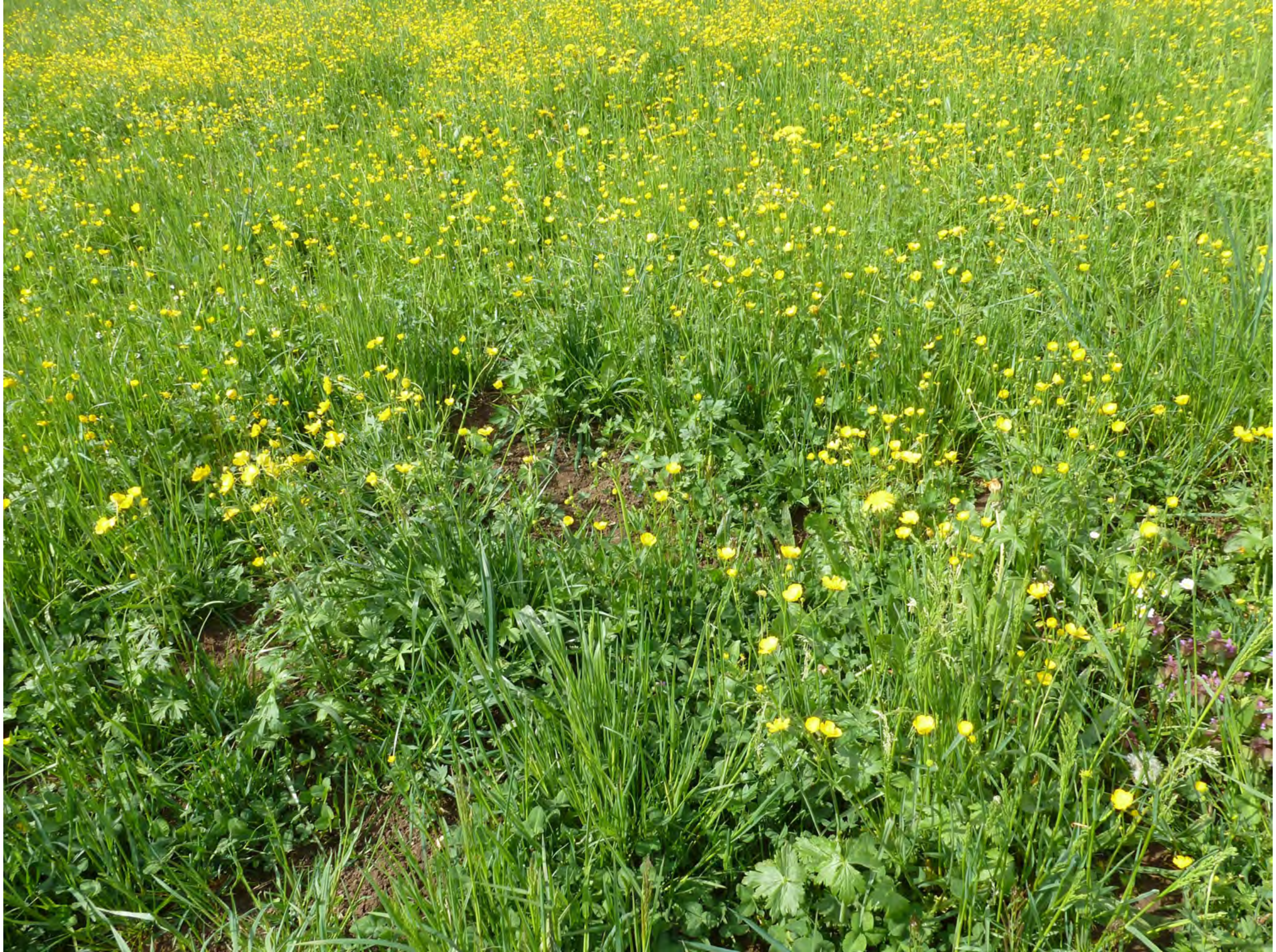

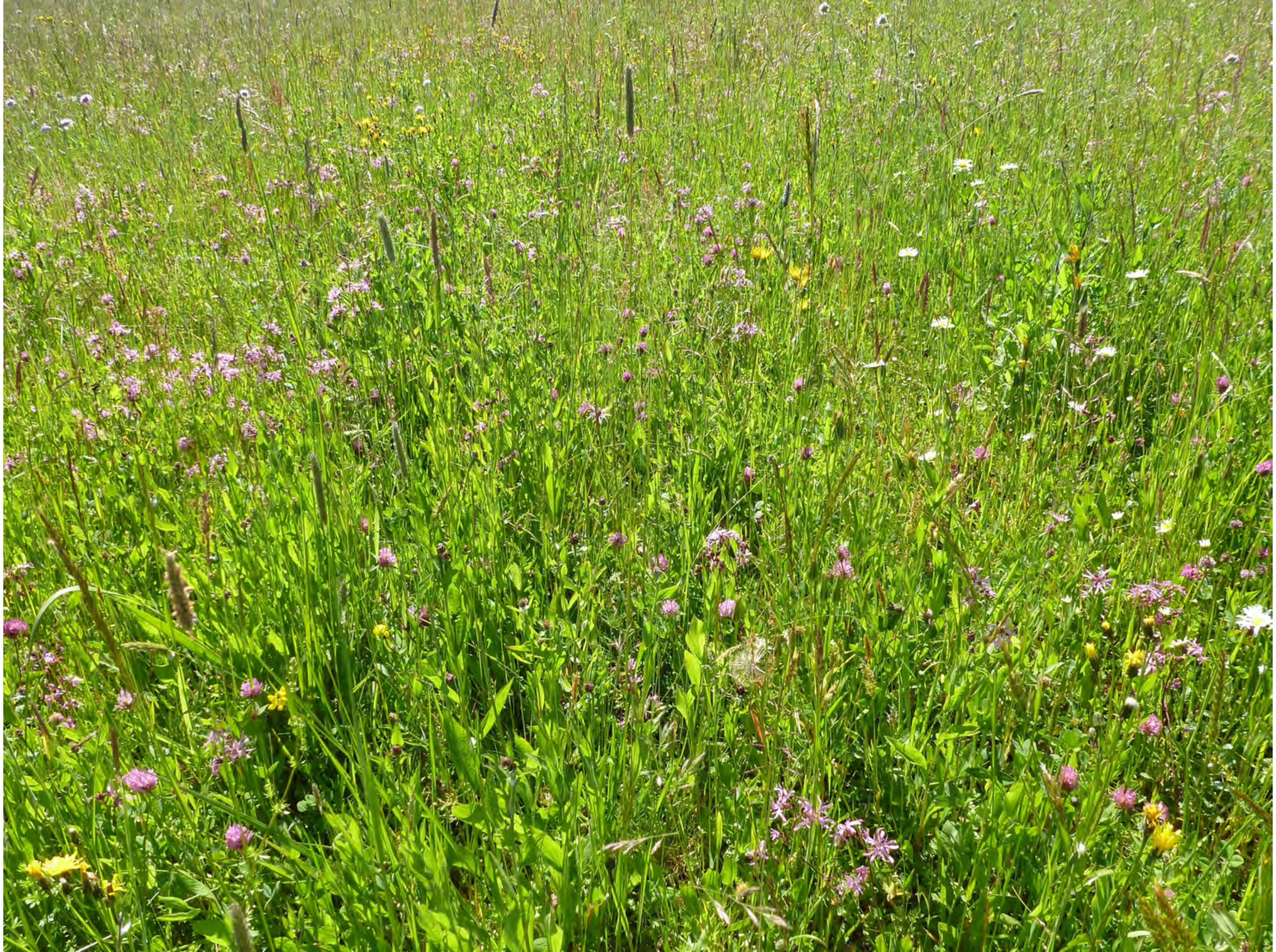

13

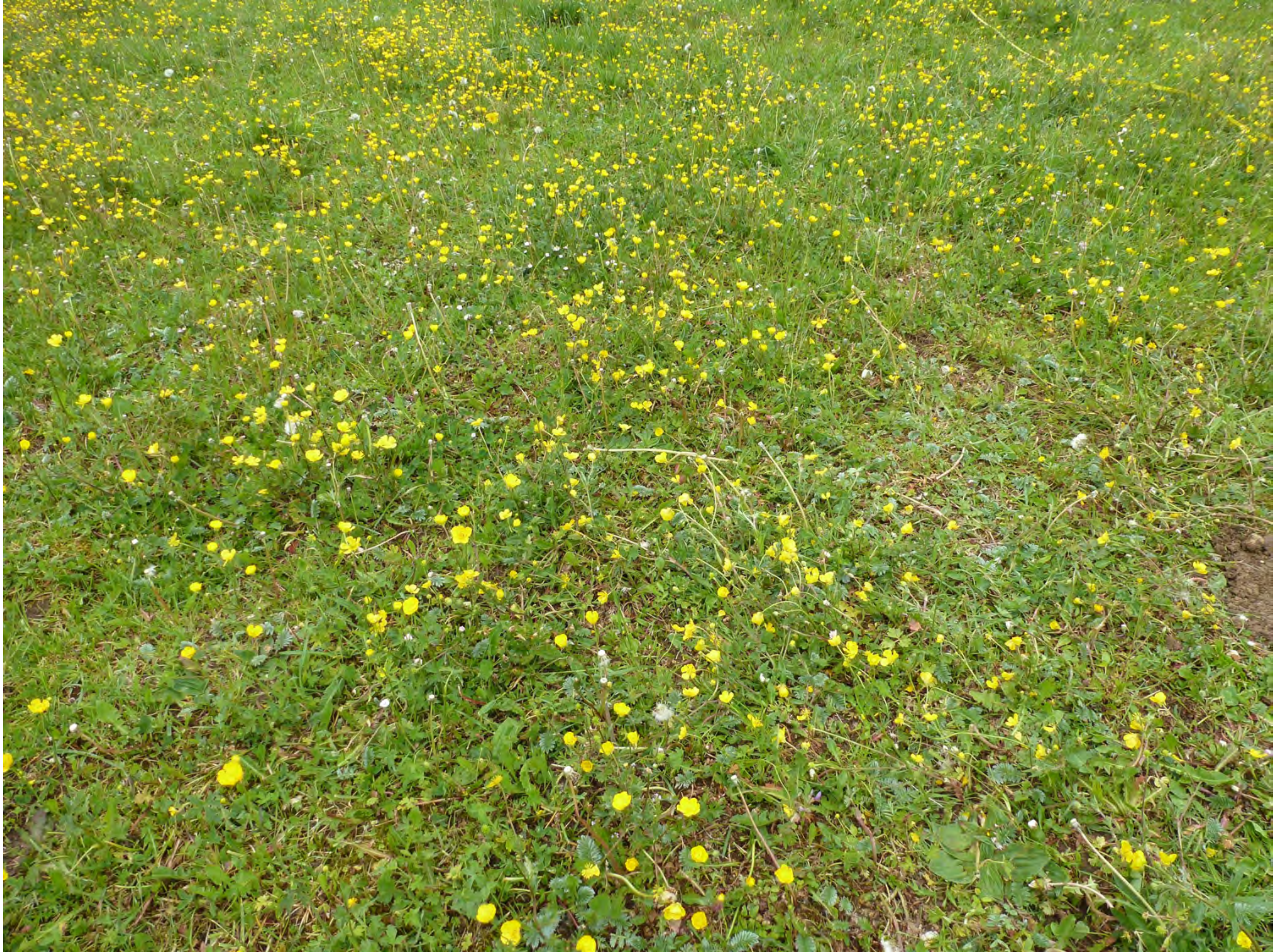

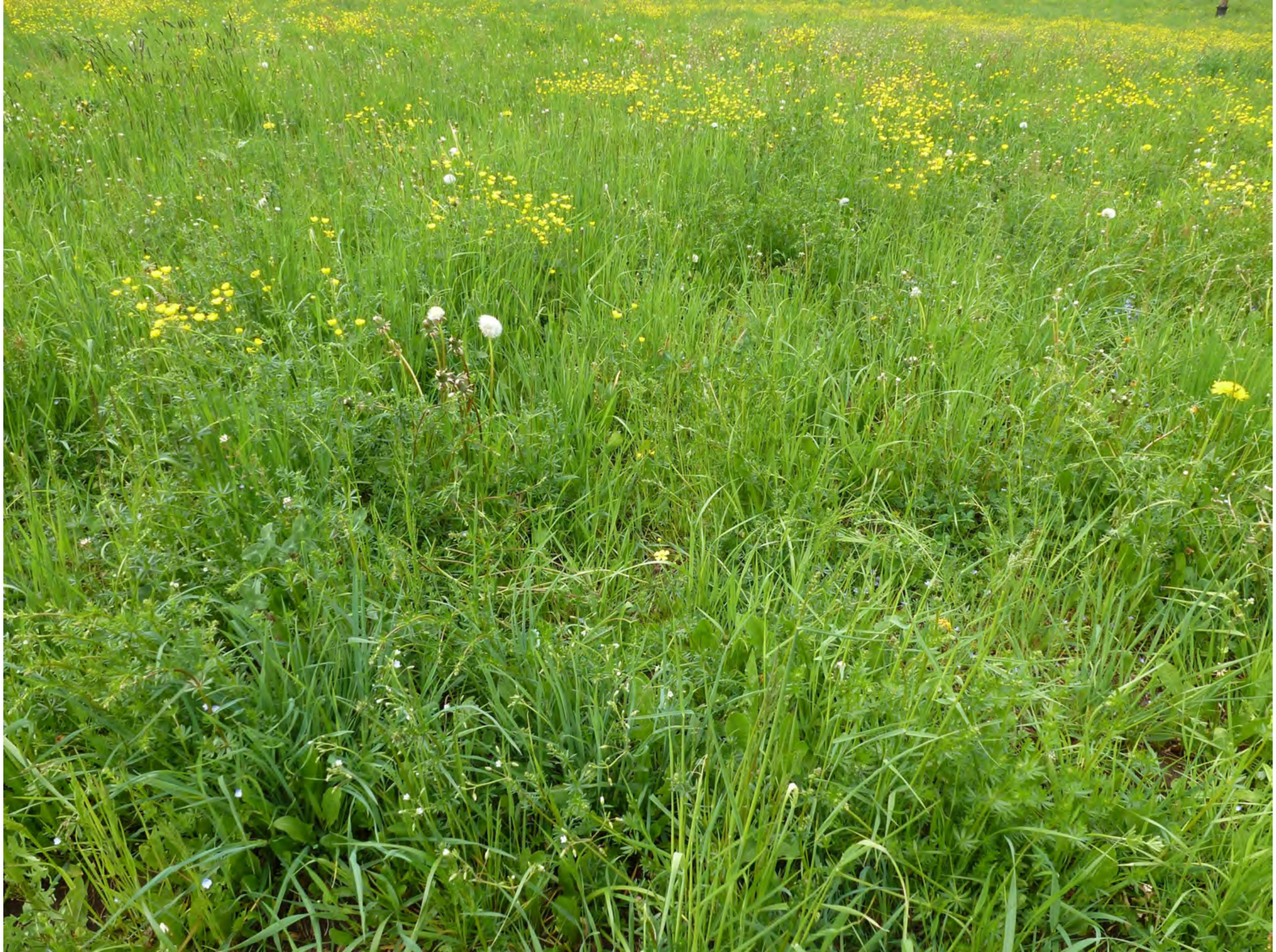

15

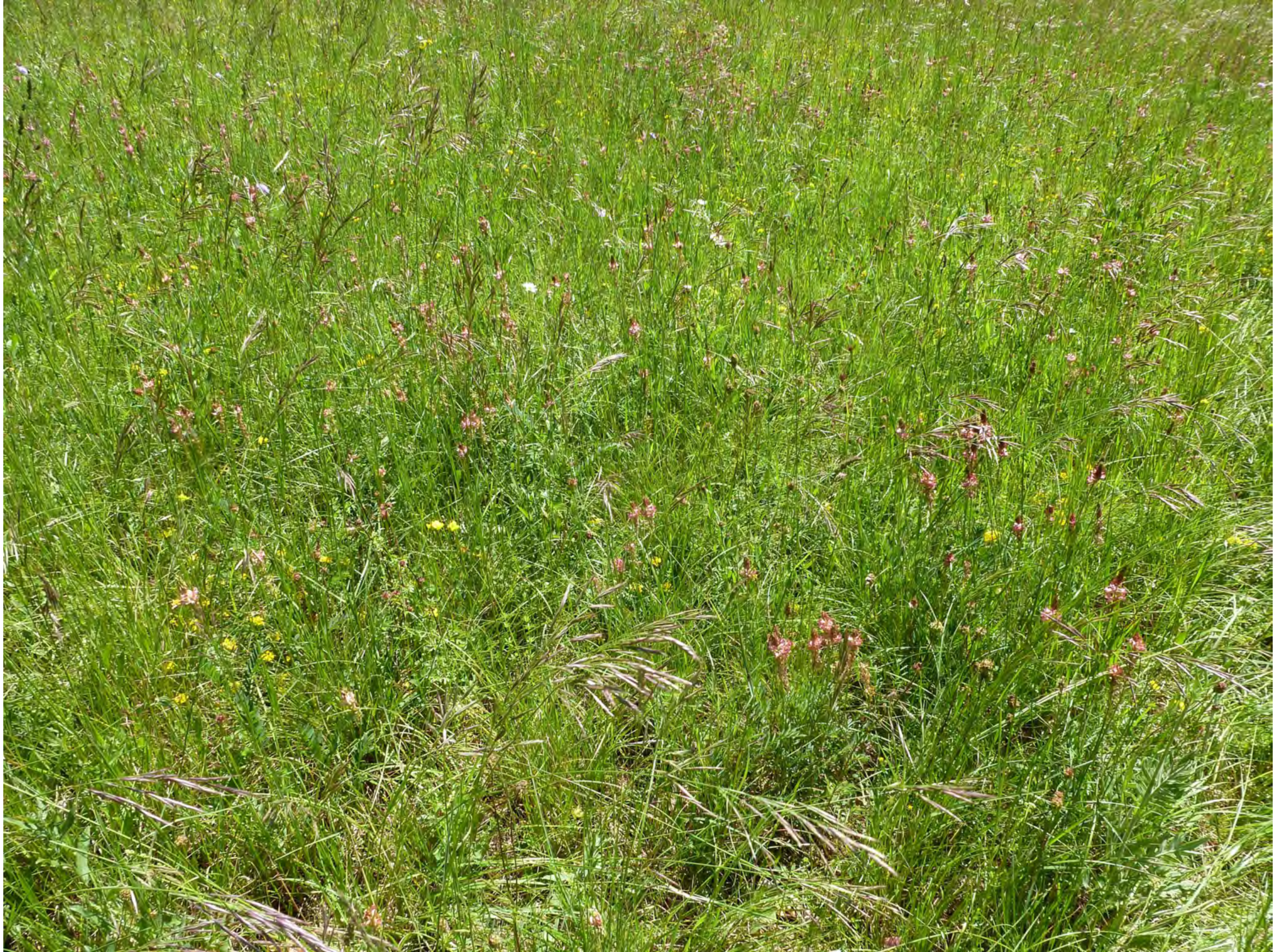

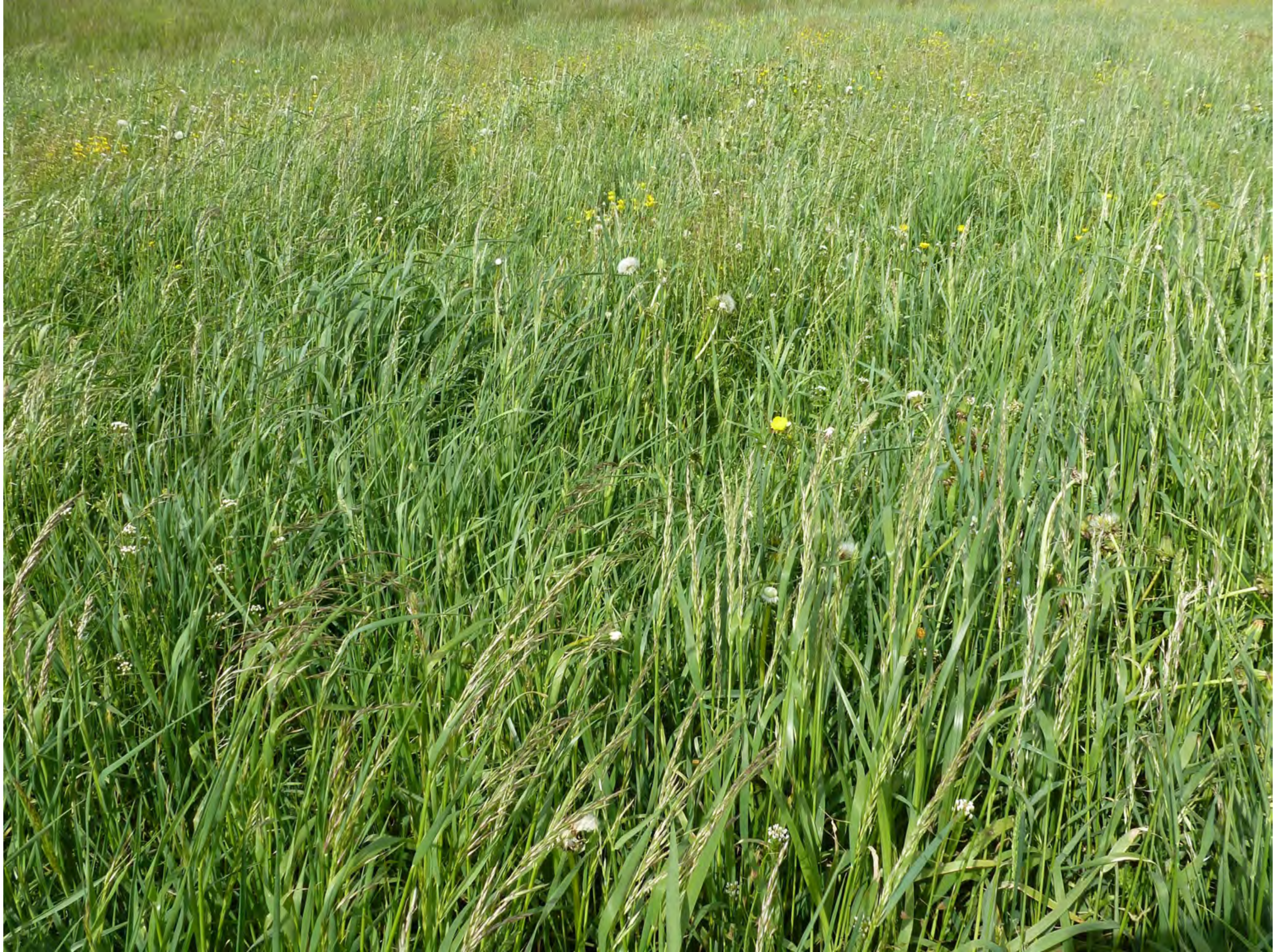

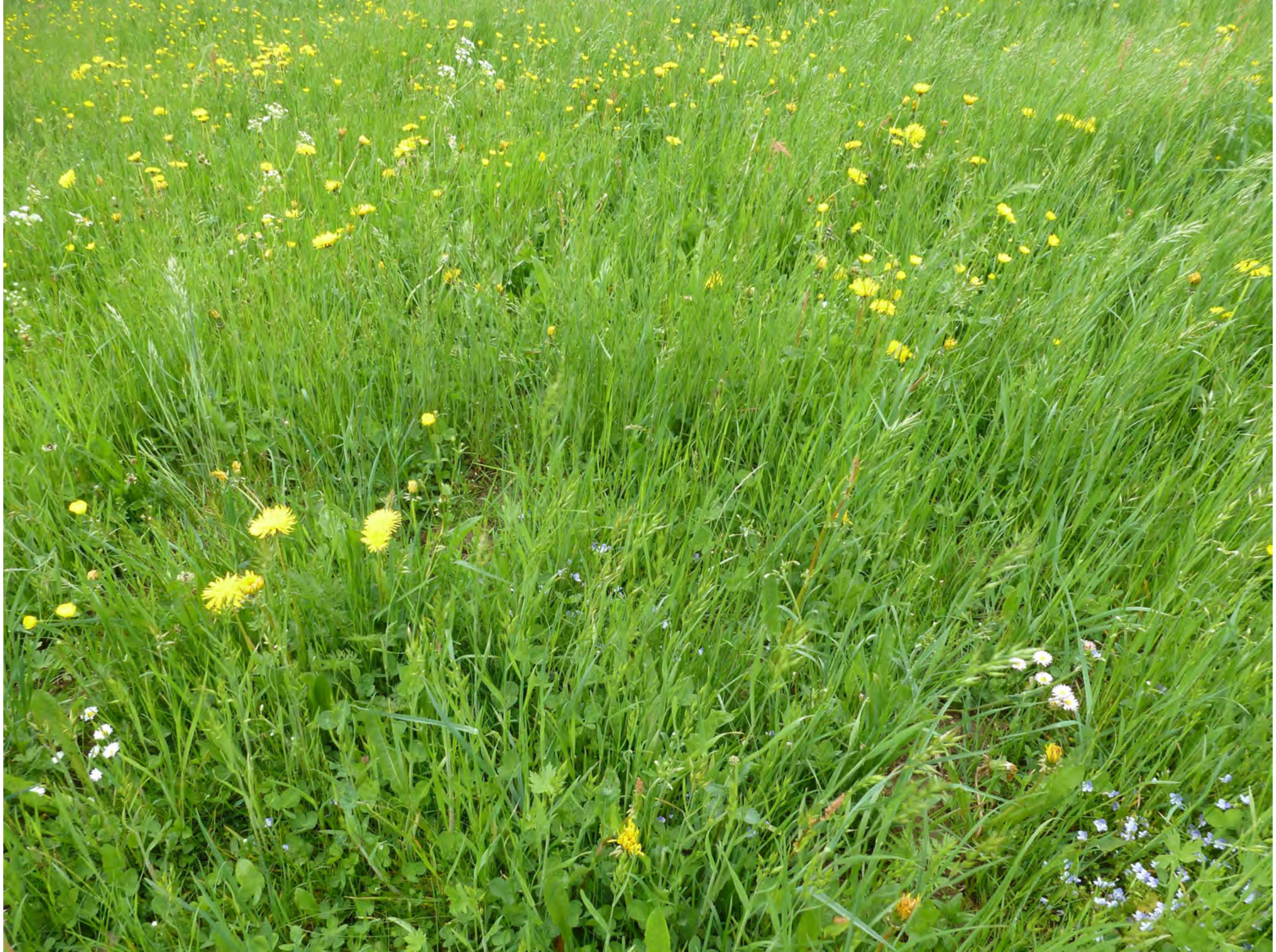

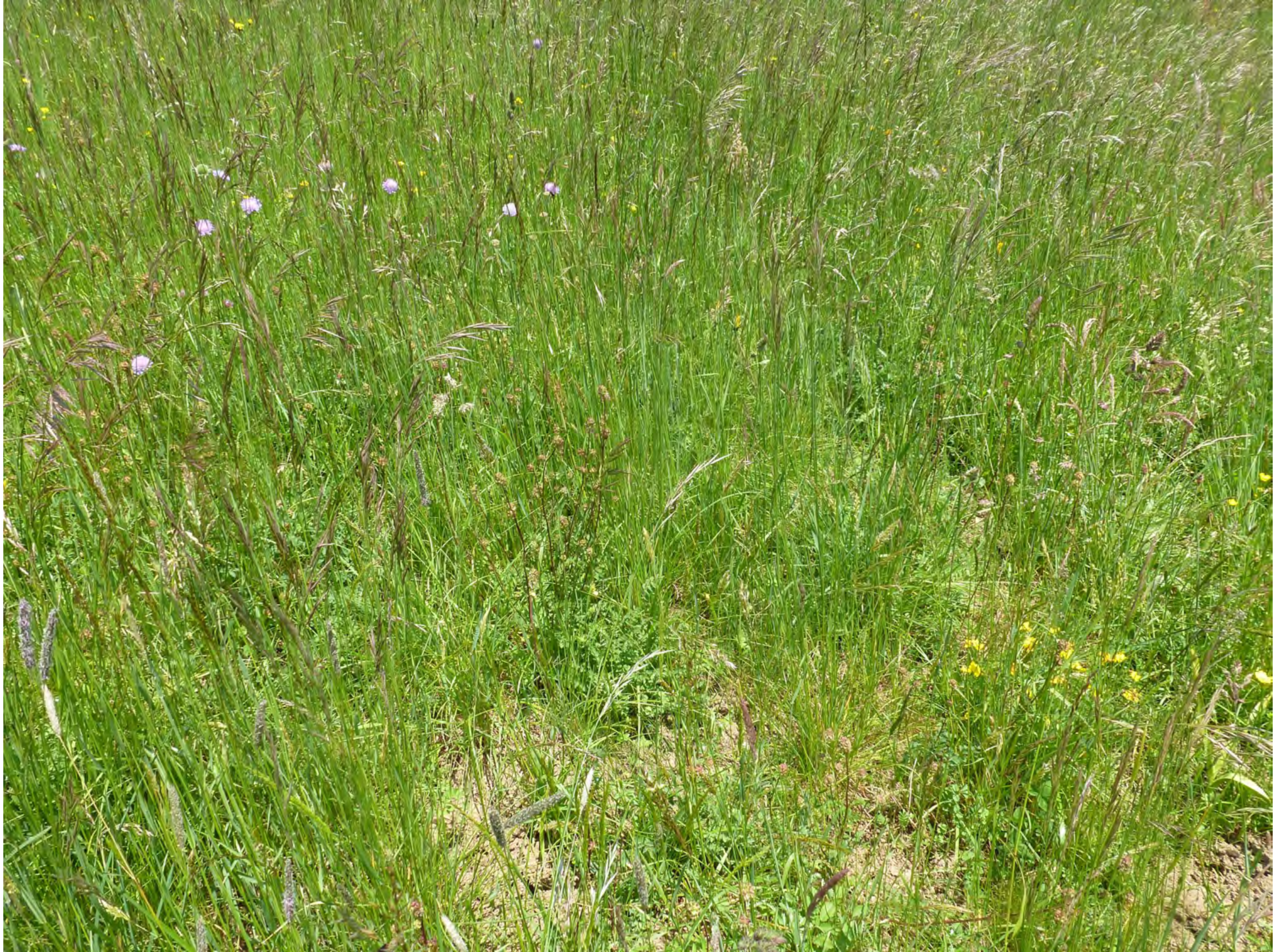

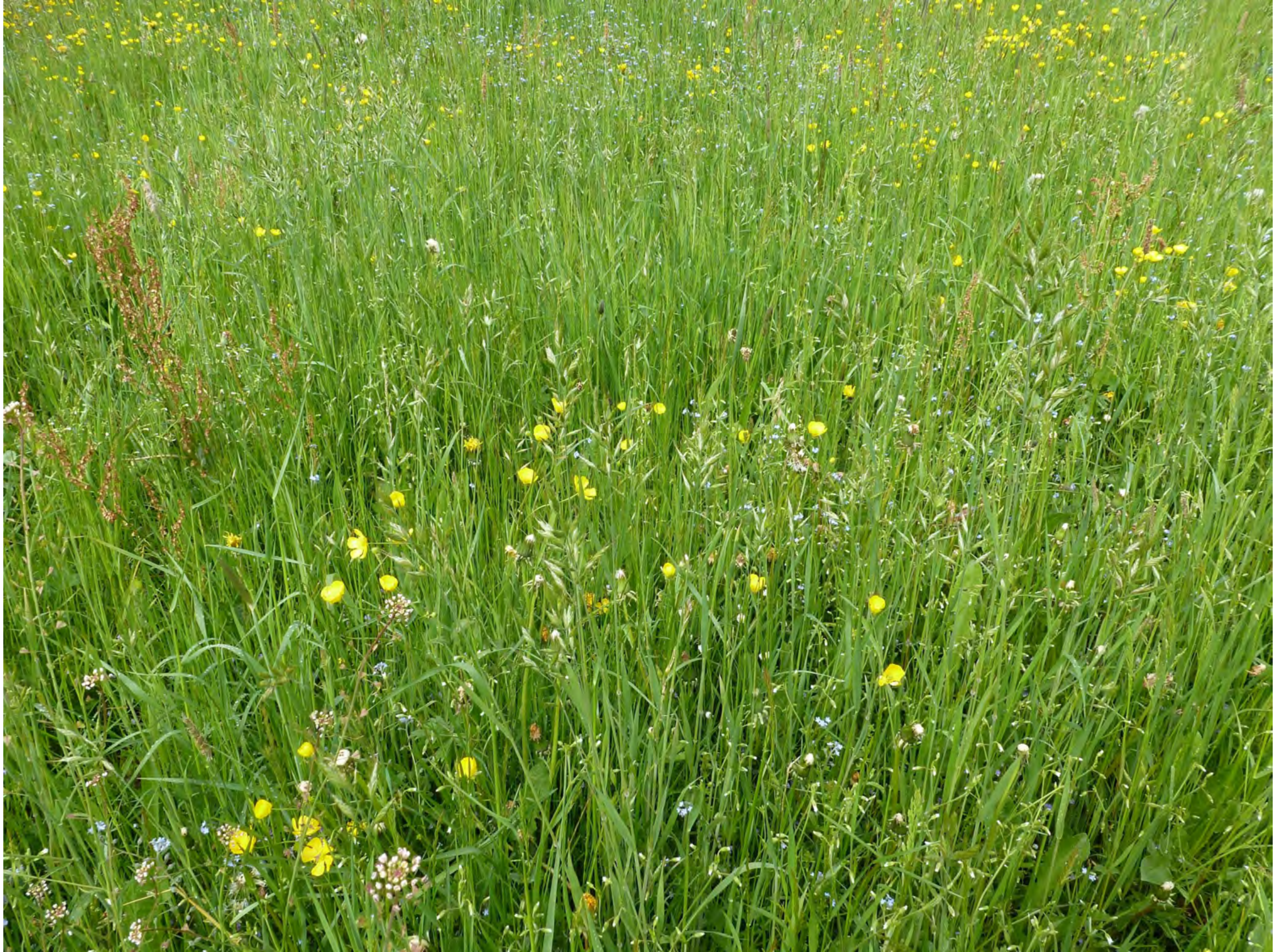

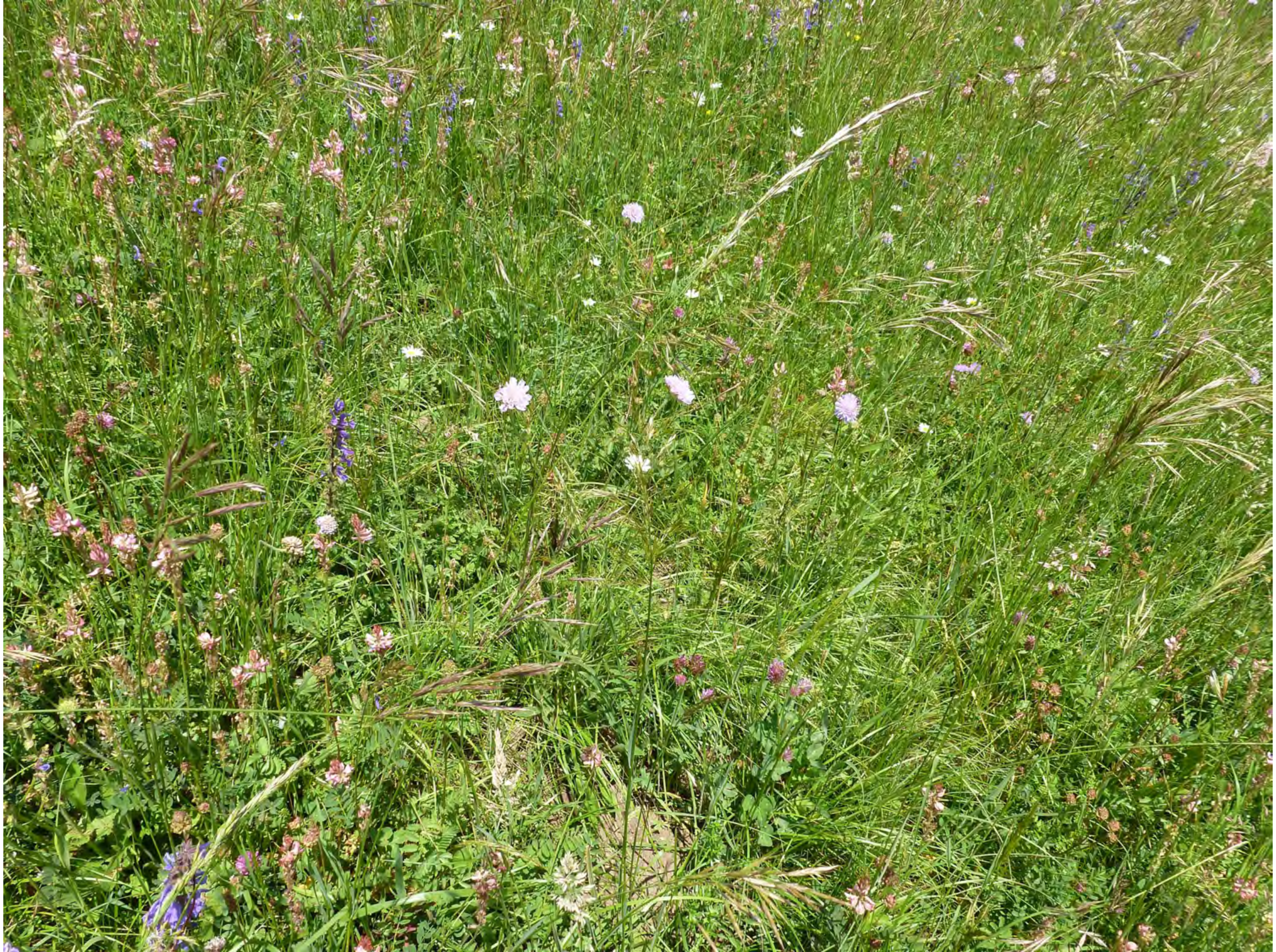

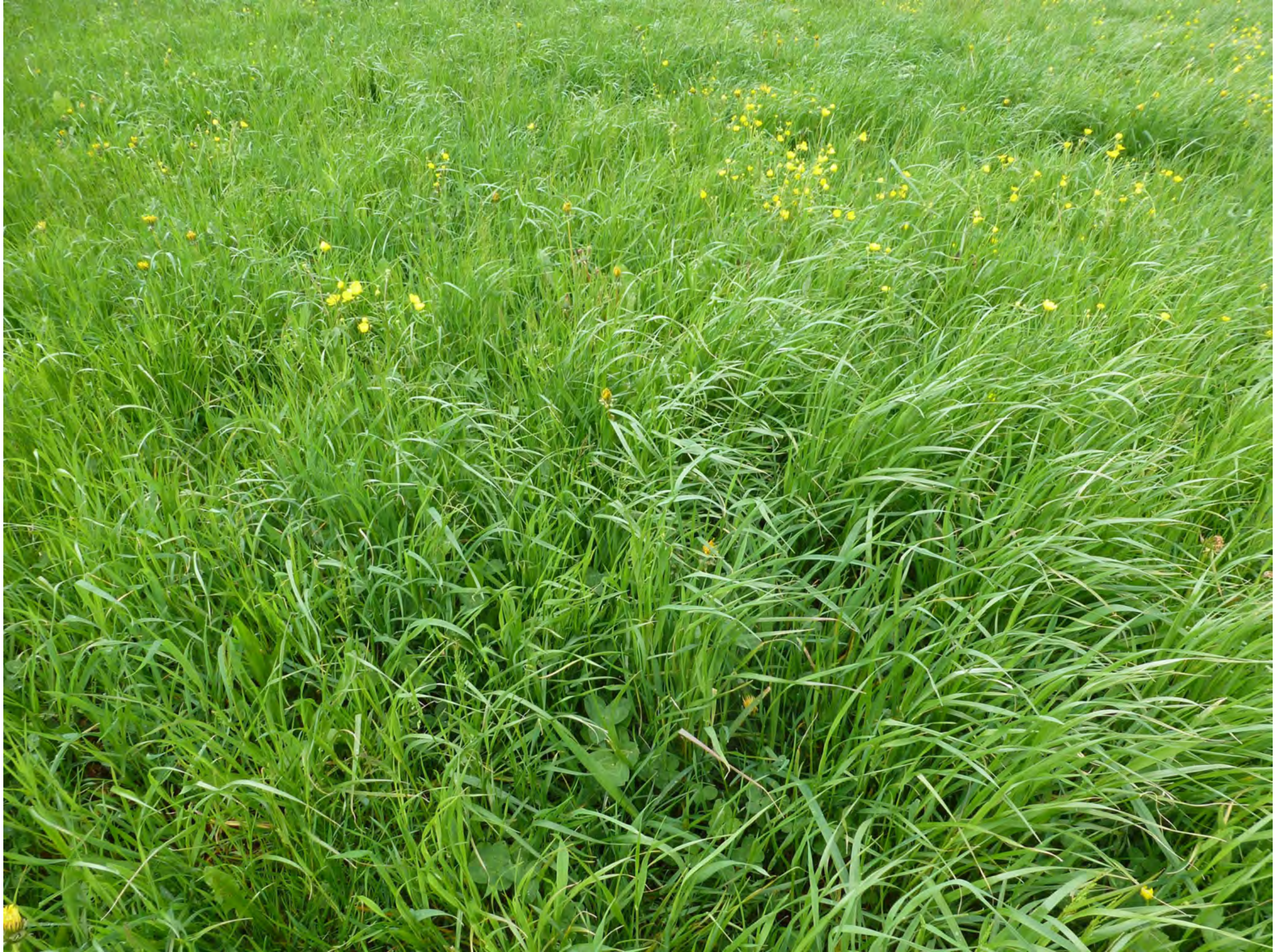

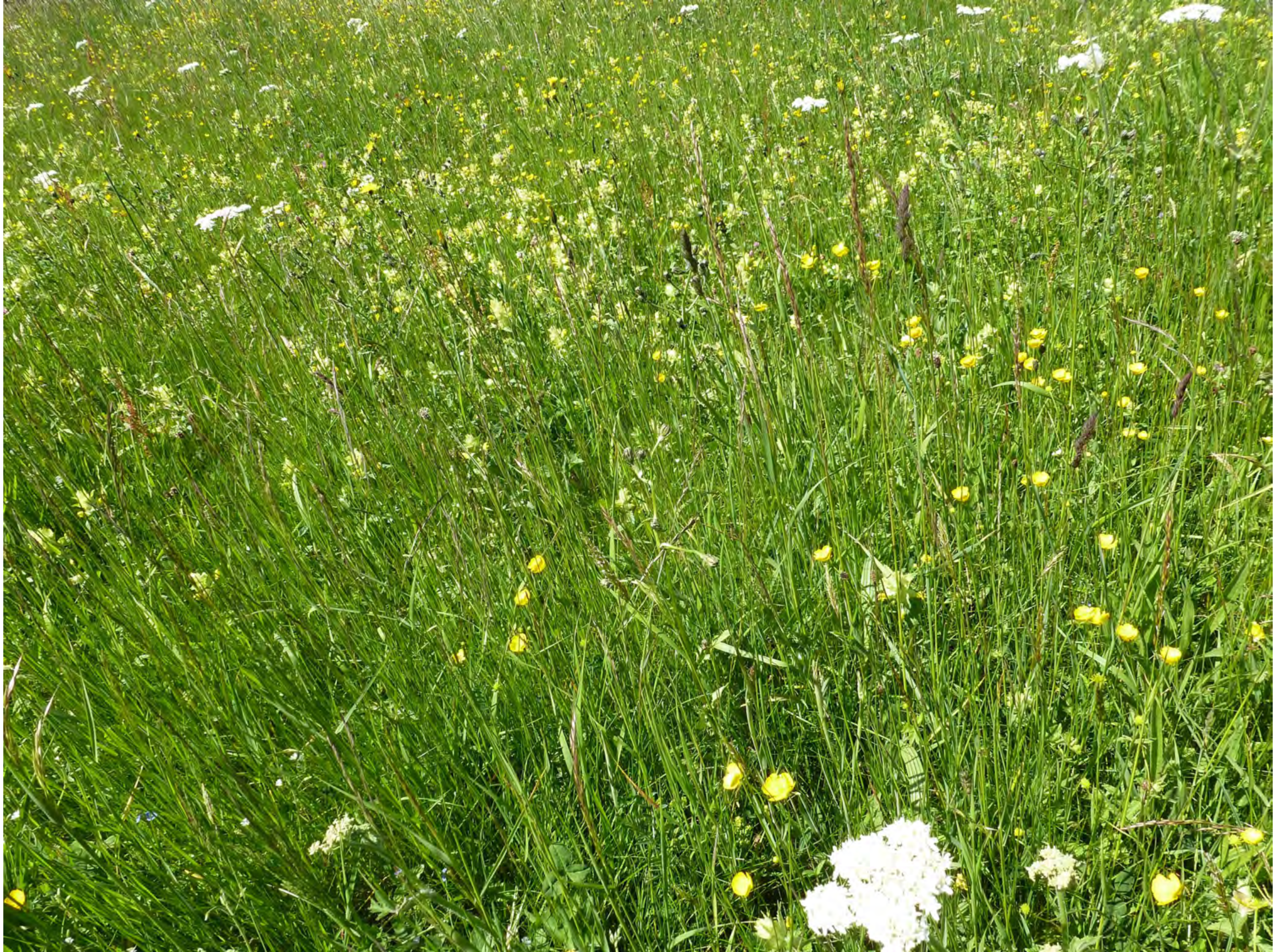

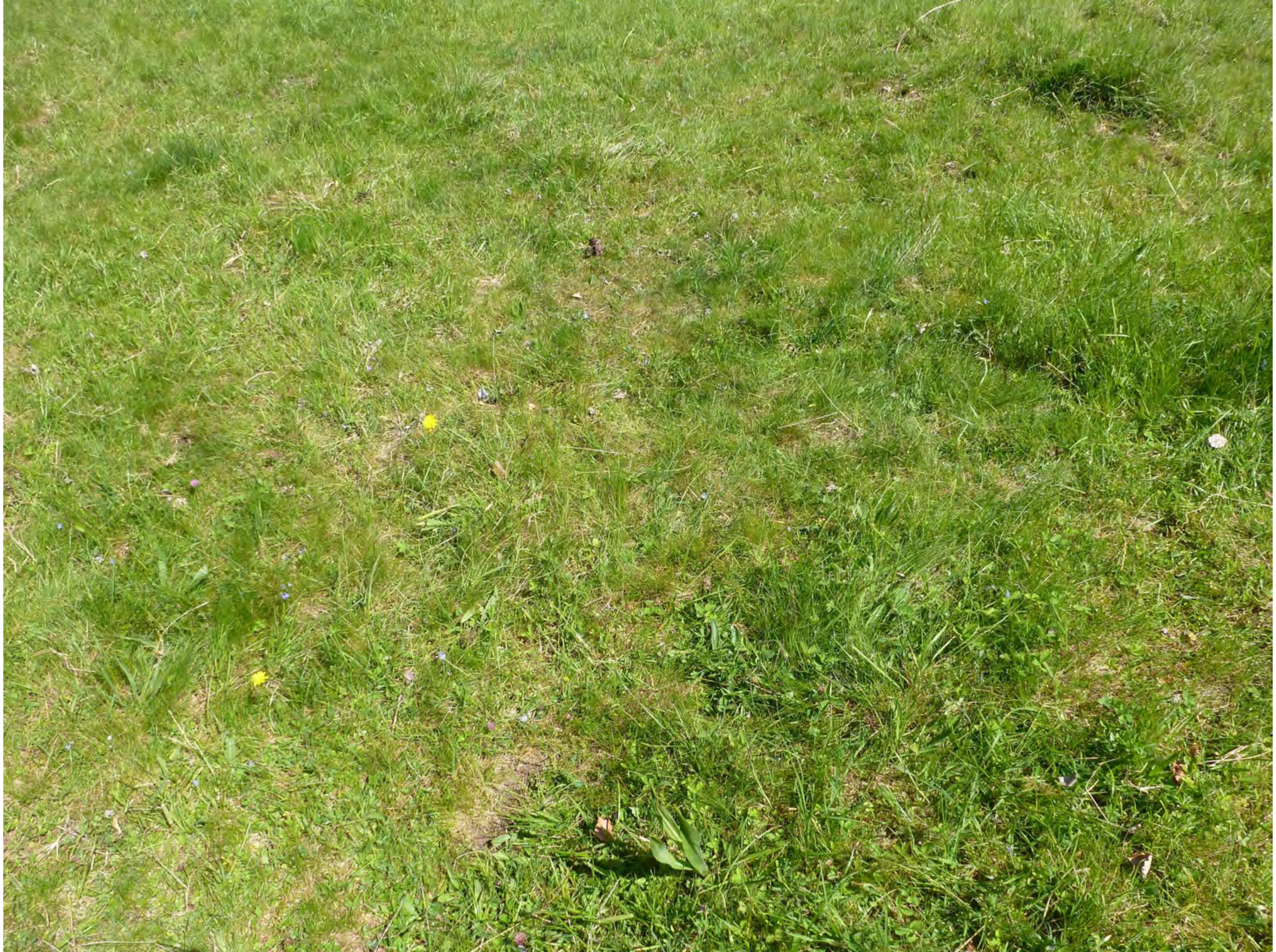

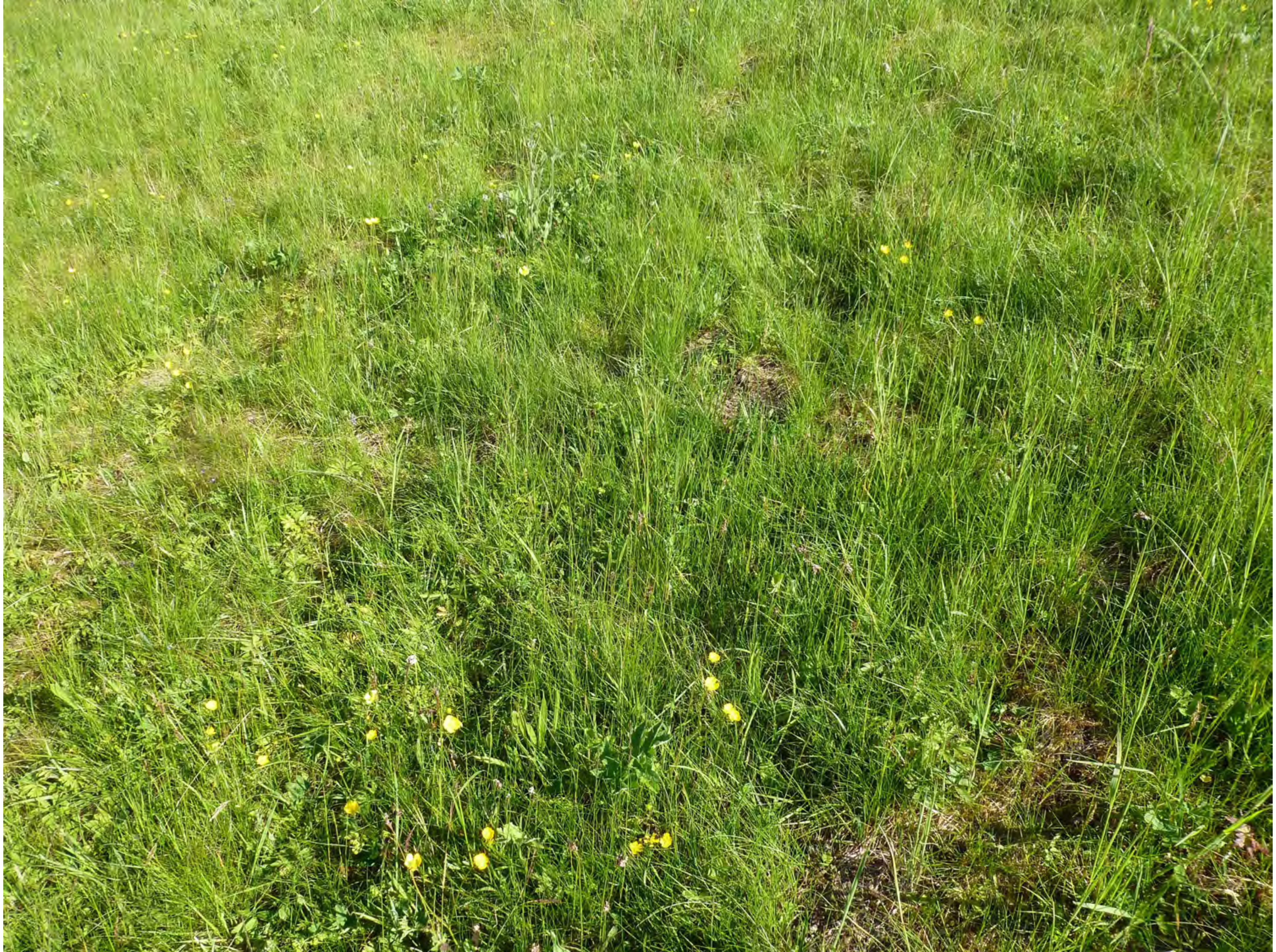

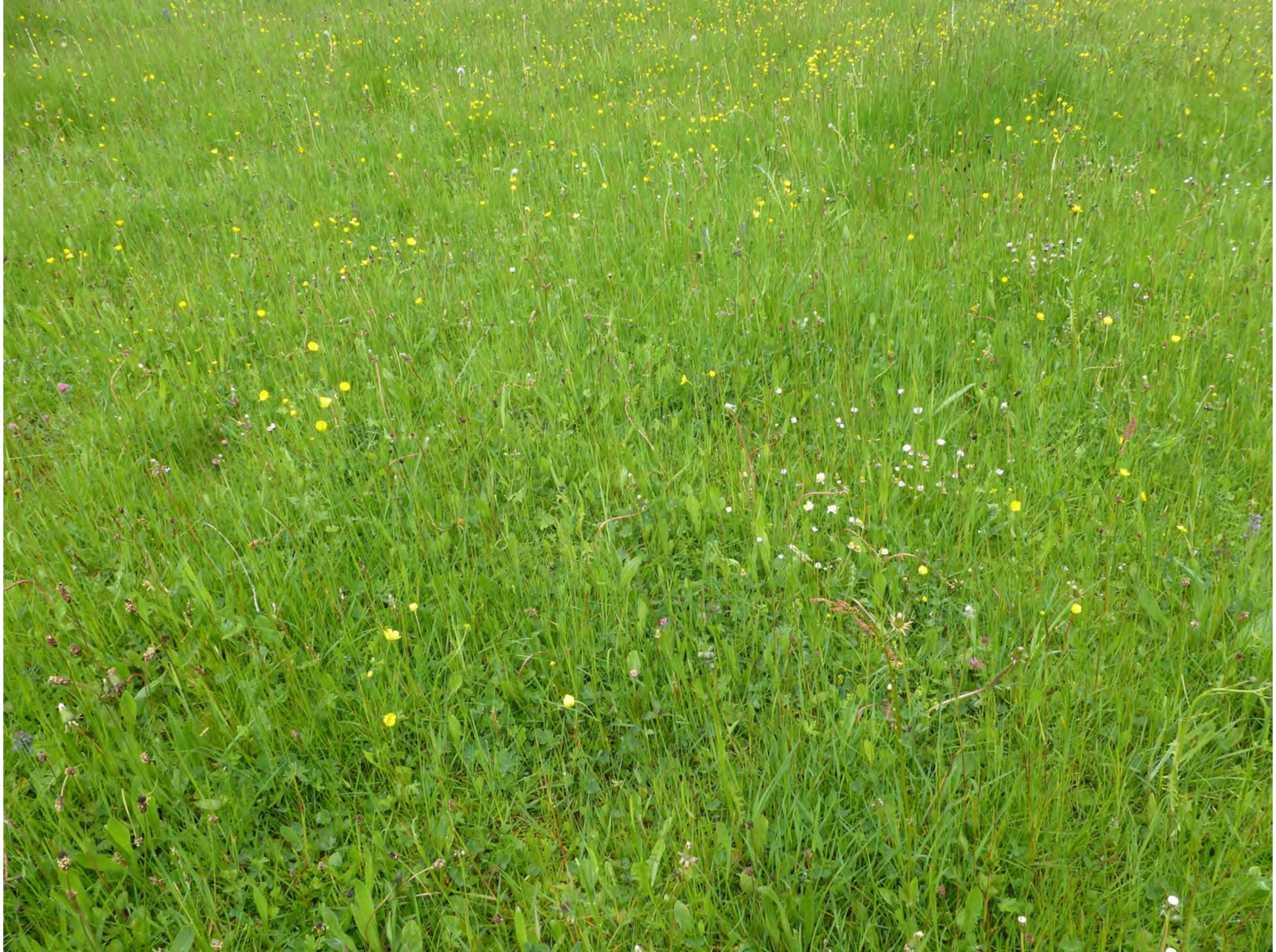

26

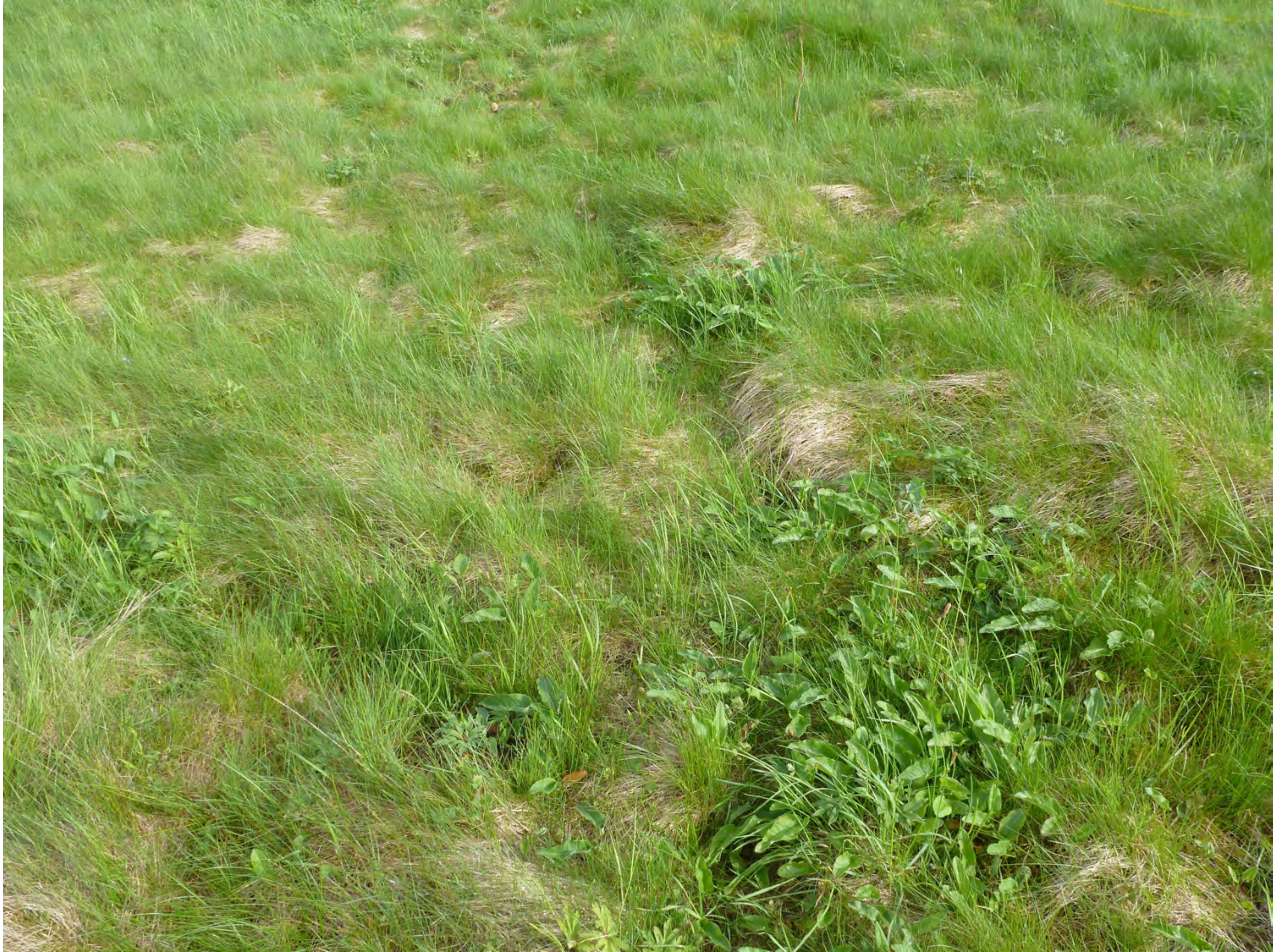

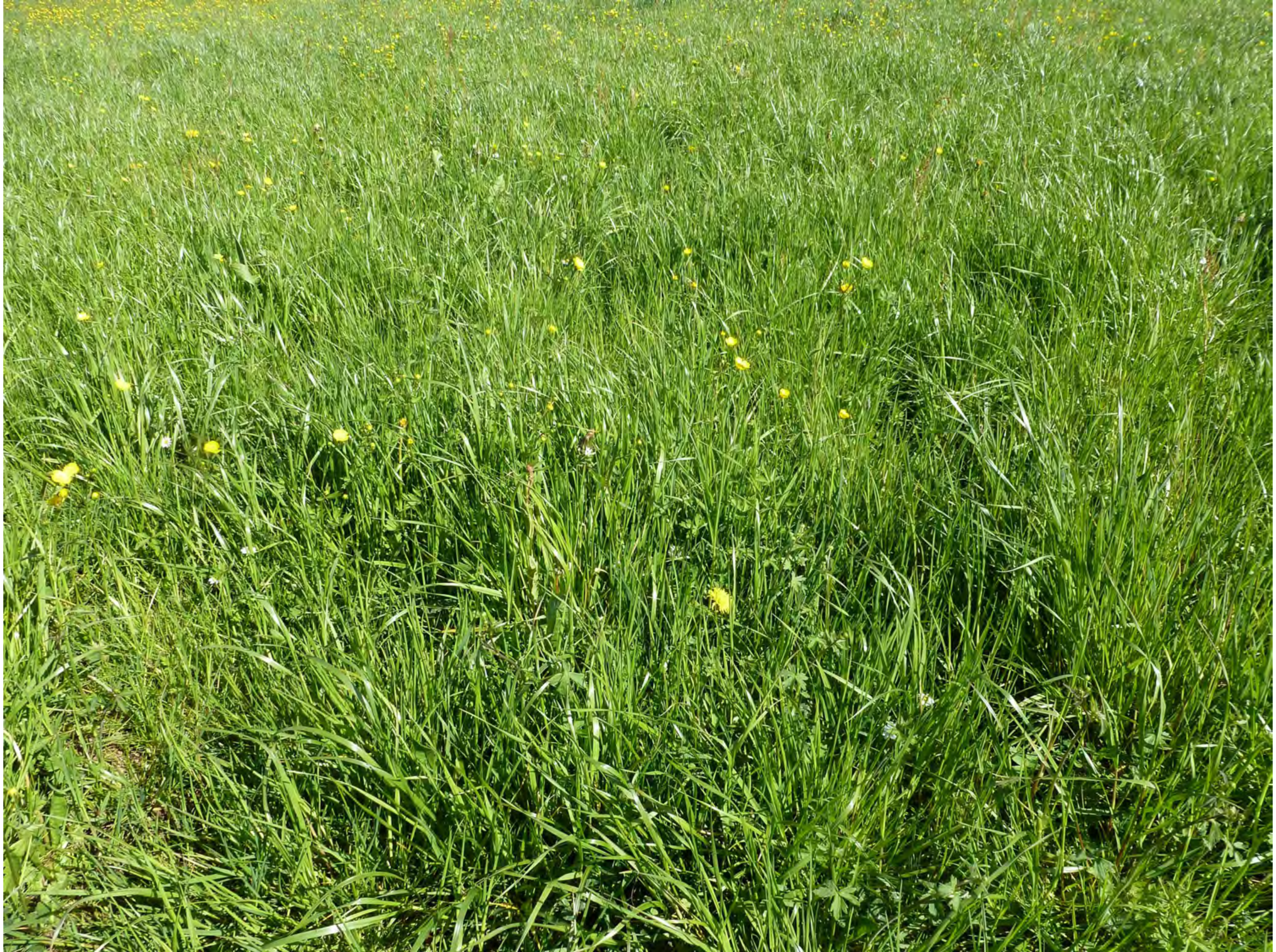

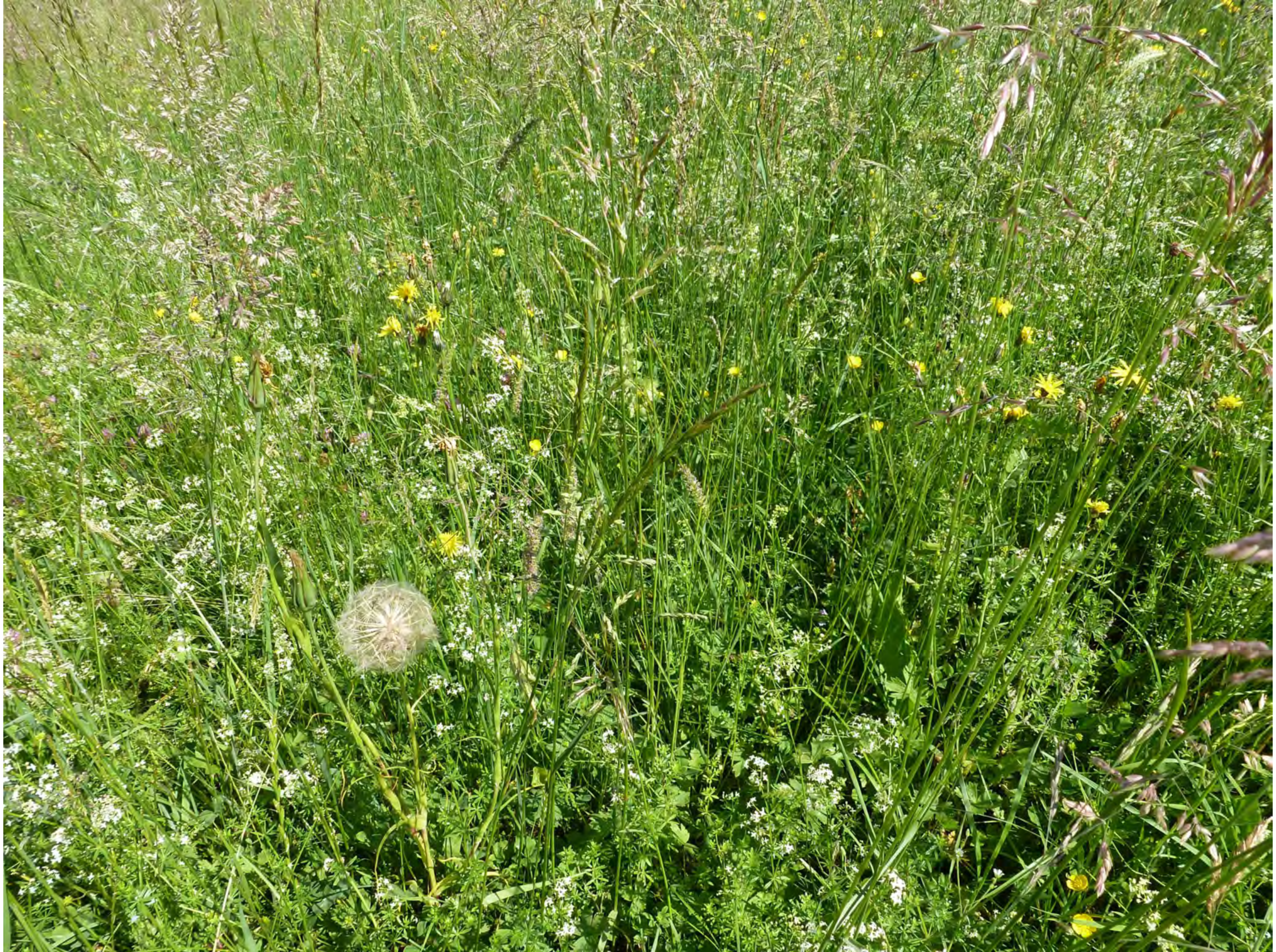

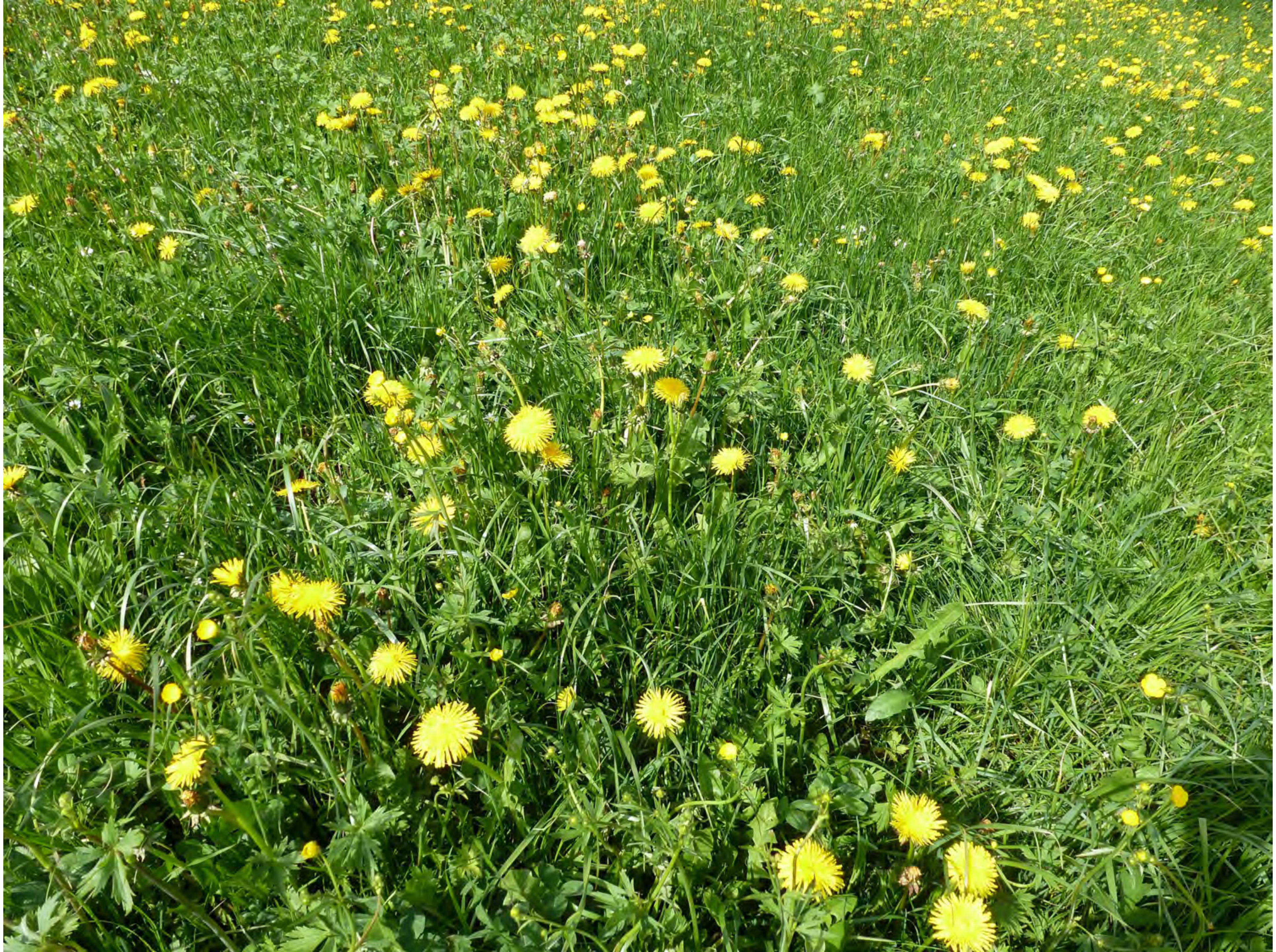

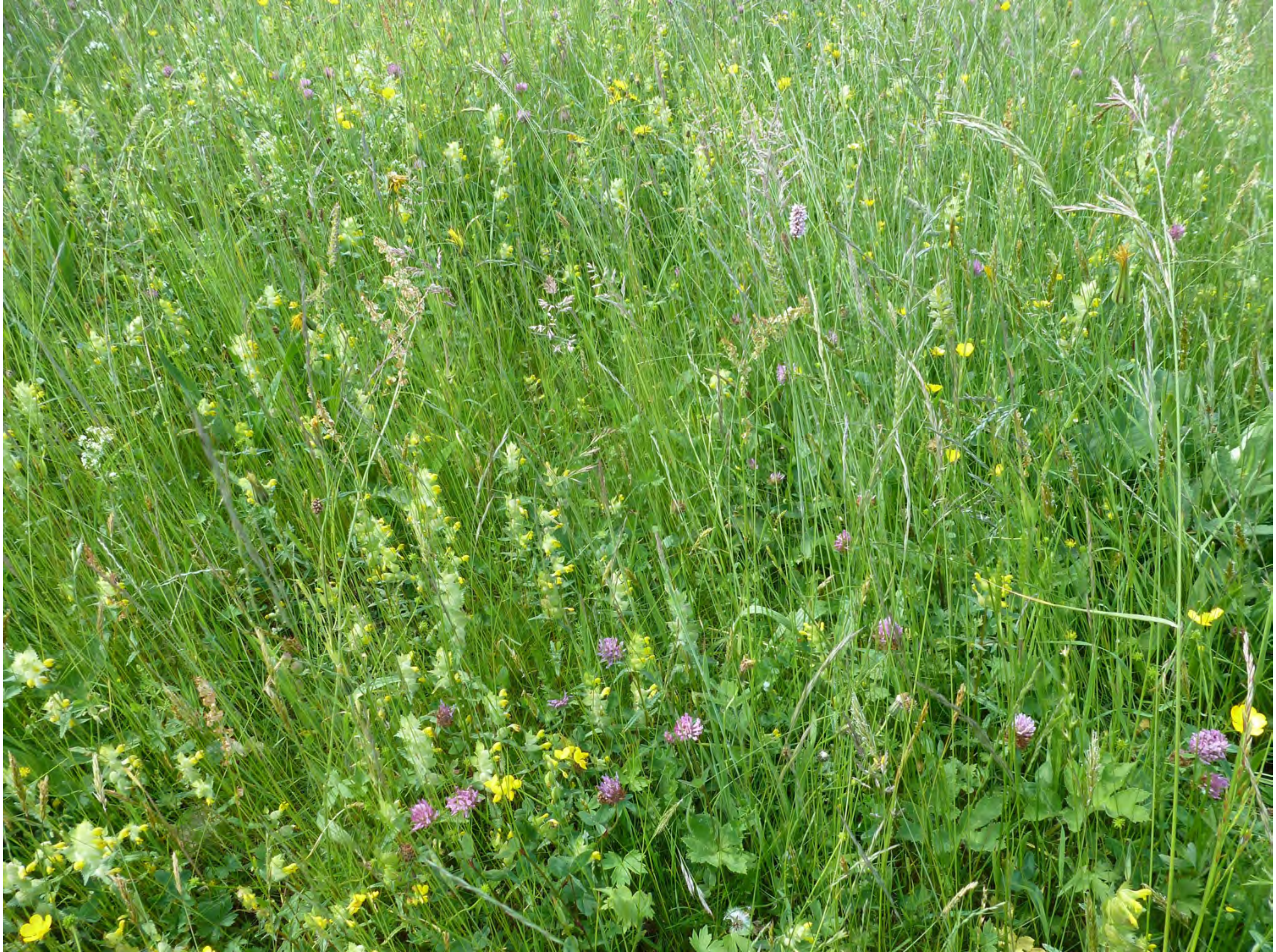

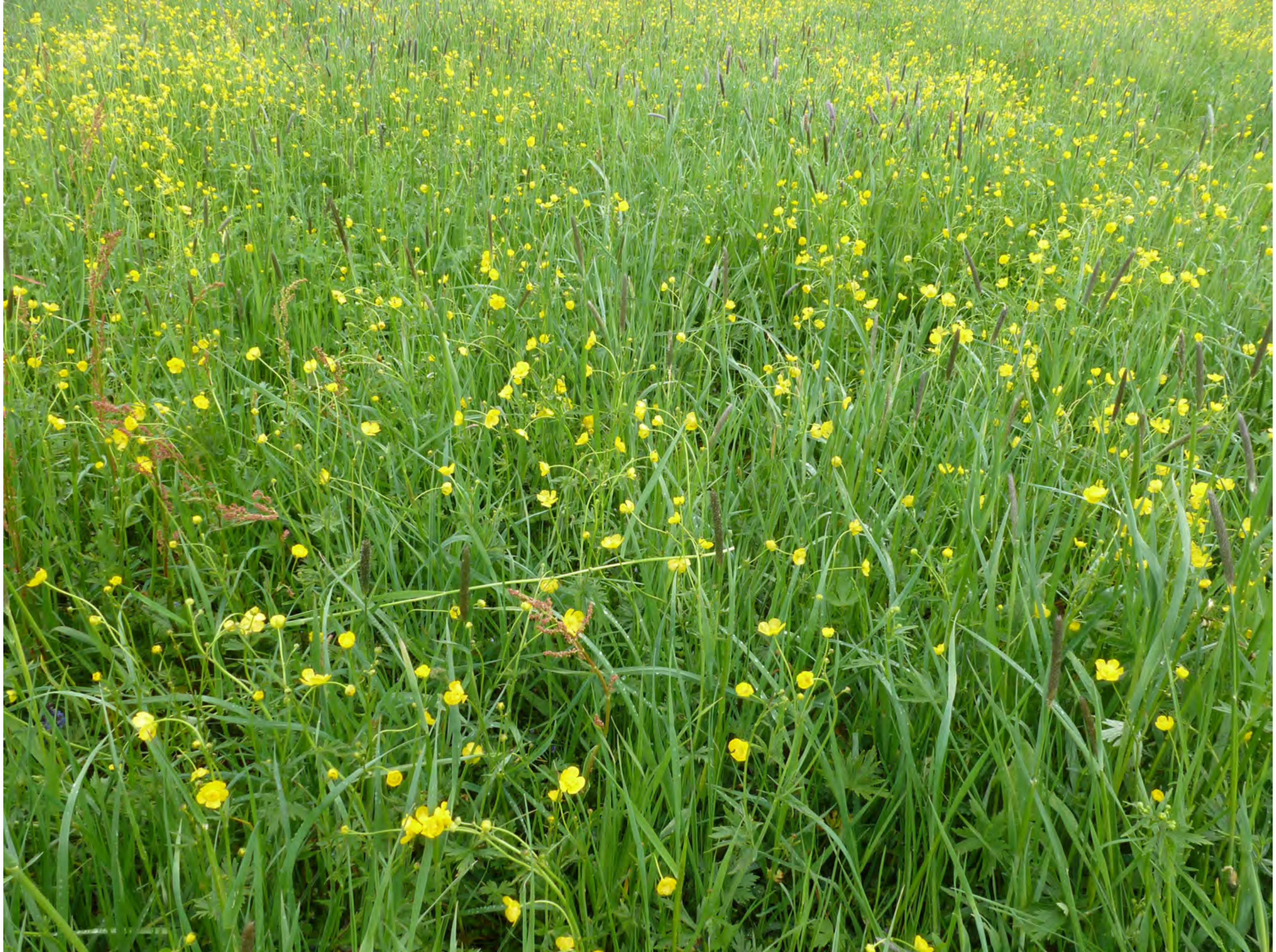

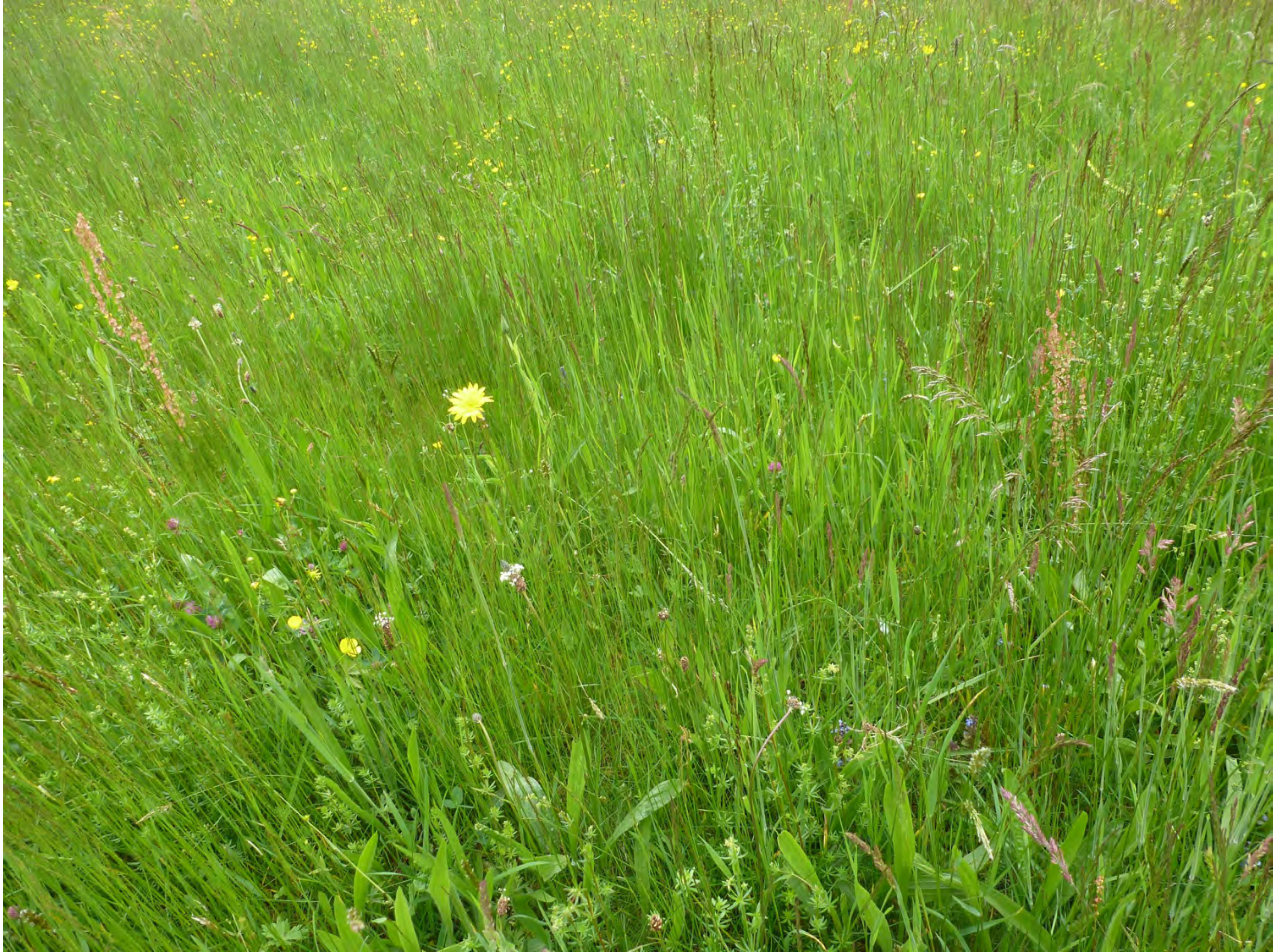

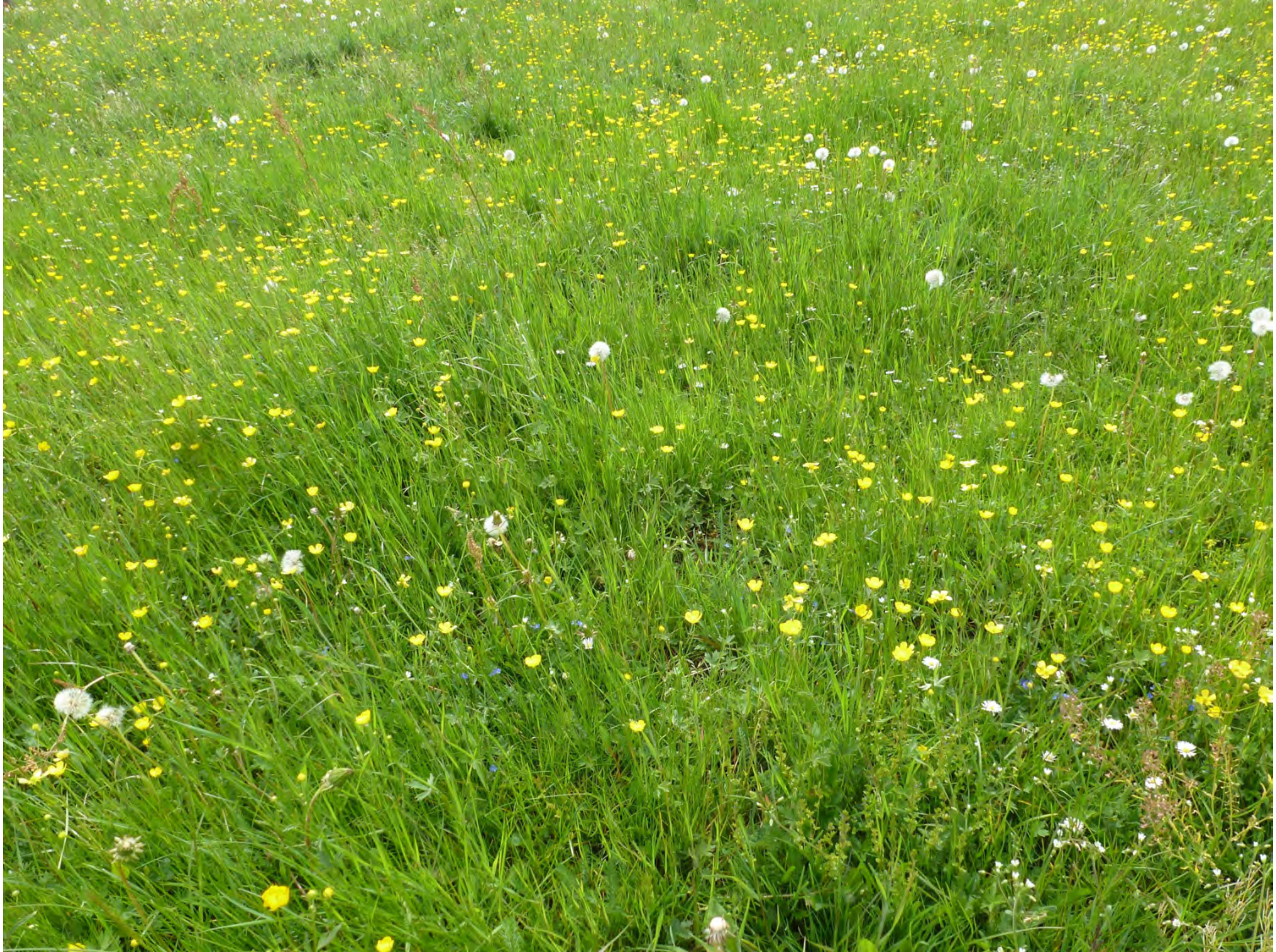

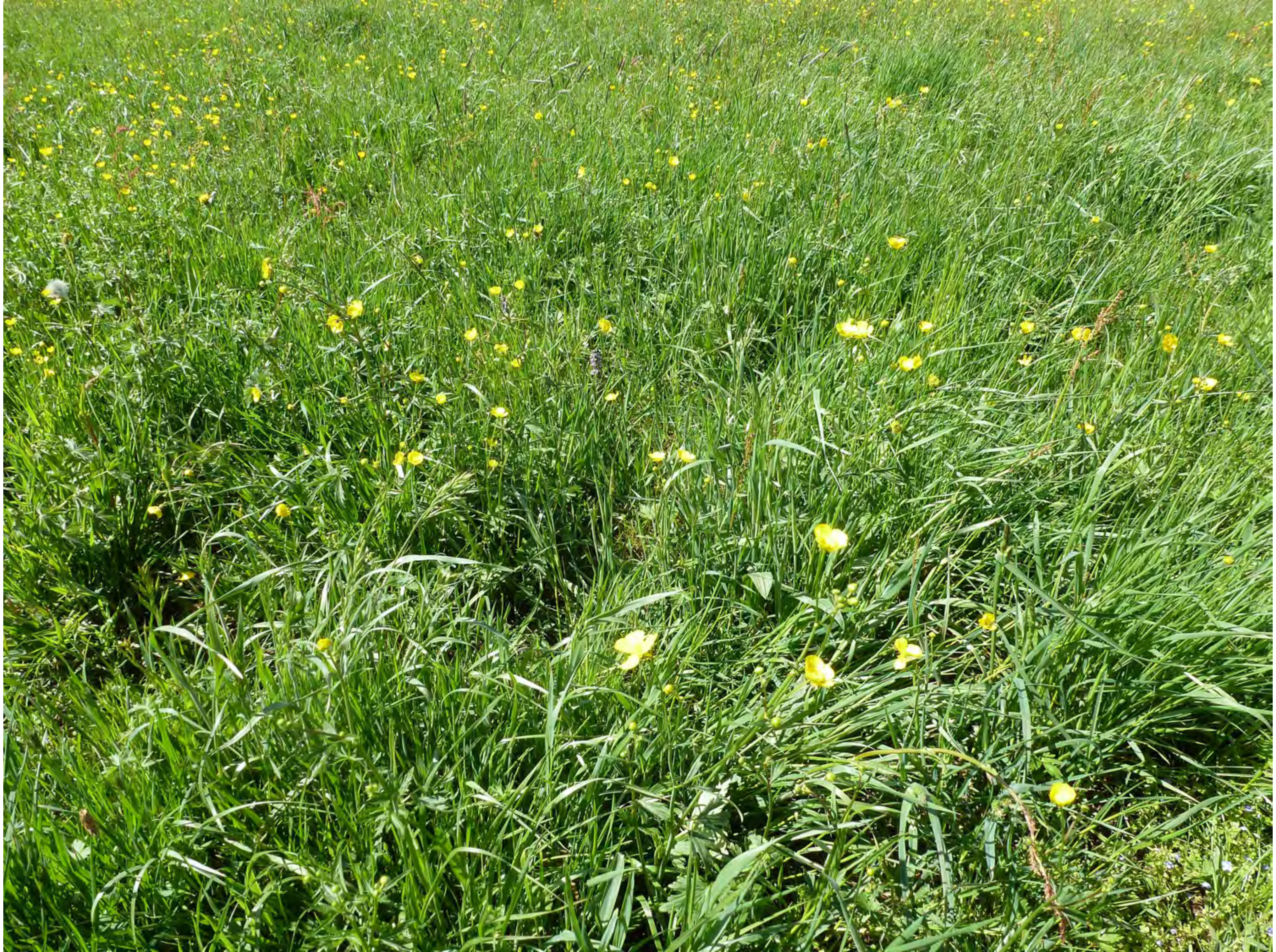

35

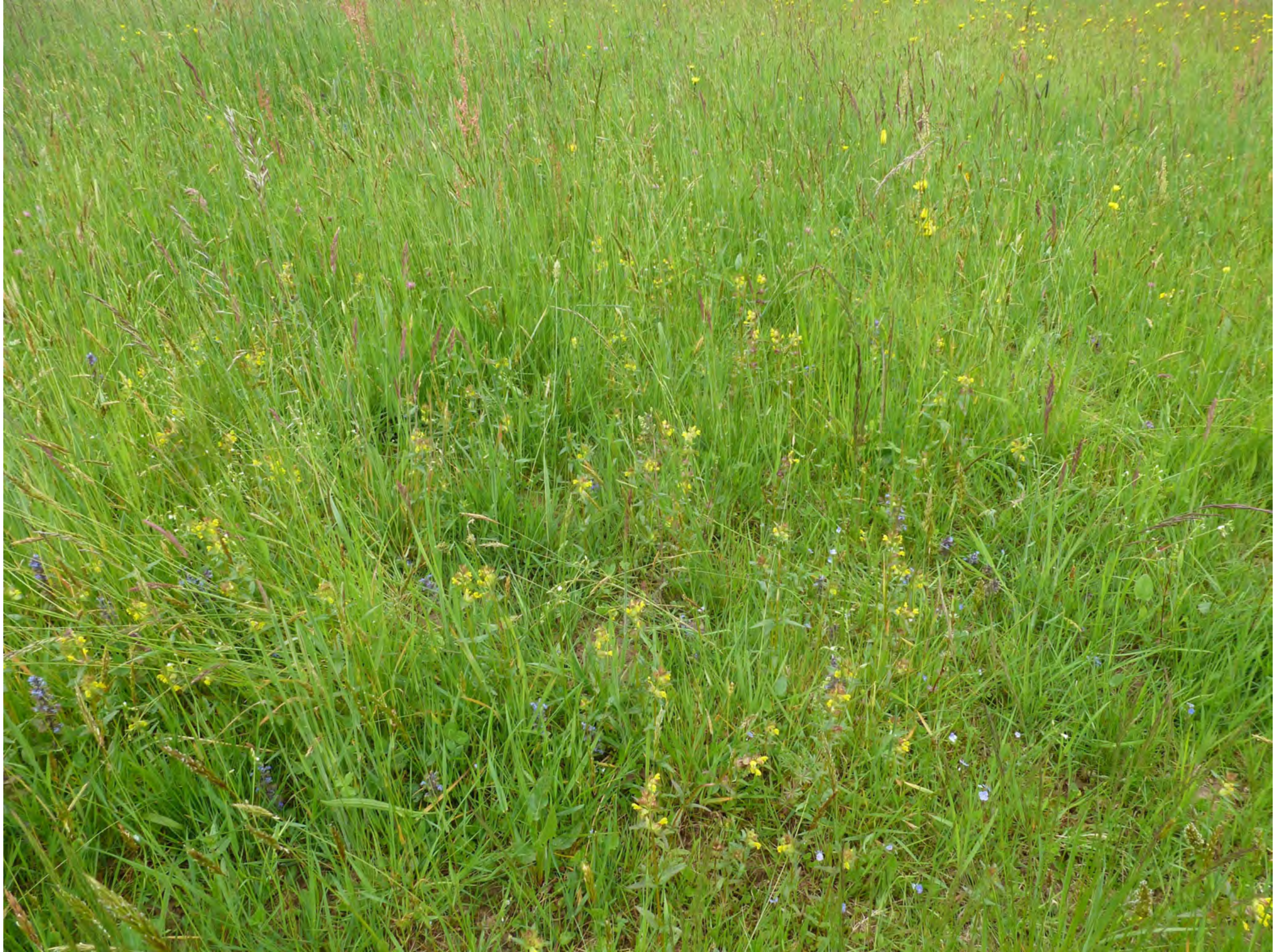

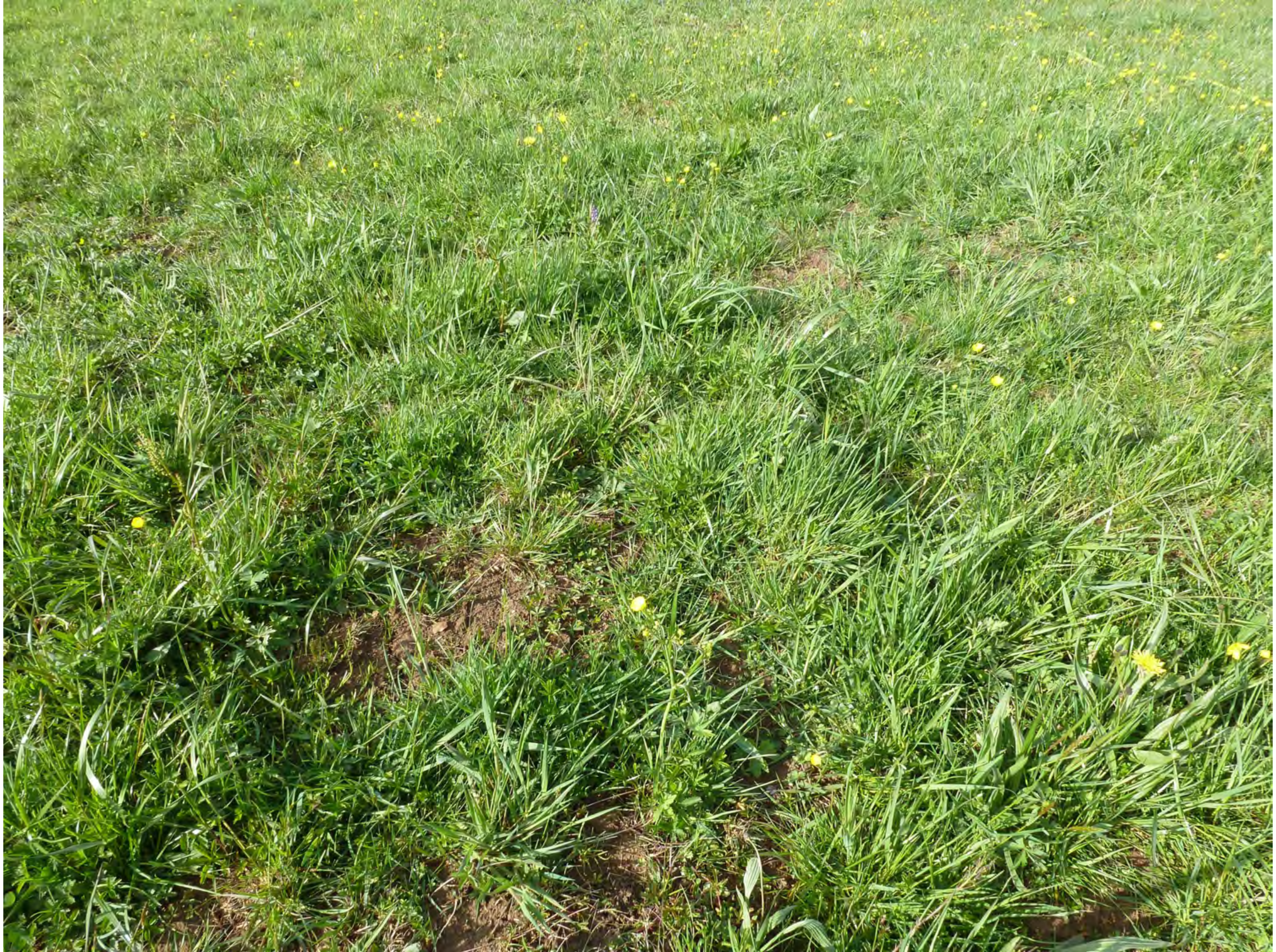

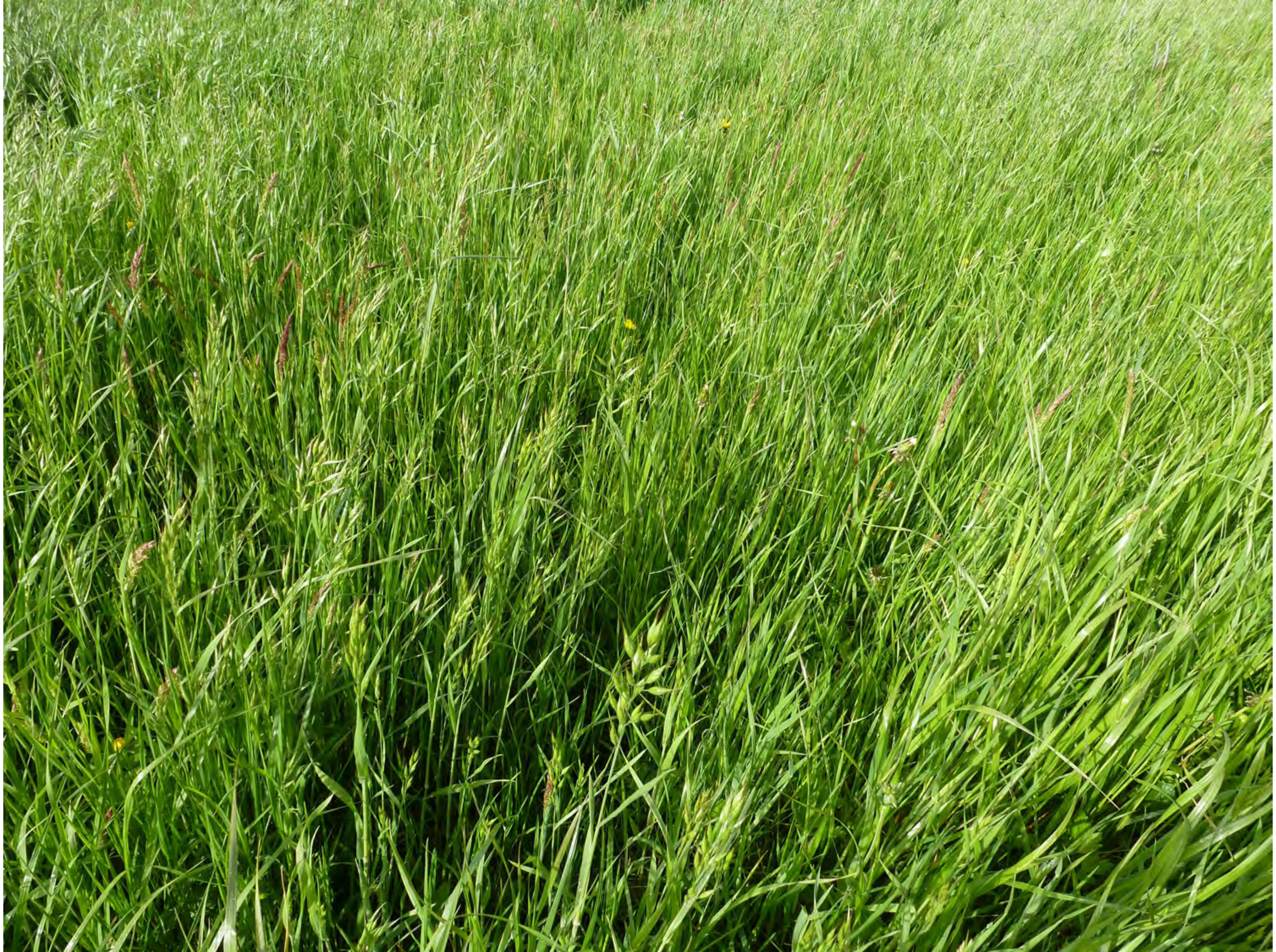

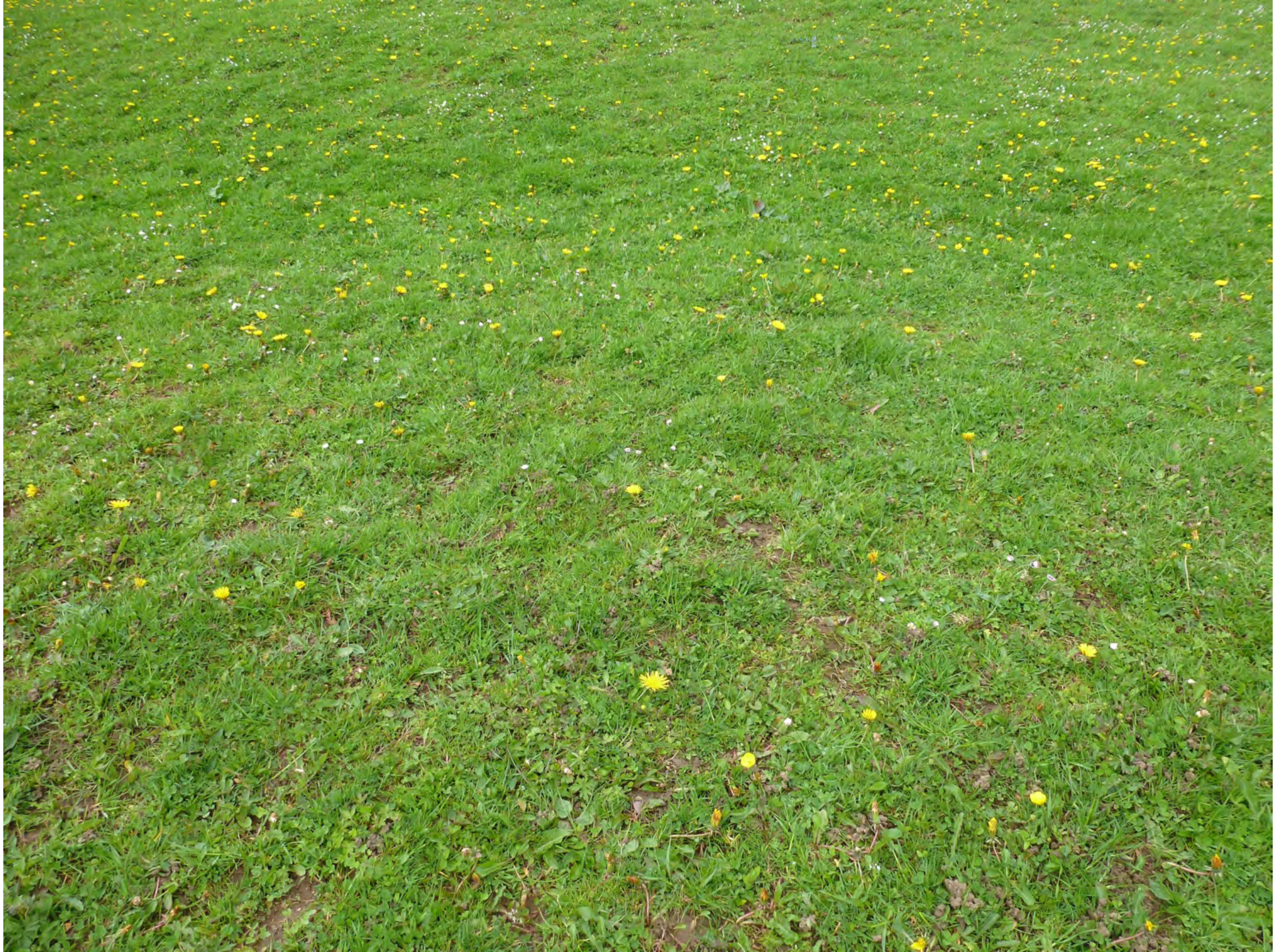

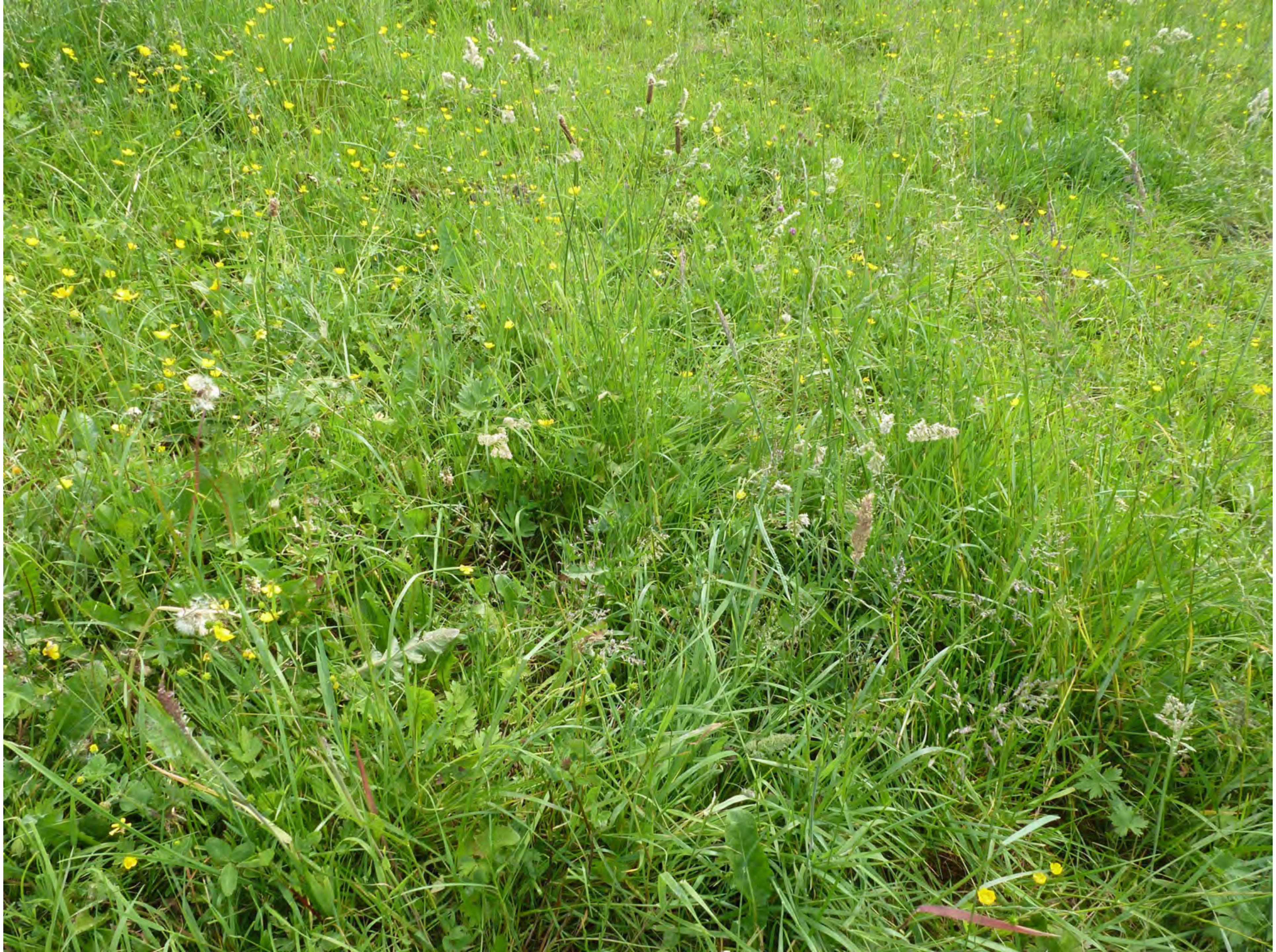

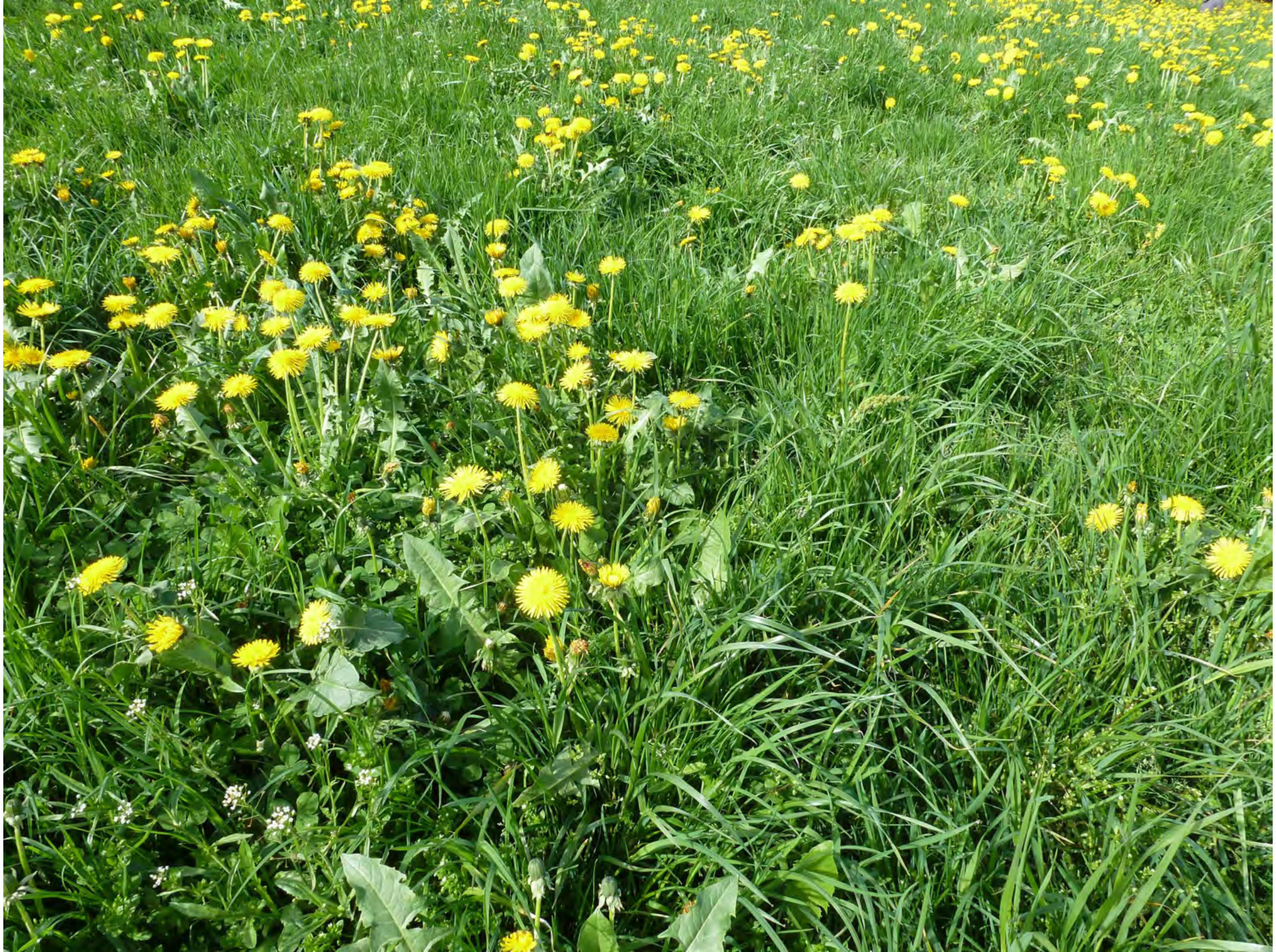

41

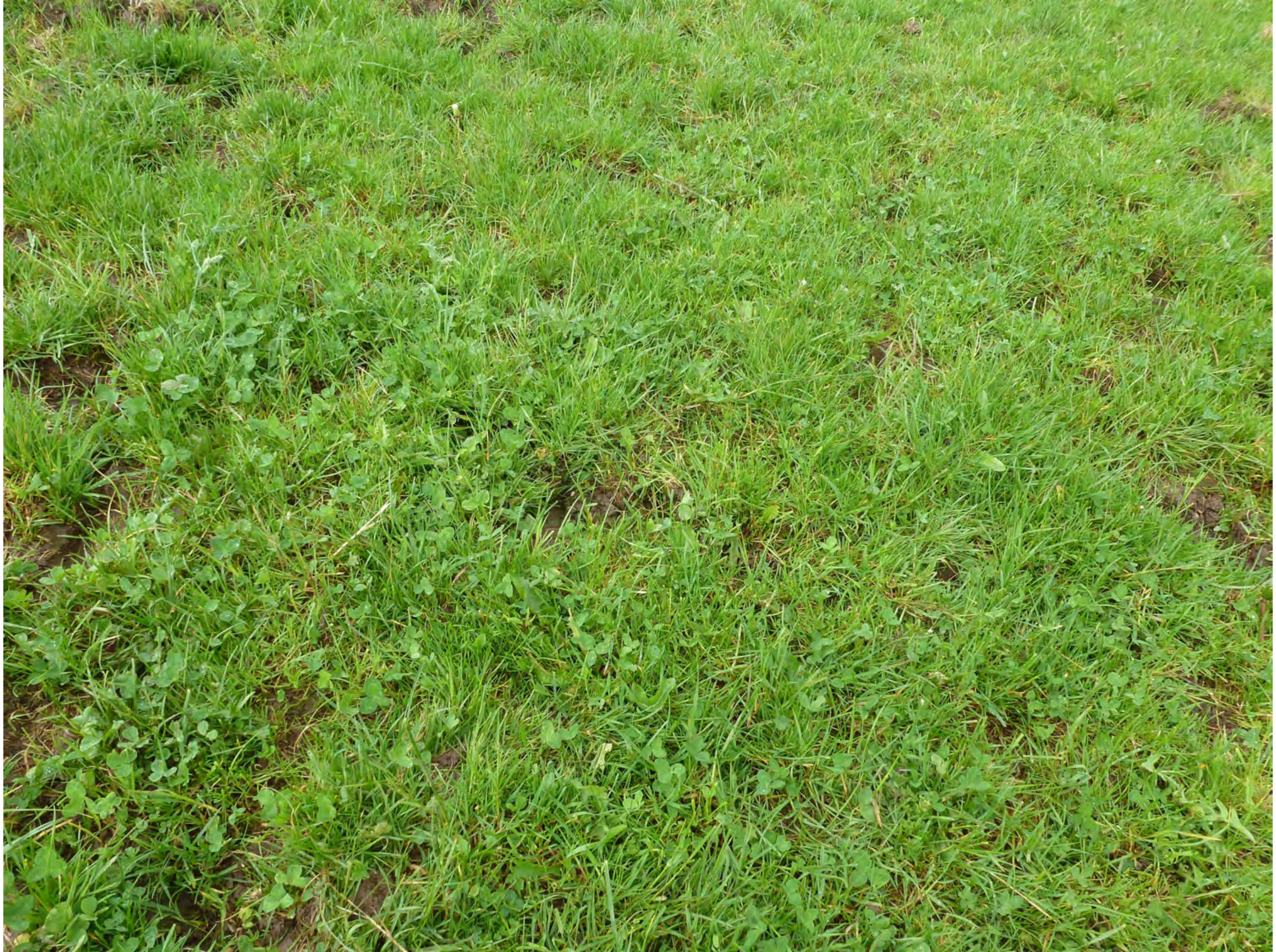

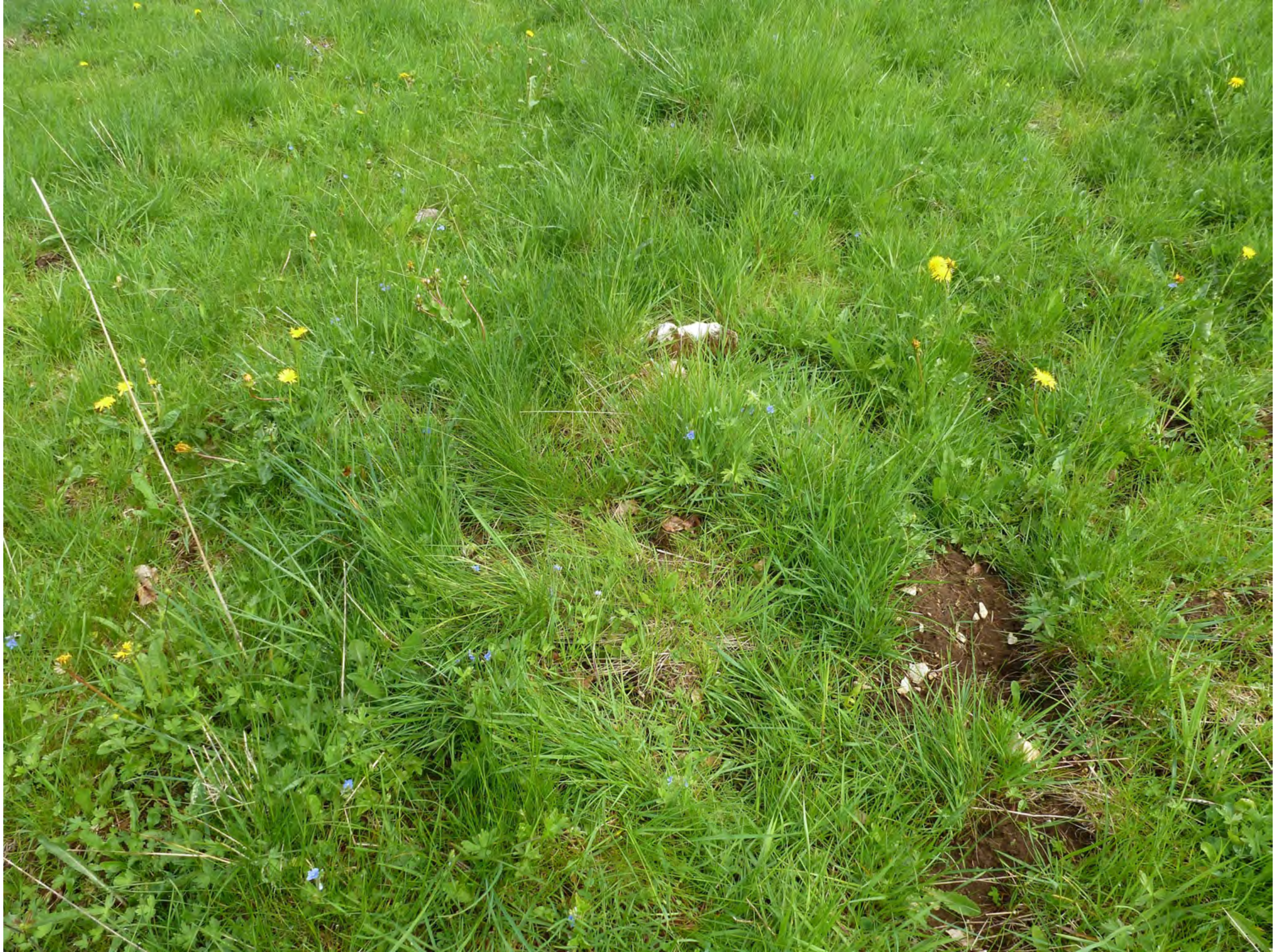

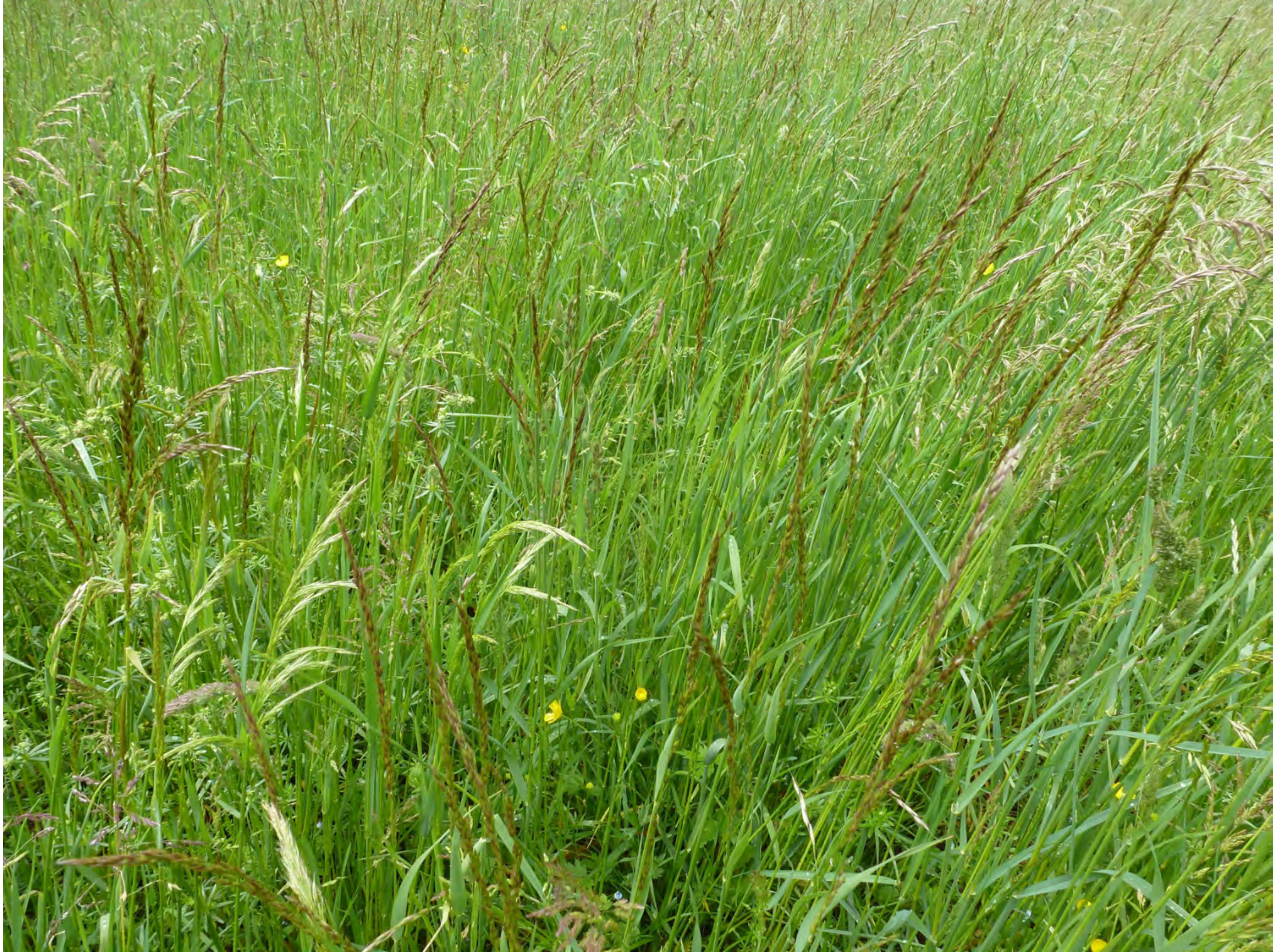

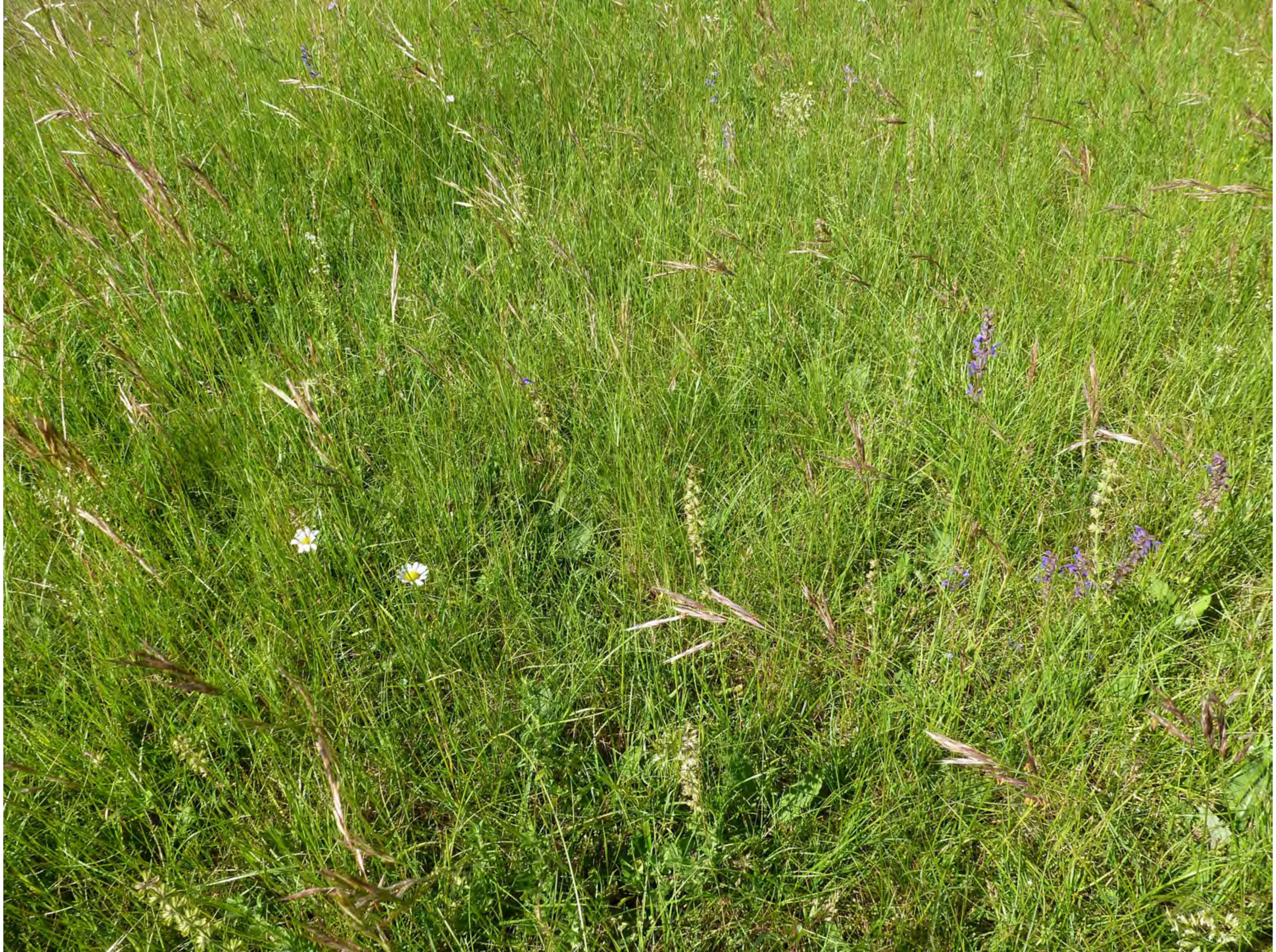

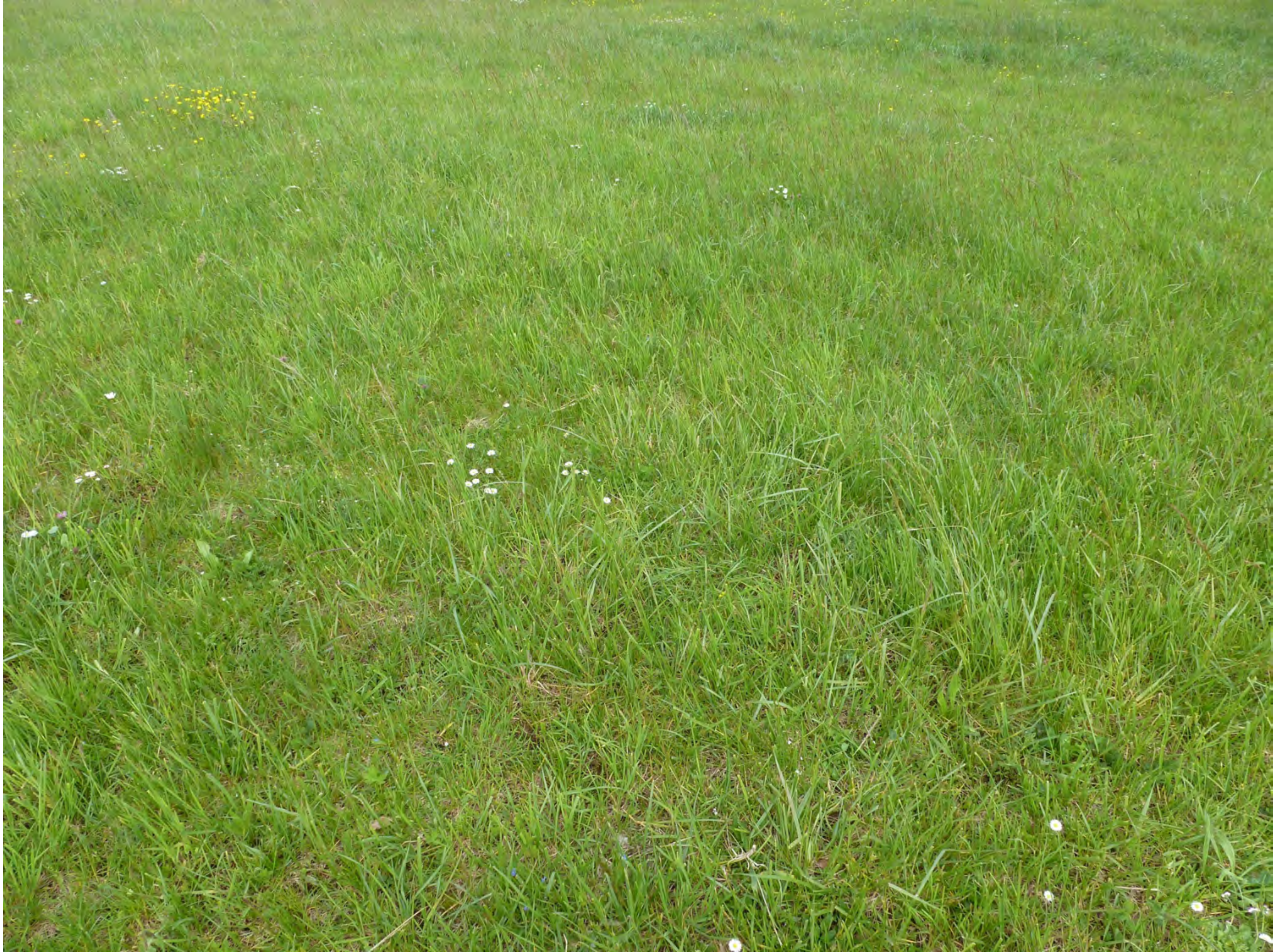

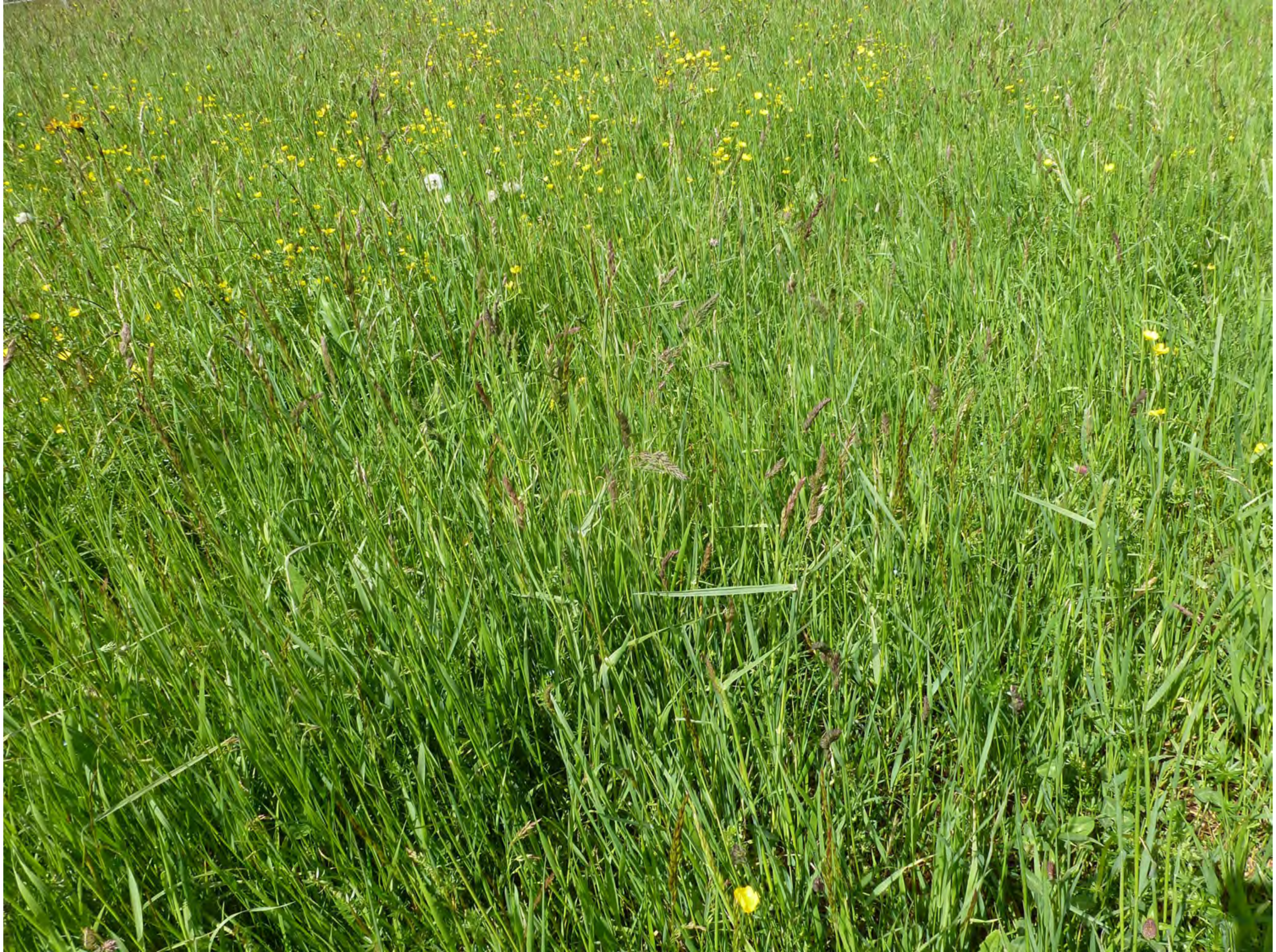

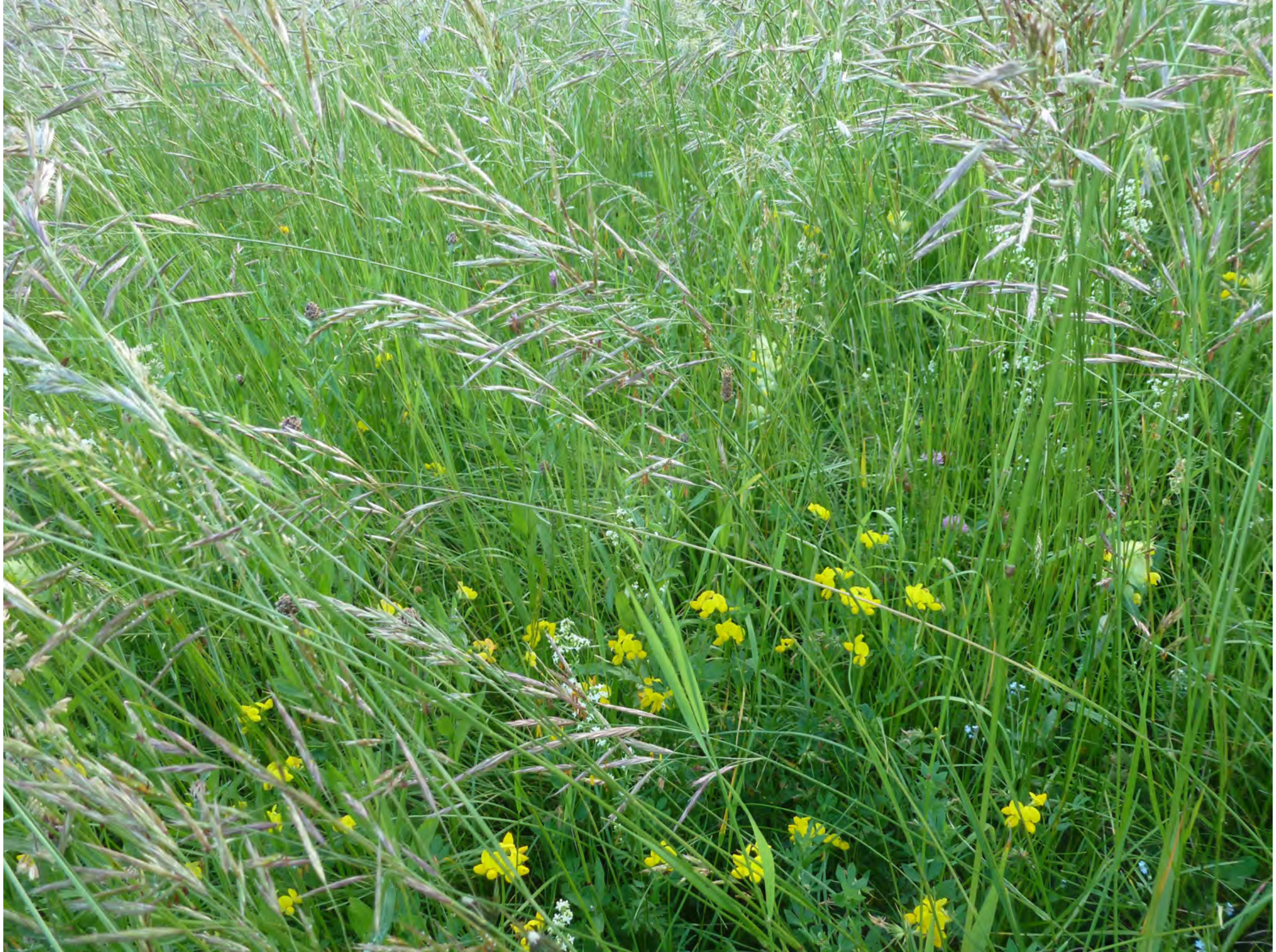

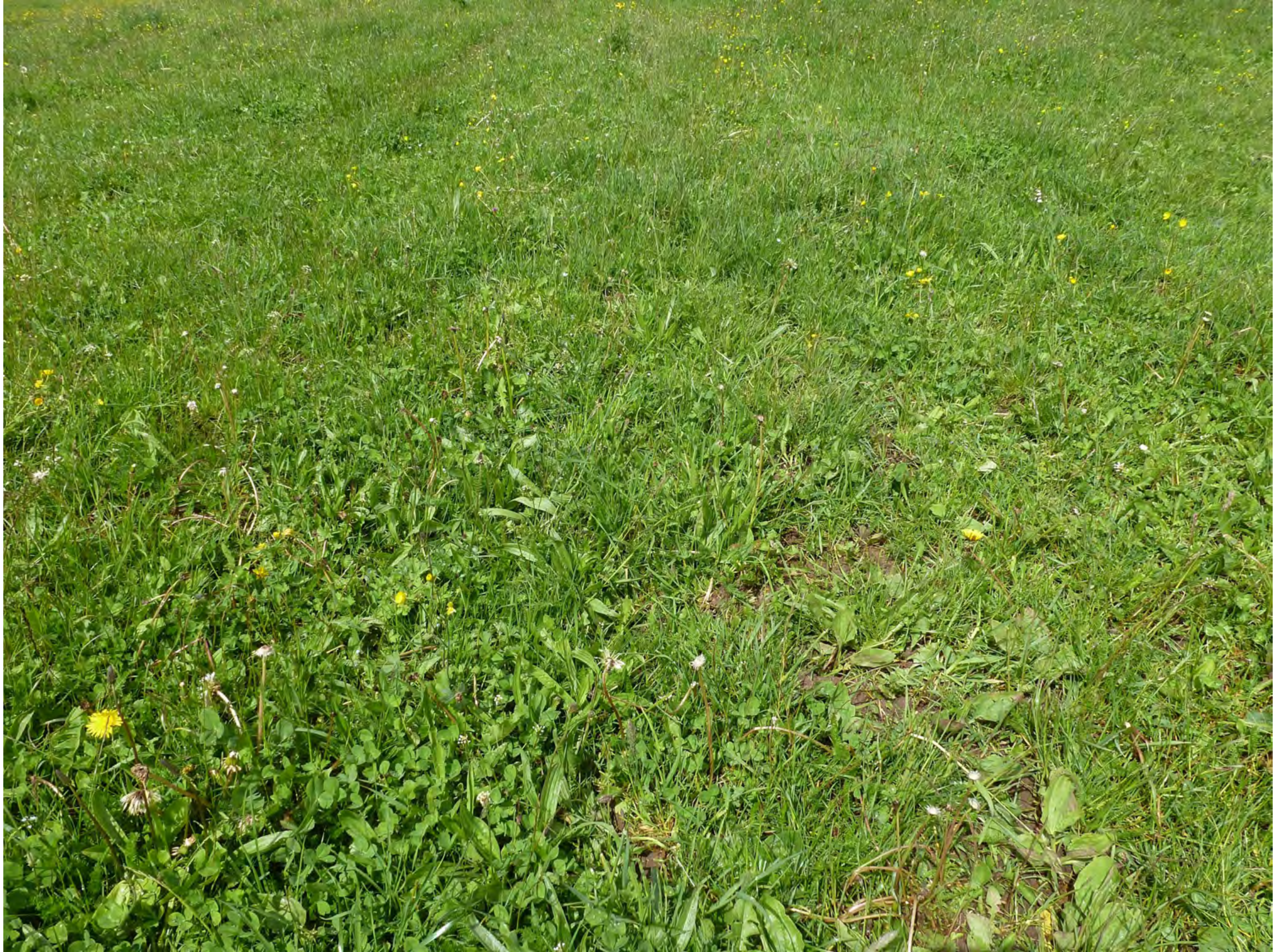

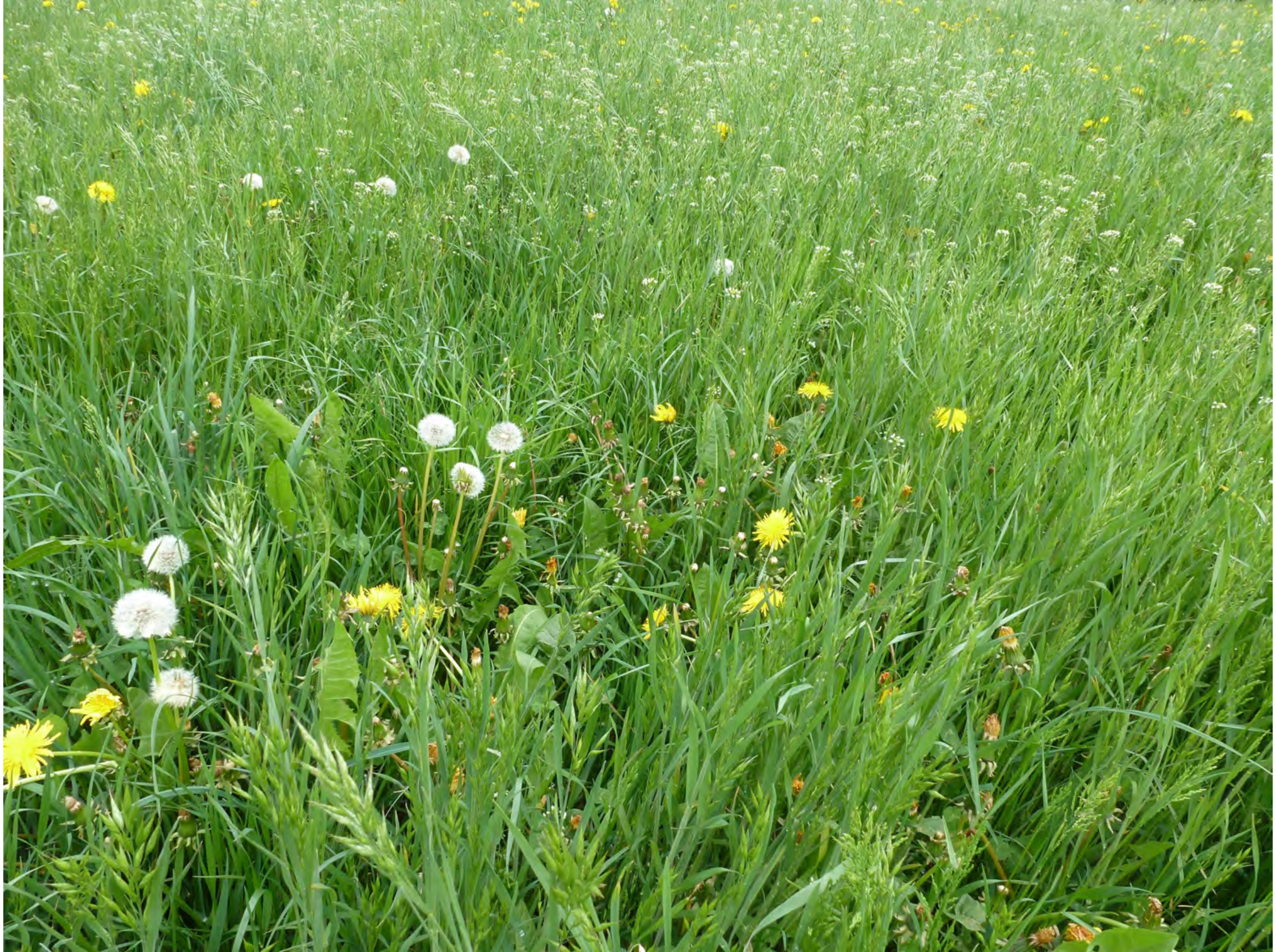

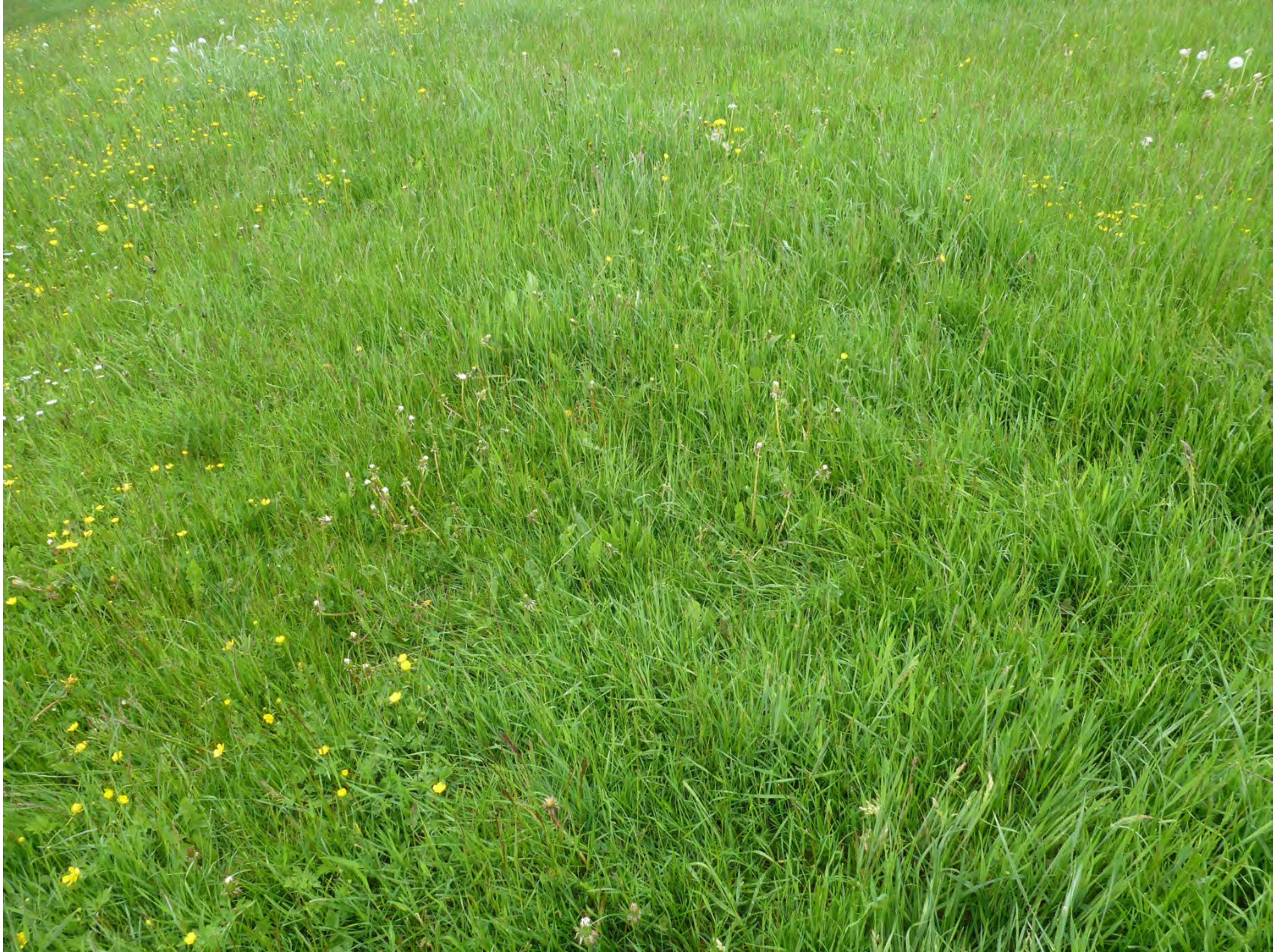

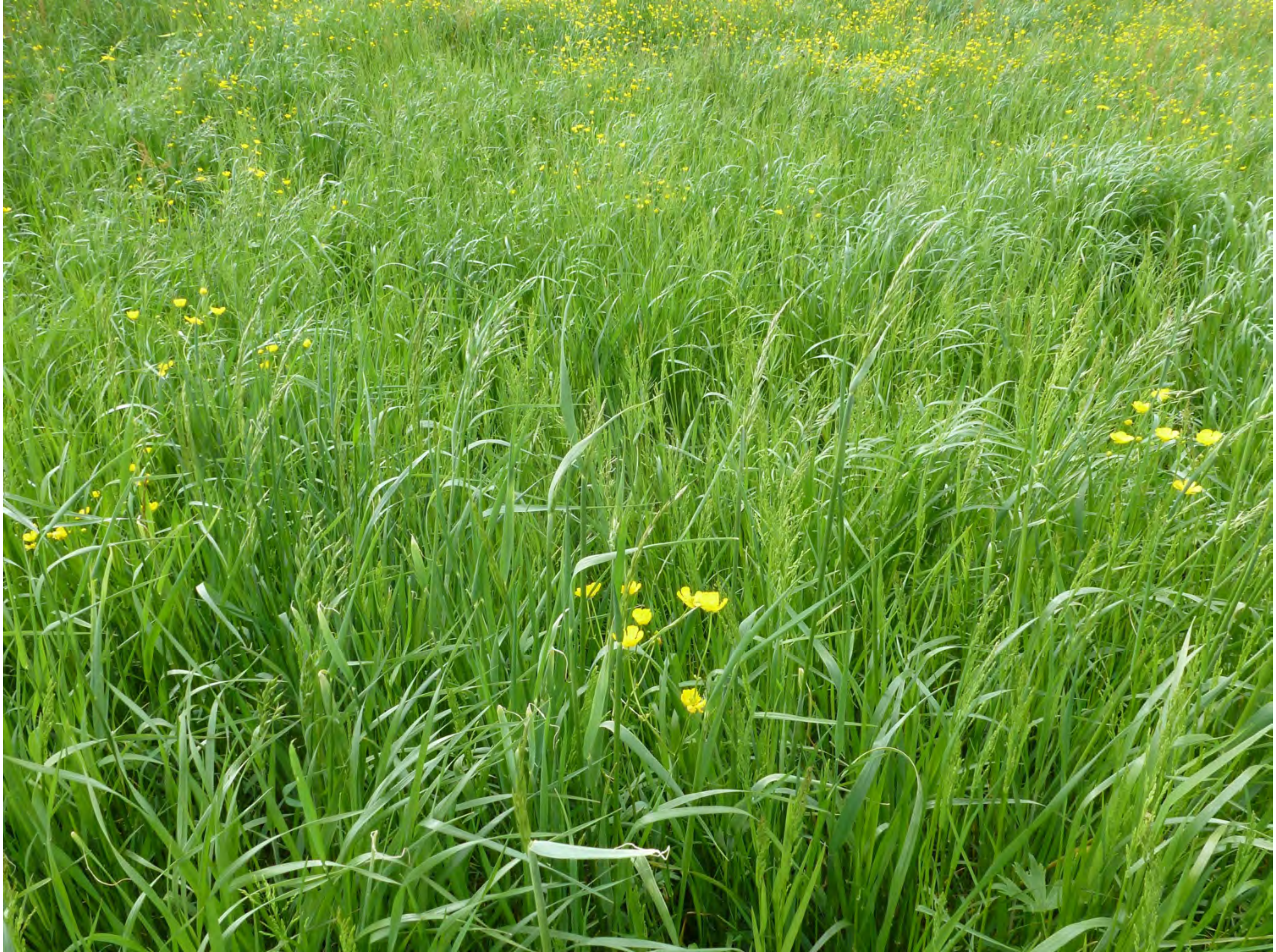

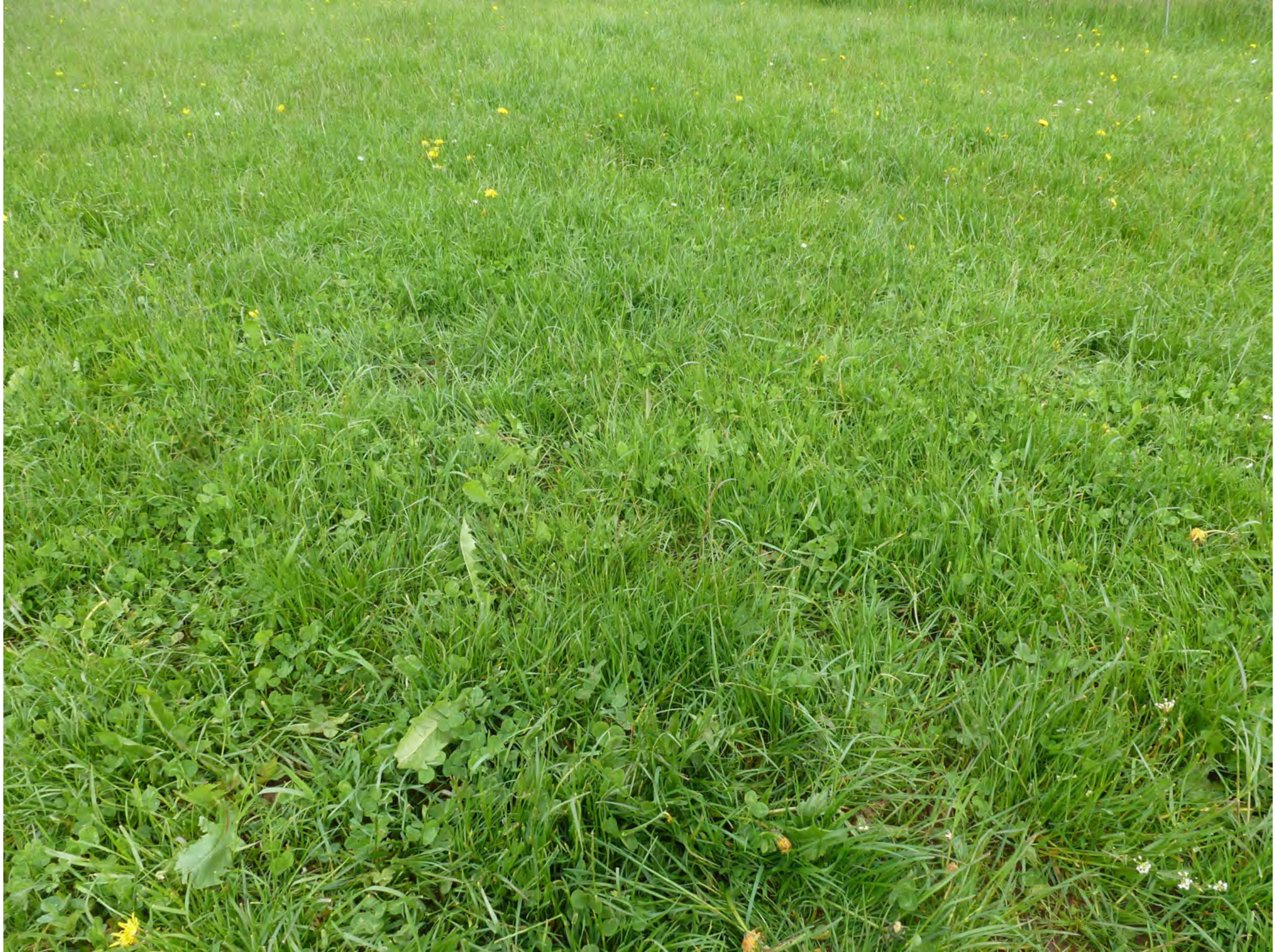

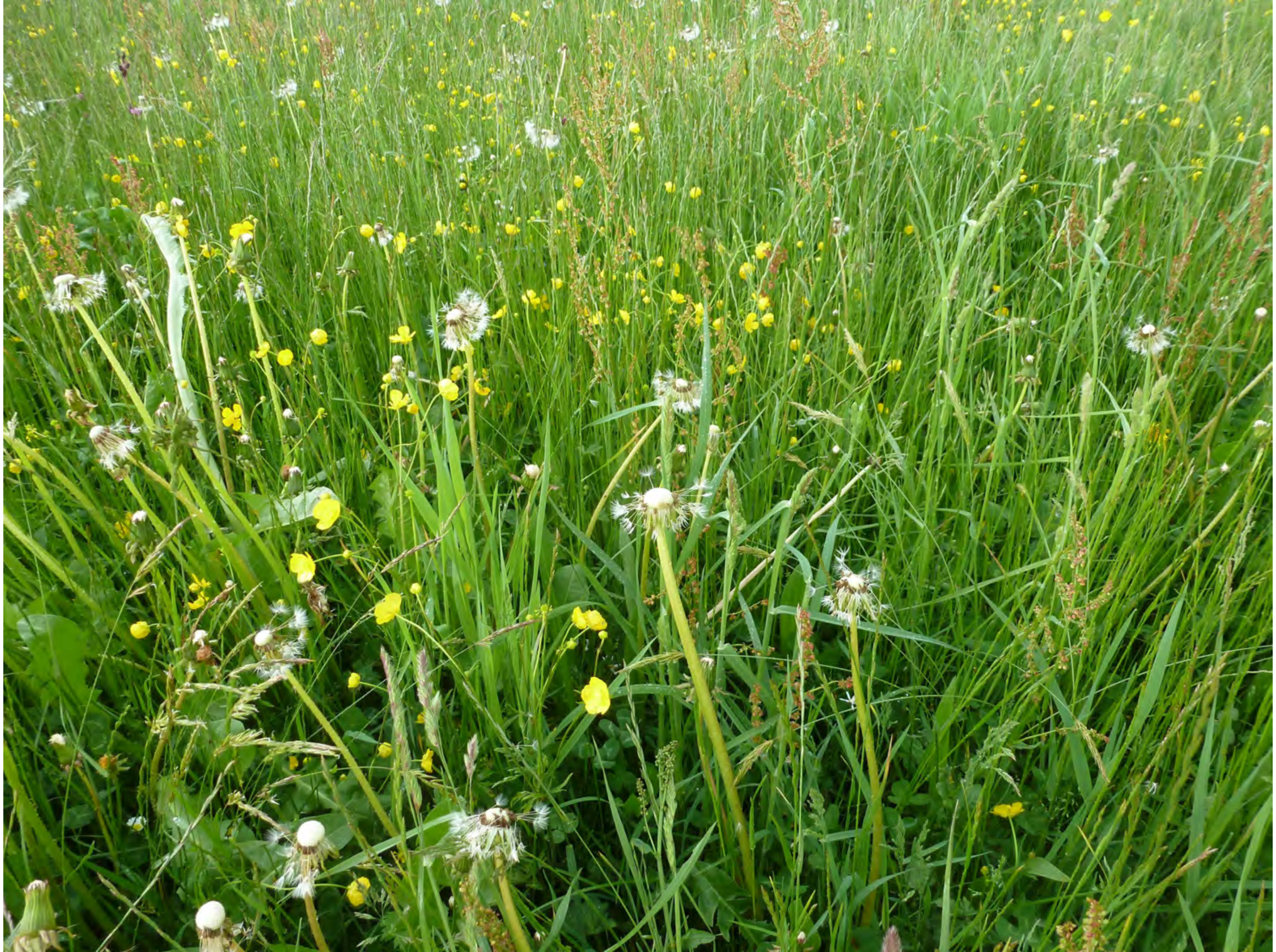

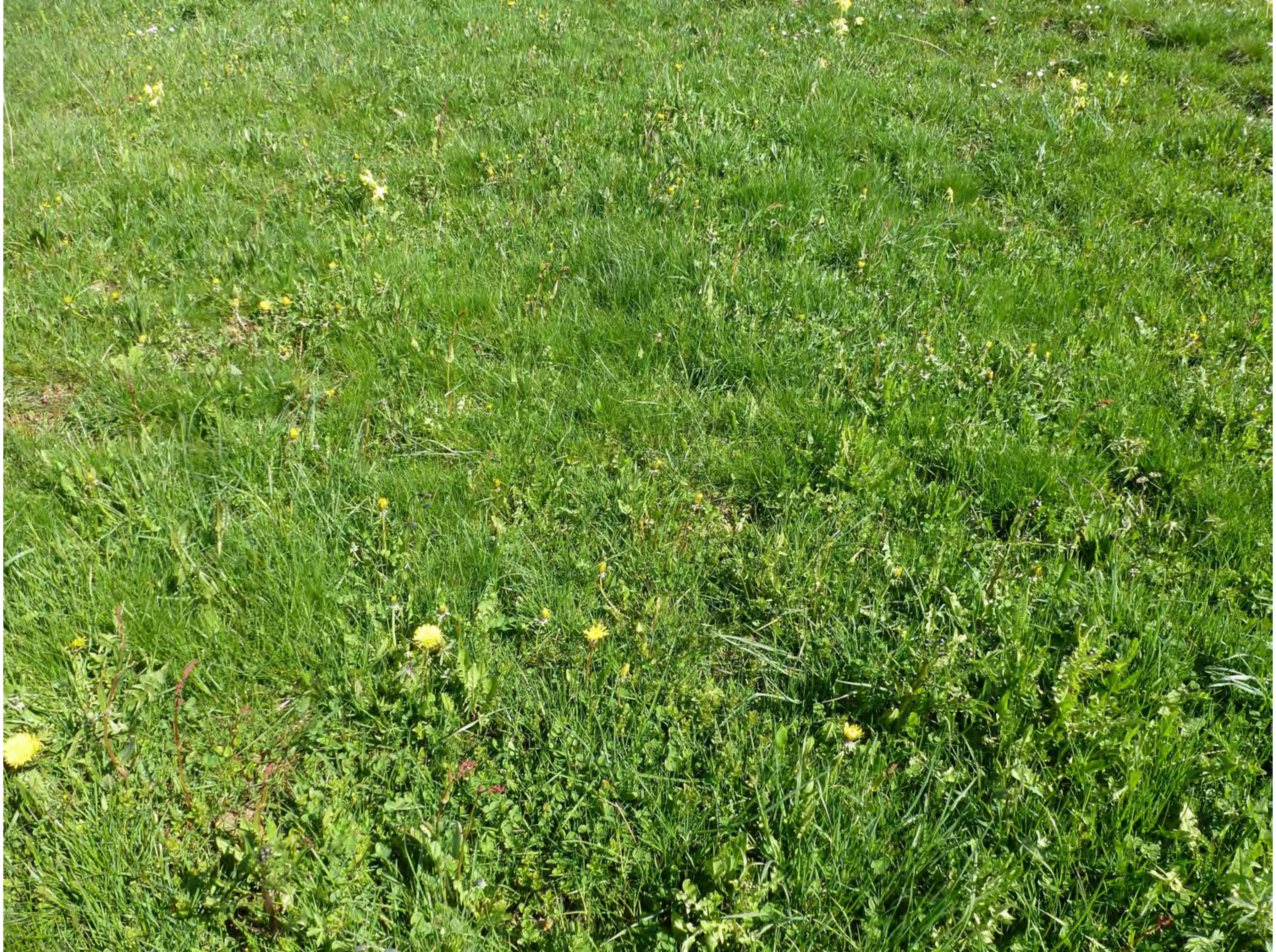

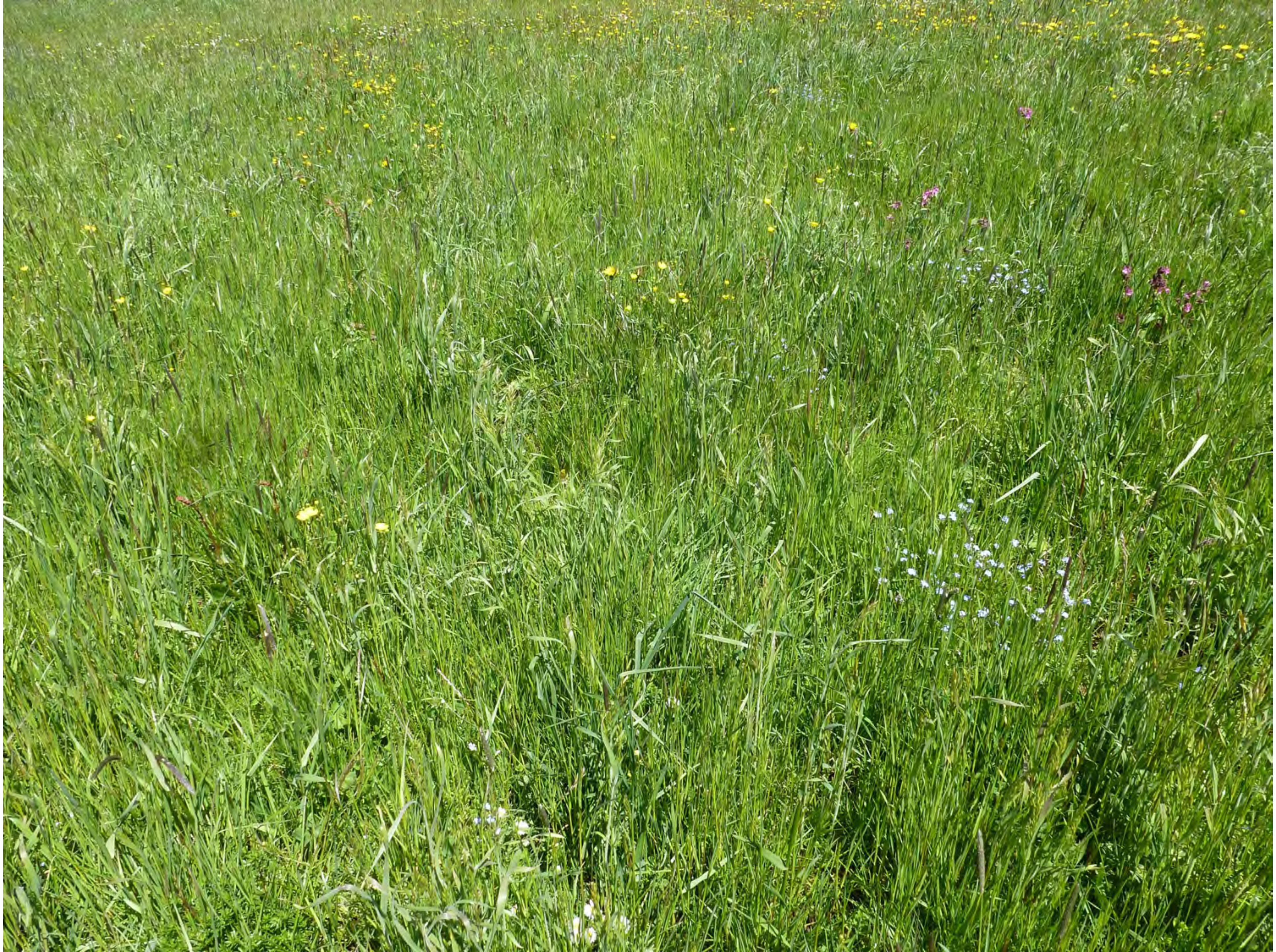

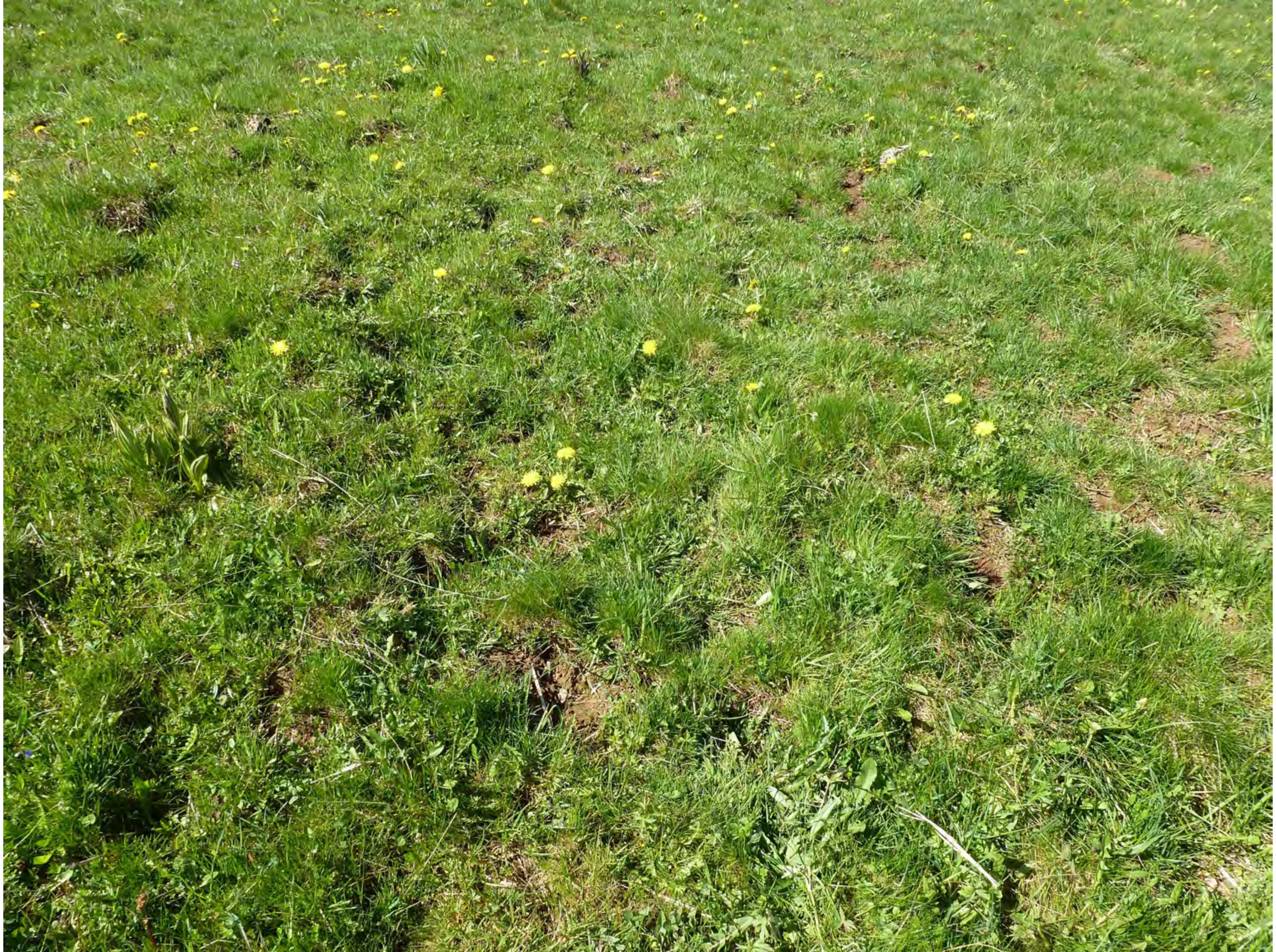

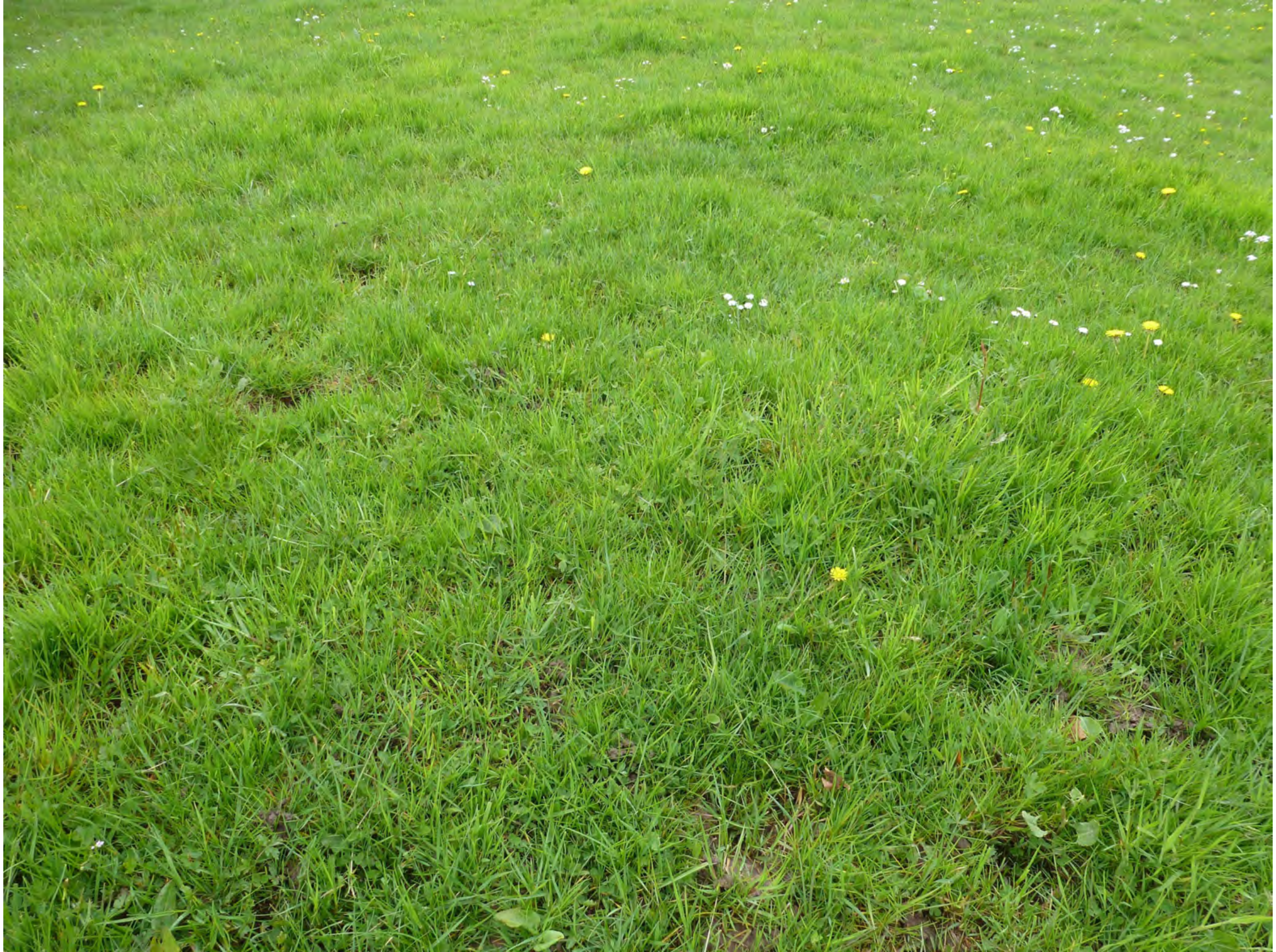

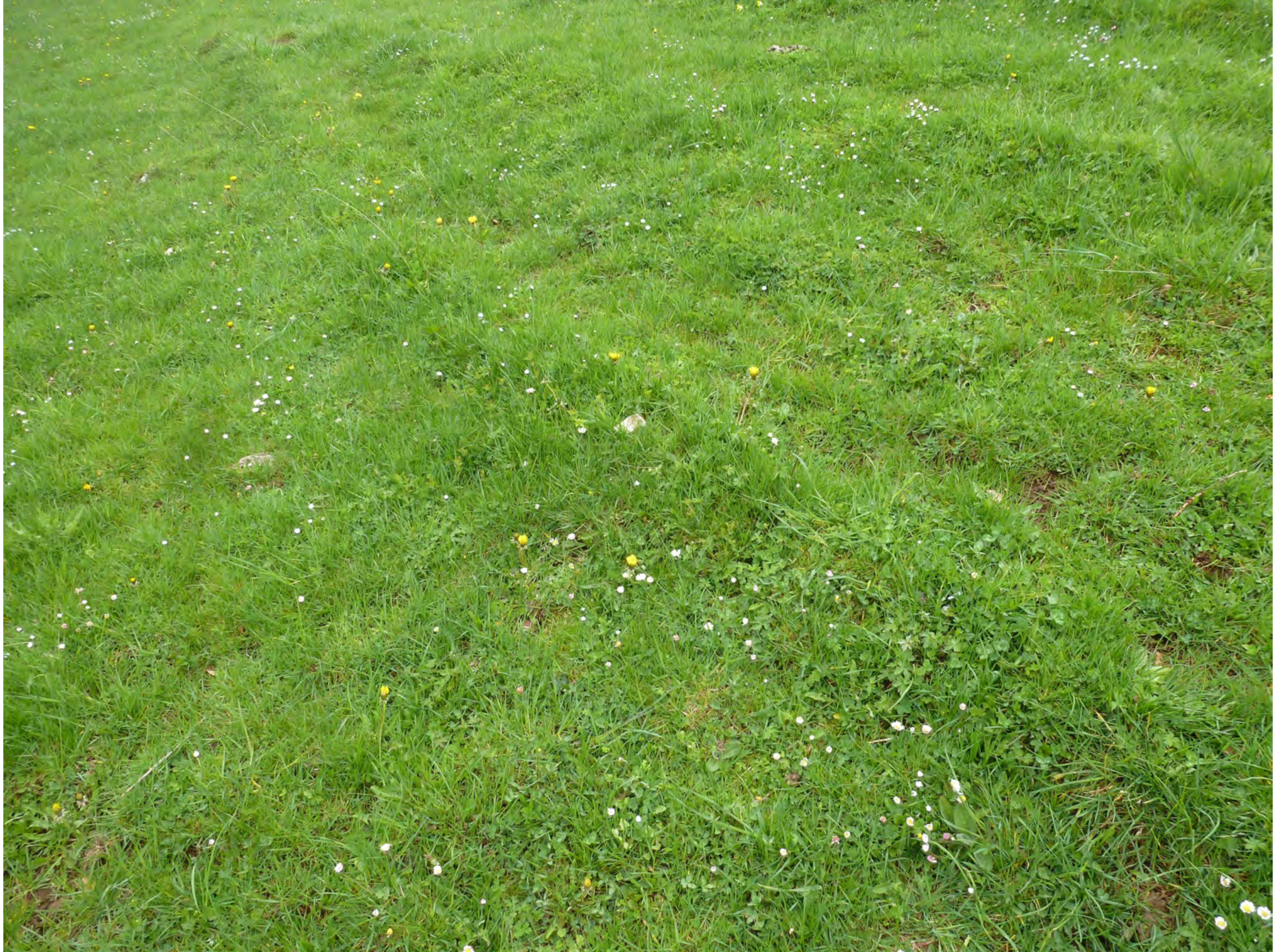

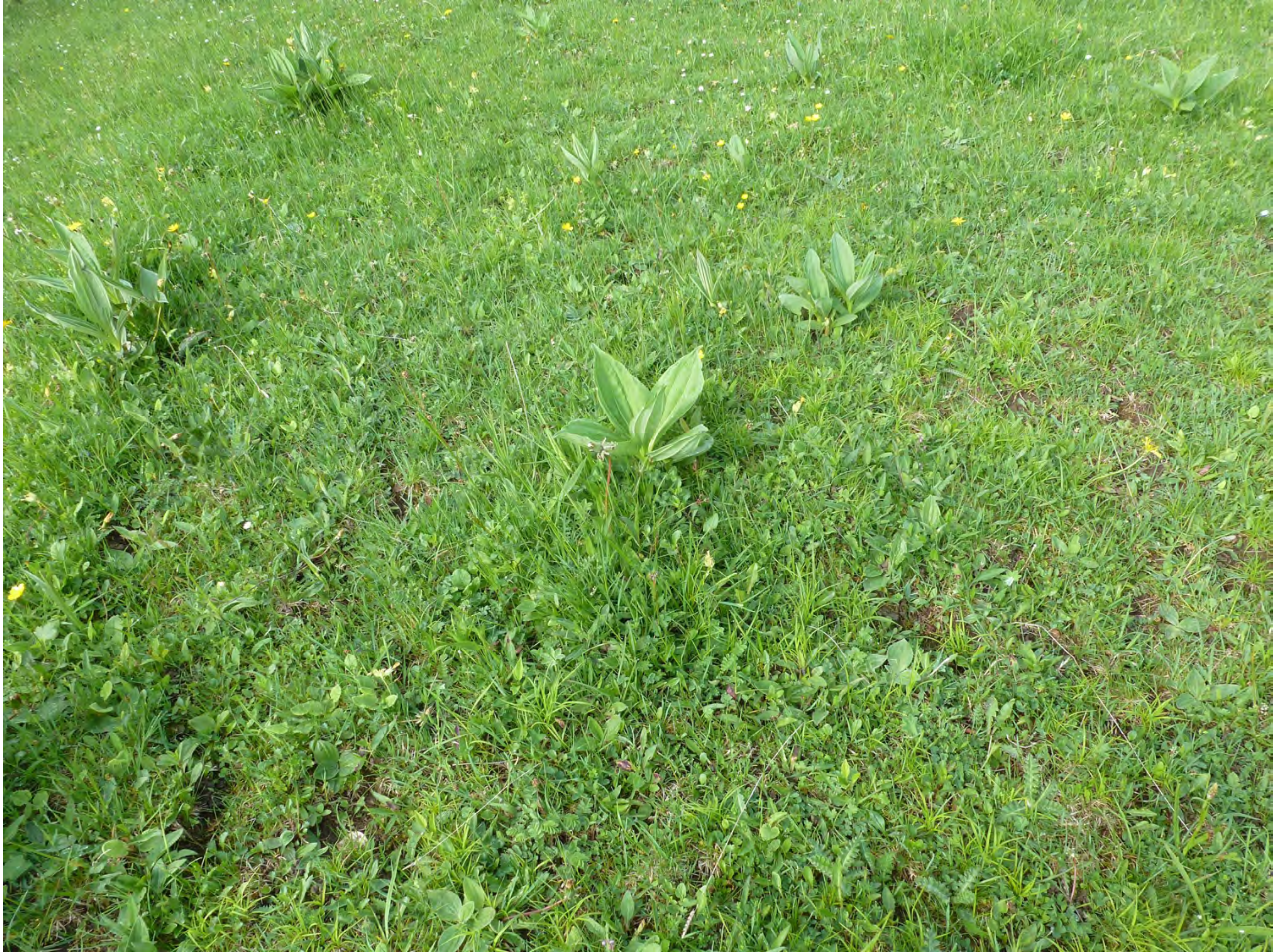

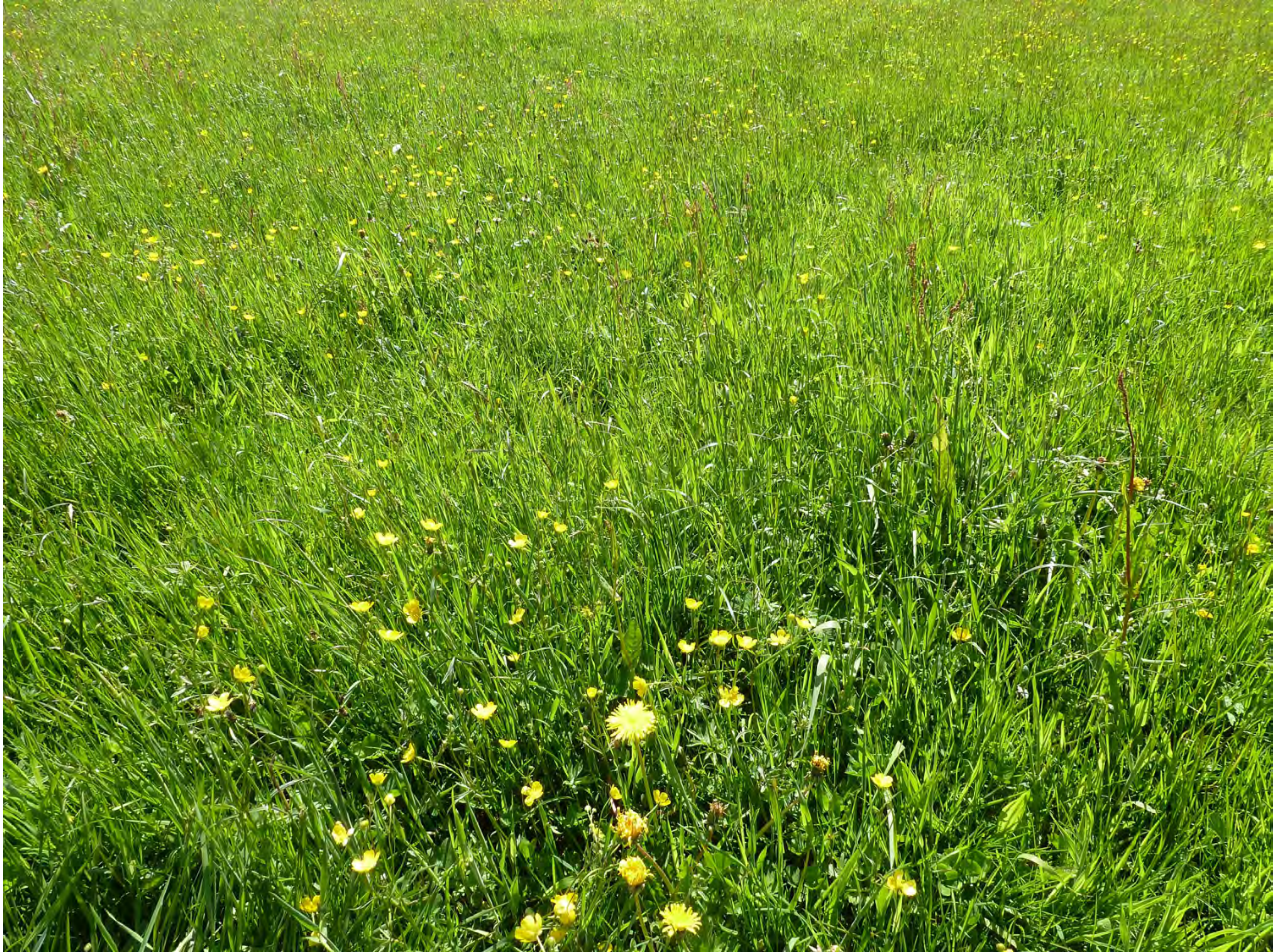

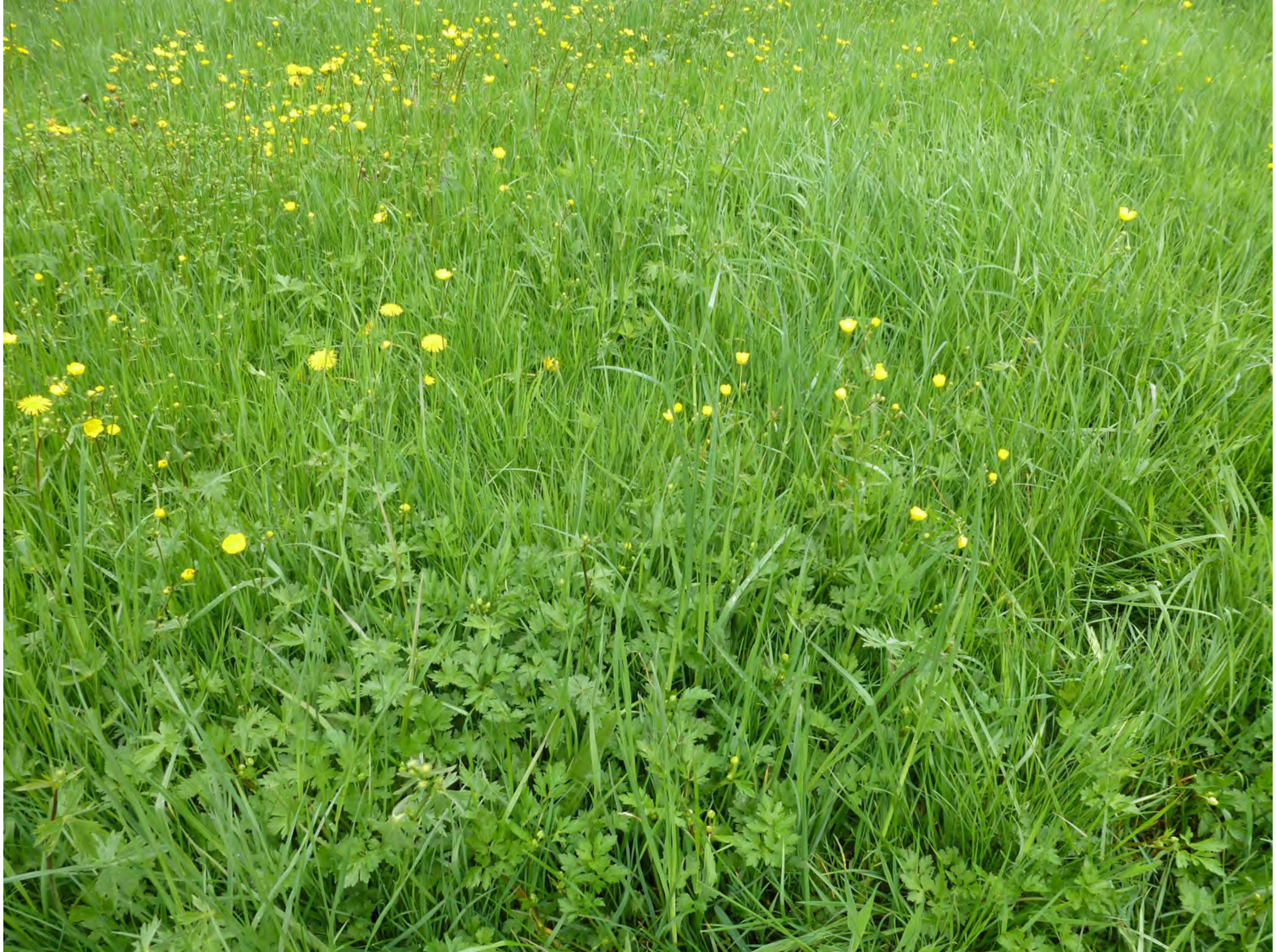

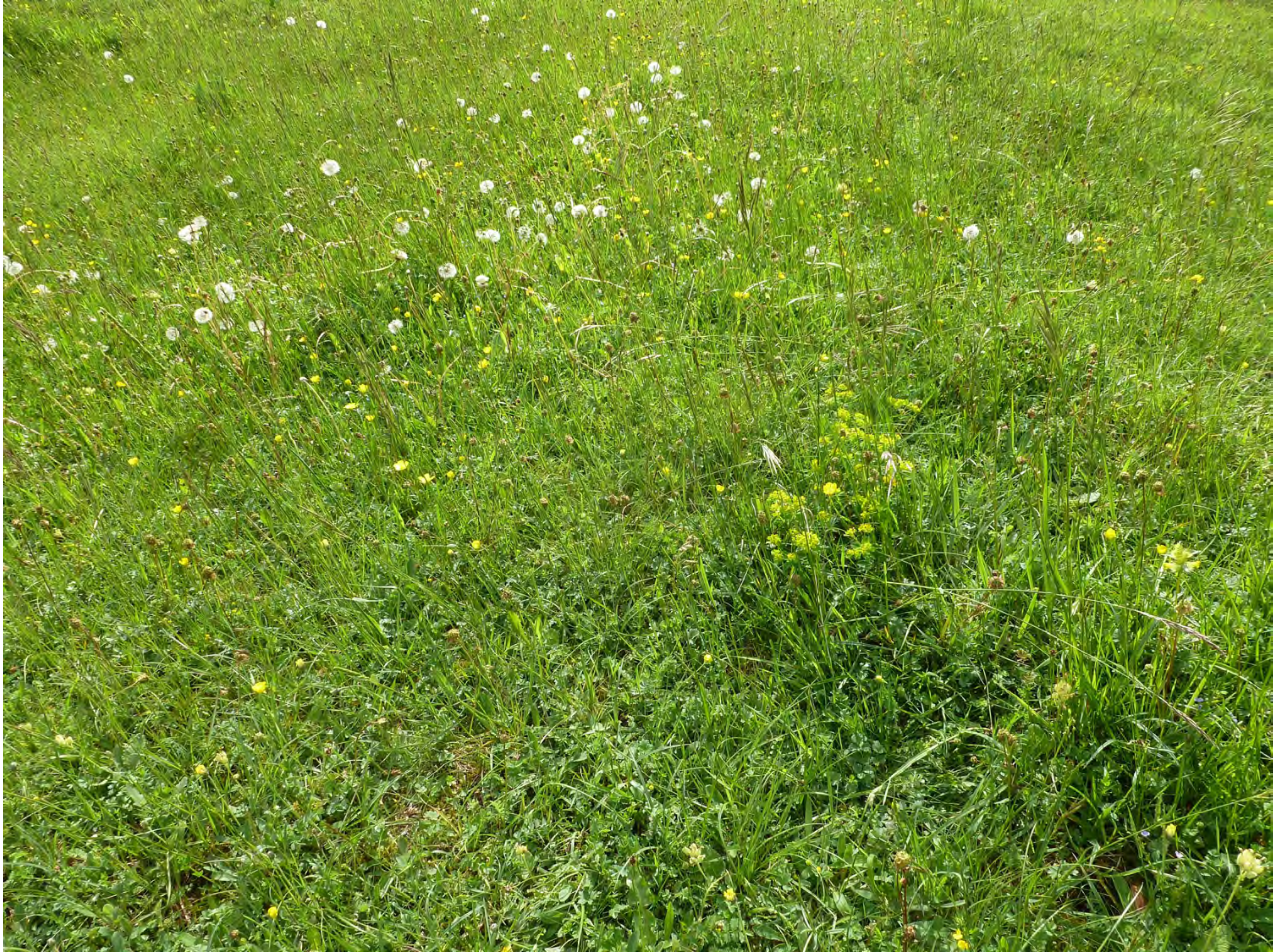

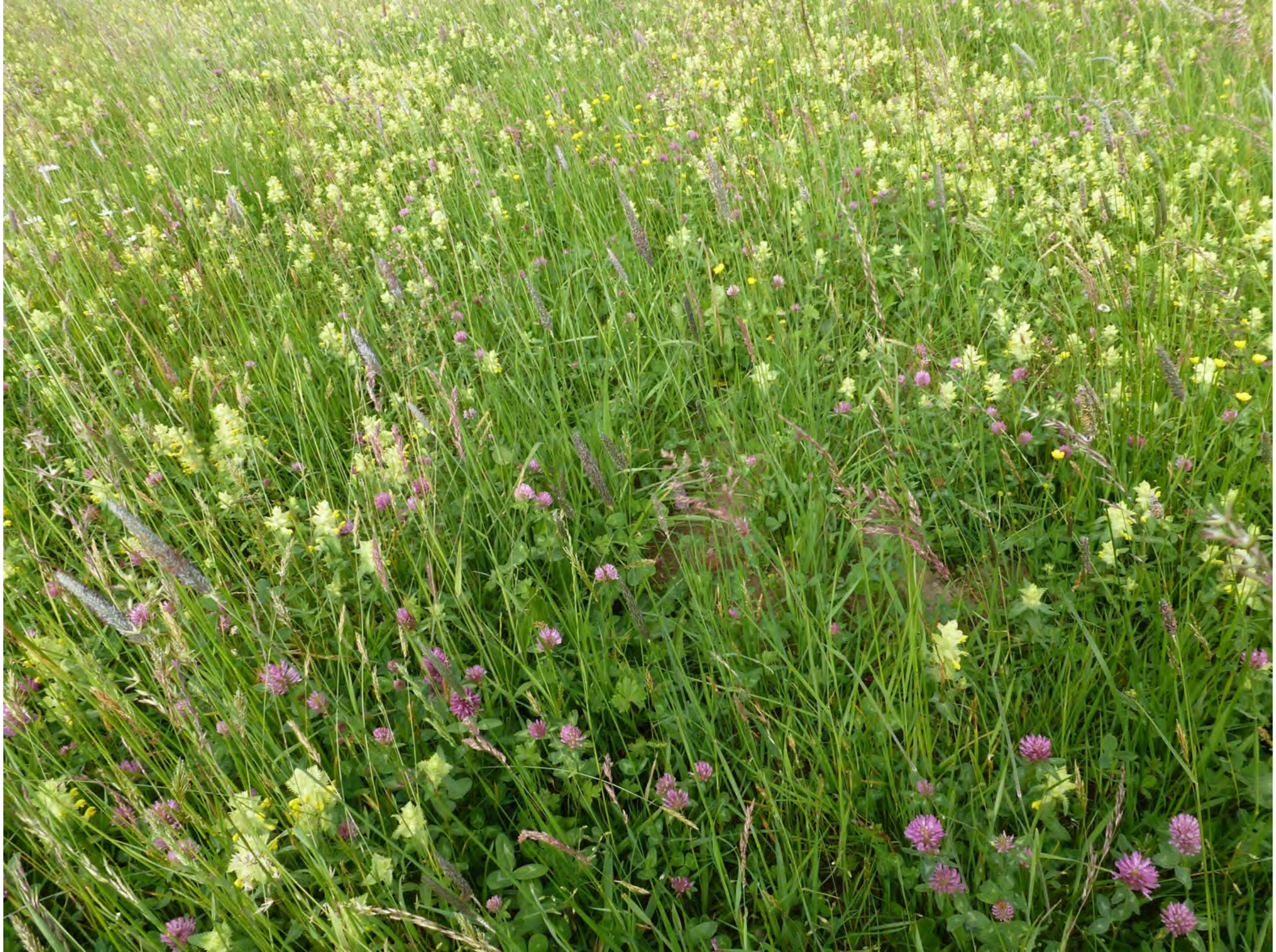

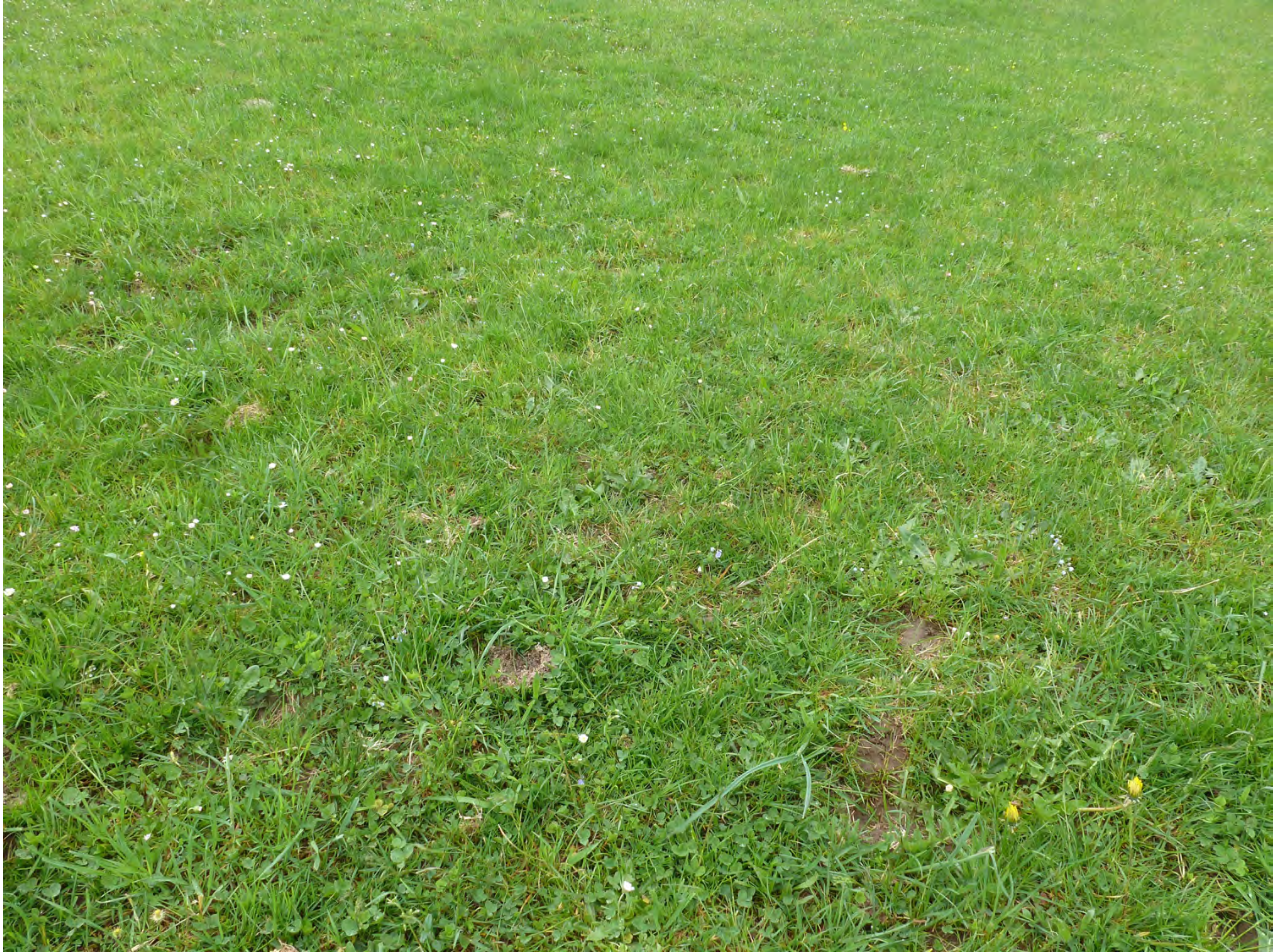

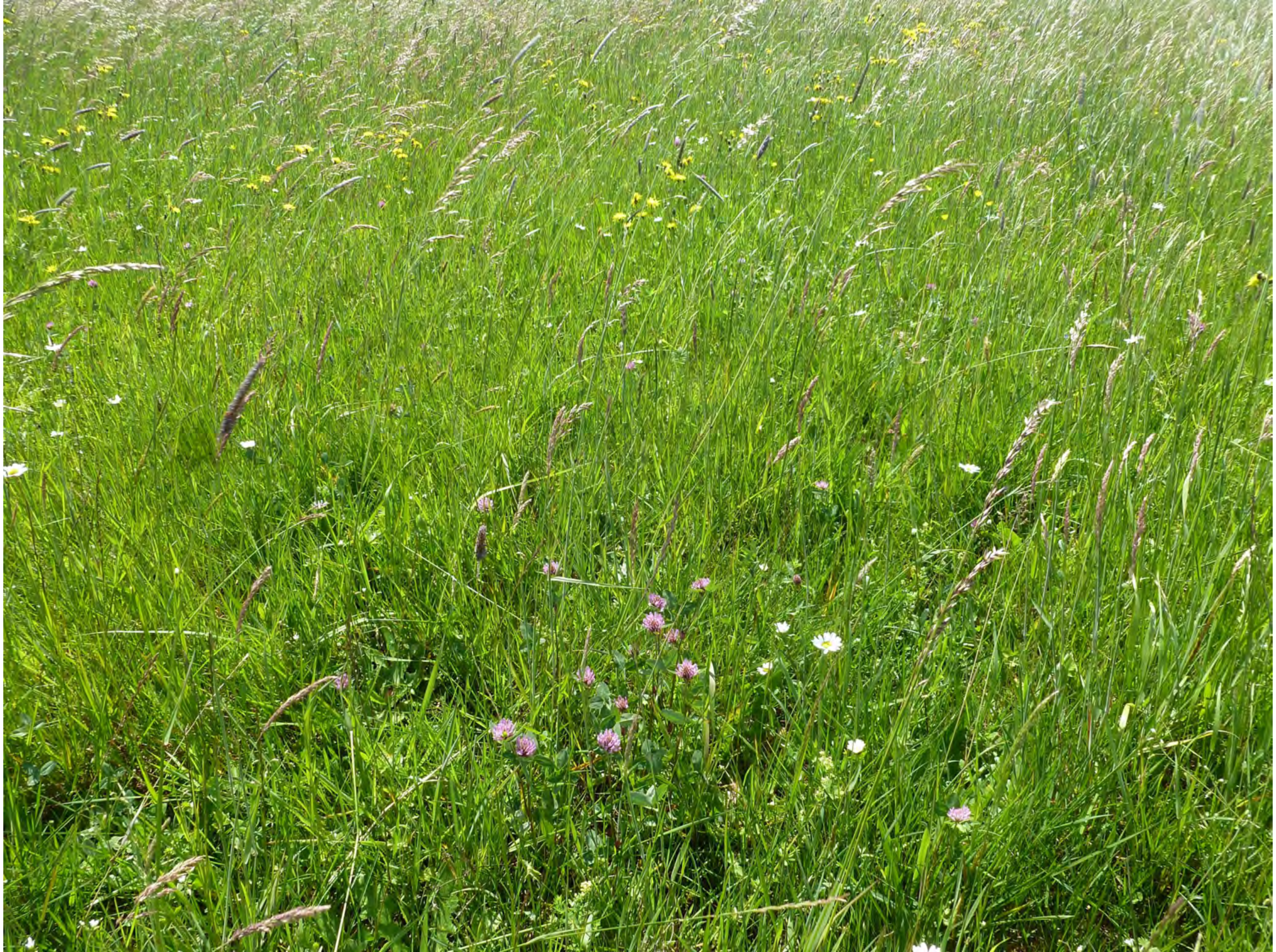

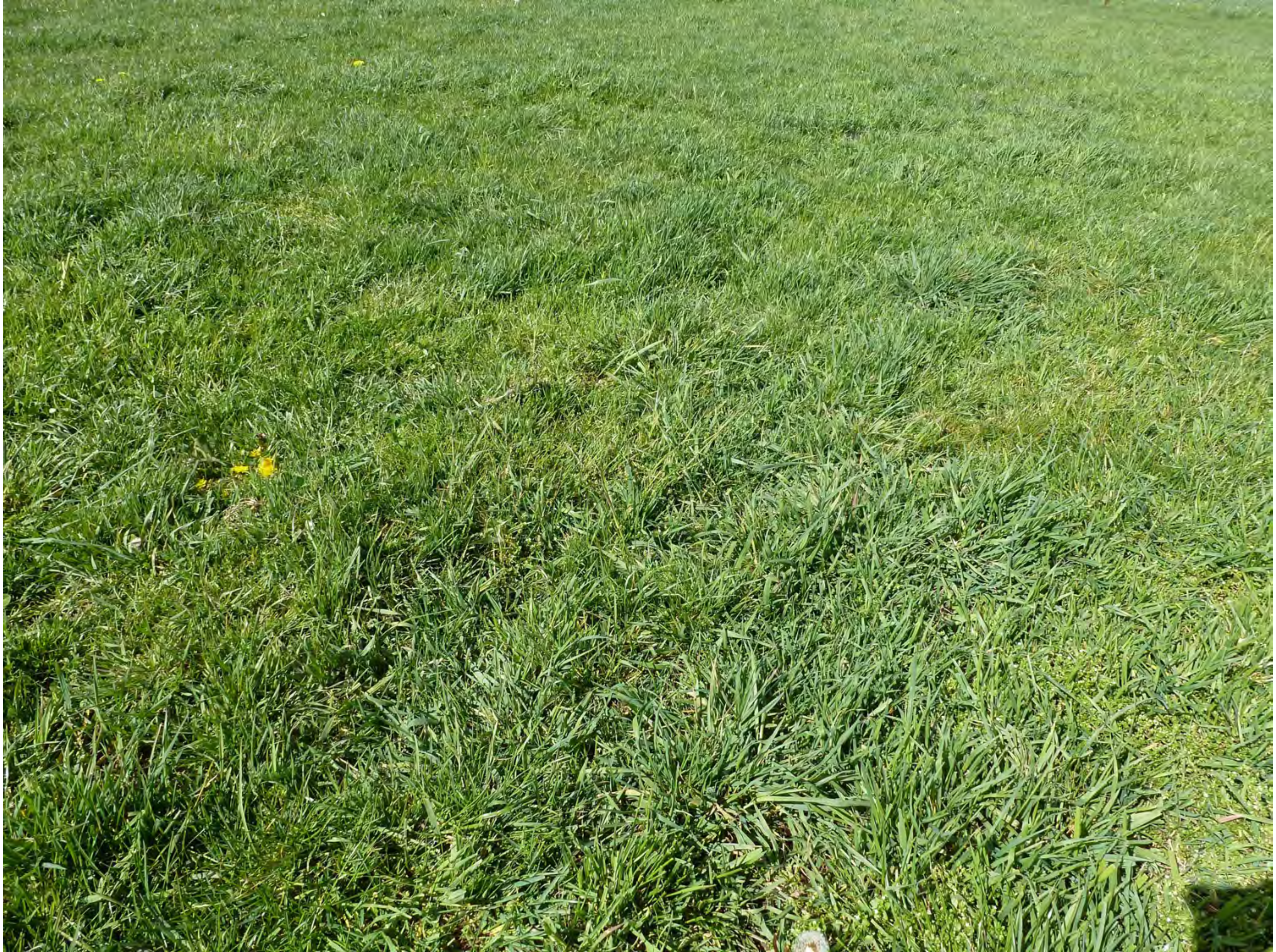

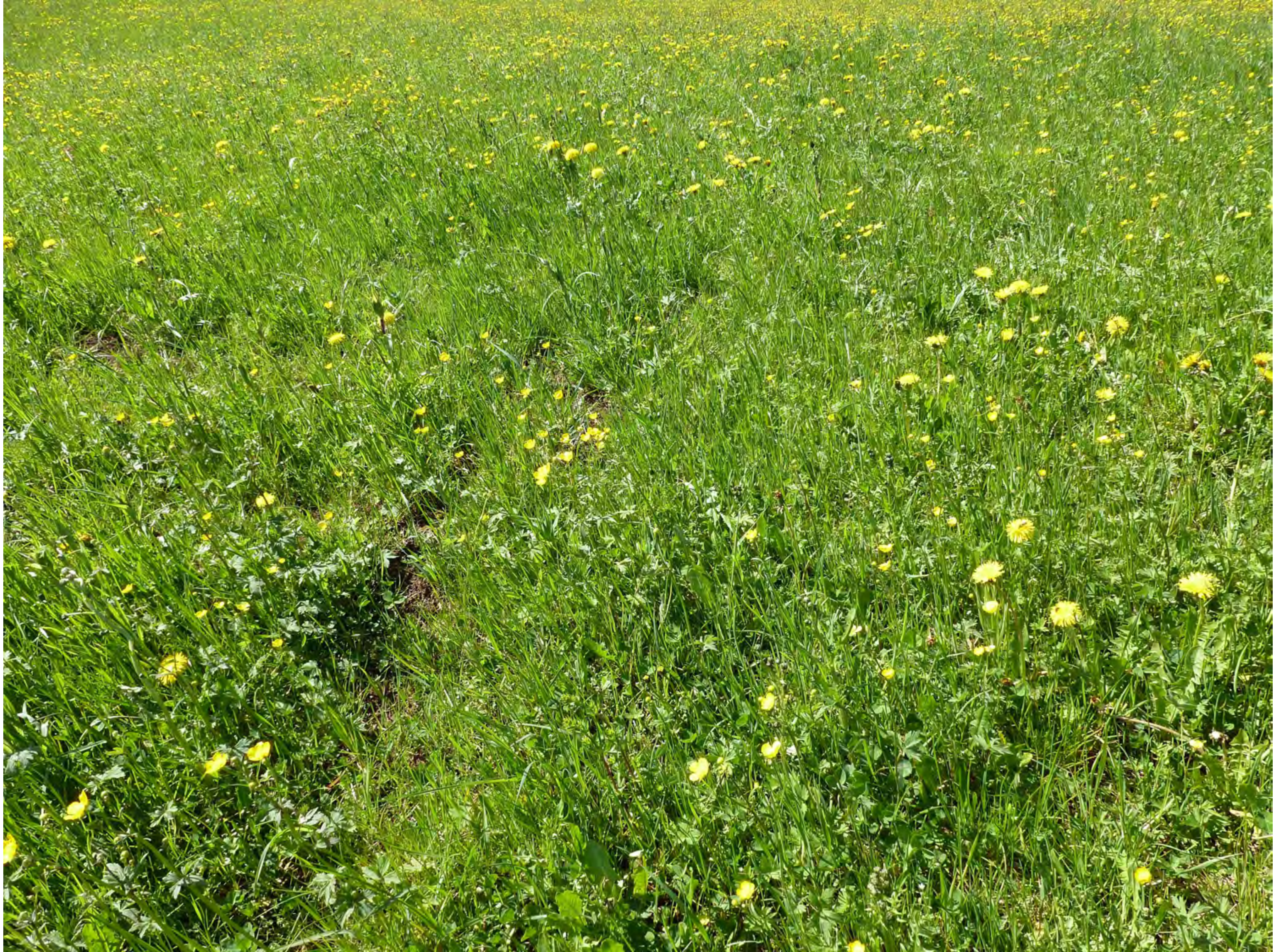

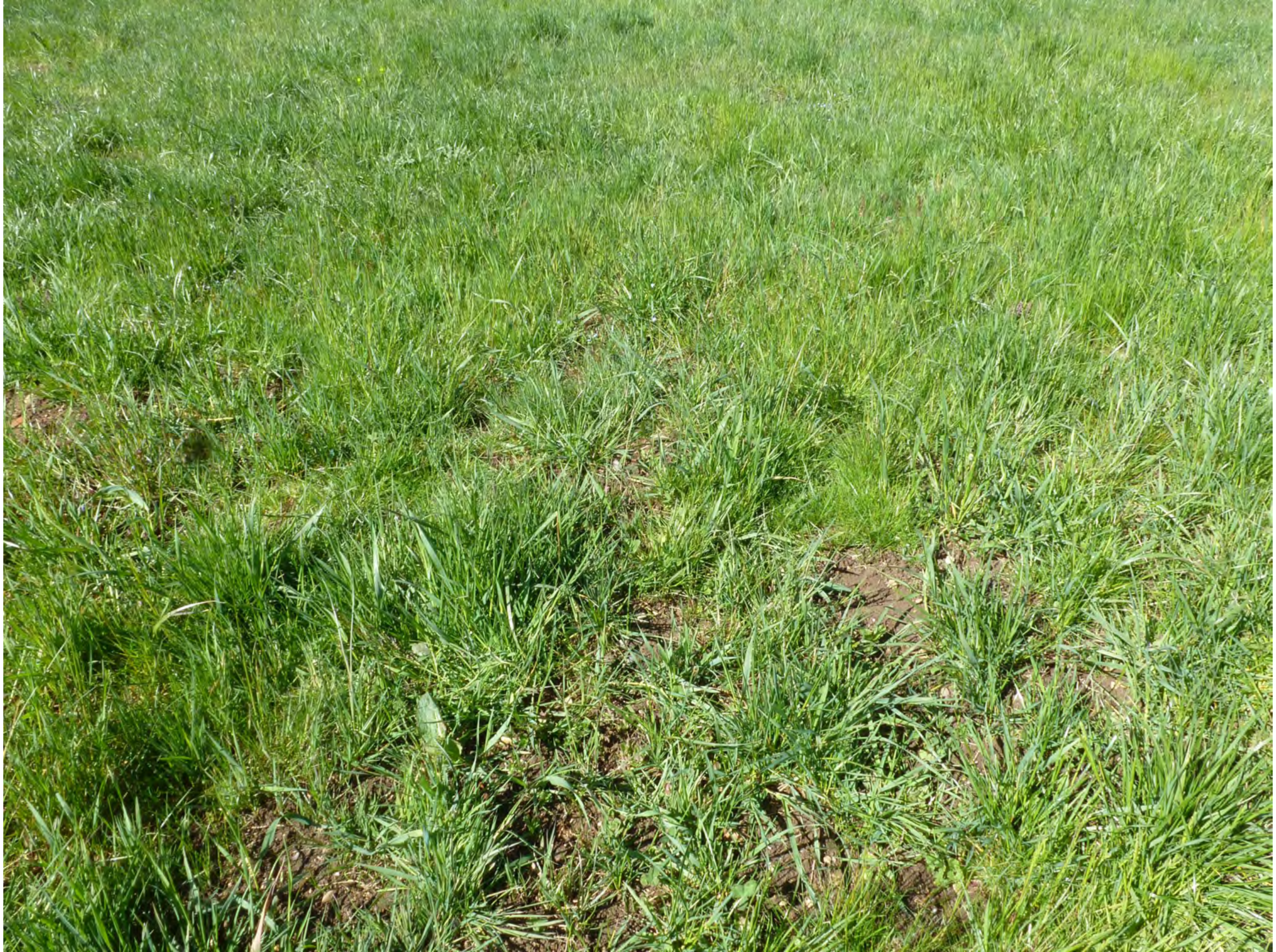

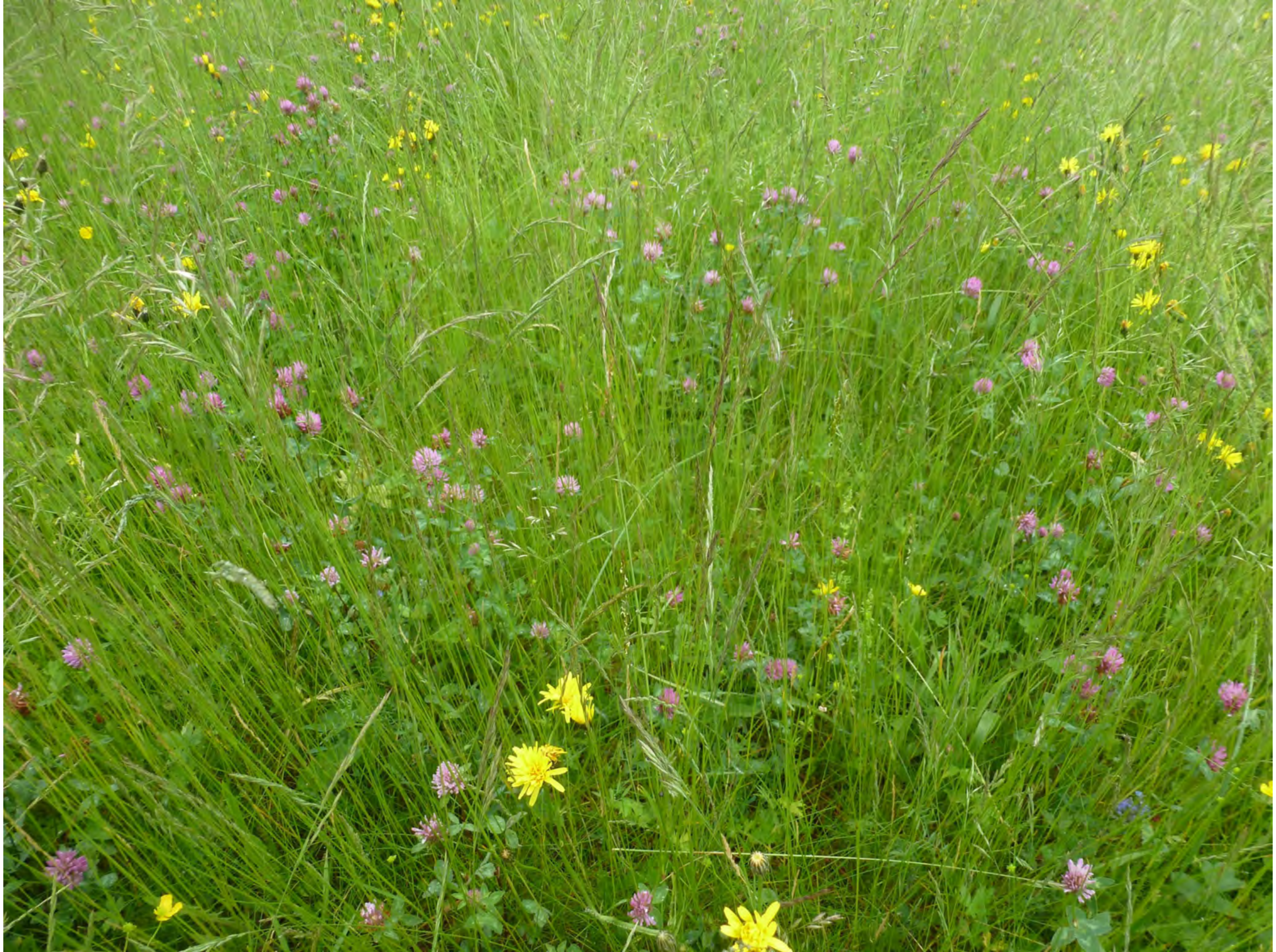

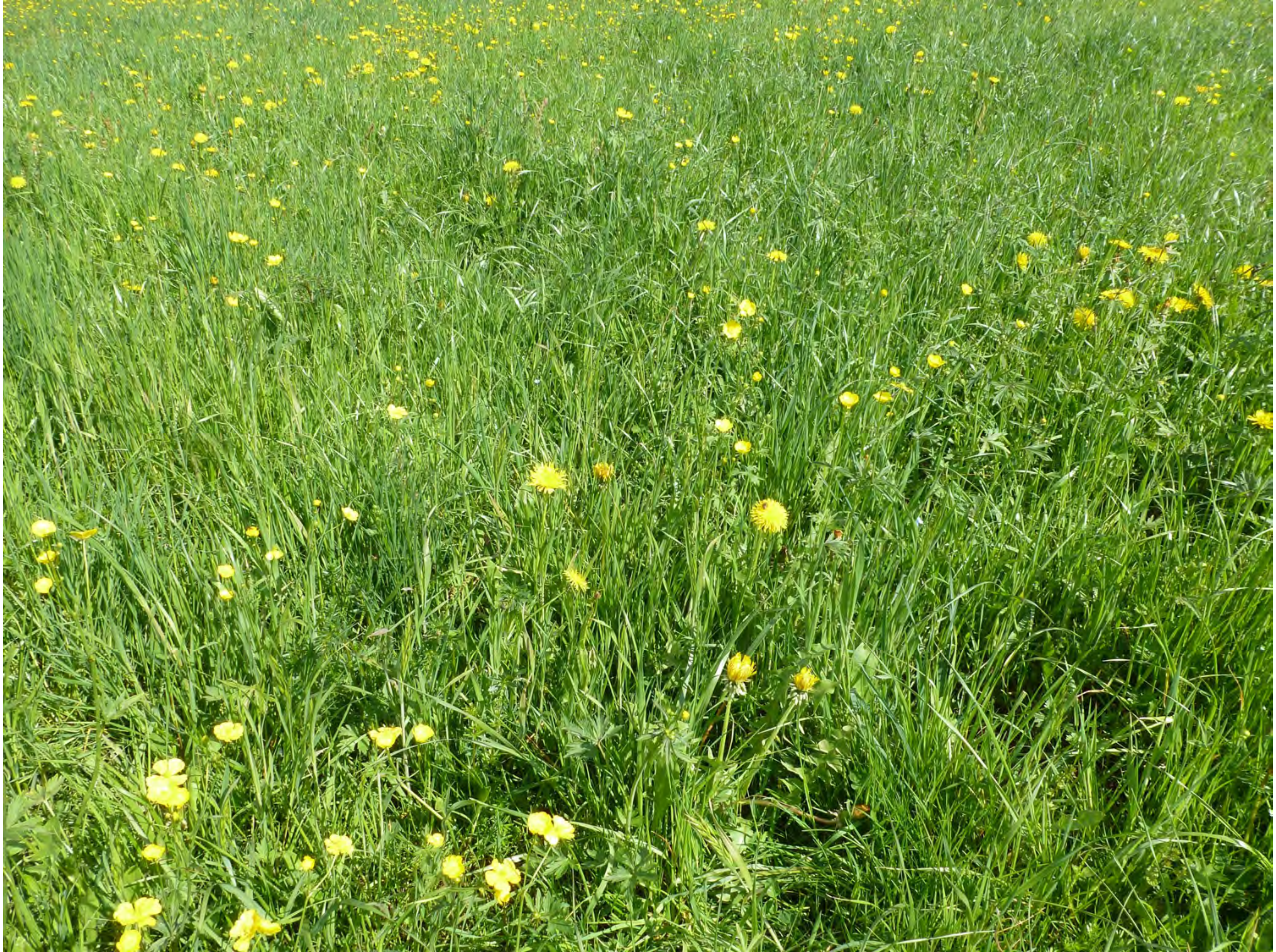

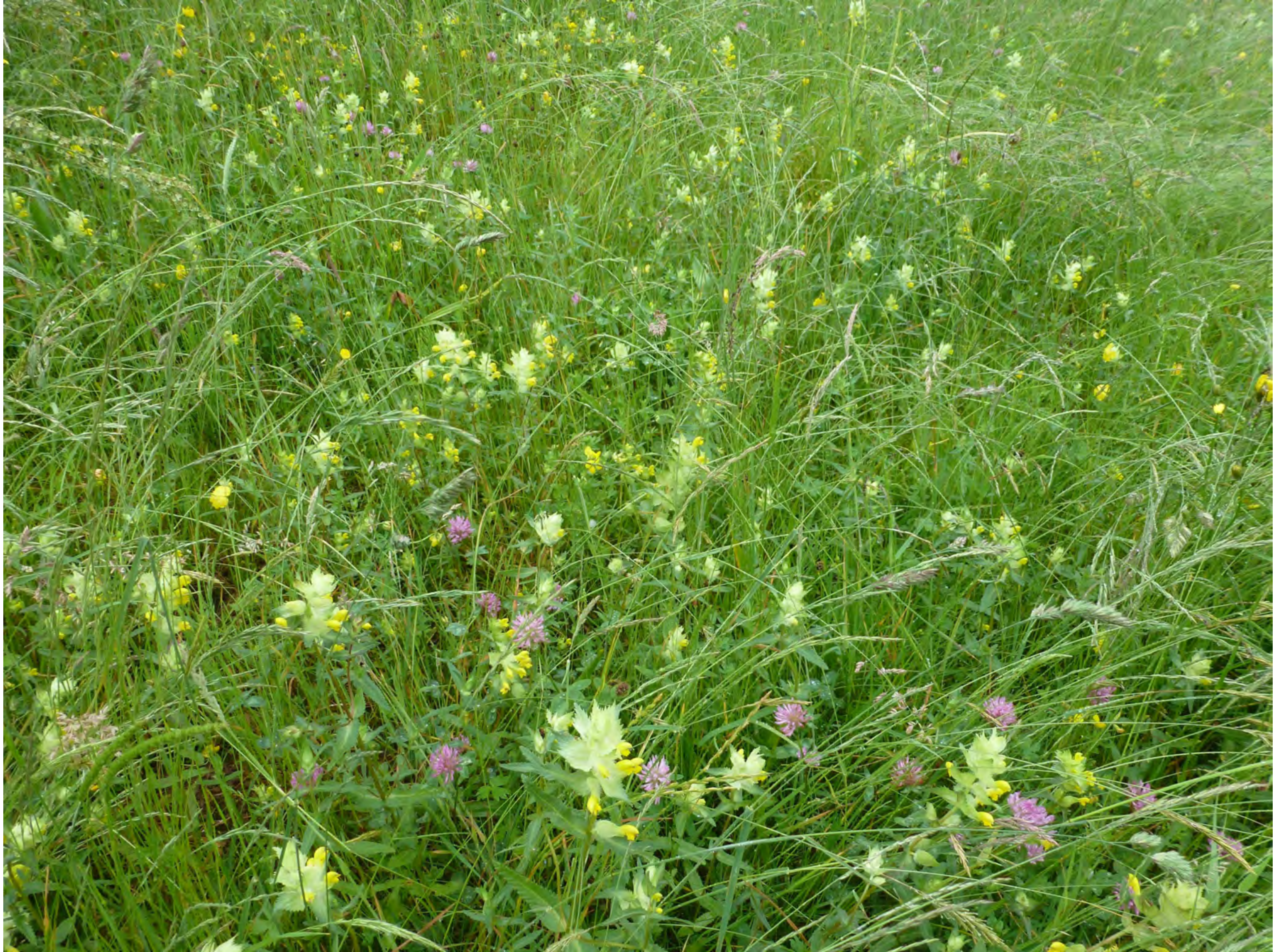

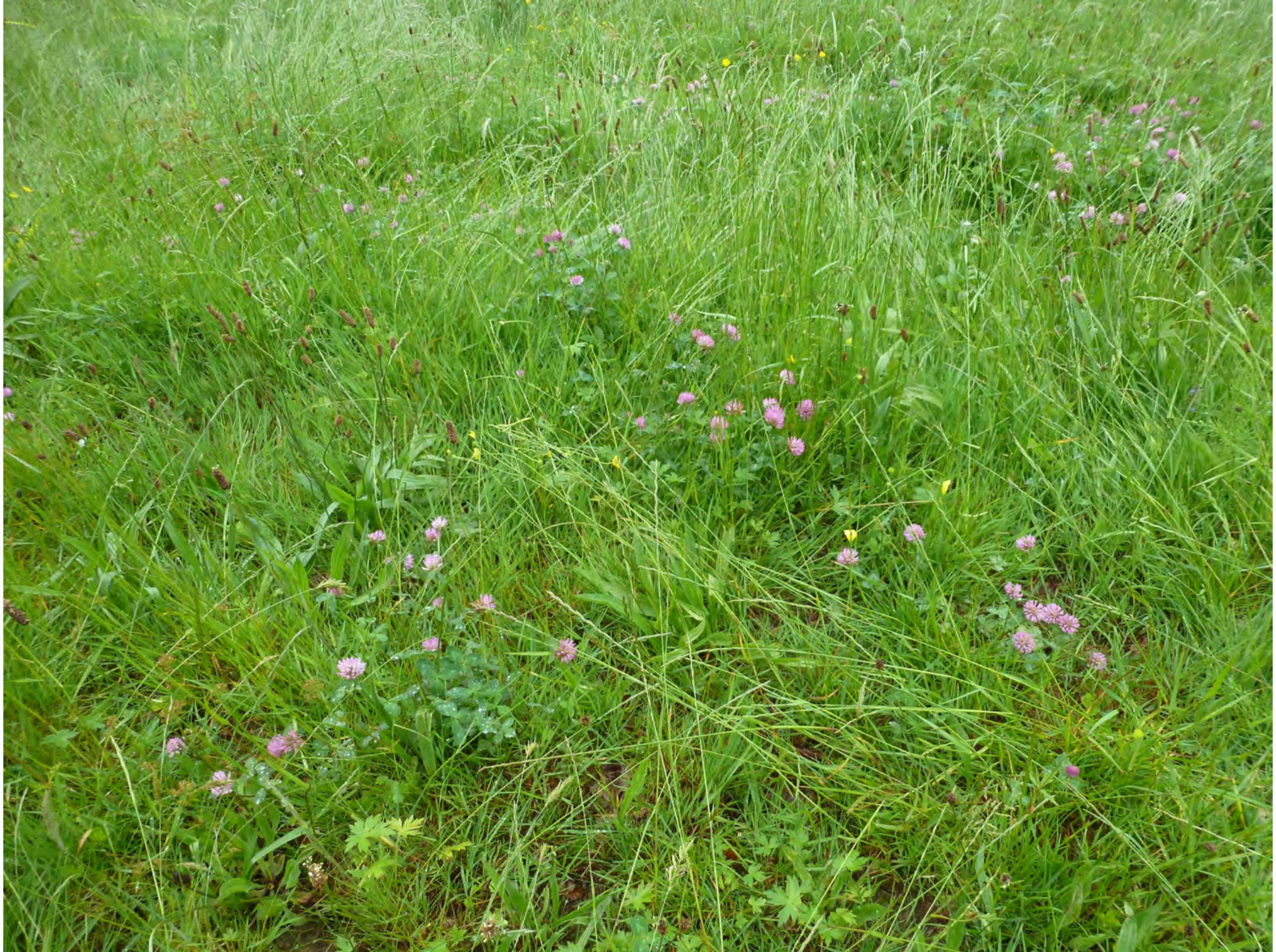

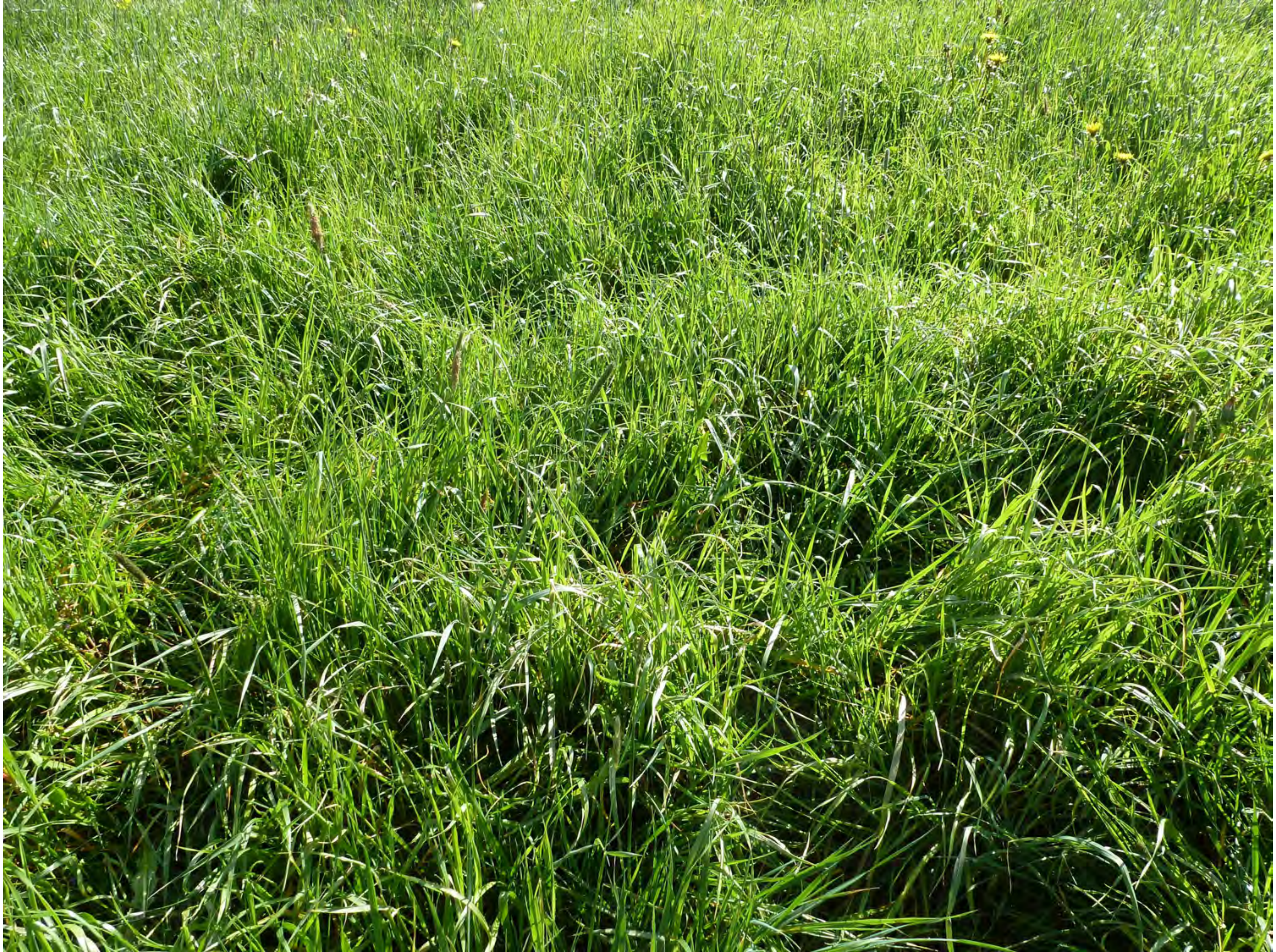

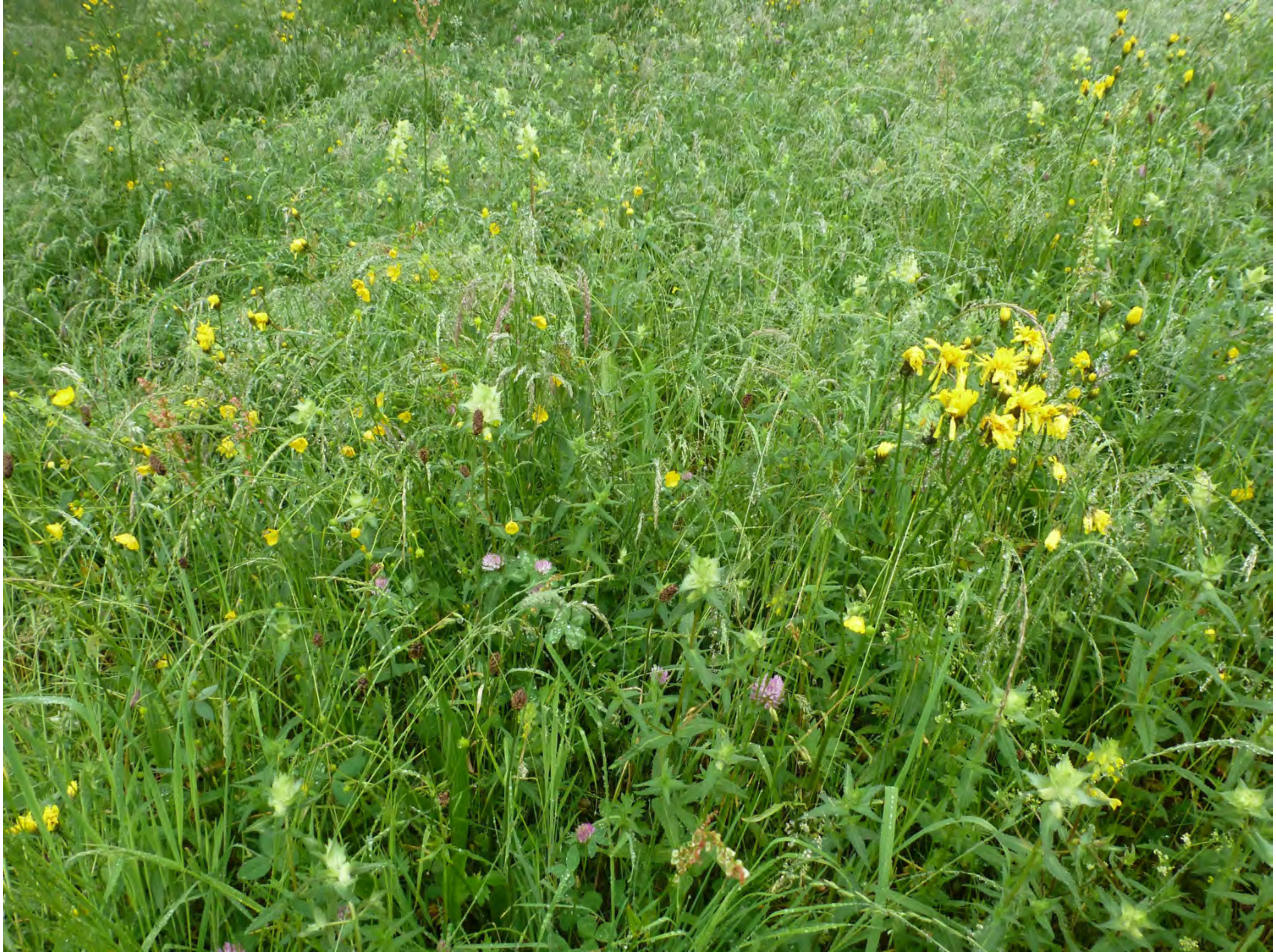

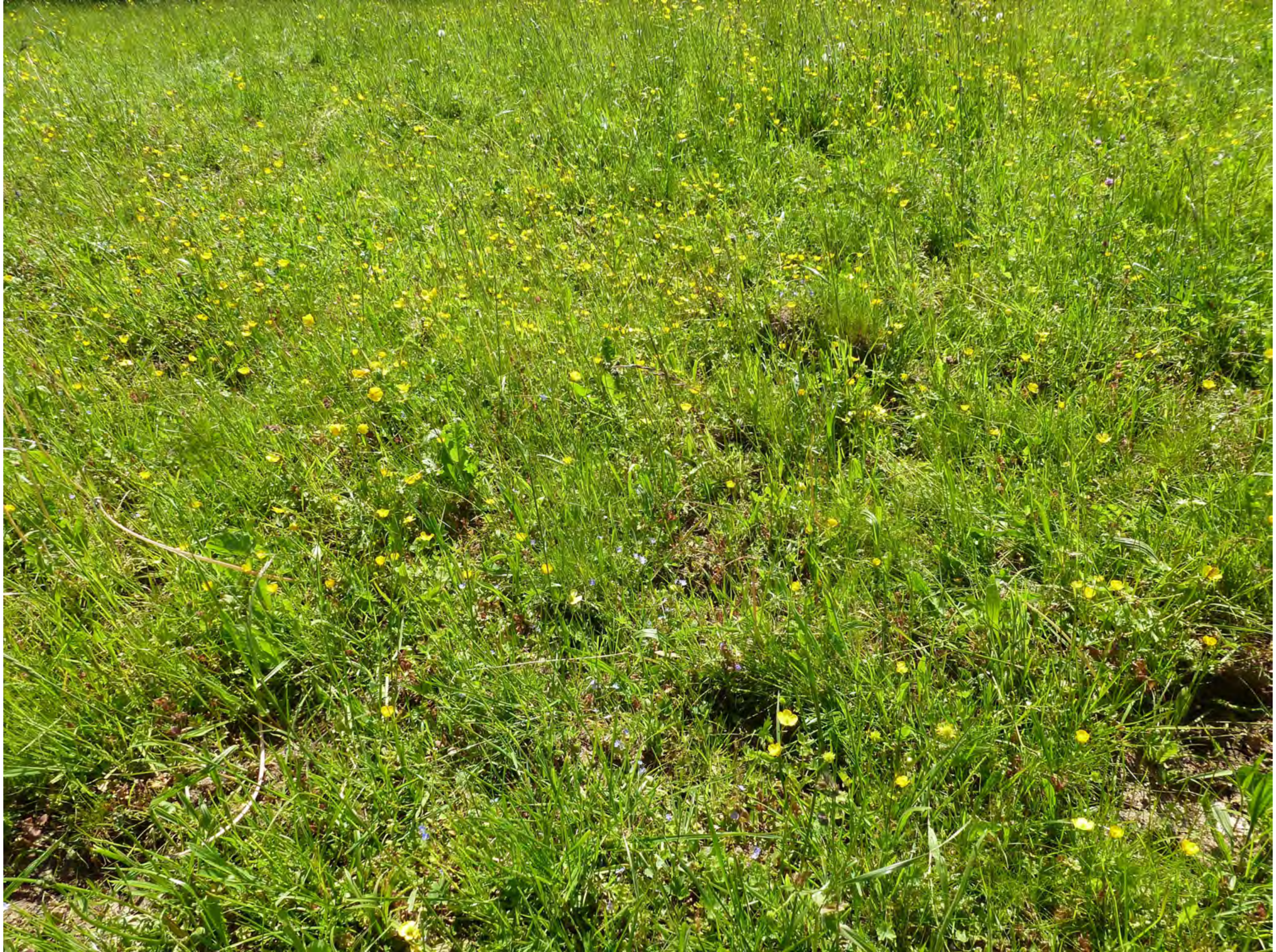

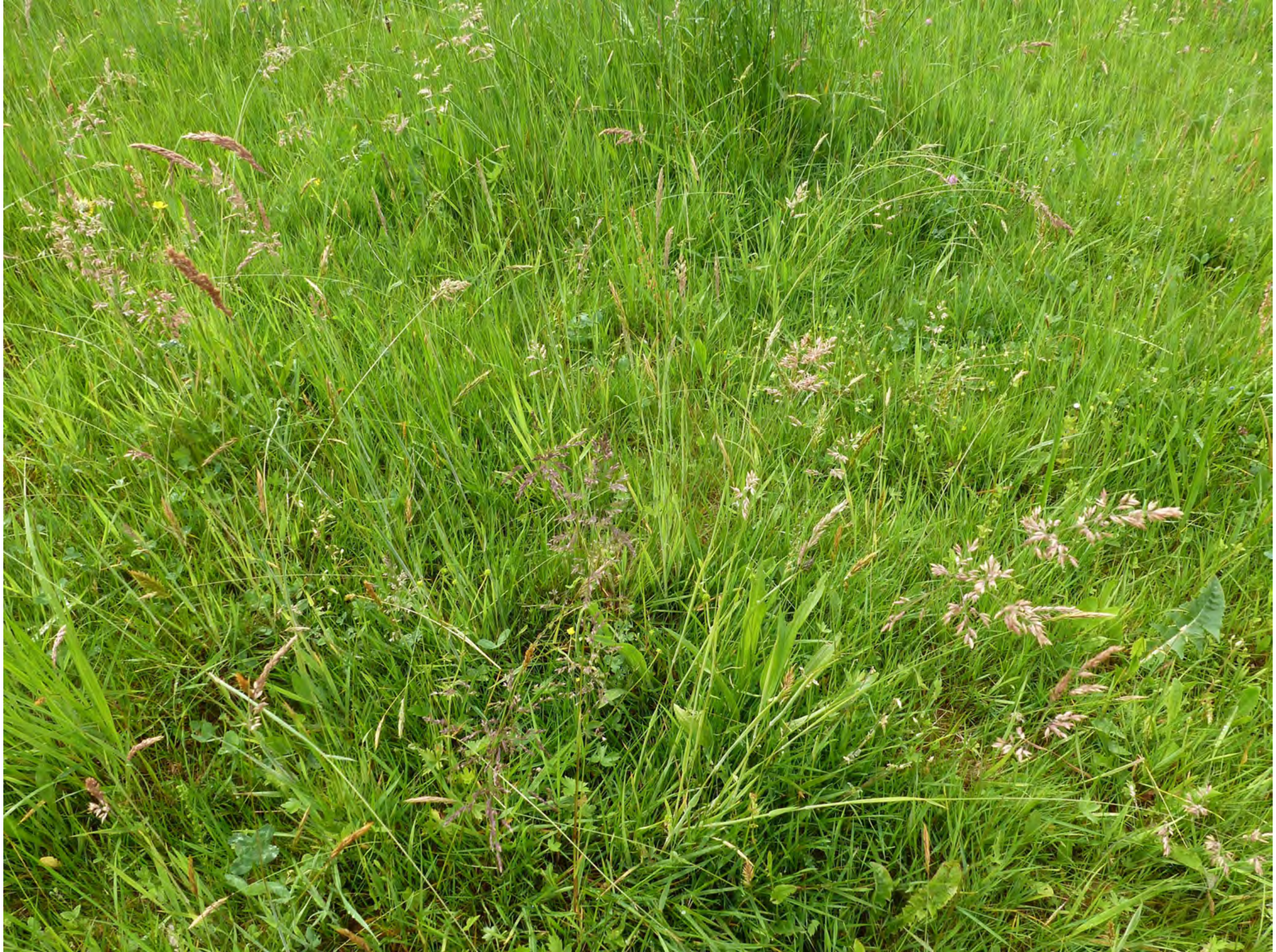

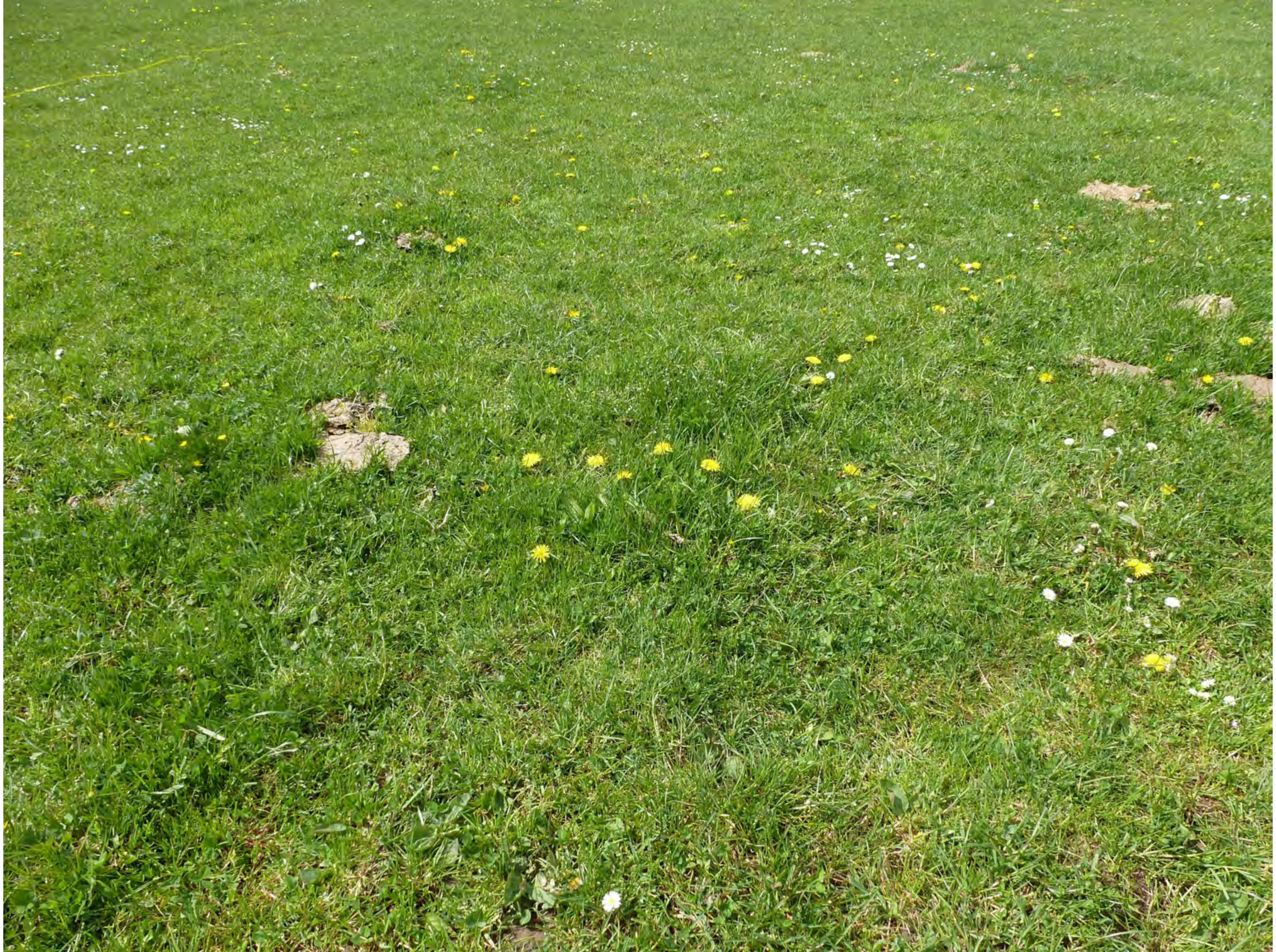

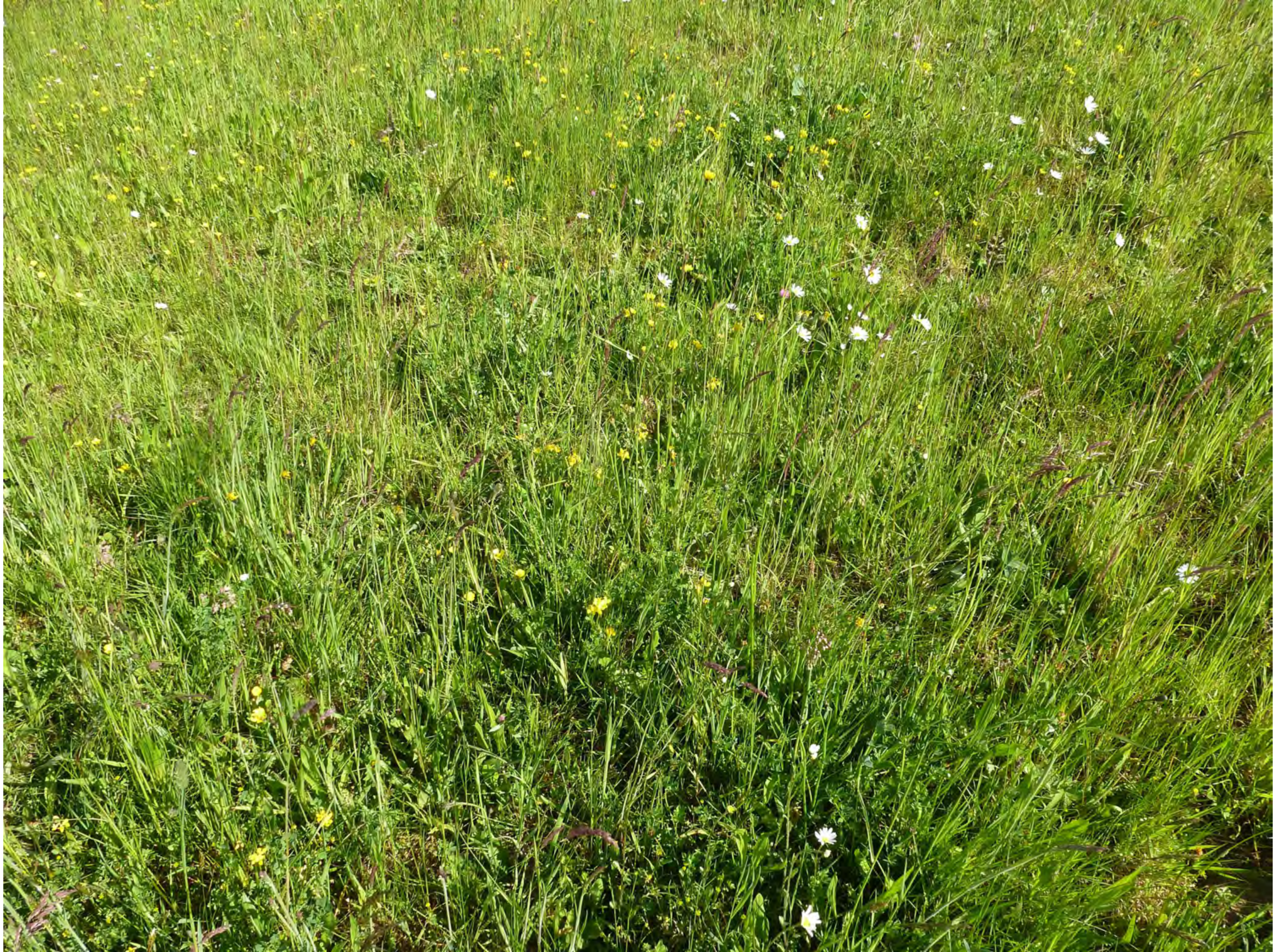

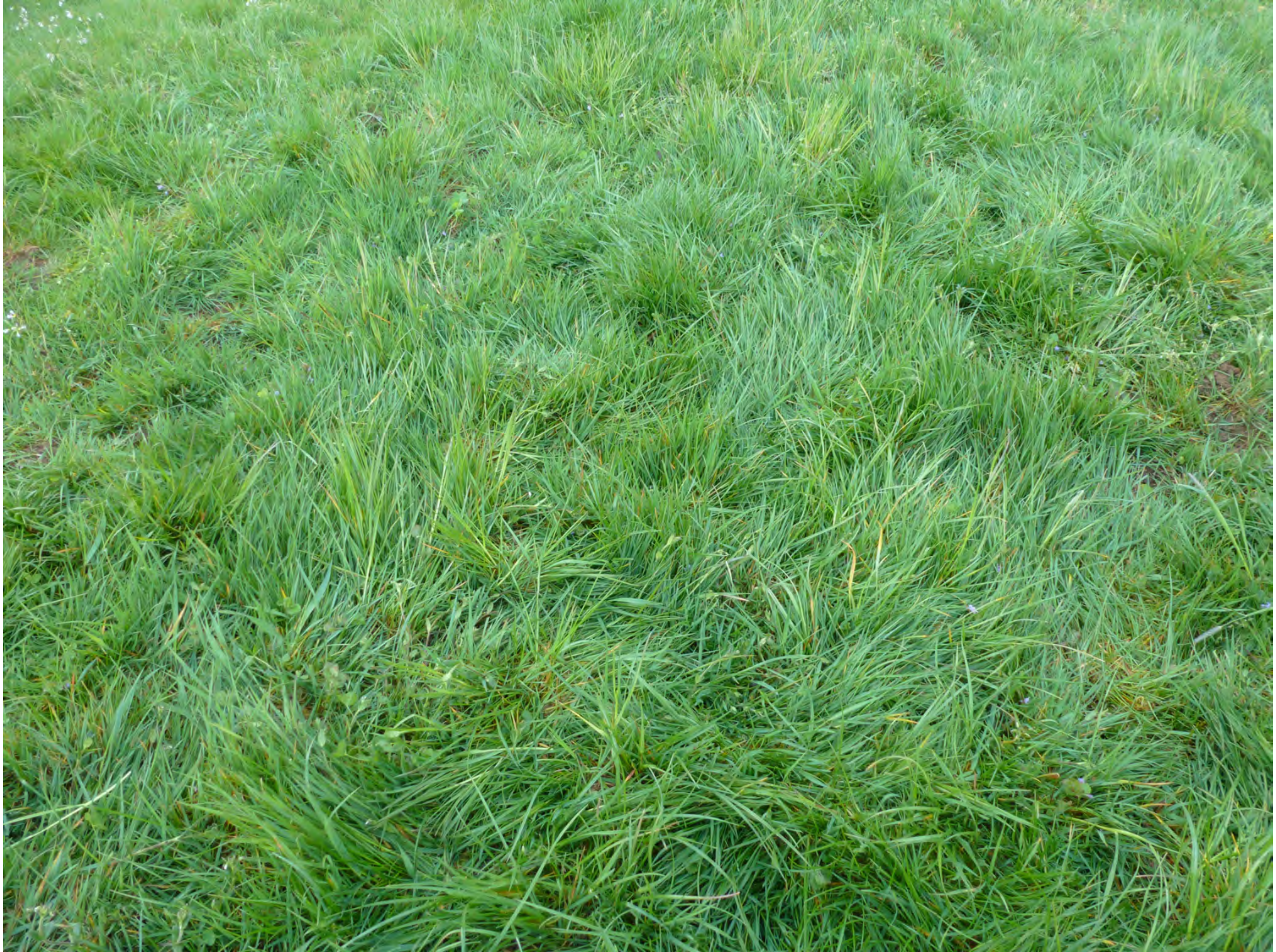

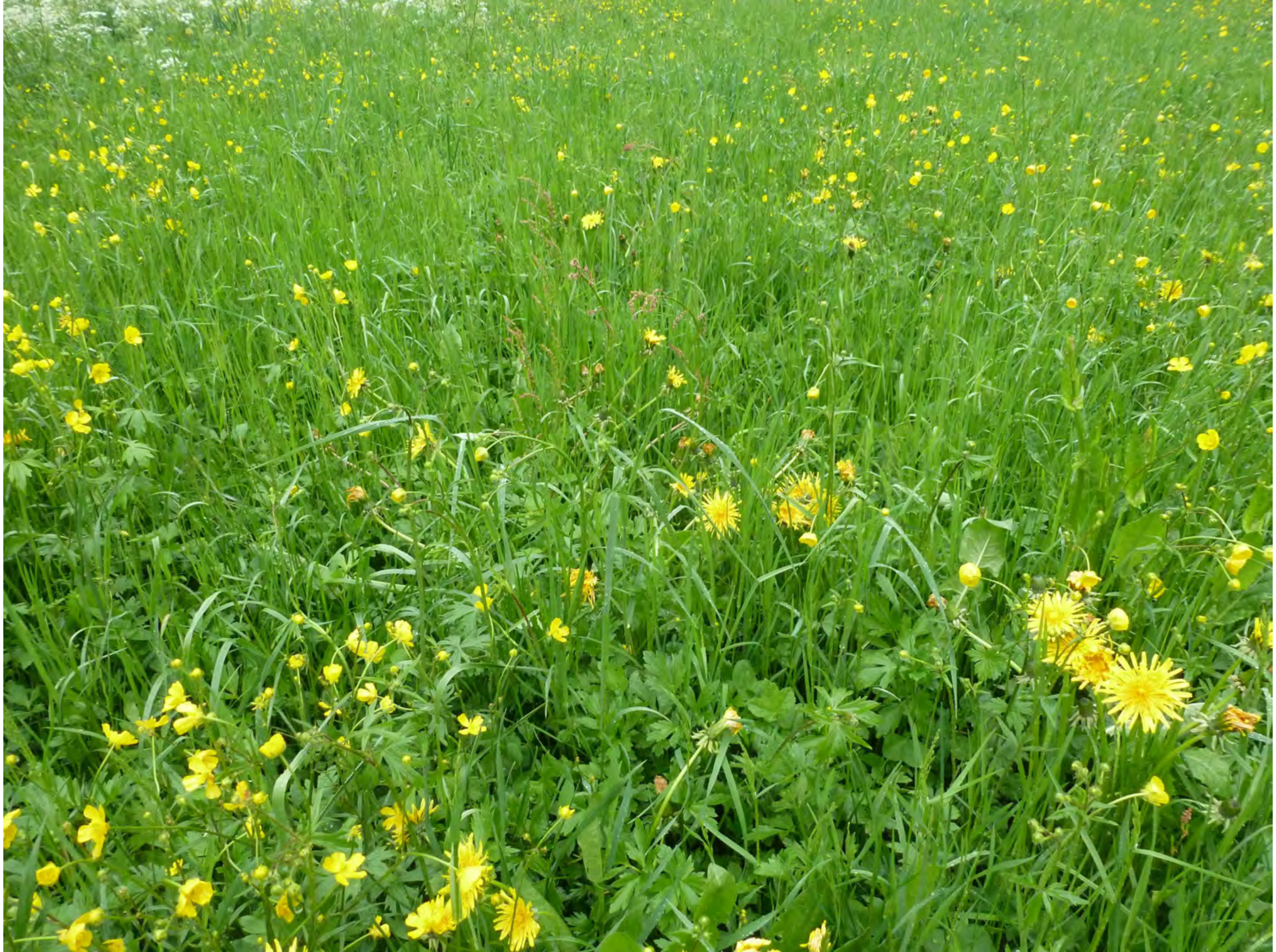

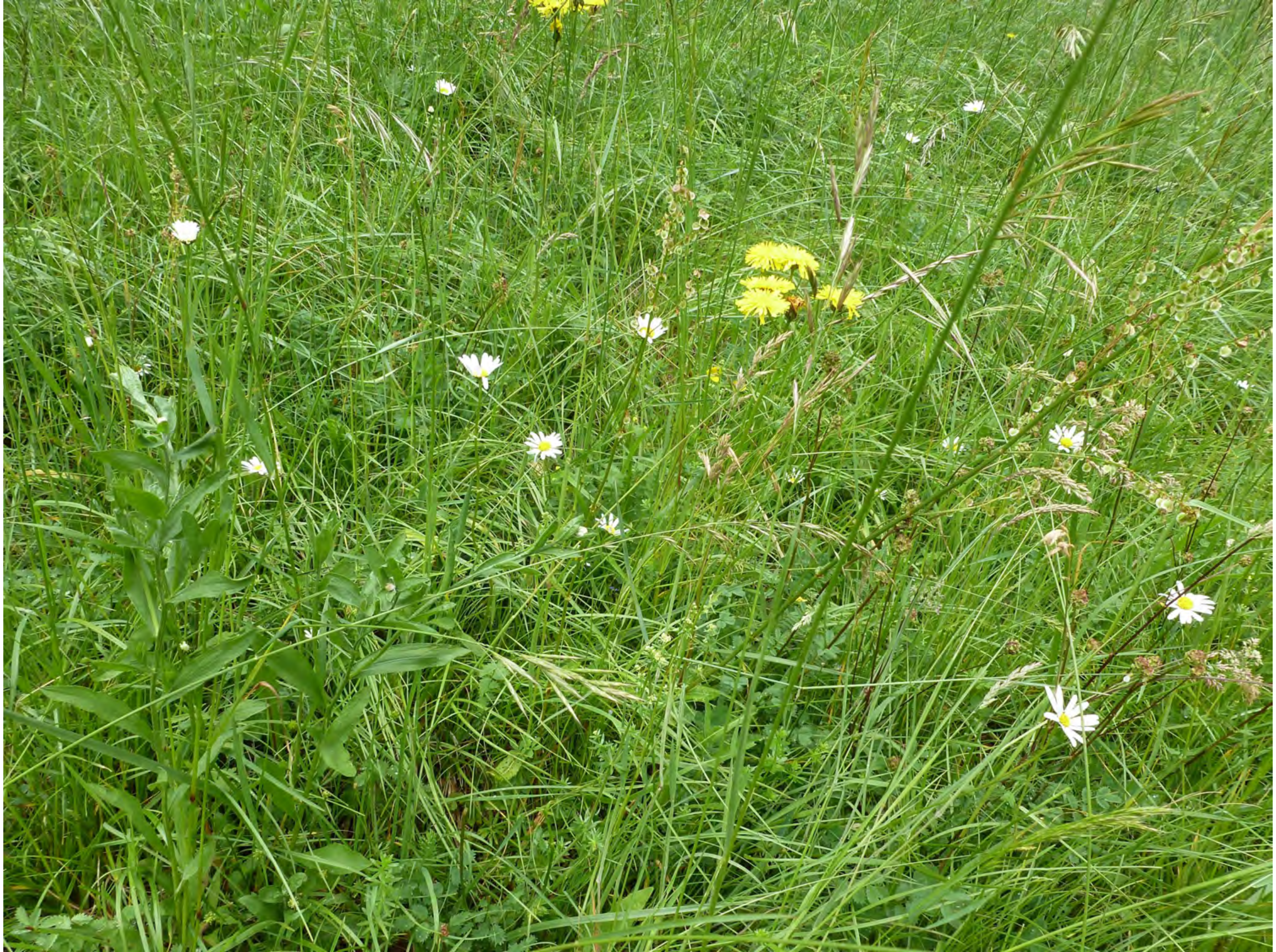

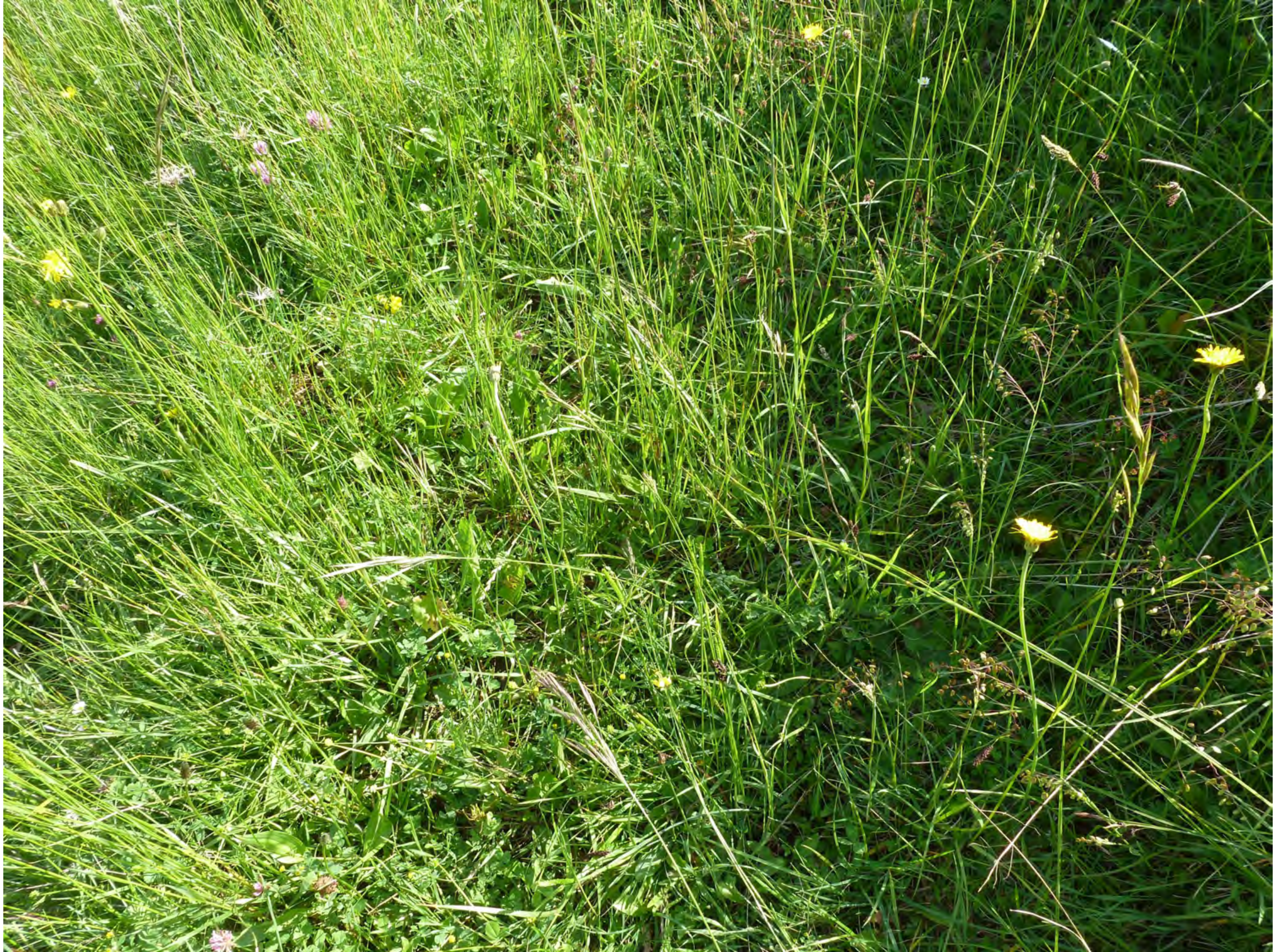

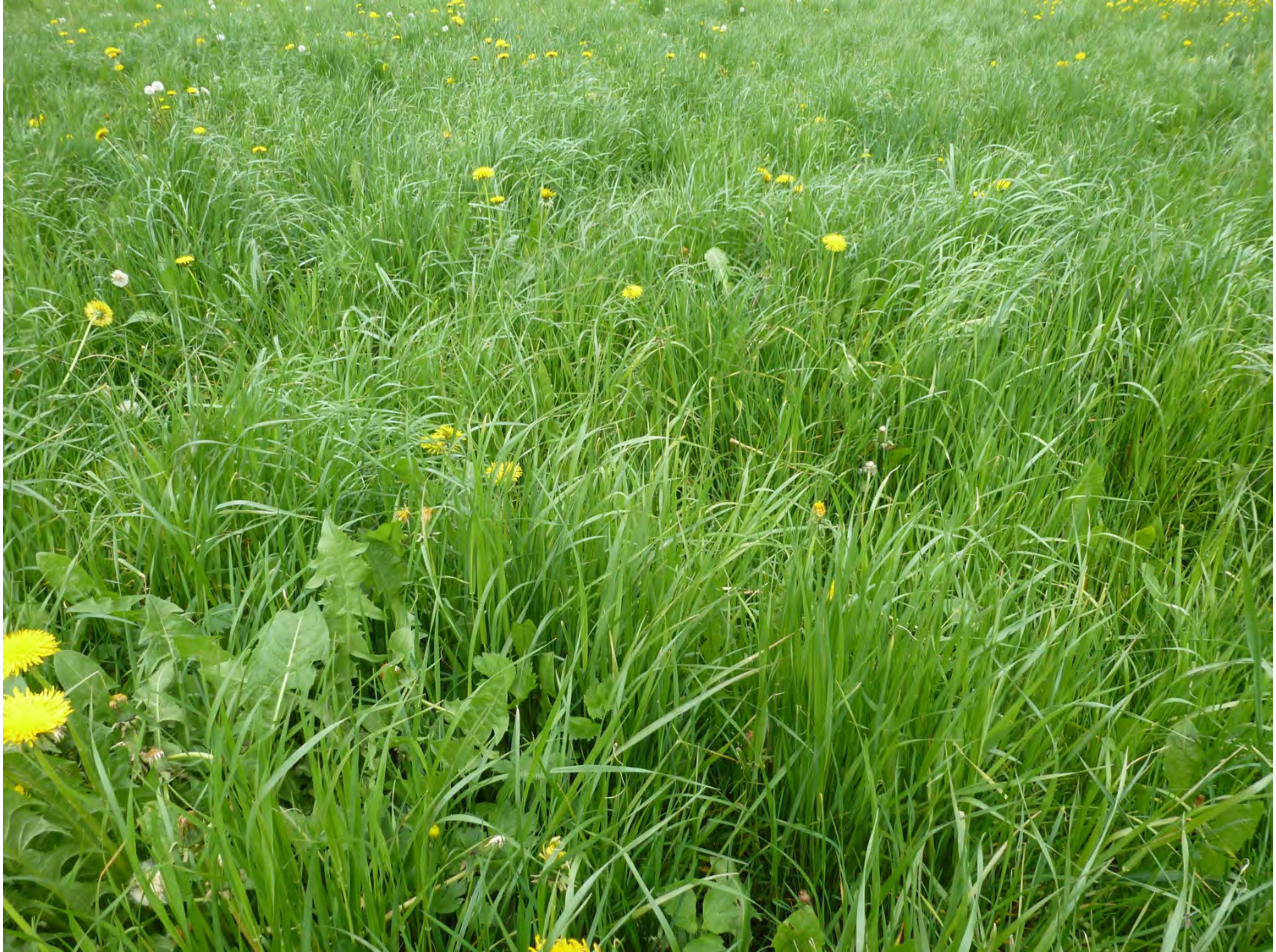

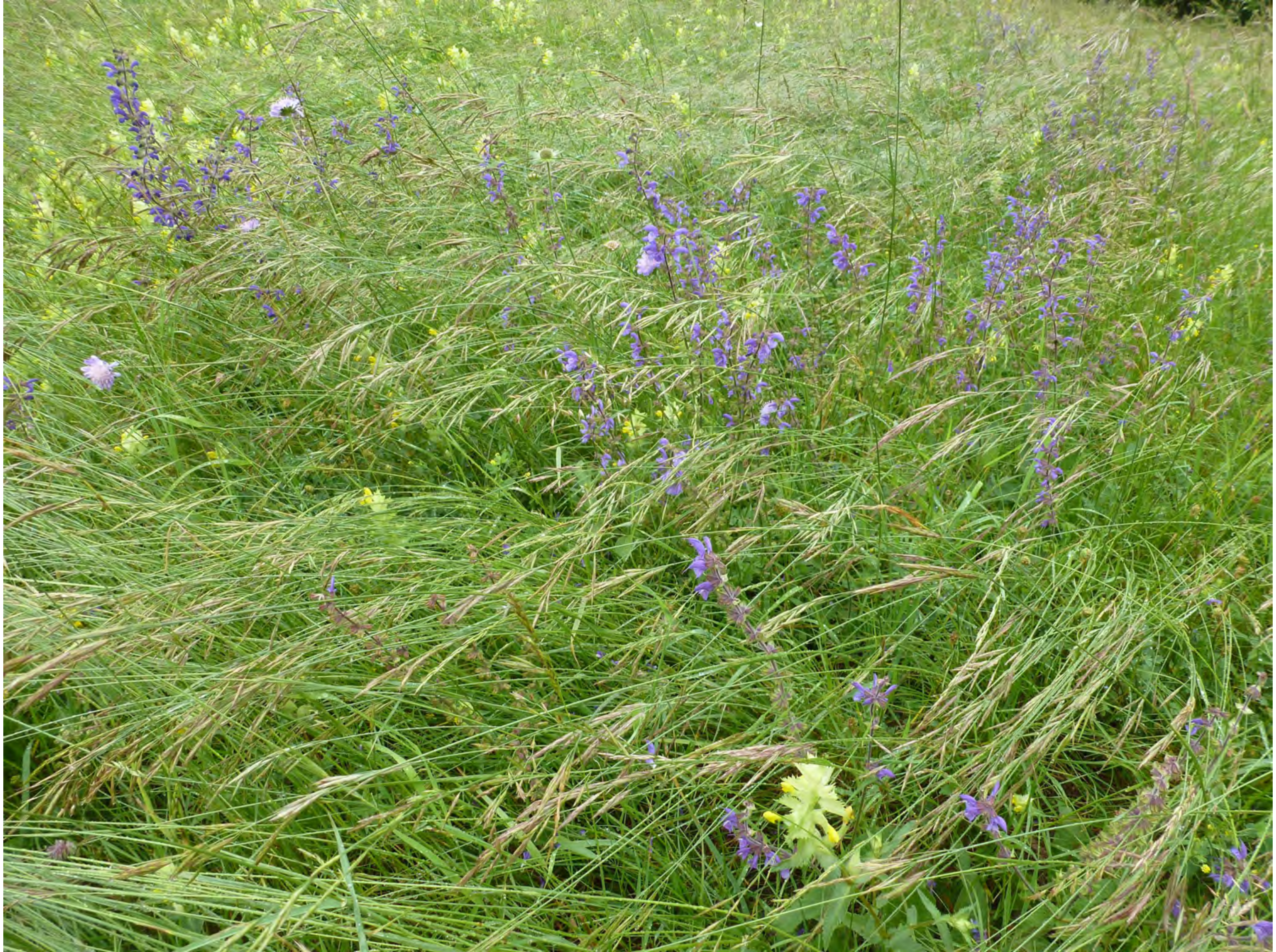

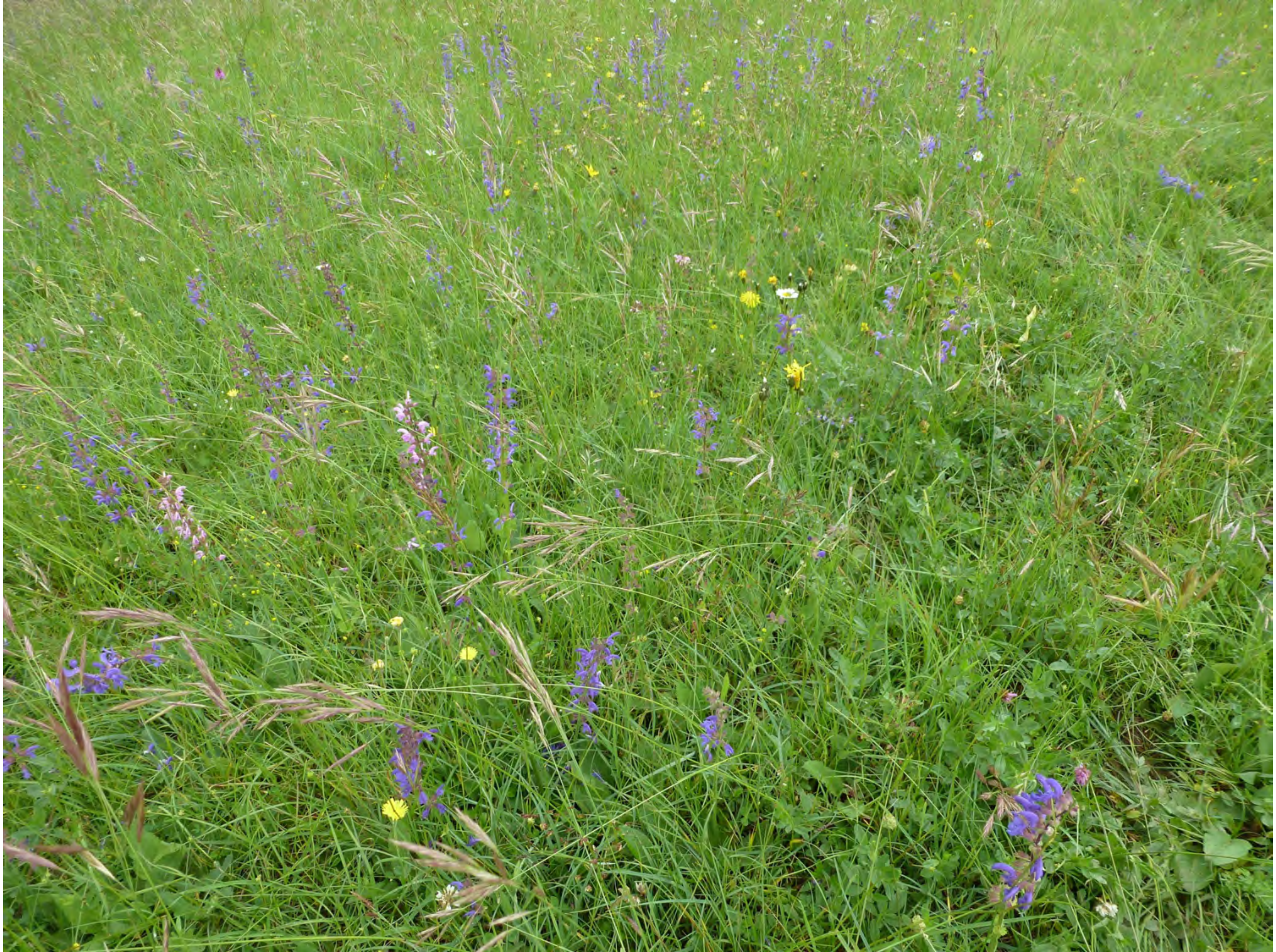

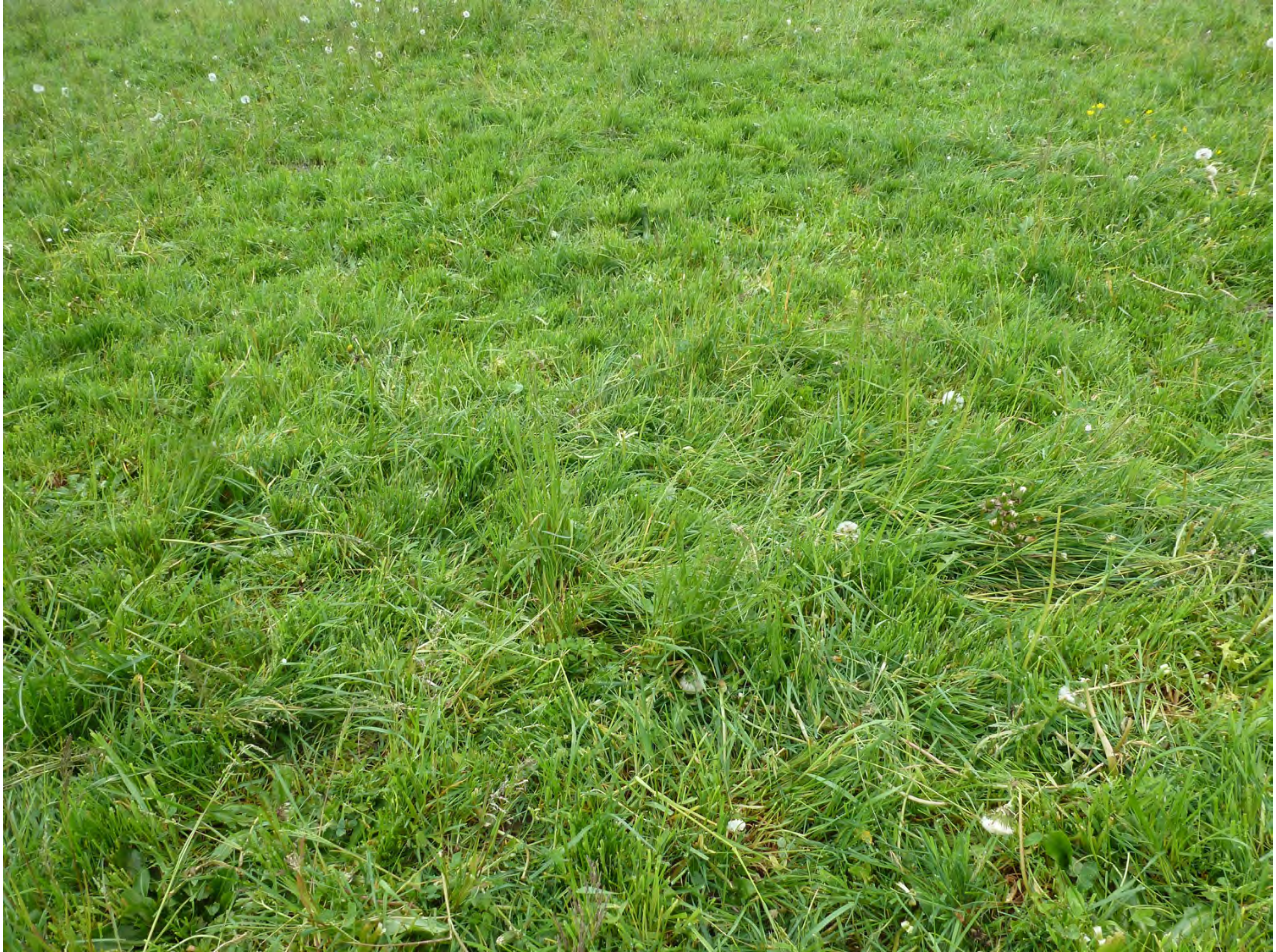

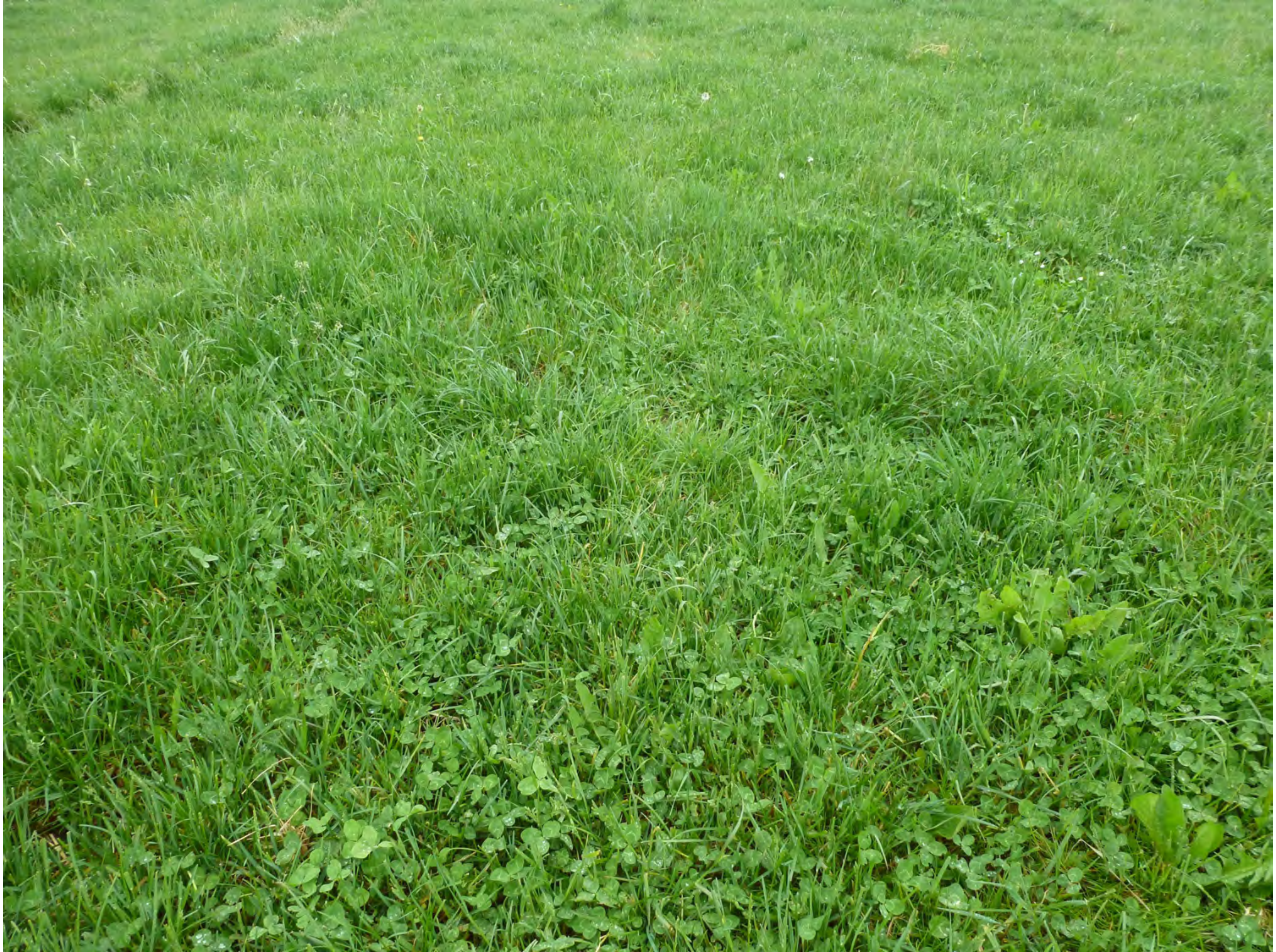

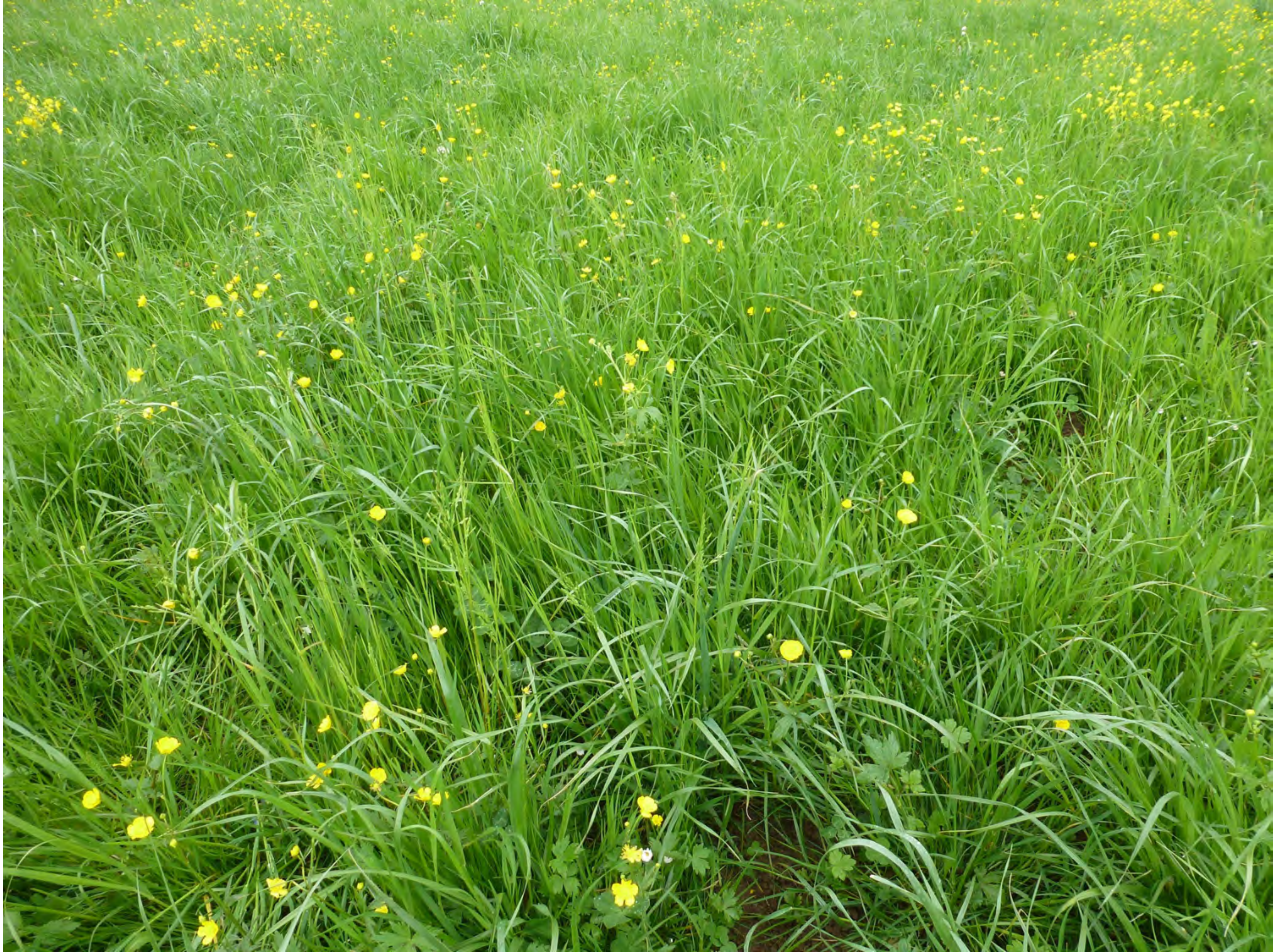

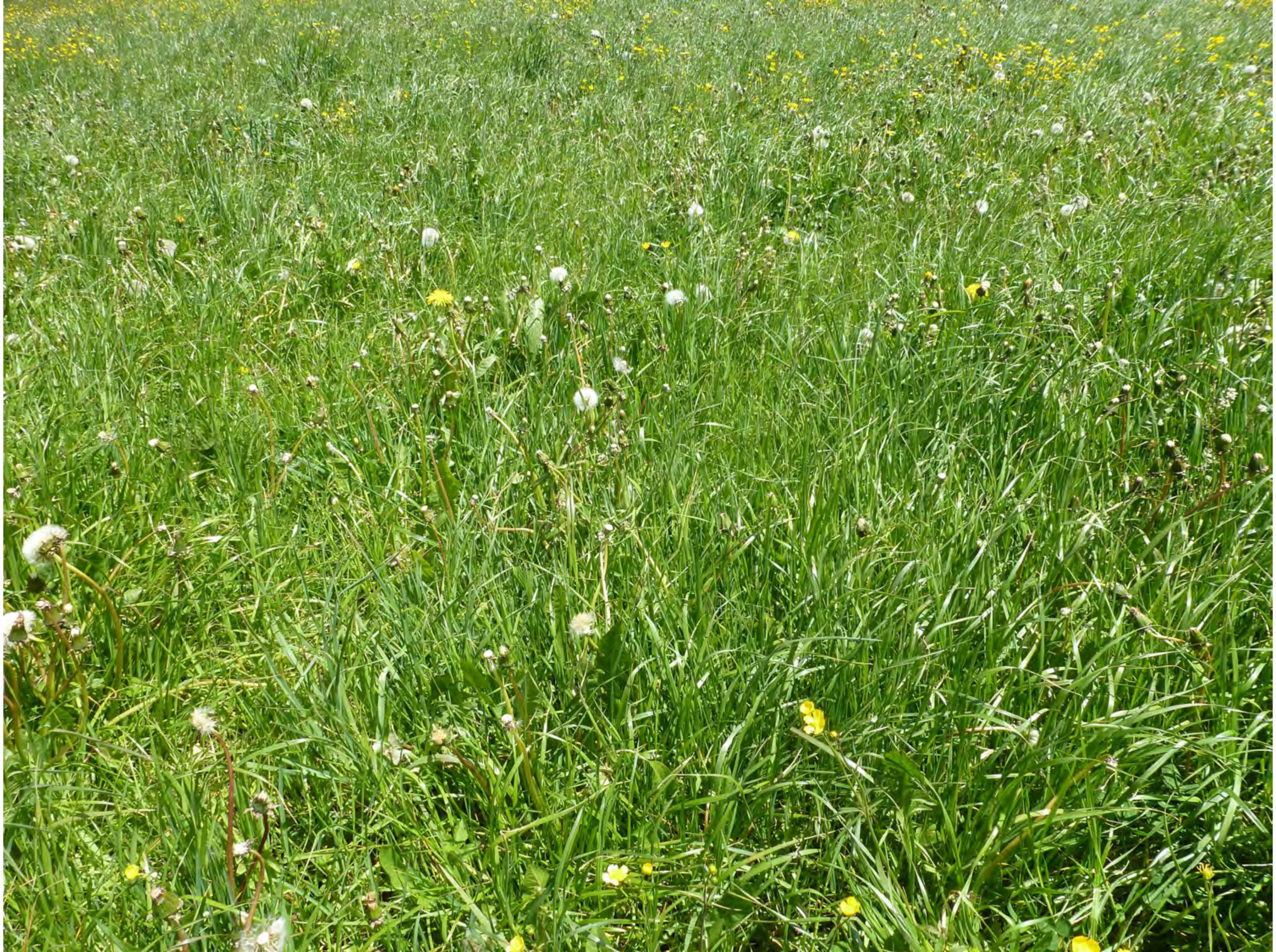

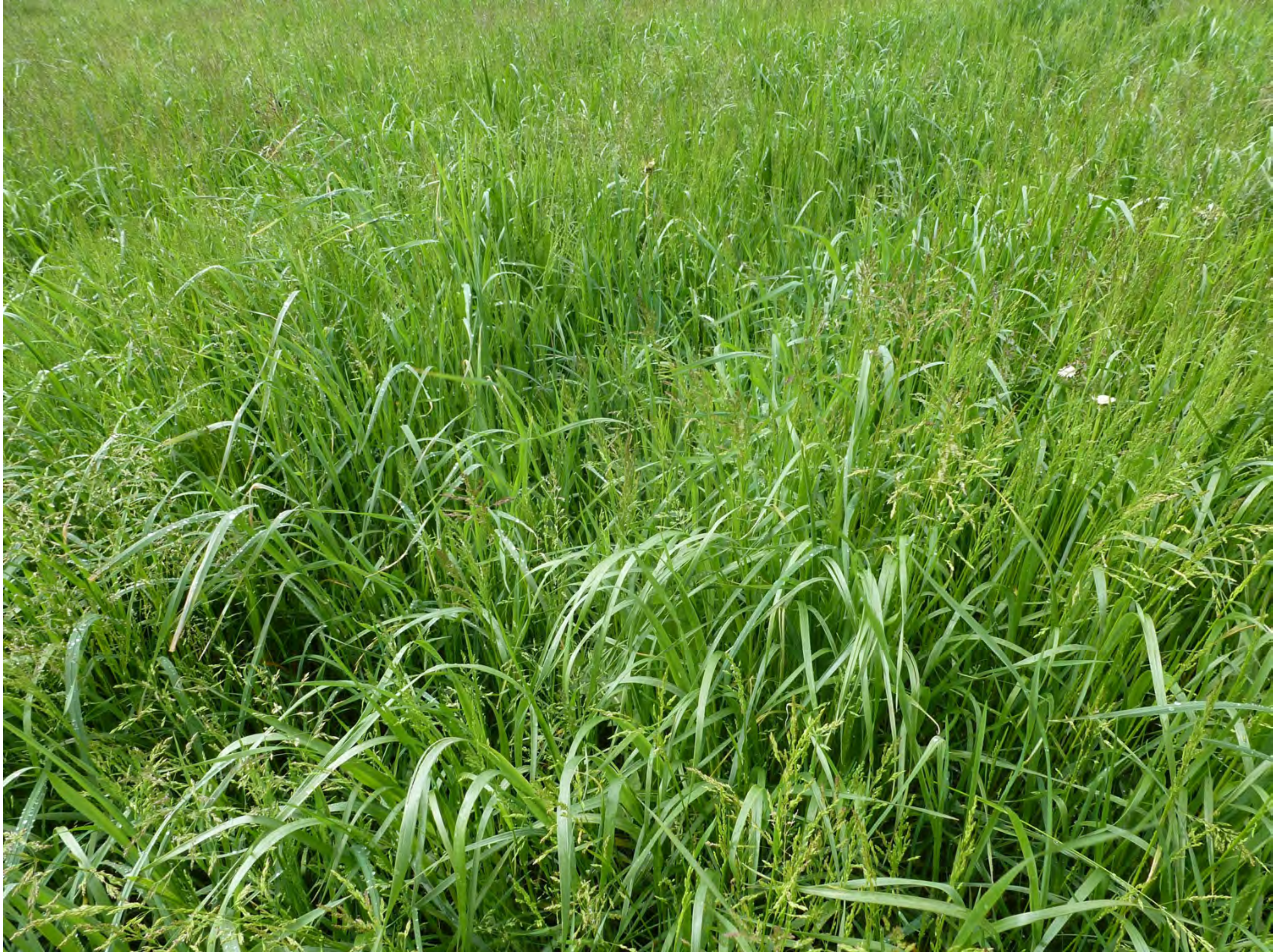

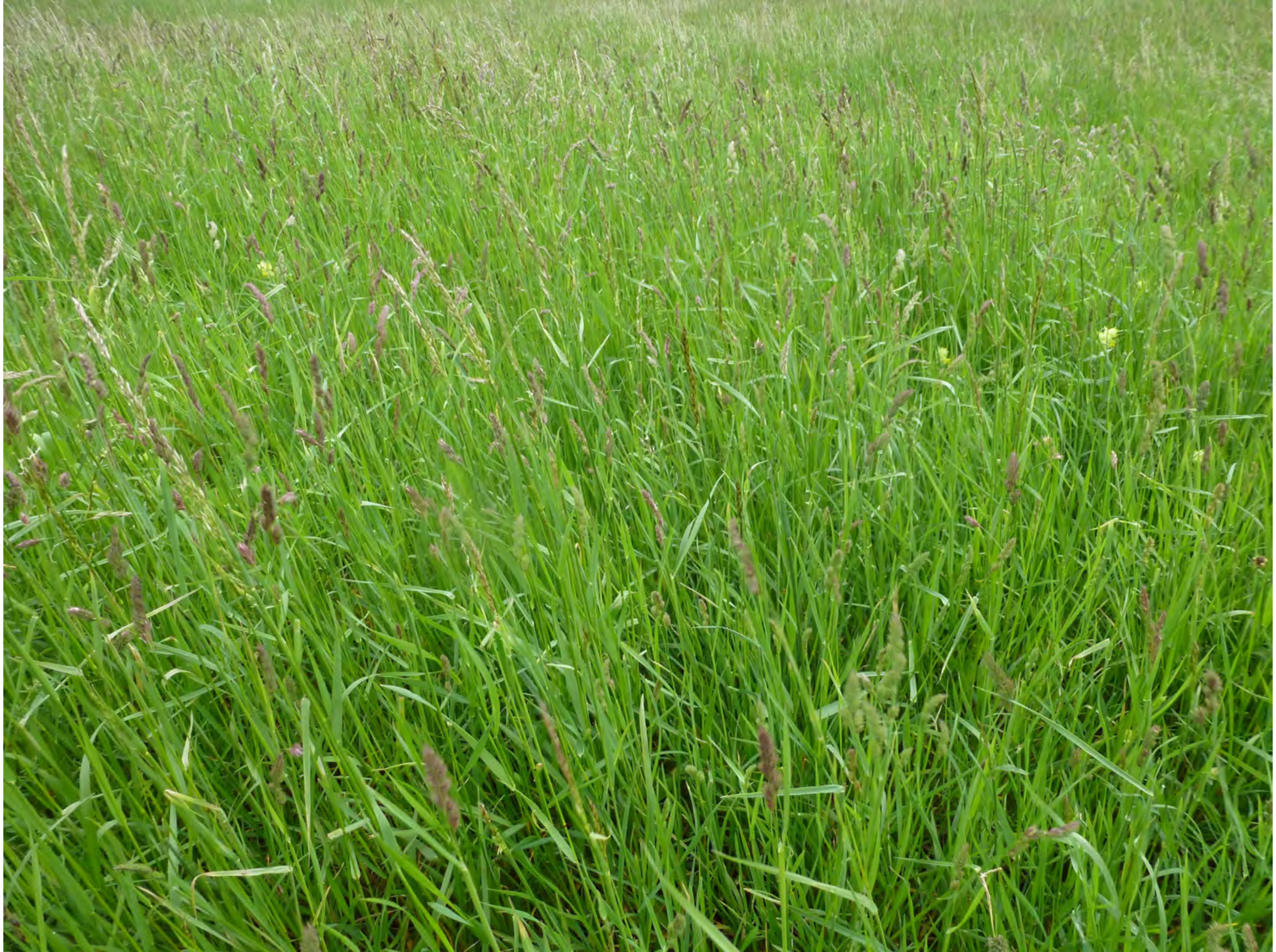

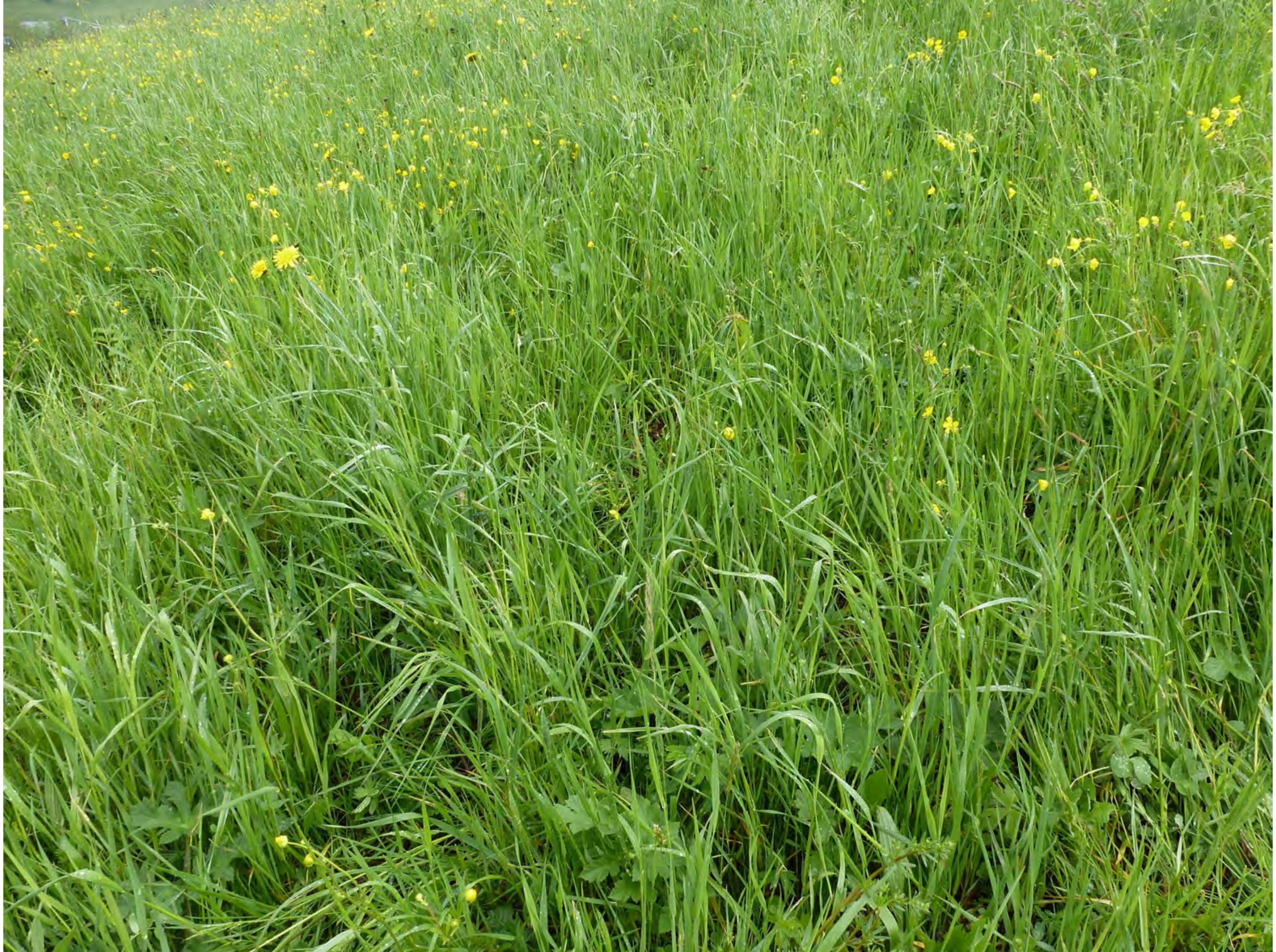

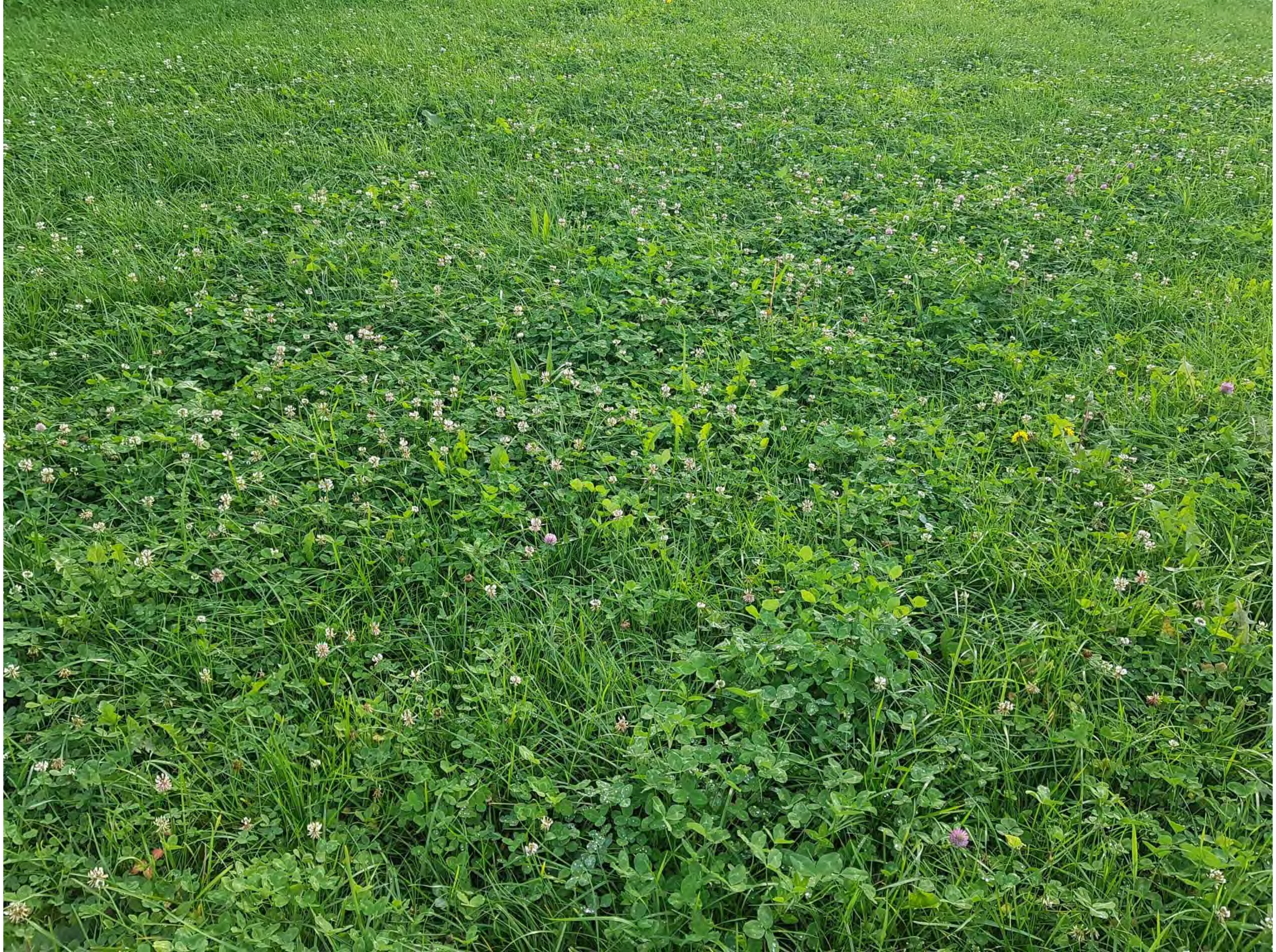

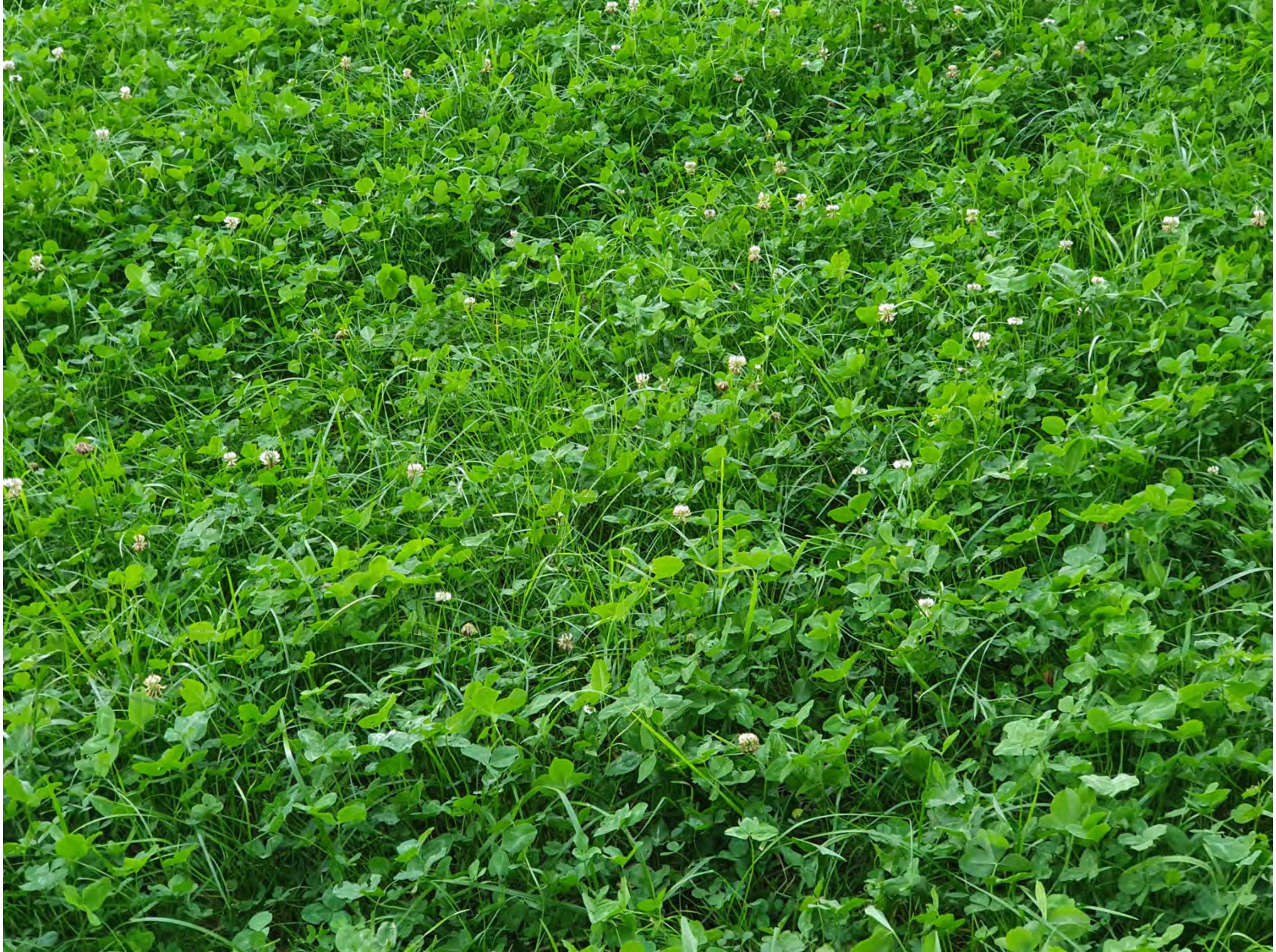
